# OPTN translocates to an ATG9A-positive compartment to regulate innate immune signalling and cytokine secretion

**DOI:** 10.1101/744672

**Authors:** Thomas O’Loughlin, Antonina J Kruppa, Andre LR Ribeiro, James R Edgar, Abdulaziz Ghannam, Andrew M Smith, Folma Buss

## Abstract

Optineurin (OPTN) is a multifunctional protein involved in autophagy, secretion as well as NF-κB and IRF3 signalling and mutations are associated with several human diseases including primary open-angle glaucoma (POAG), amyotrophic lateral sclerosis (ALS), Paget’s disease of bone (PDB) and Crohn’s disease (CD). Here we show that, in response to viral RNA, OPTN translocates to foci in the perinuclear region, where it negatively regulates NF-κB and IRF3 signalling pathways and downstream pro-inflammatory cytokine secretion. These OPTN foci consist of a tight cluster of small membrane vesicles, which are positive for marker proteins of the *trans*-Golgi network/recycling compartment – most notably ATG9A. Disease mutations linked to POAG cause aberrant formation of this compartment in the absence of stimuli, which correlates with the ability of OPTN to inhibit signalling. Using proximity labelling proteomics (BioID), we identify the linear ubiquitin assembly complex (LUBAC), CYLD and TBK1 as part of the OPTN interactome and show that these proteins, along with NEMO, are recruited to this OPTN-positive perinuclear compartment. Together, we propose OPTN might be responsible for dampening the NF-κB and IRF3 signalling responses through the sequestration of LUBAC and other positive regulators of these pathways in this dsRNA-induced compartment leading to altered pro-inflammatory cytokine secretion.

**Summary:** Disease associated OPTN mutations impact on the formation of the perinuclear compartment and result in hypo- or hyper-activation of the immune response, which could drive the development of human diseases such as POAG, ALS and also Paget’s disease of bone.

## Introduction

Pathogen associated molecular patterns (PAMPs) are recognised by pattern recognition receptors (PRRs), such as Toll-like receptors (TLRs), and trigger a range of adaptive and innate immune responses in the host [5]. For example, activation of TLR3 or RIG-I by double-stranded viral RNA activates signalling cascades culminating in the activation of transcription factors including NF-κB and IRF3 and gene expression programs composed of pro-inflammatory cytokines (e.g. IL6) and interferons (IFNs) respectively [6]. Optineurin (OPTN) appears to be a key protein in a range of pathways downstream of TLR3, participating in the innate immune response through the secretion of cytokines, acting as a selective autophagy receptor, and regulating both NF-κB as well as IRF3 signalling [7].

NF-κB signalling centres around the NF-κB transcription factor complex which, under non-stimulated conditions, is inhibited through binding to IκB proteins. In response to stimuli such as TLR3 or RIG-I ligation, the pathway is switched on leading to activation of the IKK complex, composed of two kinase subunits (IKKα and IKKβ) and a regulatory subunit IKKγ (NF-κB essential modulator [NEMO]), which phosphorylates IκB proteins and triggers their subsequent degradation. This degradation releases the NF-κB complex, allowing it to translocate to the nucleus and induce expression of numerous target genes [8]. An additional critical step in this pathway is the linear M1-linked ubiquitination of NEMO and receptor-interacting protein kinase 1 (RIPK1) by the linear ubiquitin assembly complex (LUBAC), which consists of HOIP (RNF31), HOIL-1L (RBCK1) and SHARPIN [9–12]. These linear ubiquitin chains can then function as scaffolds to recruit the IKK complex through the <u>u</u>biquitin <u>b</u>inding in <u>A</u>BIN and <u>N</u>EMO (UBAN) domain of the IKK subunit NEMO [13–15]. OPTN is highly similar to NEMO with around 52% sequence homology and sharing its linear ubiquitin-binding UBAN domain [14,16]. However, unlike NEMO, OPTN cannot bind to IKKα or IKKβ and therefore cannot rescue NF-κB activity in NEMO-deficient cells [16]. Instead, OPTN appears to antagonise NEMO function by competitively binding to ubiquitinated RIPK1 and can thereby inhibit TNFα-induced NF-κB activation [17]. In addition, OPTN interacts with CYLD, a deubiquitinase (DUB) for linear and K63 ubiquitin chains, which is able to negatively regulate NF-κB signalling via the deubiquitination of a range of NF-κB signalling proteins including NEMO and RIPK1 [18,19].

Alternatively, OPTN can bind to the IKK-related kinase TBK1 or the E3 ligase TRAF3 to regulate IRF3 activity [20,21]. A complex composed of TBK1 and IKKε is activated via TRAF3 downstream of PRRs, such as TLR3 or RIG-I. Once active, the TBK1/IKKε complex can phosphorylate its substrate, IRF3, which subsequently dimerises and translocates to the nucleus to induce expression of target genes such as Type I IFNs (IFNs). Through its interactions with both TBK1 and TRAF3, OPTN appears to attenuate IFN-β production [20].

An increasing number of perturbations in OPTN gene function have been linked to diseases including primary open-angle glaucoma (POAG), amyotrophic lateral sclerosis (ALS), Paget’s disease of bone (PDB) and Crohn’s disease (CD) [1–4]. A common feature of OPTN’s role in these diseases appears to be aberrant NF-κB signalling or cytokine secretion profiles. Many ALS mutants show a loss of OPTN-mediated NF-κB suppression [22], deficiencies in OPTN expression increase NF-κB activity and susceptibility to PDB [23], and a subset of CD patients with reduced OPTN expression display impaired secretion of TNF-α, IL6 and IFN-γ [3].

In this study, we address the role of OPTN in innate immune signalling and cytokine secretion and the mechanism by which perturbation of OPTN function in these processes may contribute to human inflammatory disease. We use a retinal pigment epithelial (RPE) cell model, which is relevant to the role of OPTN in the pathogenesis of POAG, and show these cells respond to TLR3 and RIG-I ligands, leading to upregulation of OPTN and its translocation to perinuclear foci. Our ultrastructural analysis of these foci by correlative light and electron microscopy reveals that this compartment consists of a tight cluster of small vesicles, which contain the autophagy protein ATG9A. This multispanning membrane protein is present at the Golgi complex and in clusters of small 30-40 nm vesicles, which are often found in close proximity to autophagosomes, but do not appear to be incorporated into the growing phagophore [24,25]. We demonstrate that wild-type or mutant variants of OPTN show variable recruitment to this vesicle cluster, which correlates with its ability to negatively regulate NF-κB and IRF3 signalling and therefore cytokine secretion. Using proximity-dependent proteomics (BioID) to characterise this compartment, we identify novel OPTN interacting proteins including IFT74, IFI35, a phosphoinositide phosphatase complex (MTMR6-MTMR9) and LUBAC, with the latter being recruited to OPTN-positive foci upon TLR3 ligation. Our data suggests that OPTN can inhibit the innate immune response through sequestering key components of NF-κB and IRF3 signalling pathways in a novel perinuclear compartment. Disease-associated OPTN mutations impact on the formation of the perinuclear compartment and result in hypo- or hyper-activation of the immune response, which could potentially drive the development of a number of human diseases.

## Results

### Retinal pigment epithelial (RPE) cells exhibit a robust response to double-stranded RNA

RPE cells perform a number of support functions in the inner eye including the secretion of signalling molecules and the maintenance of the immune privileged environment through communication with the immune system [26]. Previous reports have demonstrated that RPE cells express a number of TLRs including the viral RNA receptor, TLR3 [27]. OPTN mutations have been implicated in POAG [1,28], making the RPE cell line a relevant tool to study OPTN function in this disease. Furthermore, the proposed roles for OPTN in anti-viral immunity and TLR3 signalling led us to investigate the utility of this cell line as a tractable human model for OPTN function in these pathways.

RPE cells were stimulated with a range of PAMPs and the immune response determined through the quantification of CXCL8 secretion. Of all the PAMPs tested, only poly(I:C) and pppRNA induced significant CXCL8 secretion consistent with the expression and activation of TLR3 and RIG-I in RPE cells (Fig. S1A). Lipopolysaccharide (LPS), Pam3CSK4 and 2’,3’-cGAMP (cGAMP) were unable to elicit the release of CXCL8 from RPE cells illustrating a lack of activation downstream of TLR4, TLR2 and STING. To determine the complete secretory response of RPE cells downstream of poly(I:C) stimulation, we analysed conditioned medium from unstimulated or poly(I:C)-stimulated RPE cells using quantitative SILAC mass spectrometry. These experiments identified 380 proteins in the conditioned medium with 26 showing significant (p<0.05) upregulation (Fig. S1B; supplementary Table 1). Among the upregulated proteins were well-known pro-inflammatory cytokines such as CXCL8 and, to a lesser extent, IL6 (Fig. S1B). We validated this data by ELISA and found poly(I:C) stimulation resulted in the induction of both CXCL8 and IL6 protein secretion (Fig. S1C[i-ii]). To assess the contribution of NF-κB signalling in regulation of cytokine secretion, we generated an RPE cell line expressing a NF-κB luciferase reporter. We found that stimulating these cells with poly(I:C) induced NF-κB B promoter activity and a similar elevation in phospho-NF-κB p65 was observed using immunoblot analysis (Fig. S1C[iii]-D). Although no IFNs were detected in the proteomics datasets, we predicted that IRF3 signalling would also be active downstream of TLR3 [29]. Indeed, upon poly(I:C) stimulation we observed a rapid phosphorylation of IRF3, an elevation in IFN-β mRNA levels, and could detect IFN in the supernatant 2 hours post stimulation (Fig. S1C[iv-v]-D).

### OPTN translocates to a novel perinuclear compartment in response to double-stranded RNA

Transient overexpression of OPTN triggers the formation of Golgi-proximal foci [4,30–36], which have been postulated to be aggresomes [30] or organelles participating in post-Golgi membrane trafficking and the maintenance of Golgi integrity [32,34,35]. We observed that stably expressed GFP-OPTN was predominantly cytosolic in resting RPE cells but, strikingly, translocated to perinuclear foci after stimulation with both poly(I:C) or pppRNA (Fig. 1A-B), but not with other PAMPs, such as LPS, cGAMP or Pam3CSK4 (Fig. S2A). Similarly, endogenous OPTN was recruited from a diffuse cytosolic pool to bright foci in the perinuclear region in poly(I:C)-stimulated RPE cells (Fig. 1C). We assessed the rate of formation of this compartment and discovered that the foci began to form beyond 2 hours post-stimulation before peaking at approximately 24 hours (Fig. 1D). OPTN gene expression is regulated through NF-κB signalling [37] and increases upon TLR3 activation by poly(I:C) or viral infection [20]. Similarly, we observed that expression of OPTN is markedly upregulated in response to poly(I:C) stimulation in RPE cells with kinetics similar to foci formation (Fig. 1E). This data suggests that elevated OPTN expression triggers its accumulation into perinuclear foci.

**Figure 1.**
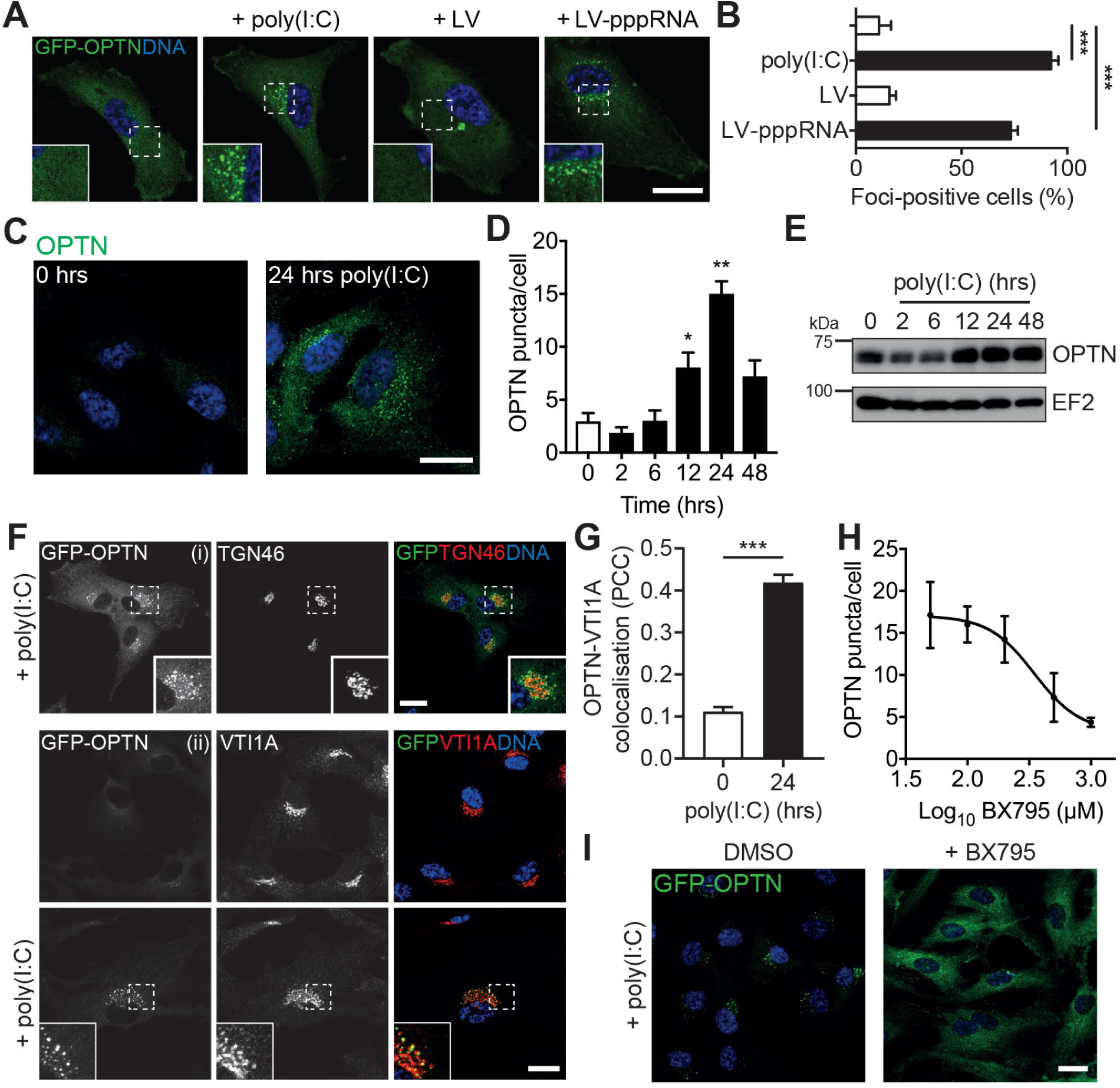
OPTN is recruited to a novel compartment in response to single- and double-stranded viral RNA. (A) Confocal microscope images of RPE cells stably expressing GFP-OPTN (green) and treated with vehicle, poly(I:C), Lyovec (LV) or pppRNA transfected with LV for 24 hours. Cells were stained with Hoechst to label DNA (blue). Scale bar, 20 µm. (B) Percentage of GFP-OPTN cells containing foci after treatment. >100 cells were manually counted per condition from *≥*10 randomly selected fields of view. Bars represent the mean of n=3 independent experiments (except for LV n=2) ±SEM. Statistical significance was determined by repeated measures ANOVA and Bonferroni post-hoc test. *** = p<0.001. (C) Confocal microscope images of RPE cells stimulated with poly(I:C) for 0 and 24 hours. Cells were immunostained with an anti-OPTN antibody (green) and DNA was visualised with Hoechst (blue). Scale bar, 20 µm. (D) Foci count/GFP-OPTN cell after treatment with poly(I:C) for the indicated times. Bars represent mean of n=3 independent experiments ±SEM. Statistical significance was determined by repeated measures ANOVA and a Bonferroni post-hoc test. * = p<0.05 and ** = p<0.01. (E) Immunoblot analysis of lysates from RPE cells stimulated with poly(I:C) for indicated times and probed with OPTN and EF2 (loading control) antibodies. (F) Confocal microscope images of RPE cells stably expressing GFP-OPTN (green) and treated with poly(I:C) as specified. Cells were immunostained with an antibody against TGN46 [i] and VTI1A [ii] (red). Scale bars, 20 µm. (G) Pearson’s correlation coefficient calculated for GFP-OPTN versus VTI1A after treatment with poly(I:C) for 0 and 24 hours. Bars represent the mean of n=3 independent experiments ±SEM. Cells were quantified from ≥20 randomly selected fields of view (1 cell/image). Statistical significance was calculated using a two-sample t-test. *** = p<0.001. (H) Dose-response curve of foci count/GFP-OPTN cell after treatment with poly(I:C) for 24 hours in combination with the indicated dose of BX795. Points represent mean of n=3 experiments ±SEM. (I) Confocal microscope images of RPE cells stably expressing GFP-OPTN (green) and treated with poly(I:C) for 24 hours in combination with DMSO (left panel) or BX795 (right panel). Scale bar, 20 µm.

To further analyse the nature of this perinuclear OPTN-positive compartment, we labelled GFP-OPTN expressing cells with a variety of organelle markers. The foci showed very little overlap with markers of the endocytic pathway including EEA1 and LAMP1 (Fig. S2B[i-ii]). Notably, the foci could be observed in close proximity to, but only showed partial colocalisation with the *trans*-Golgi marker TGN46, the cation-independent mannose-6-phosphate receptor or the autophagosomal membrane marker LC3 (Fig. 1F[i] and Fig. S2B[iii-iv]). Further observations indicated strong colocalisation with the OPTN-binding partner MYO6 (Fig. S2C) and with the Golgi SNARE VTI1A, (Fig. 1F[ii]-G) suggesting some continuation with the Golgi complex. Depletion of MYO6 by siRNA had no effect on the formation of the foci indicating that the recruitment of OPTN to these structures and the formation of the foci was not dependent on MYO6 (Fig. S2D).

Importantly, TLR3 was not recruited into OPTN foci suggesting that this compartment is distinct from the route of receptor trafficking (Fig. S3A). Furthermore, perturbation of TLR3 expression using CRISPRi largely blocked poly(I:C)-induced OPTN foci formation indicating that this phenotype is dependent on TLR3 receptor-driven signalling (Fig S3B-D).

### TBK1 activity is necessary for OPTN recruitment to foci but not their long-term stability

Given the well-established role of TBK1 and OPTN in the antiviral response [38], we next assessed the role of TBK1 in foci formation. TBK1 activity measured through the increase in phosphorylation (p-TBK1) was evident 30 mins post-TLR3 stimulation and returned to baseline levels after 8 hours (Fig. S4A). Using a specific inhibitor of TBK1, BX795, OPTN foci formation could be abolished in a dose-dependent fashion downstream of TLR3 activation (Fig. 1H-I). Interestingly, addition of BX795 six hours after poly(I:C) stimulation did not influence foci formation (Fig. S4B-C), which indicates that TBK1 kinase activity is required for initiation of the foci but is dispensable for the subsequent maintenance of the structure.

### OPTN disease mutants show perturbed foci formation

Previous work has linked the OPTN E50K mutant to POAG and has shown that OPTN overexpression causes the formation of large perinuclear foci in cells [1,31,32,34,35]. Conversely, the E478G mutation, which is linked to ALS, appears to lack this capacity [4,31]. We predicted that these mutants might show a perturbed ability to form foci in response to poly(I:C) stimulation. Strikingly, ∼95% of RPE cells expressing GFP-OPTN E50K exhibited a constitutive formation of this compartment even in the absence of stimuli, compared to around 5% of cells expressing wild-type GFP-OPTN (Fig. 2A-B). TLR3 stimulation resulted in ∼80% of wild-type GFP-OPTN expressing cells making foci, whereas stimulation had minimal effect on the GFP-OPTN E50K cells that retained foci in ∼95% of cells. By contrast, cells expressing GFP-OPTN E478G were completely unable to generate foci even after 24 hours of poly(I:C) stimulation (Fig. 2A-B).

**Figure 2.**
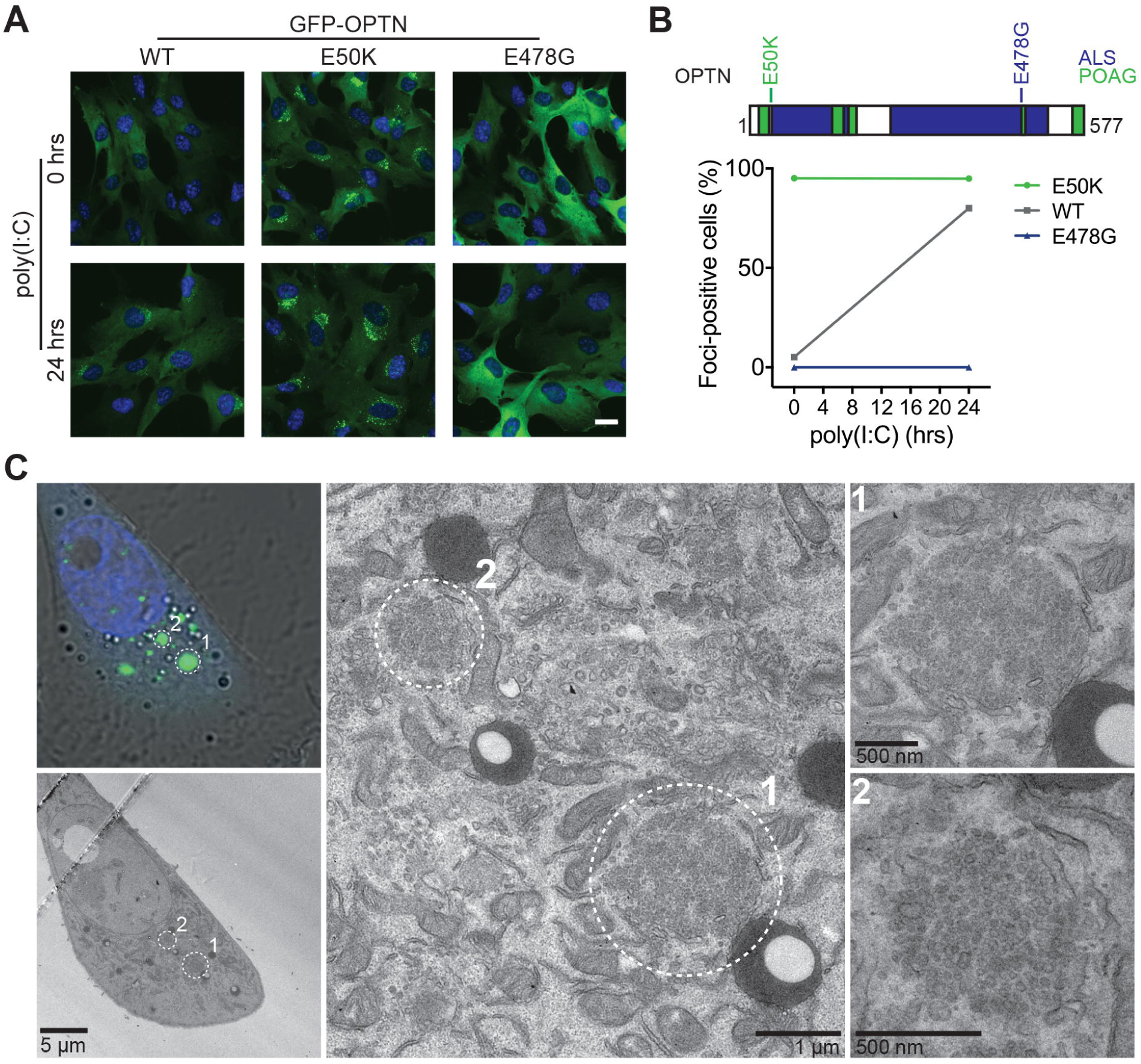
OPTN disease mutants promote aberrant foci formation. (A) Widefield microscope images of RPE cells stably expressing GFP-OPTN wild-type (WT), E50K and E478G (green) and treated for 0 hours or 24 hours with poly(I:C). Cells were stained with Hoechst to label DNA (blue). Scale bar, 20 µm. (B) Top, schematic cartoon of OPTN domain structure with mutations highlighted. Bottom, graph depicting the percentage of GFP-OPTN cells containing foci after 0 or 24 hours of poly(I:C) treatment from n=3 independent experiments ±SEM. >100 cells were manually counted per condition from ≥10 randomly selected fields of view. (C) Correlative Light Electron Microscopy (CLEM) micrographs of RPE cells stably expressing GFP-OPTN E50K. Left column shows fluorescence microscope image of cell (top) and corresponding electron micrograph (bottom). Two GFP-positive foci are highlighted by the circled regions 1 and 2. The central panel shows a magnified area around regions 1 and 2 and the right column shows further magnification of regions 1 (top) and 2 (bottom).

To visualise the nature and further define the composition of the OPTN-positive compartment, we performed correlative light electron microscopy (CLEM) on foci generated by the OPTN E50K mutant. Cells were first imaged by confocal microscopy to determine the localisation of the GFP-OPTN E50K-positive foci and then processed for electron microscopy. CLEM images showed that the foci were composed of tightly-packed small membrane vesicles contained within a spherical area void of any further delimiting membrane (Fig. 2C). As aggresomes are typically membrane-less, electron dense structures [39], our data would appear to rule out the possibility that OPTN foci are simply protein inclusions, but are clearly a membranous compartment consisting of a cluster of small vesicles of uniform size.

### OPTN foci are composed of tightly-packed ATG9A-positive membrane vesicles

ATG9A has been implicated in the innate immune response to cytosolic DNA where it regulates the assembly of STING and TBK1 on a vesicular Golgi-associated perinuclear compartment [40]. To determine whether the cellular response to viral RNA involves a similar ATG9A compartment, we determined whether the OPTN-positive vesicles colocalise with ATG9A. In unstimulated RPE cells, ATG9A is present at the Golgi complex, however, after poly(I:C) stimulation the newly formed GFP-OPTN foci are positive for ATG9A (Fig. 3A-C). Observations of OPTN mutants revealed that GFP-OPTN E50K appeared to trap ATG9A on Golgi-proximal foci even in the absence of stimuli, while GFP-OPTN E478G failed to colocalise with ATG9A even after stimulation (Fig. 3A-C). High resolution microscopy reveals the presence of distinct ATG9A-vesicle clusters within the OPTN-positive foci. Interestingly, the ATG9A-vesicles are occasionally in close proximity but show only limited overlap with autophagosomes (Fig. S2C), confirming previous data that the ATG9A-vesicles might interact with but do not appear to be incorporated into the growing phagophore.

**Figure 3.**
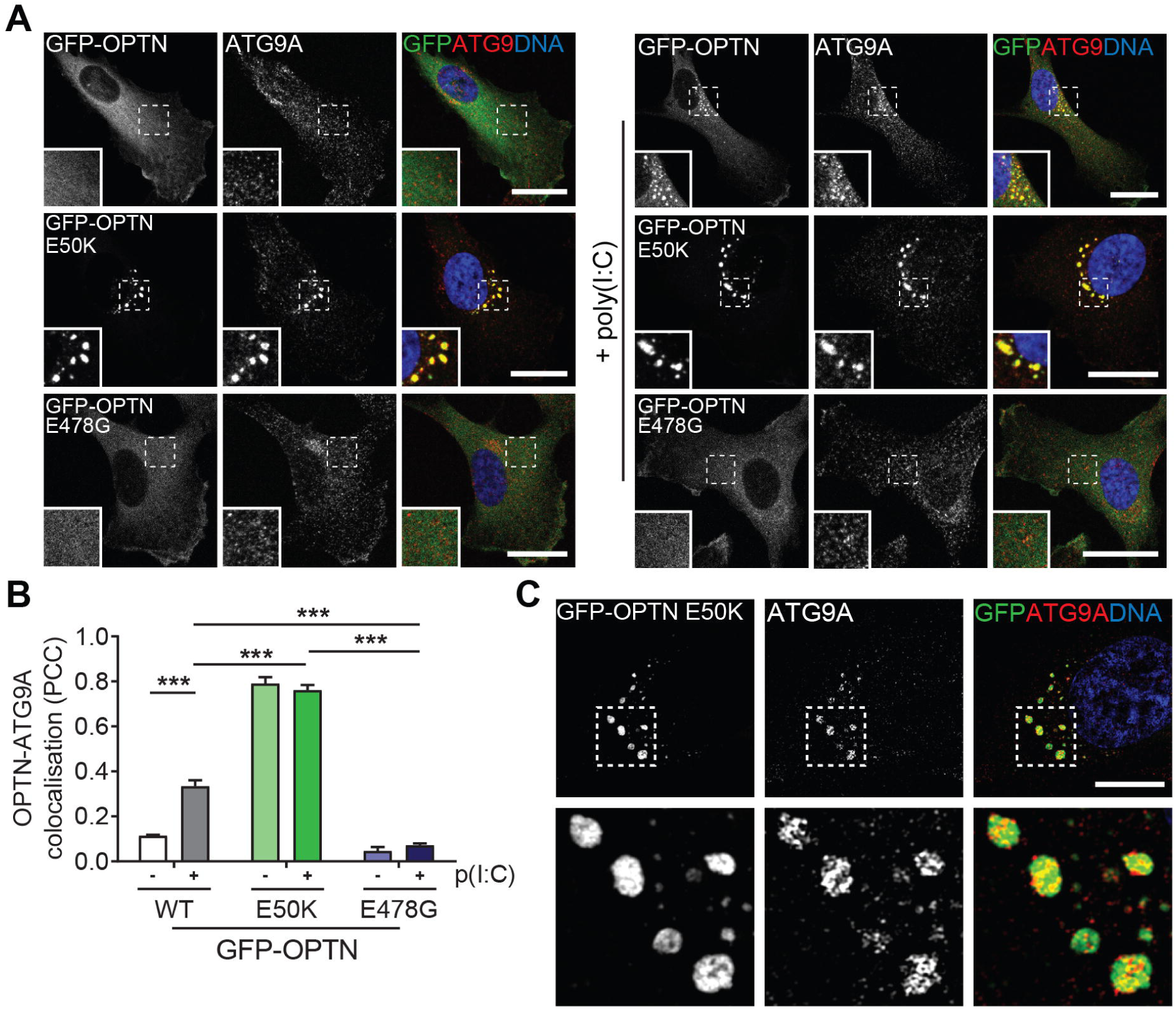
OPTN foci are composed of ATG9A vesicles. (A) Confocal microscope images of RPE cells stably expressing GFP-OPTN, E50K or E478G (green) and treated with poly(I:C) for 0 hours or 24 hours. Cells were immunostained with an ATG9A antibody (red) and Hoechst to label DNA (blue). (B) Pearson’s correlation coefficient calculated for GFP-OPTN (WT, E50K or E478G) versus ATG9A after treatment with poly(I:C) for 0 and 24 hours. Bars represent the mean of n=3 independent experiments ±SEM. Cells were quantified from ≥10 randomly selected fields of view. Statistical significance was determined by repeated measures ANOVA and a Bonferroni post-hoc test. *** = p<0.001. (C) Structured illumination microscopy image of RPE cell stably expressing GFP-OPTN E50K (green), immunostained with ATG9A antibody (red) and Hoechst to label DNA (blue). Scale bar, 10 µm. Lower panels are magnifications of the insets highlighted above.

### BioID reveals novel OPTN partners and foci proteins

To gain further insight into both OPTN function and the composition of the foci, we determined the OPTN interactome using *in situ* proximity labelling. We generated RPE stable cell lines expressing full-length OPTN tagged at the N- or C-terminus with the promiscuous biotin ligase, BirA R118G (BirA*). Expression of the BirA*-OPTN or OPTN-BirA* fusion proteins was verified by immunoblotting and the localisation assessed by immunofluorescence (Fig. S5A-B). After labelling with biotin overnight, streptavidin pulldowns were performed and enriched proteins identified by mass spectrometry.

Replicates were analysed against a bank of BirA* only controls using the online tool at Crapome.org and using a threshold FC-B score of ≥3, we identified 25 significantly enriched proteins (Table 2) [41]. Among the proteins we identified were a number of known OPTN-interacting proteins and complexes such as TBK1, CYLD, TBC1D17, and the LUBAC component HOIP (RNF31) in addition to novel putative interactors such as the myotubularin-related (MTMR) lipid phosphatase complex components MTMR6 and 9, intraflagellar transport 74 (IFT74) and Interferon Induced Protein 35 (IFI35) (Fig. 4A-B).

**Figure 4.**
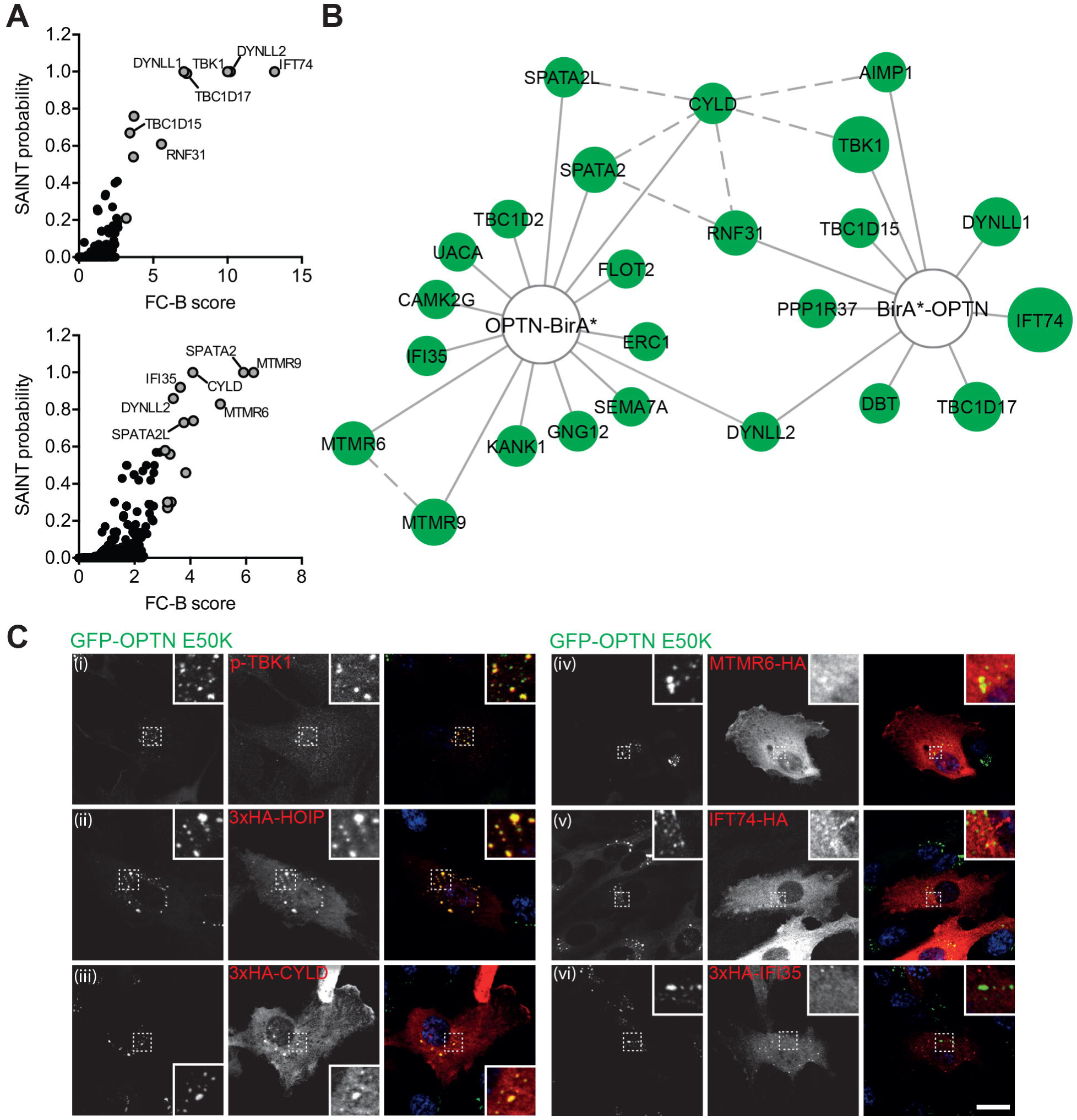
OPTN BioID reveals novel partners and proteins localised to foci. (A) Graphs depicting SAINT probability and fold change (FC-B) scores for BirA*-OPTN (top) and OPTN-BirA* (bottom) pull down experiments. Selected high-confidence OPTN interactors are labelled. (B) Network diagram of high-confidence OPTN interactors identified by BioID. Node size corresponds to FC-B score (higher confidence = larger node). Solid lines indicate interactions identified in this study and dashed lines interactions imported from publicly available protein-protein interaction databases. (C) Confocal microscope images of RPE cells stably expressing GFP-OPTN E50K (green) and immunostained with a p-TBK1(i; red) or HA antibody (ii-vi; red). Scale bar, 20 µm.

We screened a selection of these candidates for their ability to localise to GFP-OPTN E50K-induced foci (Fig. 4C) including p-TBK1, which has been shown previously to colocalise with OPTN (Fig. 4C[i]) [20]. Interestingly, the E3 ligase, HOIP, as well as the DUB, CYLD, both showed colocalisation on OPTN foci, although CYLD only showed recruitment in a small subpopulation of cells (Fig. 4C[ii]-[iii]). In contrast, MTMR6, IFT74 and IFI35 showed little recruitment to OPTN foci (Fig. 4C[iv]-[vi]), and might interact with OPTN within other cellular pathways such as autophagy.

### The linear ubiquitin assembly complex (LUBAC) is recruited to OPTN foci

NEMO interacts with, and is linearly ubiquitinated by, LUBAC to induce the activation of the IKK complex [9,15]. OPTN also binds LUBAC components HOIP and HOIL-1L and regulates the interaction of RIPK1 and NEMO with the TNF receptor (TNFR) complex in response to TNF-α [22]. Our BioID experiments are consistent with the concept that HOIP interacts with OPTN but also indicate a possible corecruitment to OPTN foci. Furthermore, the potential cooperation of HOIP and OPTN in TLR3 signalling remains unexplored.

We investigated the role of LUBAC at OPTN foci by assessing recruitment of HOIP to wild-type GFP-OPTN foci. Initially, HOIP showed little colocalisation with OPTN in unstimulated cells; however, poly(I:C) stimulation led to the recruitment of HOIP to GFP-OPTN positive vesicles (Fig. 5A). Quantification of the colocalisation confirmed this, but notably, unstimulated cells expressing the GFP-OPTN E50K mutant showed much higher HOIP recruitment than wild-type GFP-OPTN even after wild-type GFP-OPTN cells were treated with poly(I:C) (Fig. 5B). Next, we tested whether other components of the LUBAC complex were also recruited to the OPTN-positive foci and found that upon TLR3 stimulation both SHARPIN and HOIL-1L showed strong colocalisation (Fig. 5C). To confirm the interaction between OPTN and HOIP, we performed GFP immunoprecipitations from HEK293T transiently transfected with wild-type, E50K or E478G GFP-OPTN and HA-HOIP. Wild-type and E50K of GFP-OPTN coimmunoprecipitated HA-HOIP, but GFP-OPTN E478G, which lacks foci, failed to do so (Fig. 5D).

**Figure 5.**
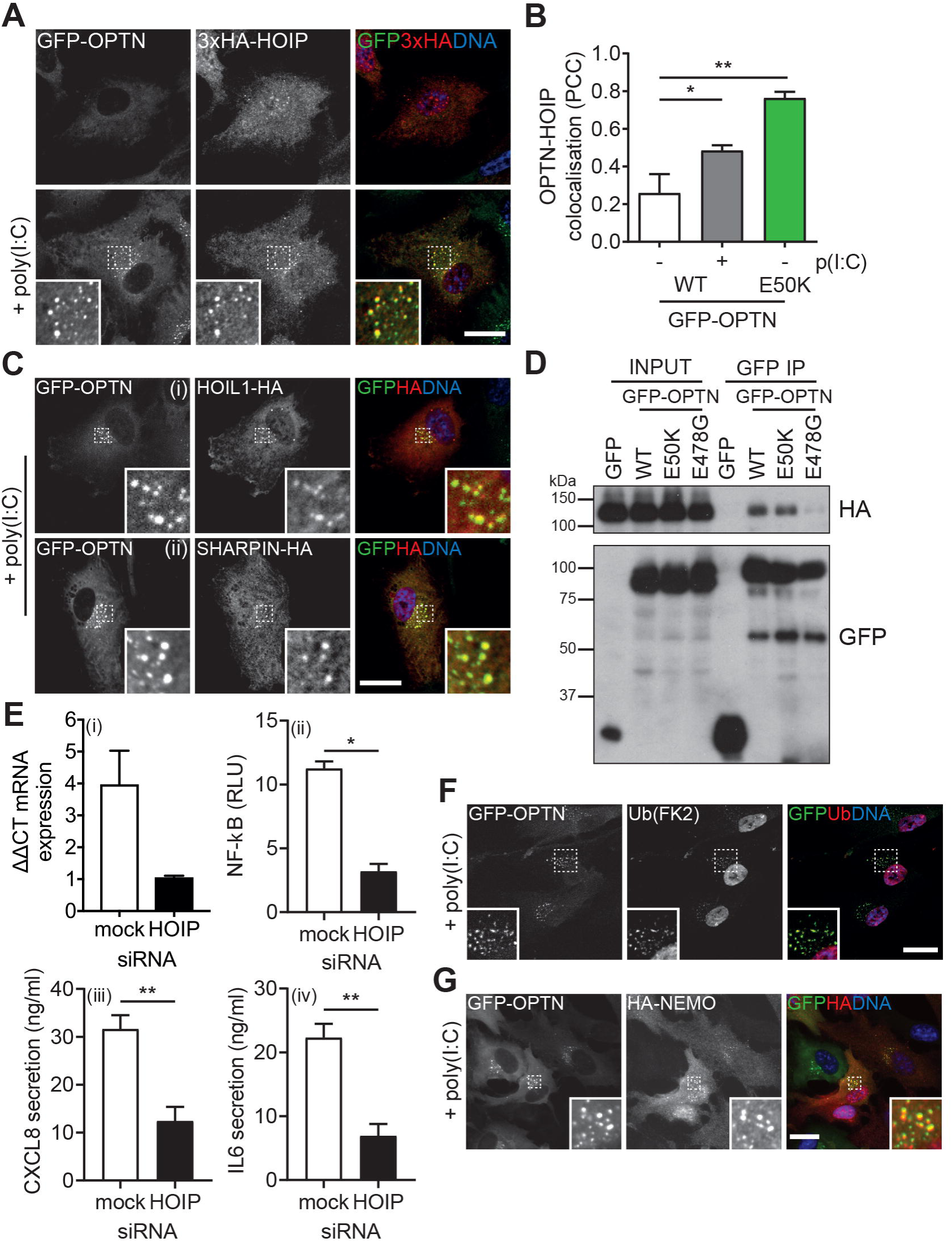
The LUBAC complex is recruited to foci. (A) Confocal microscope images of RPE cells stably expressing GFP-OPTN (green) and 3xHA-HOIP and treated with poly(I:C) for 0 hours and 24 hours. Cells were immunostained with a HA antibody (red) and Hoechst to label DNA (blue). Scale bar, 20 µm. (B) Pearson’s correlation coefficient calculated for GFP-OPTN versus HOIP after treatment with poly(I:C) for 0 and 24 hours. Bars represent the mean of n=3 independent experiments ±SEM. Cells were quantified from ≥5 randomly selected fields of view (1-2 cells/image). Statistical significance was calculated by one-way ANOVA and a Bonferroni post-hoc test. * = p<0.05 & ** = p<0.01. (C) Confocal microscope images of RPE cells stably expressing GFP-OPTN (green) and HOIL1-HA (top) or SHARPIN-HA (bottom) and treated with poly(I:C) for 24 hours. Cells were immunostained with a HA antibody (red) and Hoechst to label DNA (blue). Scale bar, 20 µm. (D) Immunoblot of GFP immunoprecipitations from HEK293T transiently transfected with GFP, GFP-OPTN wild-type (WT), E50K and E478G probed with GFP and HA antibodies. (E) Graphs of HOIP mRNA expression [i], NF-κB luciferase activity [ii] and CXCL8 [iii] and IL6 secretion [iv] in RPE cells transfected with mock or HOIP siRNA and treated with poly(I:C) for 24 hours. Bars depict mean of n=3 independent experiments ±SEM. Statistical significance was determined by two-sample t-test. * = p<0.05, ** = p<0.01. (F) Confocal microscope images of RPE cells stably expressing GFP-OPTN (green) and treated with poly(I:C) for 24 hours. Cells were immunostained with an anti-ubiquitin antibody (clone FK2; red) and Hoechst to label DNA (blue). Scale bar, 20 µm. (G) Confocal microscope images of RPE cells stably expressing GFP-OPTN (green) and HA-NEMO. Cells were treated with poly(I:C) for 24 hours and immunostained with a HA antibody (red) and Hoechst to label DNA (blue). Scale bar, 20 µm.

We assessed the contribution of HOIP (and LUBAC) to NF-κB signalling upon TLR3 activation in RPE cells. Depletion of HOIP by siRNA diminished poly(I:C)-induced NF-κB luciferase activity and secretion of CXCL8 and IL6 (Fig. 5E[i-iv]), confirming that HOIP plays a critical role in NF-κB activation downstream of TLR3 in these cells. This data suggests that OPTN can sequester positive regulators of NF-κB signalling in perinuclear foci.

### OPTN foci formation and stabilisation require ubiquitination

The presence of LUBAC on OPTN foci implied the presence of linear ubiquitin chains on this compartment. Indeed, the OPTN E478G mutant, which is characterised by its inability to bind ubiquitin [42] or HOIP, is no longer able form foci (Fig. 2 & 4C). To ascertain the role of ubiquitin on this compartment we labelled poly(I:C)-induced OPTN foci with an antibody against ubiquitin (FK2), which recognises a variety of chain types including linear [43]. In unstimulated cells, antibody staining was very weak and nuclear but after poly(I:C) treatment the ubiquitin FK2 signal was present on GFP-OPTN-positive foci (Fig. 5G). OPTN has a ubiquitin binding domain that is homologous to NEMO and which binds to linear ubiquitin chains. We cloned a previously described probe composed of 3 tandem repeats of the NEMO UBAN domain (RFP-3xUBAN), which shows a 100-fold specificity for M1-linked linkages over other chain types [44]. This probe was recruited to the perinuclear foci upon poly(I:C) stimulation and could be blocked by introduction of the F312A point mutation known to abolish ubiquitin binding (Fig. S6A). The presence of ubiquitin chains, LUBAC, OPTN and the 3xUBAN probe on the foci prompted us to investigate whether NEMO itself was also recruited. Indeed, poly(I:C) treatment of RPE cells stably expressing HA-tagged NEMO triggered its recruitment to OPTN foci (Fig. 5C). The presence of both LUBAC and NEMO on these OPTN-positive foci is highly suggestive of a regulatory role in NF-κB signalling by sequestering these components downstream of TLR3.

Despite the requirement for ubiquitin binding in the recruitment of OPTN as demonstrated by the E478G mutant, siRNA depletion of HOIP had little effect on OPTN relocalisation to foci (Fig. S6B), and suggests that OPTN recruitment is not solely dependent on LUBAC-synthesised linear ubiquitin. As the OPTN UBAN domain is capable of binding to both K63-linked and linear ubiquitin, but not to K48-linked ubiquitin [22], we hypothesised other chain types might also be present. Expression of a ubiquitin mutant construct containing a single lysine residue at K63 was also present on the foci, indicating they are likely to be a mixture of both linear and K63 chains (Fig. S6C), and thus it is possible that K63 chains are sufficient for the initial recruitment of OPTN.

### OPTN foci formation correlates with innate immune signalling and cytokine secretion

The rate of foci formation correlated well with time courses for both the induction of cytokine secretion and the inhibition of NF-κB or IRF3 signalling. In addition, the presence of multiple regulators of NF-κB and IRF3 signalling (LUBAC, NEMO and TBK1) suggested a link between OPTN-induced foci and regulation of these signalling pathways. Previous work has shown that OPTN is a negative regulator of NF-κB and IRF3 signalling and that ALS mutations or loss of ubiquitin binding perturb these functions [20,22]. Therefore, we investigated NF-κB activity in parental RPE cells or RPE cells expressing E50K or E478G and observed a negative correlation between NF-κB activation and OPTN foci formation. Cells expressing GFP-OPTN E50K markedly inhibited NF-κB activity and GFP-OPTN E478G cells showed elevated activity relative to non-expressing control cells (Fig. 6A). Next, we assessed the effect of these mutations on cytokine secretion downstream of NF-κB signalling. RPE cells overexpressing GFP-OPTN E50K showed a reduction in CXCL8 and IL6 secretion, whereas OPTN E478G cells displayed a dramatic increase in secretion of both (Fig. 6B-C). These results were consistent with data obtained by immunoblotting (Fig. 6D).

**Figure 6.**
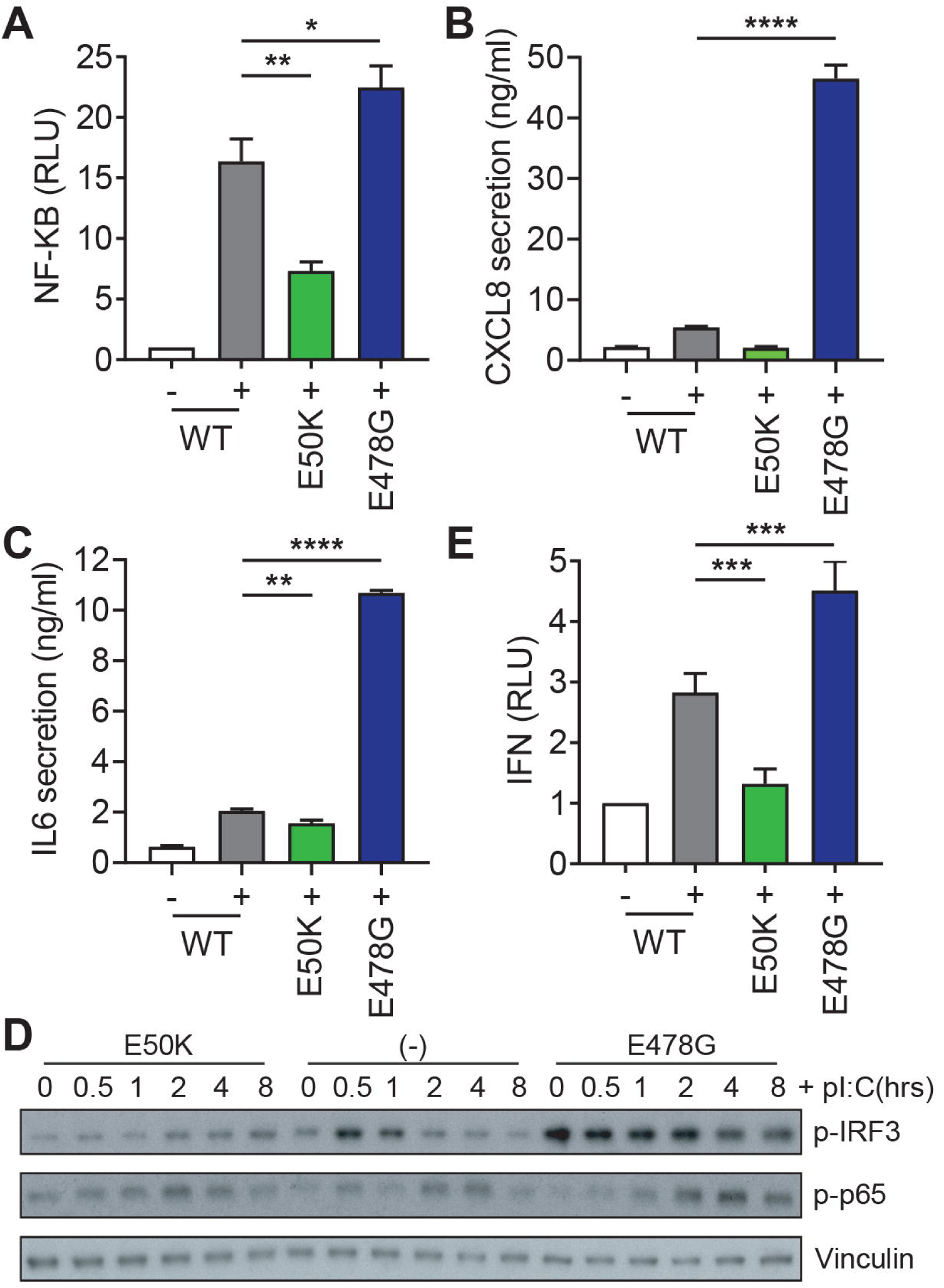
OPTN mutations regulate innate immune signalling and cytokine secretion. (A) Relative NF-κB luciferase activity in RPE cells expressing an NF-κB luciferase reporter, coexpressing GFP-OPTN E50K and E478G and stimulated with poly(I:C) as indicated. Graphs depicts mean of n=6 independent experiments ±SEM. Statistical significance was calculated by one-way ANOVA and a Bonferroni post-hoc test. * = p<0.05 and ** = p<0.01. (B-C) CXCL8 (B) and IL6 (C) secretion from RPE cells expressing GFP-OPTN E50K and E478G and stimulated with poly(I:C) as indicated. Graphs depicts mean of n=6 independent experiments ±SEM. Statistical significance was calculated by one-way ANOVA and a Bonferroni post-hoc test. ** = p<0.01, *** = p<0.001 and **** = p<0.0001. (D) Western blots of lysates from RPE cells expressing GFP-OPTN E50K or E478G, stimulated with poly(I:C) and probed with the indicated antibodies. (E) IFNα/β secretory levels from RPE cells coexpressing GFP-OPTN E50K and E478G and stimulated with poly(I:C) for 6 hours determined from luciferase activity induced in the ISRE-reporter cell line 3C11. Graph depicts mean of n=5 independent experiments ±SEM. Statistical significance was calculated by one-way ANOVA and a Bonferroni post-hoc test. *** = p<0.001.

As RIG-I stimulation with pppRNA also induced OPTN foci formation, we next investigated whether OPTN regulated cytokine secretion in this context. As with TLR3-stimulation, the E50K mutant reduced CXCL8 and IL6 secretion in response to pppRNA, while the converse is true for the E478G mutant (Fig. S7A-B). Thus, OPTN appears to regulate the innate immune response to viral RNA generally.

Since OPTN has also been implicated in IRF3 signalling, we next determined the impact of OPTN mutations on this pathway. We investigated the activity of this pathway in mutant cells lines and, again, found that overexpression of the OPTN E50K mutant blunted the IRF3 response, as determined by p-IRF3 immunoblot and IFNα/β release assays (Fig. 6D-E). Conversely, the OPTN E478G mutant showed high levels p-IRF3 prior to stimulation, which remained elevated, and a concomitant increase in IFNα/β secretion (Fig. 6D-E). Thus, the propensity to form foci correlates well with NF-κB and IRF3 signalling output and appears to indicate that the formation or presence of foci is refractory to both signalling pathways.

## Discussion

In order to establish an appropriate immune response and prevent chronic inflammation cells must tightly regulate innate immune signalling and cytokine secretion. The central role of OPTN in negatively regulating these signalling pathways is becoming increasingly clear and different mutations, which modify the ability of OPTN to modify these pathways, appear to lead to distinct diseases. Here we establish an RPE cell model to investigate the role of OPTN in innate immune signalling. Using this system, we show that OPTN translocates to Golgi-proximal foci in response to exogenous RNA and that this compartment negatively regulates downstream signalling responses. Expression of different disease-causing OPTN mutants leads to either constitutive foci formation in the absence of stimulation, and a concurrent attenuation of IRF3 and NF-κB signalling and cytokine secretion, or the converse.

Our ultrastructural characterisation of the OPTN-positive foci reveals that they are not aggresomes as previously suggested [30], but clusters of tightly packed small vesicles of around 30-40 nm. This vesicle cluster is concentrated in a concise space despite lacking an outer limiting membrane. Our double-labelling experiments suggest that the OPTN foci strongly overlap with ATG9A, a transmembrane protein with a key role in autophagy. ATG9A has a very dynamic trafficking itinerary cycling between the Golgi complex and the endocytic pathway [45]. Thus, OPTN might regulate post-Golgi trafficking and sorting of ATG9A-containing vesicles to the phagophore. In addition, as a selective autophagy receptor, OPTN may control the spatiotemporal recruitment of ATG9A vesicles to the site of autophagosome formation. This agrees with the recent finding that autophagy receptors cooperate with TBK1 to recruit the ULK1 complex to initiate autophagosome formation [46]. Therefore, the OPTN- and ATG9A-positive foci could be a compartment that accumulates post-Golgi trafficking intermediates or marks the site of autophagosome biogenesis.

Our work also highlights the correlation between the formation of OPTN foci and the role of OPTN in negatively regulating of NF-κB and IRF3 signalling. Our data demonstrates that OPTN expression is upregulated in poly(I:C)-stimulated RPE cells and occurs with kinetics similar to those of both NF-κB inactivation and OPTN foci formation. Furthermore, we were able to identify and localise several key mediators of NF-κB and IRF3 signalling to OPTN foci, including TBK1, NEMO, CYLD and components of the LUBAC complex. Taken together, this data suggests a model in which NF-κB signalling generates a negative feedback mechanism to prevent excessive signalling via upregulation of OPTN expression. We propose that the expression of OPTN is tied to its propensity to oligomerise via dimer or tetramerisation or polyubiquitin chain binding leading to foci formation, sequestration of NF-κB or IRF3 signalling machinery and the inhibition of further signalling, possibly via autophagy. The OPTN E50K mutant displays a heightened propensity to form oligomers [47], and this property may explain the observed constitutive foci. Alternatively, the loss of ubiquitin binding seen with the OPTN E478G mutant might prevent foci formation by blocking oligomerisation through polyubiquitin chain binding. Other disease-associated mutations may also alter the ability of OPTN to oligomerise or to recruit proteins into foci and lead to perturbed downstream outputs.

Notably, the OPTN foci described here also show striking similarity to those described in a number of other situations. In particular, activation of the cGAS-STING pathway by cytosolic DNA leads to the trafficking of STING from the ER to an ER-Golgi intermediate compartment (ERGIC), which is positive for ATG9A [40,48]. Trafficking of STING from the ER to this compartment is required for the induction of IRF3 signalling, while ATG9A negatively regulates this process [40,49]. Recent work has also defined a role for STING in the induction of autophagy in response to cGAMP, cytosolic DNA or DNA viruses and that the ERGIC serves a membrane source for autophagosome formation in this context [50]. As an important mediator of autophagy and innate immune signalling, it is tempting to speculate that OPTN might participate in an analogous process in response to exogenous dsRNA or RNA viruses. Other proteins including the NLRP3 inflammasome or OPTN binding partners TRAF3 and TBK1 have also been found to localise to similar Golgi-proximal perinuclear microsomes upon stimulation [38,51,52], suggesting that this Golgi-proximal platform might be a common mechanism to regulate signalling, cytokine secretion and autophagy induction in response to diverse PAMPs.

## Materials and Methods

### Antibodies, plasmids and reagents

Antibodies used in this study were CIMPR (sc-53146; Santa Cruz), EEA1 (610457; BD Biosciences), EF2 (sc-13004; Santa Cruz), GFP (A11122; Life Technologies), HA (11867423001; Roche), HA (H9658; Sigma), LC3 (M152-3; MBL), LAMP1 (H4A3; Developmental Studies Hybridoma Bank, University of Iowa), myc (05-724; Millipore), OPTN (HPA003360; Sigma), p-IRF3 (4947; Cell Signalling), p-p65 (3033; Cell Signalling), p-TBK1 (5483; Cell Signalling; 5483), TGN46 (AHP500; Bio-Rad), ubiquitin (BML-PW8810; Enzo Life Sciences), vinculin (MAB3574; Millipore) and Vti1a (611220; BD Biosciences). The ATG9A antibody (ab108338; Abcam) was a kind gift from Professor Margaret S. Robinson (CIMR). Rabbit polyclonal antibodies raised against GFP and MYO6 were generated in house as described previously [53].

Cells were treated with poly(I:C) (Enzo Life Sciences) at 10 μg/mL, LPS (Enzo Life Sciences) at 200 ng/ml, 2’,3’-cGAMP (Invivogen) at 10 μg/mL, Pam3CSK4 (Invivogen) at 10 μg/mL and BX795 (Sigma) at 500 nM. All treatments were for 24 hours unless specified otherwise.

GFP-OPTN pEGFPC2 has been described previously [54] and was subcloned into the pLXIN retroviral packaging plasmid (Clontech) for stable cell line production. GFP-OPTN E50K and E478G pLXIN mutants were generated by site-directed mutagenesis. The myc-BirA*-OPTN pLXIN vector was created by subcloning OPTN into the myc-BirA*-OPTN pLXIN plasmid described previously [55]. For OPTN-BirA*-HA pLXIN, BirA* was amplified by PCR, introducing a C-terminal HA tag, and inserted into pLXIN. OPTN was subcloned into this vector 5’ to the BirA* tag.

HA-Ub K63 pRK5 and NF-κB-TA-LUC-UBC-GFP-W pHAGE were obtained from Addgene (17606 and 49343 respectively). NEMO, TLR3, CYLD, SHARPIN, HOIP and RBCK1 were obtained from Addgene (13512, 13641, 15506, 50014, 50015 and 50016 respectively) and subcloned into pLXIN. Full-length IFT74 was generated by Gibson assembly of MGC clones 8322576 and 6614193 (Dharmacon, GE Healthcare). MTMR6 was obtained from Sino Biologicals (HG15192) and the IFI35 open reading frame was synthesised as a Gblock from Integrated DNA technologies. All were subcloned into pLXIN with HA tags.

The CRISPRi lentiviral vector pU6-sgRNA EF1Alpha-puro-T2A-BFP was a kind gift from Luke Gilbert. Protospacer sequences targeting TLR3 5’-GATTTCATCAGGGAAGTGTG-3’ or a control non-targeting sequence (GAL4) 5’-GAACGACTAGTTAGGCGTGTA-3’ were inserted by restriction cloning.

3xUBAN pRFPC3 was generated as a gBlock (Integrated DNA technologies) comprising the UBAN sequence of NEMO flanked by a 5’ SalI site and 3’ XhoI-BamHI sites. Plasmid DNA was linearised with XhoI and BamHI and ligated with gBlock DNA digested with SalI and BamHI. Complementary SalI and XhoI overhangs were ligated, destroying the restriction sites and leaving a unique XhoI site at the 3’ end of the UBAN open reading frame which could be used in subsequent cloning steps. This process was repeated 3 times to generate 3 tandem duplicates of the UBAN sequence.

### Cell lines and transfection

RPE1 cells were cultured in DMEM:F12-HAM (Sigma) mixed in a 1:1 ratio and supplemented with 10% FBS (Sigma), 2 mM L-glutamine (Sigma), 100 U/ml penicillin and 100 µg/ml streptomycin (Sigma). HEK293T and Phoenix cells were cultured in DMEM containing GlutaMAX (Thermo Fisher Scientific) and supplemented with 10% FBS and 100 U/ml penicillin and 100 µg/ml streptomycin.

Stably expressing cell lines were generated using retrovirus or lentivirus produced in the Phoenix retroviral packaging cell line or HEK293T cells respectively. Cells growing in 100 mm dishes were transfected with 10 µg retroviral transfer vector DNA and 25 µl Lipofectamine 2000 (Thermo Fisher Scientific) or 8 µg lentiviral transfer vector DNA, 8 µg pCMV-dR8.91 and 1 µg pMD2.G packaging plasmids using 48 µl TransIT-LT1 (Mirus). Plasmid DNA was mixed with transfection reagent in Opti-MEM (Thermo Fisher Scientific) and incubated for 30 minutes before addition to cells. After 48 hours, conditioned medium was harvested, filtered and added to the relevant cells. Cells were subsequently selected with 500 µg/ml G418 (Gibco), 1 µg/ml puromycin or sorted by FACS. RPE1 dCas9-KRAB cells were a kind gift from Ron Vale.For immunoprecipitation experiments, HEK293T cells were transfected in 100 mm dishes using 8 µg plasmid DNA and 24 µl PEI (Polysciences, Inc). DNA was mixed with PEI in Opti-MEM (Thermo Fisher Scientific), incubated for 20 minutes and added to cells. For siRNA-mediated gene silencing, RPE cells were transfected with ON-TARGETplus SMARTpool oligonucleotides (Dharmacon, GE Healthcare) targeting MYO6 or HOIP using Oligofectamine (Invitrogen). Cells were transfected on both day 1 and day 3 and assayed on day 5.

For RIG-I stimulation experiments, 1 μg pppRNA (Invivogen) was added to 100 μl LyoVec (Invivogen), incubated for 15 minutes at RT, transfected into cells at a final concentration of 500 ng/ml, and incubated for 24 hours.

### Cytokine assays

Cytokine (IL6 and CXCL8) levels in tissue culture supernatants were determined by ELISA assay (DY206 and DY208; R&D Systems). All assays were performed according to the manufacturer’s instructions and read on a CLARIOstar microplate reader (BMG Labtech). ELISA data was normalized to viable cell number determined by MTT assay (Boehringer Ingelheim) or CellTiter-Blue (Promega). IFN levels were determined using a HEK293T IFN reporter cell line (clone 3C11) which was obtained from Prof. Jan Rehwinkel (University of Oxford, UK) [56]. For the IFN assay, IFN reporter cells were cultured on a 96-well plate with 70 μL DMEM medium overlaid with 30 μL of cell culture supernatant. After 24 hours, luciferase expression was quantified using a Pierce™ Firefly Luc One-Step Glow Assay Kit (Thermo Fisher Scientific) according to manufacturer’s instructions and read on a FLUOstar Omega microplate reader (BMG Labtech).

### qPCR

Total RNA was harvested using a RNeasy Mini Kit and RNase-free DNase treatment (Qiagen), in accordance with the manufacturer’s instructions. RNA (1 μg) was converted to cDNA using oligo d(T) primers and Promega reverse transcription kit. Quantitative real time PCR (qRT-PCR) was performed in duplicate using a SYBR® Green PCR kit (Qiagen) on a Mastercycler® ep realplex (Eppendorf) or Quantstudio 7 flex (Life Technologies). The PCR mix was annealed/extended at 60 °C for 60 seconds, for a total of 40 cycles, then a melting curve was performed. Primers for HOIP were 5’-AGACTGCCTCTTCTACCTGC-3’ and 5’-CTTCGTCCCTGAGCCCATT-3’, TLR3 set 1 5’-TCAACTCAGAAGATTACCAGCCG-3’ and 5’-AGTTCAGTCAAATTCGTGCAGAA-3’, TLR3 set 2 5’-CAAACACAAGCATTCGGAATCTG-3’ and 5’-AAGGAATCGTTACCAACCACATT-3’ and the housekeeper gene peptidylprolyl isomerase A (PPIA) 5’-GTGTTCTTCGACATTGCCGT-3’ and 5’-CCATTATGGCGTGTGAAGTCA-3’ or Actin 5’-GCTACGAGCTGCCTGACG-3’ and 5’-GGCTGGAAGAGTGCCTCA-3’. Relative expression was compared between groups using the ΔΔCt method [57].

### Cell lysate preparation

Cells were plated in a 6-well plate and stimulated at ∼80% confluent. Cells were washed and lysed in RIPA buffer (50 mM Tris-HCl, 150 mM NaCl, 0.5% sodium deoxycholate, 1 mM EDTA, 0.5 mM EGTA, 1% IGEPAL® CA-630, and 0.1% SDS) containing protease inhibitor cocktail (Roche) and PhosSTOP™ (Sigma). Cell lysates were sonicated and clarified at 20,000 xg for 10 minutes at 4°C. Total protein concentration was measured using a Pierce™ BCA Protein Assay Kit (Thermo Fisher Scientific) and used to normalise sample loading.

### Immunoprecipitation

48 hours post-transfection, cells were lysed with 1% NP-40 lysis buffer (50 mM Tris-HCl pH 7.5, 150 mM NaCl, 1 mM EDTA, 1% NP-40) containing complete protease inhibitor cocktail (Roche), passed repeatedly through a 25G needle to homogenise and clarified by centrifugation at 20,000 x g for 10 minutes at 4°C. Subsequently, clarified lysates were incubated with 10 µl of GFP-nanobody Affi-gel resin [55] for 3 hours with mixing. Beads were washed with 1% NP-40 buffer, then TBS and were eluted in SDS sample loading buffer at 95°C.

### Western blotting

Cell lysates and immunoprecipitations were resolved using precast Novex 4-12% Bis-Tris Midi Protein Gels (Thermo Fisher Scientific) and transferred to methanol-activated Immobilon-P PVDF Membranes (Millipore) using a wet transfer system. Membranes were blocked with 5% BSA (Sigma) or 5% milk in TBS containing 1% Tween-20 and incubated with primary antibody overnight at 4°C. Membranes were subsequently probed with HRP-conjugated secondary antibody, washed and bound antibody was detected using enhanced chemiluminescence (ECL) substrate.

### Immunofluorescence

Cells were grown on sterilised coverslips and fixed in 4% formaldehyde. In the case of structured illumination microscopy experiments, cells were grown on acid-washed, high performance, No. 1.5 (170±5 µm), 18 mm square coverslips (Schott). Post-fixation cells were permeabilised in 0.2% Triton X-100 and blocked with 1% BSA. Coverslips were incubated with primary antibody and then fluorescently-labelled secondary antibodies (Molecular probes). Hoechst was used to visualise DNA and biotin with AlexaFluor®568-conjugated streptavidin (Molecular probes). Images were acquired on a Zeiss Axioimager M1, a Zeiss LSM710 confocal microscope, a Zeiss Elyra PS1 super-resolution microscope or Thermo Fisher CellInsight CX7 high-content microscope. To measure colocalisation, images from randomly selected fields were background subtracted and manually segmented before calculating the Pearson’s correlation coefficient using ImageJ and the coloc2 plugin. Alternatively, confocal images from randomly selected fields of view were automatically thresholded using the Costes et al. method [58] before calculating the Pearson’s correlation coefficient using Volocity software v6.3 (PerkinElmer). Counts of OPTN puncta were performed using the HCS Studio 3.0 software packaged with the Cell Insight CX7 Microscope and the SpotDetector V4 application. Foci-positive cells were scored manually. All statistical analysis was performed in GraphPad Prism.

### CLEM

Cells were plated on alpha-numeric gridded glass-bottom coverslips (P35G-1.5-14-C-GRID, MatTek, MA, USA) at ∼40-50% confluency and fixed with 2% formaldehyde, 2.5% glutaraldehyde and 0.1 M cacodylate buffer for 30 minutes at room temperature. The reaction was quenched with 1% sodium borohydride for 20 minutes and cells were stained with Hoechst before washing with 0.1 M cacodylate. Cells were imaged on an LSM780 confocal microscope (Zeiss) and the coordinates of cells selected for imaging were recorded. After confocal image acquisition, cells were secondarily fixed with 1% osmium tetroxide and 1.5% potassium ferrocyanide before being washed and incubated with 1% tannic acid in 0.1 M cacodylate to enhance membrane contrast. Samples were washed with dH_2_O, dehydrated through an ethanol series (70%, 90%, 100%, and absolute 100%) and infiltrated with epoxy resin (Araldite CY212 mix, Agar Scientific) mixed at 1:1 with propylene oxide for one hour, before replacement with neat Epoxy resin. Excess resin was removed from the coverslip before pre-baked resin stubs were inverted over coordinates of interest and the resin cured overnight. Stubs were removed from the coverslip by immersing the coverslip in liquid nitrogen. Areas of interest were identified by alpha-numeric coordinates and 70 nm ultrathin sections were collected using a Diatome diamond knife attached to an ultracut UCT ultramicrotome (Leica). Sections were collected onto piloform-coated slot grids, stained with lead citrate and imaged on a FEI Tecnai Spirit transmission electron microscope at an operating voltage of 80kV.

### NF-κB luciferase assay

NF-κB luciferase reporter cells were plated onto 24-well plates and, at ∼80% confluency, were stimulated with 10 µg/ml poly(I:C) for 6 hours. Cells were washed with PBS, lysed in 100 µl Glo lysis buffer and clarified at 20,000 xg for 10 mins. Clarified supernatants were mixed 1:1 with ONE-GLO luciferase reagent and luminescence was analysed on a CLARIOstar microplate reader (BMG Labtech). To normalise the data, GFP fluorescence of the clarified supernatant was also determined using the same plate reader.

### Secretomics

RPE cells were cultured in SILAC DMEM:F12 (Thermo Fisher Scientific) supplemented 10% dialysed FBS (Gibco) and the heavy amino acids L-Arginine ^13^C_6_ ^15^N_4_ (147.5 mg/l) and L-Lysine ^13^C_6_ ^15^N_2_ (91.25 mg/l; Cambridge Isotope Laboratories), or equal amounts of light arginine and lysine (Sigma). Cells were taken through 3 passages to ensure complete labelling and plated onto 100 mm dishes. At ∼80% confluency, cells were incubated for 18 hours in the presence or absence of 10 µg/ml poly(I:C). Subsequently, cells were washed thoroughly with PBS and serum-free medium and incubated for 6 hours in 10 ml serum-free medium containing poly(I:C) or vehicle. Conditioned medium was harvested and clarified at 4000 xg at 4°C. Cell counts were used to normalise loading and equivalent volumes of heavy and light medium were pooled. The volume of the medium was reduced using low molecular weight spin concentrators (Sartorius) and the samples were resolved approximately 1.5 cm into a pre-cast 4-12% Bis-Tris polyacrylamide gel. The lanes were excised, cut into chunks and the proteins reduced, alkylated and digested in-gel. The resulting tryptic peptides were analysed by LC-MSMS using a Q Exactive coupled to an RSLCnano3000 (Thermo Scientific). Peptides were resolved on a 50 cm EASY-spray column (Thermo Scientific) with MSMS data acquired in a DDA fashion. Spectra were searched against a *Homo sapiens* Uniprot reference database in the MaxQuant proteomics software package [59]. Cysteine carbamidomethlyation set as a fixed modification and methionine oxidation and N-terminal acetylation as variable modifications. Peptide and protein false discovery rates (FDRs) were set to 0.01, the minimum peptide length was set at 7 amino acids and up to 2 missed cleavages were tolerated. Protein differential abundance was evaluated using the Limma package [60], within the R programming environment [61]. Differences in protein abundances were statistically determined using the Student’s t-test with variances moderated by Limma’s empirical Bayes method. P-values were adjusted for multiple testing by the Benjamini Hochberg method [62]. Gene ontology cellular component enrichment analysis was performed using the PANTHER online web tool [63].

### BioID proteomics

BioID experiments were performed as described previously [55]. Data was processed using the online tool at CRAPome.org using the default settings and a threshold of ≥3 FC-B was established to determine candidate OPTN interacting proteins. Data was visualised in Cytoscape and merged with protein-protein interaction data mined from MIMIx or IMEx curated databases [64–66].

## Supporting information

Figure Legends for Supplemental Figures

## Acknowledgements

We thank John Kendrick-Jones for generation of antibodies and reagents, Robin Antrobus and Yagnesh Umrania for help with the proteomics and Alexandra Davies and Paul Manna for helpful discussions and advice. This work was funded by a Wellcome Trust PhD studentship to T.O., the Isaac Newton Trust Cambridge and project grants from the BBSRC (BB/R001316/1) and Medical Research Council (MR/N000048/1) to F.B. CIMR is supported by the Wellcome Trust with a strategic award (100140) and an equipment grant (093026). AMS was supported by the Medical Research Council (MR/L000261/1), ALRR by the CAPES Foundation of the Ministry of Education of Brazil (0698130) and AG by the Umm Al Qura University, Faculty of Dentistry, Ministry of Education of Kingdom of Saudi Arabia (156780).

