## Supplemental Table 2 for "OPTN translocates to an ATG9A-positive compartment to regulate innate immune signalling and cytokine secretion"

BirA*-OPTN RPE BioID

| | | Spectral counts | | | | | | | |
|---|---|---|---|---|---|---|---|---|---|
| | | BirA*-OPTN | | | BirA* | | | | |
| Gene name | Fold change (FC-B) | A | B | C | A | B | C | D | E |
| OPTN | 354.7 | 1006 | 1051 | 1069 | 2 | 1 | 2 | 4 | 1 |
| IFT74 | 13.17 | 20 | 18 | 21 | 1 | 0 | 0 | 0 | 2 |
| DYNLL2 | 10.2 | 11 | 9 | 9 | 0 | 0 | 0 | 1 | 0 |
| TBK1 | 10.01 | 25 | 27 | 24 | 0 | 3 | 1 | 3 | 1 |
| TBC1D17 | 7.26 | 11 | 8 | 11 | 0 | 0 | 0 | 2 | 1 |
| DYNLL1 | 7.07 | 37 | 45 | 48 | 6 | 8 | 4 | 5 | 5 |
| RNF31 | 5.55 | 2 | 2 | 10 | 0 | 0 | 0 | 0 | 0 |
| DBT | 3.71 | 3 | 3 | 3 | 0 | 0 | 0 | 1 | 0 |
| AIMP1 | 3.67 | 18 | 11 | 10 | 3 | 2 | 2 | 2 | 6 |
| TBC1D15 | 3.43 | 50 | 31 | 40 | 10 | 13 | 13 | 16 | 10 |
| PPP1R37 | 3.18 | 7 | 7 | 7 | 1 | 1 | 0 | 2 | 4 |
| RPL10A | 2.63 | 2 | 4 | 4 | 0 | 1 | 0 | 1 | 1 |
| KIAA1033 | 2.61 | 4 | 2 | 2 | 0 | 0 | 1 | 0 | 1 |
| RIMS1 | 2.58 | 0 | 2 | 3 | 0 | 0 | 0 | 0 | 0 |
| RPL8 | 2.53 | 3 | 5 | 2 | 0 | 1 | 1 | 0 | 1 |
| ACACA | 2.44 | 257 | 234 | 312 | 102 | 123 | 146 | 108 | 116 |
| TPGS1 | 2.43 | 7 | 7 | 9 | 3 | 4 | 2 | 2 | 1 |
| DYNC1H1 | 2.42 | 55 | 69 | 73 | 31 | 35 | 23 | 23 | 25 |
| LRRC49 | 2.42 | 6 | 9 | 7 | 2 | 3 | 3 | 1 | 4 |
| PCCB | 2.38 | 40 | 42 | 60 | 15 | 25 | 20 | 28 | 25 |
| SHC1 | 2.35 | 2 | 4 | 5 | 0 | 1 | 1 | 2 | 1 |
| ILF3 | 2.3 | 5 | 1 | 3 | 0 | 0 | 0 | 2 | 1 |
| PCCA | 2.29 | 54 | 52 | 74 | 20 | 28 | 34 | 35 | 30 |
| TNKS | 2.27 | 1 | 1 | 1 | 0 | 0 | 0 | 0 | 0 |
| TXNL1 | 2.26 | 2 | 3 | 3 | 0 | 0 | 1 | 1 | 1 |
| RPS18 | 2.25 | 1 | 3 | 0 | 0 | 0 | 0 | 0 | 0 |
| UTS2 | 2.25 | 2 | 2 | 2 | 0 | 0 | 0 | 1 | 1 |
| RPL7 | 2.23 | 8 | 14 | 10 | 2 | 3 | 4 | 6 | 8 |
| SEC23IP | 2.21 | 15 | 12 | 18 | 6 | 6 | 7 | 10 | 8 |
| IGF2R | 2.18 | 23 | 23 | 36 | 4 | 3 | 6 | 26 | 19 |
| RPL38 | 2.17 | 2 | 2 | 2 | 0 | 1 | 0 | 0 | 1 |
| STX5 | 2.12 | 6 | 4 | 5 | 0 | 2 | 0 | 3 | 3 |
| HIST1H2AD | 2.11 | 5 | 3 | 4 | 1 | 2 | 2 | 1 | 0 |
| PTPN23 | 2.09 | 8 | 10 | 7 | 3 | 3 | 5 | 5 | 4 |
| STK39 | 2.07 | 3 | 4 | 4 | 1 | 0 | 2 | 2 | 1 |
| BAG2 | 2.03 | 1 | 1 | 2 | 0 | 0 | 0 | 1 | 0 |
| ZFYVE1 | 2.03 | 2 | 1 | 1 | 0 | 0 | 0 | 1 | 0 |
| TBC1D8 | 2.02 | 3 | 2 | 5 | 1 | 1 | 2 | 0 | 0 |
| WWOX | 2 | 0 | 1 | 2 | 0 | 0 | 0 | 0 | 0 |
| TRAPPC2;TRAPPC2B | 1.98 | 1 | 0 | 2 | 0 | 0 | 0 | 0 | 0 |
| RPL6 | 1.97 | 8 | 13 | 13 | 4 | 8 | 6 | 4 | 4 |

| | | | | | | | | | |
|---|---|---|---|---|---|---|---|---|---|
| C11orf49 | 1.94 | 7 | 4 | 2 | 1 | 3 | 0 | 2 | 1 |
| PC | 1.93 | 91 | 91 | 113 | 47 | 59 | 66 | 62 | 63 |
| DLST | 1.9 | 4 | 3 | 5 | 2 | 2 | 1 | 2 | 2 |
| MAGED2 | 1.89 | 2 | 7 | 4 | 1 | 1 | 1 | 4 | 2 |
| MCCC1 | 1.88 | 41 | 35 | 52 | 22 | 20 | 26 | 32 | 33 |
| CALD1 | 1.87 | 5 | 6 | 4 | 1 | 5 | 2 | 0 | 0 |
| RPL18 | 1.87 | 3 | 4 | 2 | 0 | 0 | 1 | 3 | 1 |
| DARS | 1.82 | 4 | 5 | 9 | 5 | 2 | 2 | 2 | 2 |
| HSPA5 | 1.82 | 77 | 83 | 83 | 36 | 53 | 63 | 34 | 43 |
| USO1 | 1.79 | 13 | 8 | 18 | 5 | 7 | 8 | 6 | 11 |
| XRCC6 | 1.79 | 9 | 0 | 2 | 0 | 0 | 1 | 2 | 0 |
| TRAPPC3 | 1.78 | 1 | 1 | 1 | 0 | 0 | 0 | 0 | 1 |
| HIST1H4A;HIST1H... | 1.75 | 51 | 12 | 17 | 10 | 17 | 15 | 8 | 9 |
| LMAN1 | 1.75 | 1 | 1 | 1 | 0 | 0 | 0 | 1 | 0 |
| CCDC132 | 1.74 | 1 | 1 | 0 | 0 | 0 | 0 | 0 | 0 |
| SPATA2L | 1.74 | 1 | 1 | 0 | 0 | 0 | 0 | 0 | 0 |
| CARD6 | 1.73 | 0 | 1 | 1 | 0 | 0 | 0 | 0 | 0 |
| HIST2H3A;HIST2H... | 1.72 | 8 | 4 | 3 | 0 | 2 | 1 | 4 | 4 |
| HUWE1 | 1.72 | 2 | 2 | 2 | 0 | 1 | 2 | 0 | 0 |
| MYL9 | 1.72 | 5 | 3 | 2 | 2 | 3 | 0 | 0 | 0 |
| RPL27 | 1.71 | 1 | 3 | 4 | 1 | 1 | 1 | 2 | 0 |
| TUBA1A;TUBA3C | 1.69 | 9 | 10 | 8 | 5 | 4 | 8 | 3 | 5 |
| SNAP29 | 1.68 | 5 | 5 | 5 | 2 | 2 | 3 | 5 | 3 |
| CYLD | 1.67 | 11 | 9 | 8 | 4 | 5 | 3 | 9 | 8 |
| DIABLO | 1.67 | 1 | 1 | 1 | 0 | 1 | 0 | 0 | 0 |
| HNRNPR | 1.67 | 1 | 1 | 2 | 0 | 0 | 0 | 1 | 1 |
| HIST1H1B | 1.66 | 3 | 7 | 8 | 3 | 4 | 3 | 4 | 3 |
| AZI2 | 1.65 | 1 | 2 | 1 | 0 | 0 | 0 | 2 | 0 |
| SLC16A1 | 1.63 | 1 | 2 | 1 | 0 | 0 | 1 | 0 | 1 |
| TAB2 | 1.63 | 2 | 4 | 2 | 0 | 1 | 1 | 3 | 1 |
| PRDX6 | 1.62 | 7 | 8 | 8 | 2 | 4 | 5 | 2 | 9 |
| DNAJC13 | 1.61 | 1 | 4 | 3 | 1 | 0 | 0 | 0 | 4 |
| HSPB1 | 1.61 | 8 | 9 | 10 | 5 | 6 | 7 | 4 | 4 |
| RPL29 | 1.61 | 3 | 2 | 4 | 2 | 2 | 1 | 1 | 1 |
| MRPL4 | 1.6 | 1 | 1 | 1 | 1 | 0 | 0 | 0 | 0 |
| RPLP0;RPLP0P6 | 1.58 | 4 | 6 | 6 | 3 | 2 | 1 | 5 | 4 |
| TMSB10 | 1.58 | 2 | 4 | 3 | 0 | 3 | 1 | 0 | 2 |
| PHB | 1.57 | 9 | 7 | 9 | 4 | 8 | 2 | 6 | 5 |
| CAND1 | 1.56 | 7 | 4 | 6 | 4 | 3 | 4 | 1 | 1 |
| LPP | 1.56 | 4 | 9 | 10 | 4 | 2 | 4 | 8 | 4 |
| SSR4 | 1.56 | 3 | 4 | 2 | 1 | 2 | 2 | 2 | 2 |
| CLINT1 | 1.55 | 14 | 14 | 25 | 11 | 12 | 13 | 14 | 16 |
| YWHAG | 1.55 | 4 | 6 | 4 | 2 | 3 | 0 | 5 | 3 |
| PRDX2 | 1.54 | 3 | 3 | 4 | 2 | 2 | 2 | 2 | 3 |
| RPLP2 | 1.54 | 6 | 4 | 5 | 5 | 2 | 2 | 3 | 2 |

| | | | | | | | | | |
|---|---|---|---|---|---|---|---|---|---|
| SH3PXD2A;SH3PXD2B | 1.54 | 0 | 1 | 2 | 0 | 0 | 0 | 1 | 0 |
| NSF | 1.53 | 6 | 5 | 5 | 2 | 5 | 3 | 4 | 4 |
| PSMA1 | 1.53 | 1 | 2 | 2 | 0 | 1 | 1 | 1 | 1 |
| AASDHPPT | 1.52 | 2 | 0 | 0 | 0 | 0 | 0 | 0 | 0 |
| ASPM | 1.52 | 2 | 0 | 0 | 0 | 0 | 0 | 0 | 0 |
| CDC2;CDK1 | 1.51 | 3 | 2 | 2 | 0 | 1 | 2 | 0 | 2 |
| CLIP2 | 1.51 | 0 | 0 | 2 | 0 | 0 | 0 | 0 | 0 |
| KIAA1217 | 1.51 | 4 | 4 | 5 | 0 | 2 | 0 | 4 | 5 |
| MKL2 | 1.5 | 17 | 20 | 27 | 7 | 7 | 4 | 30 | 20 |
| ZWILCH | 1.49 | 0 | 1 | 2 | 0 | 0 | 1 | 0 | 0 |
| CCT5 | 1.48 | 24 | 28 | 32 | 19 | 22 | 22 | 21 | 16 |
| ELP6 | 1.48 | 9 | 6 | 10 | 5 | 7 | 6 | 6 | 6 |
| MCCC2 | 1.48 | 28 | 29 | 38 | 19 | 22 | 26 | 31 | 30 |
| TUBB3 | 1.47 | 4 | 3 | 6 | 1 | 0 | 0 | 5 | 5 |
| RPS27L;RPS27 | 1.46 | 1 | 3 | 4 | 0 | 5 | 0 | 0 | 0 |
| TPD52L2 | 1.46 | 4 | 4 | 8 | 3 | 4 | 4 | 2 | 2 |
| VCPIP1 | 1.46 | 29 | 28 | 41 | 23 | 26 | 25 | 28 | 32 |
| AHNAK | 1.45 | 863 | 926 | 974 | 604 | 776 | 792 | 579 | 597 |
| DPM1 | 1.45 | 2 | 1 | 3 | 0 | 1 | 2 | 0 | 1 |
| KIAA0196 | 1.45 | 3 | 3 | 6 | 3 | 2 | 3 | 1 | 1 |
| NRBP1 | 1.44 | 2 | 4 | 6 | 2 | 3 | 3 | 1 | 1 |
| RPL3 | 1.44 | 8 | 9 | 5 | 1 | 3 | 0 | 10 | 7 |
| RPS27A;UBB;UBC;UBA52 | 1.44 | 12 | 10 | 13 | 1 | 3 | 2 | 15 | 16 |
| CCT4 | 1.43 | 21 | 27 | 29 | 20 | 19 | 20 | 23 | 19 |
| LARS | 1.43 | 2 | 3 | 2 | 0 | 0 | 0 | 4 | 2 |
| PLA2G4A | 1.43 | 9 | 12 | 13 | 6 | 9 | 9 | 9 | 12 |
| ATP5A1 | 1.41 | 19 | 19 | 27 | 16 | 12 | 20 | 16 | 19 |
| DVL3 | 1.41 | 8 | 9 | 8 | 4 | 6 | 9 | 5 | 7 |
| PSMD6 | 1.41 | 1 | 1 | 3 | 0 | 2 | 1 | 0 | 0 |
| HAUS6 | 1.4 | 4 | 8 | 7 | 2 | 5 | 1 | 7 | 5 |
| DPYSL3 | 1.39 | 30 | 32 | 41 | 24 | 25 | 34 | 24 | 25 |
| UBR4 | 1.39 | 1 | 1 | 1 | 0 | 1 | 0 | 0 | 1 |
| AHSA1 | 1.38 | 5 | 7 | 8 | 4 | 6 | 2 | 4 | 7 |
| SEC61A1 | 1.38 | 6 | 3 | 4 | 2 | 3 | 4 | 3 | 4 |
| SLC25A6 | 1.37 | 18 | 26 | 29 | 15 | 24 | 20 | 11 | 14 |
| TTK | 1.37 | 1 | 1 | 2 | 0 | 0 | 1 | 1 | 1 |
| DYNC1LI1 | 1.36 | 0 | 1 | 1 | 0 | 0 | 0 | 0 | 1 |
| HAUS3 | 1.36 | 0 | 2 | 2 | 0 | 2 | 0 | 0 | 0 |
| HSPA8 | 1.36 | 71 | 67 | 70 | 63 | 53 | 55 | 54 | 58 |
| RPL12 | 1.36 | 3 | 4 | 3 | 0 | 0 | 0 | 4 | 6 |
| CCT7 | 1.35 | 25 | 28 | 43 | 25 | 28 | 24 | 14 | 13 |
| RPS3 | 1.35 | 17 | 22 | 24 | 14 | 21 | 17 | 13 | 18 |
| IRS2 | 1.34 | 1 | 0 | 1 | 0 | 0 | 0 | 0 | 1 |
| NME1-NME2;NME2 | 1.34 | 10 | 7 | 6 | 4 | 6 | 8 | 6 | 7 |
| SPG20 | 1.34 | 12 | 11 | 16 | 7 | 3 | 12 | 16 | 10 |

| | | | | | | | | | |
|---|---|---|---|---|---|---|---|---|---|
| TLN2 | 1.34 | 1 | 0 | 1 | 0 | 0 | 0 | 0 | 1 |
| ARL3 | 1.33 | 0 | 1 | 1 | 0 | 0 | 0 | 1 | 0 |
| CCT6A | 1.33 | 22 | 30 | 28 | 19 | 26 | 22 | 17 | 16 |
| COPZ2 | 1.33 | 0 | 1 | 0 | 0 | 0 | 0 | 0 | 0 |
| EIF4E2 | 1.33 | 12 | 11 | 11 | 4 | 8 | 5 | 13 | 14 |
| HSPA1B;HSPA1A | 1.33 | 22 | 20 | 27 | 19 | 21 | 14 | 20 | 19 |
| LMCD1 | 1.33 | 0 | 1 | 0 | 0 | 0 | 0 | 0 | 0 |
| LRRC40 | 1.33 | 0 | 1 | 0 | 0 | 0 | 0 | 0 | 0 |
| MRPL2 | 1.33 | 0 | 1 | 0 | 0 | 0 | 0 | 0 | 0 |
| NUDT5 | 1.33 | 7 | 7 | 5 | 1 | 7 | 5 | 5 | 2 |
| SUMO1 | 1.33 | 1 | 1 | 1 | 0 | 2 | 0 | 0 | 0 |
| WASH6P | 1.33 | 0 | 1 | 1 | 0 | 0 | 0 | 1 | 0 |
| XPOT | 1.33 | 0 | 1 | 1 | 0 | 0 | 0 | 1 | 0 |
| XRCC5 | 1.33 | 7 | 1 | 2 | 1 | 3 | 1 | 2 | 2 |
| ERLIN1 | 1.32 | 2 | 2 | 2 | 1 | 2 | 0 | 1 | 2 |
| RPS2 | 1.32 | 16 | 12 | 10 | 15 | 7 | 7 | 9 | 10 |
| TUBB4B | 1.32 | 60 | 47 | 58 | 46 | 46 | 47 | 47 | 47 |
| ARPC4 | 1.31 | 7 | 6 | 6 | 3 | 7 | 5 | 3 | 6 |
| ATP6V1H | 1.31 | 0 | 0 | 1 | 0 | 0 | 0 | 0 | 0 |
| CAMK2G | 1.31 | 0 | 0 | 1 | 0 | 0 | 0 | 0 | 0 |
| CFAP54 | 1.31 | 1 | 0 | 0 | 0 | 0 | 0 | 0 | 0 |
| COMMD4 | 1.31 | 0 | 0 | 1 | 0 | 0 | 0 | 0 | 0 |
| FLRT2 | 1.31 | 0 | 0 | 1 | 0 | 0 | 0 | 0 | 0 |
| ILVBL | 1.31 | 0 | 0 | 1 | 0 | 0 | 0 | 0 | 0 |
| KIF11 | 1.31 | 0 | 0 | 1 | 0 | 0 | 0 | 0 | 0 |
| KSR1 | 1.31 | 0 | 0 | 1 | 0 | 0 | 0 | 0 | 0 |
| MYH11 | 1.31 | 0 | 0 | 1 | 0 | 0 | 0 | 0 | 0 |
| MYL1;MYL3 | 1.31 | 1 | 0 | 0 | 0 | 0 | 0 | 0 | 0 |
| OSTC | 1.31 | 2 | 1 | 1 | 0 | 1 | 1 | 1 | 1 |
| RPL22L1 | 1.31 | 1 | 0 | 0 | 0 | 0 | 0 | 0 | 0 |
| RPL39P5;RPL39 | 1.31 | 0 | 0 | 1 | 0 | 0 | 0 | 0 | 0 |
| SLC25A4 | 1.31 | 1 | 0 | 0 | 0 | 0 | 0 | 0 | 0 |
| WDR20 | 1.31 | 0 | 0 | 1 | 0 | 0 | 0 | 0 | 0 |
| CKAP5 | 1.3 | 21 | 31 | 25 | 16 | 19 | 19 | 34 | 28 |
| CNGB3 | 1.29 | 1 | 1 | 0 | 0 | 0 | 1 | 0 | 0 |
| HIST1H2BI;HIST1H2BN;HIST1H2BL | 1.29 | 10 | 6 | 6 | 4 | 7 | 5 | 7 | 8 |
| SRP14 | 1.29 | 2 | 1 | 2 | 0 | 2 | 1 | 1 | 0 |
| VPS13C | 1.29 | 0 | 0 | 3 | 0 | 0 | 0 | 1 | 0 |
| CTSC | 1.28 | 0 | 1 | 1 | 0 | 1 | 0 | 0 | 0 |
| HIST2H2BE;HIST1H2BB;HIST1H2BO | 1.28 | 6 | 2 | 2 | 1 | 3 | 3 | 2 | 1 |
| HSPA6;HSPA7 | 1.28 | 0 | 2 | 1 | 0 | 0 | 0 | 1 | 1 |
| MAST4 | 1.28 | 0 | 1 | 1 | 0 | 0 | 1 | 0 | 0 |
| OSBPL9 | 1.28 | 1 | 1 | 1 | 1 | 0 | 1 | 0 | 0 |
| RPL30 | 1.28 | 4 | 4 | 6 | 3 | 4 | 0 | 2 | 6 |
| CLTC | 1.27 | 18 | 30 | 29 | 16 | 27 | 24 | 13 | 13 |

| | | | | | | | | | |
|---|---|---|---|---|---|---|---|---|---|
| SNTB2 | 1.27 | 1 | 2 | 3 | 1 | 1 | 2 | 1 | 2 |
| SRP72 | 1.27 | 3 | 8 | 5 | 3 | 5 | 5 | 1 | 1 |
| TGM2 | 1.27 | 9 | 8 | 15 | 8 | 6 | 13 | 7 | 7 |
| VIPAS39 | 1.27 | 2 | 2 | 1 | 0 | 1 | 3 | 0 | 0 |
| AP3S1 | 1.26 | 1 | 0 | 2 | 0 | 0 | 0 | 1 | 1 |
| BRE | 1.26 | 1 | 1 | 3 | 0 | 1 | 1 | 2 | 1 |
| ERCC6L | 1.26 | 4 | 7 | 9 | 2 | 2 | 3 | 9 | 9 |
| GAPVD1 | 1.26 | 8 | 8 | 8 | 5 | 6 | 7 | 11 | 6 |
| PASK | 1.26 | 1 | 0 | 1 | 0 | 1 | 0 | 0 | 0 |
| PRKAB1 | 1.26 | 1 | 0 | 1 | 0 | 1 | 0 | 0 | 0 |
| RPL13A | 1.26 | 2 | 5 | 3 | 1 | 0 | 1 | 3 | 6 |
| S100A11 | 1.26 | 2 | 3 | 3 | 2 | 2 | 1 | 3 | 2 |
| AHNAK2 | 1.25 | 182 | 168 | 202 | 121 | 151 | 151 | 223 | 242 |
| FLNB | 1.25 | 100 | 116 | 126 | 86 | 103 | 111 | 125 | 142 |
| HAUS4 | 1.25 | 0 | 1 | 2 | 0 | 0 | 0 | 2 | 0 |
| PSMD11 | 1.25 | 2 | 6 | 3 | 1 | 4 | 4 | 0 | 1 |
| SMAP1 | 1.25 | 3 | 5 | 6 | 2 | 6 | 4 | 2 | 1 |
| KLC1 | 1.24 | 19 | 15 | 16 | 14 | 17 | 14 | 16 | 13 |
| MTAP | 1.24 | 1 | 2 | 3 | 1 | 2 | 1 | 2 | 1 |
| NUP62 | 1.24 | 3 | 2 | 3 | 3 | 0 | 1 | 3 | 1 |
| PATL1 | 1.24 | 3 | 0 | 0 | 0 | 1 | 0 | 0 | 0 |
| RPS6 | 1.24 | 10 | 12 | 11 | 7 | 11 | 11 | 11 | 11 |
| TCP1 | 1.24 | 26 | 30 | 37 | 22 | 33 | 29 | 20 | 22 |
| ANXA2;ANXA2P2 | 1.23 | 61 | 69 | 74 | 57 | 63 | 67 | 36 | 41 |
| CD2AP | 1.23 | 41 | 44 | 45 | 44 | 38 | 36 | 36 | 39 |
| ENSA | 1.23 | 3 | 2 | 2 | 1 | 2 | 3 | 0 | 0 |
| NRBF2 | 1.23 | 2 | 7 | 8 | 4 | 1 | 5 | 5 | 4 |
| PARP1 | 1.23 | 5 | 0 | 0 | 0 | 0 | 0 | 1 | 1 |
| EIF4E | 1.22 | 8 | 8 | 7 | 6 | 7 | 8 | 7 | 4 |
| RAPGEF2 | 1.22 | 1 | 0 | 2 | 0 | 1 | 0 | 0 | 1 |
| ROCK2 | 1.22 | 2 | 5 | 3 | 2 | 3 | 2 | 4 | 2 |
| FLNA | 1.21 | 786 | 897 | 865 | 626 | 852 | 915 | 559 | 480 |
| PRKDC | 1.21 | 20 | 22 | 27 | 16 | 30 | 18 | 16 | 22 |
| HIST1H1E | 1.2 | 4 | 3 | 3 | 2 | 4 | 3 | 2 | 1 |
| XRN1 | 1.2 | 40 | 43 | 49 | 36 | 40 | 45 | 52 | 53 |
| LDHB | 1.19 | 1 | 1 | 2 | 0 | 1 | 0 | 1 | 2 |
| RPLP1 | 1.19 | 3 | 1 | 2 | 1 | 3 | 1 | 1 | 0 |
| CLIC1 | 1.18 | 5 | 3 | 5 | 5 | 2 | 3 | 4 | 4 |
| HIST1H1C;HIST1H1D | 1.18 | 12 | 10 | 11 | 12 | 8 | 5 | 11 | 12 |
| TRIP6 | 1.18 | 2 | 1 | 1 | 0 | 1 | 0 | 2 | 1 |
| CRYBG3 | 1.17 | 0 | 2 | 2 | 1 | 0 | 0 | 1 | 1 |
| GCN1L1 | 1.17 | 27 | 28 | 32 | 24 | 34 | 26 | 21 | 19 |
| KIF13A | 1.17 | 3 | 7 | 2 | 3 | 4 | 3 | 2 | 2 |
| RIPK1 | 1.17 | 2 | 0 | 0 | 0 | 0 | 0 | 1 | 0 |
| YBX1 | 1.17 | 5 | 5 | 6 | 2 | 6 | 2 | 7 | 5 |

| | | | | | | | | | |
|---|---|---|---|---|---|---|---|---|---|
| ARHGAP1 | 1.16 | 3 | 5 | 4 | 0 | 2 | 3 | 4 | 7 |
| DENND4C | 1.16 | 9 | 8 | 12 | 8 | 8 | 7 | 12 | 12 |
| PDLIM5 | 1.16 | 34 | 54 | 50 | 36 | 44 | 53 | 41 | 46 |
| RPL18A | 1.16 | 6 | 5 | 4 | 3 | 3 | 2 | 7 | 7 |
| RPS7 | 1.16 | 7 | 7 | 6 | 6 | 7 | 6 | 4 | 4 |
| SLC2A1 | 1.16 | 5 | 6 | 9 | 7 | 4 | 6 | 5 | 7 |
| ARPIN | 1.15 | 4 | 2 | 8 | 3 | 3 | 2 | 5 | 6 |
| PIK3C3 | 1.15 | 1 | 1 | 2 | 1 | 0 | 0 | 2 | 1 |
| PRDX1 | 1.15 | 9 | 10 | 9 | 6 | 6 | 10 | 8 | 15 |
| CCT2 | 1.14 | 22 | 24 | 33 | 19 | 30 | 29 | 18 | 23 |
| CRK | 1.14 | 4 | 10 | 13 | 2 | 9 | 7 | 10 | 10 |
| CTTN | 1.14 | 42 | 49 | 50 | 29 | 50 | 49 | 52 | 63 |
| GANAB | 1.14 | 1 | 1 | 1 | 0 | 2 | 0 | 0 | 1 |
| IFIT5 | 1.14 | 16 | 18 | 17 | 12 | 20 | 19 | 14 | 12 |
| PHGDH | 1.14 | 2 | 3 | 4 | 2 | 1 | 4 | 3 | 1 |
| ENO1 | 1.13 | 18 | 8 | 14 | 8 | 10 | 9 | 17 | 20 |
| NT5C2 | 1.13 | 3 | 3 | 6 | 4 | 3 | 4 | 2 | 1 |
| PYGB | 1.13 | 8 | 9 | 13 | 6 | 10 | 13 | 8 | 10 |
| SLC25A3 | 1.13 | 4 | 3 | 7 | 3 | 4 | 4 | 5 | 7 |
| TUBB | 1.13 | 13 | 13 | 15 | 11 | 16 | 14 | 8 | 10 |
| RPL32 | 1.12 | 2 | 2 | 2 | 2 | 2 | 0 | 2 | 1 |
| STAT2 | 1.12 | 6 | 6 | 8 | 5 | 8 | 7 | 6 | 3 |
| TUBA1C | 1.12 | 2 | 1 | 2 | 0 | 0 | 0 | 3 | 3 |
| SEPT2 | 1.11 | 14 | 14 | 15 | 11 | 16 | 12 | 18 | 19 |
| CORO1B | 1.11 | 53 | 55 | 56 | 55 | 53 | 59 | 31 | 32 |
| GBF1 | 1.11 | 1 | 2 | 5 | 1 | 3 | 1 | 2 | 3 |
| LIMS1 | 1.11 | 1 | 1 | 0 | 0 | 0 | 0 | 1 | 1 |
| SYNJ2 | 1.11 | 37 | 34 | 41 | 35 | 37 | 39 | 47 | 40 |
| TNS3 | 1.11 | 13 | 17 | 20 | 11 | 18 | 16 | 22 | 21 |
| CRKL | 1.1 | 5 | 5 | 9 | 5 | 5 | 5 | 7 | 10 |
| HAUS7 | 1.1 | 0 | 1 | 1 | 0 | 0 | 0 | 1 | 1 |
| PHB2 | 1.1 | 6 | 8 | 9 | 7 | 9 | 7 | 8 | 8 |
| RPL4 | 1.1 | 9 | 9 | 10 | 4 | 5 | 5 | 18 | 13 |
| ARHGEF5 | 1.09 | 1 | 1 | 1 | 1 | 1 | 0 | 1 | 1 |
| S100A6 | 1.09 | 2 | 2 | 2 | 2 | 2 | 2 | 1 | 1 |
| SLC25A11 | 1.09 | 2 | 5 | 4 | 4 | 3 | 2 | 4 | 3 |
| VBP1 | 1.09 | 1 | 0 | 1 | 0 | 0 | 0 | 1 | 1 |
| YIPF5 | 1.09 | 1 | 0 | 1 | 0 | 0 | 0 | 1 | 1 |
| CAP1 | 1.08 | 11 | 20 | 19 | 16 | 18 | 16 | 19 | 19 |
| DTD1 | 1.08 | 0 | 1 | 1 | 0 | 0 | 0 | 2 | 0 |
| GAK | 1.08 | 3 | 4 | 3 | 3 | 2 | 3 | 5 | 1 |
| GART | 1.08 | 7 | 6 | 10 | 9 | 5 | 7 | 8 | 9 |
| MOB2 | 1.08 | 2 | 2 | 2 | 1 | 1 | 3 | 1 | 3 |
| PDLIM4 | 1.08 | 10 | 12 | 16 | 7 | 14 | 12 | 18 | 15 |
| TUBA1B | 1.08 | 69 | 72 | 72 | 68 | 76 | 78 | 49 | 53 |

| | | | | | | | | | |
|---|---|---|---|---|---|---|---|---|---|
| CPNE3;CPNE8;CPNE2 | 1.07 | 1 | 1 | 1 | 1 | 1 | 1 | 0 | 1 |
| H3F3B;H3F3A;H3F3C | 1.07 | 2 | 0 | 0 | 1 | 0 | 0 | 0 | 0 |
| HSPA4 | 1.07 | 4 | 3 | 6 | 3 | 4 | 5 | 5 | 6 |
| LRCH3 | 1.07 | 1 | 1 | 1 | 1 | 1 | 1 | 1 | 0 |
| MAPKAP1 | 1.07 | 1 | 1 | 1 | 1 | 1 | 1 | 0 | 0 |
| NME7 | 1.07 | 2 | 0 | 0 | 1 | 0 | 0 | 0 | 0 |
| RPS10 | 1.07 | 1 | 1 | 1 | 1 | 1 | 1 | 1 | 0 |
| SYNJ1 | 1.07 | 4 | 4 | 3 | 1 | 1 | 4 | 6 | 4 |
| ARF4 | 1.06 | 9 | 9 | 14 | 11 | 10 | 12 | 6 | 7 |
| EEF1E1 | 1.06 | 2 | 1 | 1 | 1 | 2 | 1 | 0 | 1 |
| TXNDC5 | 1.06 | 0 | 0 | 2 | 1 | 0 | 0 | 0 | 0 |
| CCT3 | 1.05 | 17 | 17 | 18 | 14 | 15 | 24 | 22 | 22 |
| GEN1 | 1.05 | 1 | 1 | 1 | 0 | 0 | 0 | 3 | 1 |
| KIF13B | 1.05 | 0 | 1 | 1 | 0 | 0 | 1 | 1 | 0 |
| PSMC2 | 1.05 | 3 | 7 | 4 | 2 | 5 | 5 | 6 | 3 |
| STK38 | 1.05 | 9 | 10 | 9 | 7 | 9 | 9 | 12 | 16 |
| TACC2 | 1.05 | 2 | 2 | 3 | 1 | 1 | 2 | 5 | 2 |
| TANC1 | 1.05 | 4 | 13 | 11 | 5 | 6 | 7 | 10 | 17 |
| WASF2 | 1.05 | 5 | 7 | 5 | 5 | 7 | 6 | 3 | 1 |
| ACACB | 1.04 | 0 | 1 | 0 | 0 | 0 | 0 | 0 | 1 |
| LRBA | 1.04 | 18 | 19 | 28 | 17 | 22 | 21 | 36 | 23 |
| RAN | 1.04 | 13 | 10 | 10 | 14 | 9 | 12 | 10 | 11 |
| VKORC1 | 1.04 | 0 | 1 | 0 | 0 | 0 | 0 | 0 | 1 |
| YWHAE | 1.04 | 10 | 8 | 8 | 8 | 6 | 11 | 11 | 10 |
| ZYX | 1.04 | 22 | 19 | 23 | 19 | 24 | 27 | 23 | 25 |
| ANXA1 | 1.03 | 14 | 13 | 16 | 14 | 16 | 17 | 12 | 11 |
| HIST3H2A;HIST1H2AB | 1.03 | 16 | 6 | 7 | 8 | 11 | 10 | 6 | 6 |
| HMGN2 | 1.03 | 1 | 0 | 0 | 0 | 0 | 0 | 0 | 1 |
| IDH1 | 1.03 | 1 | 2 | 2 | 3 | 1 | 1 | 1 | 1 |
| MB21D2 | 1.03 | 1 | 0 | 0 | 0 | 0 | 0 | 0 | 1 |
| NEK9 | 1.03 | 19 | 27 | 28 | 22 | 30 | 22 | 27 | 36 |
| PSMA3 | 1.03 | 1 | 0 | 0 | 0 | 0 | 0 | 0 | 1 |
| RELA | 1.03 | 1 | 3 | 3 | 2 | 1 | 2 | 2 | 4 |
| RPS13 | 1.03 | 1 | 2 | 2 | 1 | 2 | 1 | 0 | 3 |
| CCDC25 | 1.02 | 1 | 1 | 1 | 0 | 1 | 0 | 2 | 1 |
| CDC16 | 1.02 | 0 | 1 | 0 | 0 | 0 | 0 | 1 | 0 |
| DPYSL2 | 1.02 | 78 | 68 | 74 | 75 | 78 | 91 | 42 | 42 |
| FSIP2 | 1.02 | 0 | 0 | 1 | 0 | 0 | 0 | 0 | 1 |
| OGDH | 1.02 | 1 | 1 | 0 | 0 | 1 | 1 | 0 | 0 |
| OSBPL11 | 1.02 | 2 | 4 | 4 | 2 | 2 | 2 | 4 | 7 |
| PSMB6 | 1.02 | 0 | 1 | 0 | 0 | 0 | 0 | 1 | 0 |
| SNRPD1 | 1.02 | 0 | 1 | 0 | 0 | 0 | 0 | 1 | 0 |
| STAMBP | 1.02 | 0 | 1 | 0 | 0 | 0 | 0 | 1 | 0 |
| TLN1 | 1.02 | 152 | 162 | 179 | 155 | 184 | 212 | 153 | 165 |
| TTC1 | 1.02 | 0 | 1 | 1 | 1 | 0 | 0 | 0 | 1 |

| | | | | | | | | | |
|---|---|---|---|---|---|---|---|---|---|
| CCT8 | 1.01 | 33 | 34 | 39 | 34 | 40 | 45 | 44 | 48 |
| DERL1 | 1.01 | 1 | 0 | 0 | 0 | 0 | 0 | 1 | 0 |
| FASN | 1.01 | 281 | 260 | 287 | 300 | 307 | 314 | 170 | 177 |
| MTMR9 | 1.01 | 1 | 0 | 0 | 0 | 0 | 0 | 1 | 0 |
| NBEA | 1.01 | 1 | 3 | 8 | 1 | 2 | 8 | 0 | 0 |
| PLEC | 1.01 | 29 | 41 | 52 | 17 | 27 | 30 | 65 | 72 |
| PSMB2 | 1.01 | 1 | 0 | 0 | 0 | 0 | 0 | 1 | 0 |
| SMG8 | 1.01 | 1 | 0 | 0 | 0 | 0 | 0 | 1 | 0 |
| UBTF | 1.01 | 1 | 0 | 0 | 0 | 0 | 0 | 1 | 0 |
| USP15 | 1.01 | 24 | 20 | 21 | 20 | 27 | 25 | 26 | 32 |
| YWHAH | 1.01 | 2 | 2 | 1 | 0 | 0 | 0 | 5 | 2 |
| SEPT9 | 1 | 21 | 25 | 26 | 23 | 26 | 30 | 34 | 35 |
| CHMP5 | 1 | 4 | 3 | 3 | 3 | 3 | 5 | 4 | 1 |
| COPZ1 | 1 | 1 | 1 | 1 | 0 | 0 | 2 | 1 | 1 |
| FAM114A1 | 1 | 9 | 14 | 12 | 11 | 10 | 12 | 16 | 19 |
| GPHN | 1 | 0 | 0 | 1 | 0 | 0 | 0 | 1 | 0 |
| NEK7 | 1 | 0 | 0 | 1 | 0 | 0 | 0 | 1 | 0 |
| PLOD2 | 1 | 0 | 0 | 1 | 0 | 0 | 0 | 1 | 0 |
| SPATA2 | 1 | 0 | 0 | 1 | 0 | 0 | 0 | 1 | 0 |
| AAK1 | 0.99 | 12 | 14 | 14 | 13 | 14 | 14 | 16 | 24 |
| DFNA5 | 0.99 | 8 | 8 | 12 | 10 | 11 | 10 | 12 | 12 |
| DNMBP | 0.99 | 32 | 33 | 37 | 35 | 38 | 44 | 41 | 41 |
| EIF4A1 | 0.99 | 10 | 7 | 10 | 6 | 11 | 3 | 13 | 13 |
| PABPC4 | 0.99 | 1 | 0 | 2 | 0 | 1 | 1 | 1 | 1 |
| PARVA | 0.99 | 2 | 3 | 2 | 2 | 3 | 3 | 1 | 2 |
| PSME2 | 0.99 | 2 | 2 | 3 | 2 | 3 | 3 | 1 | 3 |
| AMPD2 | 0.98 | 1 | 1 | 1 | 1 | 1 | 1 | 0 | 2 |
| AP3B1 | 0.98 | 6 | 4 | 6 | 6 | 7 | 4 | 6 | 6 |
| CDR2L | 0.98 | 0 | 1 | 0 | 0 | 0 | 1 | 0 | 0 |
| LPCAT2 | 0.98 | 0 | 1 | 2 | 1 | 0 | 1 | 0 | 1 |
| LRP1 | 0.98 | 0 | 1 | 0 | 0 | 1 | 0 | 0 | 0 |
| PCBP1 | 0.98 | 9 | 7 | 10 | 6 | 7 | 4 | 14 | 16 |
| PLS1 | 0.98 | 2 | 0 | 0 | 0 | 0 | 0 | 0 | 2 |
| POLR2H | 0.98 | 1 | 1 | 0 | 1 | 1 | 0 | 0 | 0 |
| PSMA5 | 0.98 | 0 | 1 | 0 | 0 | 1 | 0 | 0 | 0 |
| RAD50 | 0.98 | 2 | 1 | 1 | 1 | 2 | 1 | 1 | 2 |
| SERPINH1 | 0.98 | 7 | 6 | 4 | 4 | 6 | 5 | 4 | 12 |
| SNAPIN | 0.98 | 0 | 1 | 0 | 0 | 1 | 0 | 0 | 0 |
| TMEM33 | 0.98 | 1 | 1 | 1 | 1 | 1 | 1 | 1 | 2 |
| VIM | 0.98 | 90 | 121 | 113 | 111 | 139 | 123 | 85 | 87 |
| ASPSCR1 | 0.97 | 1 | 1 | 2 | 0 | 2 | 2 | 1 | 0 |
| CCP110 | 0.97 | 0 | 0 | 1 | 0 | 0 | 1 | 0 | 0 |
| CNN2 | 0.97 | 7 | 9 | 5 | 1 | 8 | 9 | 10 | 10 |
| COL12A1 | 0.97 | 1 | 0 | 0 | 0 | 1 | 0 | 0 | 0 |
| H2AFY | 0.97 | 1 | 0 | 0 | 0 | 1 | 0 | 0 | 0 |

| | | | | | | | | | |
|---|---|---|---|---|---|---|---|---|---|
| HOOK3 | 0.97 | 0 | 0 | 1 | 0 | 0 | 1 | 0 | 0 |
| LOXL2 | 0.97 | 1 | 0 | 0 | 0 | 1 | 0 | 0 | 0 |
| PSMB5 | 0.97 | 0 | 0 | 1 | 0 | 0 | 1 | 0 | 0 |
| SFXN3 | 0.97 | 1 | 2 | 2 | 0 | 2 | 1 | 3 | 2 |
| SUGT1 | 0.97 | 4 | 4 | 4 | 5 | 4 | 5 | 4 | 5 |
| GTPBP1 | 0.96 | 0 | 0 | 1 | 0 | 1 | 0 | 0 | 0 |
| HSPH1 | 0.96 | 10 | 8 | 10 | 5 | 8 | 9 | 15 | 17 |
| PPIA | 0.96 | 1 | 1 | 2 | 1 | 2 | 0 | 2 | 1 |
| PSMC6 | 0.96 | 2 | 3 | 2 | 2 | 3 | 0 | 4 | 3 |
| PSMD2 | 0.96 | 12 | 9 | 9 | 10 | 10 | 6 | 15 | 16 |
| SEC23A | 0.96 | 6 | 6 | 8 | 5 | 7 | 11 | 5 | 9 |
| HN1L | 0.95 | 3 | 5 | 3 | 2 | 3 | 4 | 6 | 6 |
| HNRNPUL1 | 0.95 | 2 | 3 | 1 | 3 | 2 | 1 | 2 | 1 |
| MARCKS | 0.95 | 6 | 2 | 4 | 0 | 2 | 1 | 7 | 8 |
| PPP1R13L | 0.95 | 6 | 3 | 5 | 6 | 3 | 7 | 3 | 4 |
| RPL31 | 0.95 | 1 | 5 | 1 | 2 | 1 | 1 | 3 | 3 |
| RPL7A | 0.95 | 3 | 4 | 5 | 0 | 3 | 3 | 6 | 9 |
| ACTR1A | 0.94 | 11 | 12 | 5 | 13 | 7 | 9 | 10 | 12 |
| AIMP2 | 0.94 | 1 | 3 | 3 | 1 | 2 | 1 | 4 | 4 |
| EHD1 | 0.94 | 1 | 2 | 3 | 2 | 3 | 2 | 2 | 1 |
| FRYL | 0.94 | 0 | 2 | 2 | 1 | 2 | 1 | 1 | 0 |
| H2AFV;H2AFZ | 0.94 | 3 | 0 | 1 | 1 | 1 | 1 | 1 | 2 |
| RHEB | 0.94 | 2 | 1 | 2 | 2 | 2 | 2 | 1 | 1 |
| SCYL3 | 0.94 | 0 | 2 | 2 | 1 | 1 | 2 | 1 | 1 |
| UGP2 | 0.94 | 0 | 3 | 2 | 2 | 1 | 1 | 1 | 2 |
| VDAC1 | 0.94 | 3 | 3 | 4 | 4 | 4 | 4 | 1 | 4 |
| ADD1 | 0.93 | 4 | 8 | 12 | 8 | 6 | 10 | 11 | 10 |
| DTX3L | 0.93 | 0 | 0 | 2 | 0 | 0 | 1 | 0 | 1 |
| EIF4G1 | 0.93 | 36 | 39 | 42 | 42 | 50 | 45 | 59 | 55 |
| HNRNPAB | 0.93 | 1 | 2 | 1 | 1 | 3 | 1 | 0 | 1 |
| PCBP2 | 0.93 | 3 | 6 | 4 | 5 | 4 | 3 | 6 | 7 |
| WIPI2 | 0.93 | 8 | 6 | 10 | 6 | 6 | 11 | 13 | 11 |
| SEPT7 | 0.92 | 10 | 17 | 19 | 16 | 19 | 20 | 23 | 23 |
| ABI1 | 0.92 | 3 | 2 | 2 | 1 | 1 | 2 | 4 | 5 |
| CBL | 0.92 | 2 | 1 | 4 | 0 | 2 | 1 | 2 | 6 |
| ELP5 | 0.92 | 4 | 4 | 5 | 6 | 5 | 5 | 4 | 5 |
| LYPLA2 | 0.92 | 4 | 4 | 5 | 4 | 4 | 5 | 8 | 7 |
| MFSD10 | 0.92 | 1 | 0 | 0 | 1 | 0 | 0 | 0 | 0 |
| MSTO1 | 0.92 | 0 | 0 | 2 | 0 | 0 | 1 | 1 | 0 |
| RPS15A | 0.92 | 4 | 5 | 6 | 3 | 8 | 7 | 5 | 6 |
| RPS5 | 0.92 | 2 | 3 | 1 | 0 | 2 | 2 | 3 | 4 |
| TOM1L2 | 0.92 | 0 | 0 | 2 | 0 | 0 | 1 | 1 | 0 |
| TUBB6 | 0.92 | 6 | 9 | 7 | 8 | 11 | 8 | 7 | 9 |
| ALG13 | 0.91 | 1 | 1 | 1 | 1 | 2 | 1 | 1 | 1 |
| CCDC50 | 0.91 | 1 | 1 | 1 | 0 | 0 | 1 | 2 | 2 |

| | | | | | | | | | |
|---|---|---|---|---|---|---|---|---|---|
| EIF3G | 0.91 | 1 | 3 | 3 | 2 | 2 | 1 | 4 | 4 |
| ENAH | 0.91 | 0 | 1 | 2 | 0 | 0 | 1 | 1 | 2 |
| IARS | 0.91 | 3 | 7 | 9 | 3 | 7 | 8 | 9 | 10 |
| PSMC4 | 0.91 | 3 | 3 | 2 | 1 | 4 | 5 | 2 | 2 |
| RPS4X | 0.91 | 4 | 4 | 3 | 4 | 5 | 5 | 1 | 1 |
| SEC24B | 0.91 | 1 | 4 | 2 | 2 | 2 | 3 | 3 | 4 |
| SNX24 | 0.91 | 0 | 2 | 2 | 2 | 1 | 1 | 1 | 1 |
| TACC3 | 0.91 | 2 | 2 | 3 | 4 | 1 | 2 | 3 | 1 |
| TBC1D2B | 0.9 | 1 | 2 | 1 | 0 | 0 | 3 | 2 | 1 |
| TRIM25 | 0.9 | 9 | 7 | 8 | 7 | 10 | 11 | 13 | 13 |
| ATG2B | 0.89 | 10 | 9 | 9 | 13 | 5 | 6 | 14 | 14 |
| CYFIP1 | 0.89 | 9 | 12 | 12 | 12 | 15 | 15 | 12 | 16 |
| DNAJB1 | 0.89 | 7 | 4 | 6 | 9 | 6 | 6 | 7 | 7 |
| FKBP15 | 0.89 | 1 | 2 | 1 | 1 | 1 | 0 | 4 | 1 |
| G6PD | 0.89 | 7 | 6 | 8 | 7 | 11 | 9 | 8 | 7 |
| MSH6 | 0.89 | 1 | 1 | 1 | 2 | 1 | 1 | 0 | 0 |
| PSMC5 | 0.89 | 2 | 2 | 1 | 1 | 1 | 0 | 4 | 3 |
| RPS27 | 0.89 | 2 | 1 | 2 | 1 | 1 | 0 | 4 | 3 |
| TUBG1;TUBG2 | 0.89 | 0 | 0 | 2 | 0 | 1 | 1 | 0 | 0 |
| KIAA0368;ECM29 | 0.88 | 4 | 14 | 23 | 10 | 19 | 15 | 1 | 0 |
| MGEA5 | 0.88 | 4 | 6 | 8 | 6 | 9 | 8 | 4 | 8 |
| TANK | 0.88 | 2 | 0 | 1 | 0 | 0 | 1 | 3 | 0 |
| TSG101 | 0.88 | 1 | 1 | 3 | 2 | 1 | 1 | 1 | 4 |
| UBE2O | 0.88 | 7 | 10 | 7 | 9 | 10 | 9 | 11 | 15 |
| BAG3 | 0.87 | 9 | 8 | 10 | 6 | 9 | 5 | 18 | 17 |
| DDX39A | 0.87 | 2 | 5 | 3 | 1 | 3 | 1 | 6 | 7 |
| EEF2 | 0.87 | 22 | 17 | 19 | 26 | 21 | 28 | 21 | 19 |
| RBMX;RBMXL1 | 0.87 | 3 | 0 | 0 | 0 | 1 | 0 | 1 | 1 |
| RPL15 | 0.87 | 5 | 5 | 4 | 1 | 5 | 1 | 9 | 9 |
| RPS24 | 0.87 | 2 | 2 | 2 | 1 | 2 | 3 | 4 | 3 |
| SYNCRIP | 0.87 | 3 | 4 | 3 | 4 | 7 | 1 | 1 | 3 |
| GORASP2 | 0.86 | 2 | 2 | 5 | 2 | 1 | 4 | 5 | 5 |
| PTK2 | 0.86 | 2 | 3 | 4 | 4 | 4 | 2 | 4 | 5 |
| RPL22 | 0.86 | 1 | 3 | 1 | 0 | 2 | 4 | 1 | 1 |
| RPS20 | 0.86 | 4 | 3 | 4 | 4 | 5 | 6 | 2 | 4 |
| SLC25A5 | 0.86 | 7 | 7 | 6 | 7 | 10 | 10 | 7 | 5 |
| SMG7 | 0.86 | 0 | 1 | 1 | 0 | 1 | 1 | 1 | 1 |
| TTC28 | 0.86 | 1 | 1 | 1 | 0 | 1 | 2 | 2 | 0 |
| CAD | 0.85 | 2 | 4 | 3 | 3 | 5 | 4 | 2 | 5 |
| CEP170 | 0.85 | 42 | 49 | 43 | 33 | 49 | 49 | 88 | 85 |
| ILF2 | 0.85 | 1 | 0 | 1 | 0 | 1 | 1 | 1 | 0 |
| RAB3GAP2 | 0.85 | 3 | 6 | 3 | 3 | 6 | 6 | 3 | 6 |
| RPS23 | 0.85 | 3 | 2 | 2 | 1 | 3 | 2 | 3 | 6 |
| SDC1 | 0.85 | 2 | 0 | 0 | 1 | 1 | 0 | 0 | 0 |
| STAT3 | 0.85 | 12 | 4 | 9 | 8 | 9 | 10 | 11 | 17 |

| | | | | | | | | | |
|---|---|---|---|---|---|---|---|---|---|
| TOP2A | 0.85 | 4 | 0 | 1 | 0 | 0 | 0 | 6 | 0 |
| WIPI1 | 0.85 | 2 | 4 | 4 | 2 | 4 | 6 | 5 | 3 |
| COPB1 | 0.84 | 7 | 7 | 11 | 6 | 11 | 10 | 14 | 16 |
| DVL2 | 0.84 | 3 | 5 | 8 | 3 | 4 | 2 | 10 | 12 |
| EIF4G2 | 0.84 | 13 | 13 | 8 | 12 | 12 | 18 | 19 | 17 |
| ETF1 | 0.84 | 3 | 4 | 3 | 3 | 3 | 5 | 4 | 8 |
| FAM101B | 0.84 | 0 | 1 | 0 | 0 | 0 | 0 | 1 | 1 |
| FAM91A1 | 0.84 | 7 | 5 | 7 | 5 | 7 | 7 | 13 | 12 |
| KARS | 0.84 | 2 | 5 | 3 | 3 | 5 | 4 | 6 | 3 |
| MMP14 | 0.84 | 0 | 1 | 0 | 0 | 0 | 0 | 1 | 1 |
| PLIN3 | 0.84 | 25 | 30 | 25 | 31 | 34 | 44 | 26 | 28 |
| SLC1A5 | 0.84 | 5 | 1 | 1 | 3 | 2 | 2 | 3 | 3 |
| SNCA | 0.84 | 6 | 4 | 3 | 7 | 5 | 5 | 3 | 6 |
| TXNDC9 | 0.84 | 3 | 2 | 1 | 3 | 3 | 2 | 2 | 2 |
| AP2A1 | 0.83 | 9 | 15 | 11 | 15 | 19 | 6 | 15 | 13 |
| ARFIP1 | 0.83 | 0 | 0 | 1 | 0 | 0 | 0 | 1 | 1 |
| ARHGAP18 | 0.83 | 13 | 10 | 12 | 15 | 15 | 18 | 15 | 13 |
| CENPC | 0.83 | 0 | 1 | 0 | 0 | 0 | 0 | 2 | 0 |
| DYNC1LI2 | 0.83 | 2 | 2 | 4 | 1 | 2 | 2 | 5 | 7 |
| EIF5 | 0.83 | 17 | 9 | 13 | 16 | 18 | 16 | 19 | 23 |
| EPS8L2 | 0.83 | 1 | 0 | 1 | 1 | 0 | 1 | 1 | 0 |
| HIST2H2AA3;HIST2H2AC | 0.83 | 1 | 0 | 0 | 0 | 0 | 0 | 1 | 1 |
| HNRNPC | 0.83 | 2 | 1 | 3 | 1 | 3 | 0 | 3 | 4 |
| KIDINS220 | 0.83 | 0 | 1 | 0 | 0 | 0 | 0 | 2 | 0 |
| KIF5B | 0.83 | 26 | 28 | 24 | 38 | 33 | 33 | 37 | 44 |
| NDUFA4 | 0.83 | 1 | 0 | 0 | 0 | 0 | 0 | 1 | 1 |
| PDAP1 | 0.83 | 4 | 6 | 6 | 3 | 2 | 4 | 11 | 13 |
| RNF213 | 0.83 | 0 | 0 | 1 | 0 | 0 | 0 | 1 | 1 |
| SDCBP | 0.83 | 1 | 0 | 2 | 1 | 1 | 0 | 2 | 1 |
| SMC3 | 0.83 | 1 | 0 | 0 | 0 | 0 | 0 | 1 | 1 |
| ANKRD52 | 0.82 | 1 | 1 | 0 | 1 | 1 | 1 | 1 | 0 |
| CBX5 | 0.82 | 1 | 0 | 0 | 0 | 0 | 0 | 2 | 0 |
| EXOC4 | 0.82 | 3 | 2 | 3 | 4 | 1 | 2 | 5 | 4 |
| KIAA1524 | 0.82 | 9 | 6 | 6 | 10 | 7 | 9 | 12 | 9 |
| NFKBIE | 0.82 | 1 | 0 | 0 | 0 | 0 | 0 | 2 | 0 |
| PAICS | 0.82 | 14 | 24 | 22 | 24 | 25 | 33 | 22 | 27 |
| RPS11 | 0.82 | 5 | 8 | 10 | 8 | 14 | 7 | 11 | 10 |
| RPS8 | 0.82 | 4 | 7 | 5 | 4 | 9 | 6 | 8 | 10 |
| SNX1 | 0.82 | 6 | 7 | 6 | 10 | 8 | 6 | 10 | 10 |
| TAB1 | 0.82 | 4 | 1 | 3 | 4 | 4 | 0 | 2 | 3 |
| TIPRL | 0.82 | 7 | 8 | 7 | 10 | 8 | 11 | 9 | 13 |
| VPS33B | 0.82 | 3 | 1 | 1 | 1 | 3 | 3 | 1 | 0 |
| ARPC1B | 0.81 | 5 | 6 | 8 | 5 | 9 | 11 | 5 | 11 |
| ASAP1 | 0.81 | 4 | 3 | 3 | 2 | 6 | 4 | 4 | 7 |
| EEF1D | 0.81 | 5 | 8 | 6 | 8 | 11 | 7 | 9 | 10 |

| | | | | | | | | |
|---|---|---|---|---|---|---|---|---|
| ITGB6 | 0.81 | 0 | 1 | 1 | 1 | 1 | 1 | 0 | 1 |
| PEA15 | 0.81 | 0 | 1 | 1 | 1 | 1 | 1 | 0 | 0 |
| PIK3R4 | 0.81 | 0 | 1 | 0 | 0 | 1 | 0 | 0 | 1 |
| PSMD13 | 0.81 | 1 | 3 | 1 | 2 | 3 | 2 | 1 | 0 |
| PTPN12 | 0.81 | 12 | 10 | 10 | 13 | 11 | 11 | 20 | 20 |
| RPS9 | 0.81 | 8 | 6 | 4 | 7 | 10 | 8 | 6 | 8 |
| RSL1D1 | 0.81 | 0 | 1 | 1 | 1 | 2 | 0 | 0 | 0 |
| SH3KBP1 | 0.81 | 21 | 19 | 11 | 18 | 21 | 16 | 34 | 30 |
| TAGLN2 | 0.81 | 13 | 13 | 20 | 17 | 19 | 28 | 13 | 17 |
| TMEM165 | 0.81 | 1 | 2 | 1 | 2 | 2 | 2 | 1 | 0 |
| TUBB8 | 0.81 | 2 | 0 | 3 | 1 | 0 | 0 | 3 | 3 |
| UNC45A | 0.81 | 10 | 8 | 8 | 11 | 8 | 7 | 16 | 16 |
| ABCF3 | 0.8 | 2 | 0 | 2 | 1 | 1 | 0 | 3 | 2 |
| AP2S1 | 0.8 | 1 | 1 | 1 | 1 | 1 | 1 | 2 | 3 |
| COPG2 | 0.8 | 29 | 36 | 37 | 46 | 45 | 52 | 48 | 56 |
| DCTN1 | 0.8 | 2 | 9 | 9 | 4 | 4 | 4 | 15 | 11 |
| DDX3X;DDX3Y | 0.8 | 10 | 14 | 11 | 9 | 15 | 20 | 21 | 18 |
| EFHD2 | 0.8 | 14 | 10 | 9 | 15 | 13 | 17 | 17 | 19 |
| HDLBP | 0.8 | 15 | 21 | 24 | 18 | 28 | 29 | 29 | 42 |
| NAMPT | 0.8 | 1 | 0 | 0 | 0 | 0 | 1 | 1 | 0 |
| RUFY1 | 0.8 | 1 | 2 | 1 | 1 | 2 | 2 | 3 | 1 |
| SKIV2L | 0.8 | 1 | 0 | 1 | 1 | 1 | 1 | 0 | 0 |
| SPAG9 | 0.8 | 6 | 10 | 12 | 7 | 10 | 7 | 19 | 19 |
| STAT6 | 0.8 | 1 | 0 | 1 | 1 | 1 | 1 | 1 | 0 |
| ATP6V1E1 | 0.79 | 1 | 0 | 0 | 0 | 1 | 0 | 1 | 0 |
| CLNS1A | 0.79 | 1 | 1 | 0 | 0 | 1 | 0 | 1 | 2 |
| EIF2A | 0.79 | 8 | 12 | 9 | 13 | 11 | 16 | 15 | 14 |
| HSP90AB1 | 0.79 | 13 | 10 | 11 | 15 | 16 | 8 | 20 | 22 |
| ITPR3 | 0.79 | 0 | 0 | 1 | 0 | 1 | 0 | 1 | 0 |
| MYL6 | 0.79 | 22 | 12 | 7 | 20 | 15 | 18 | 5 | 10 |
| NCL | 0.79 | 5 | 10 | 10 | 7 | 14 | 6 | 14 | 14 |
| SQSTM1 | 0.79 | 1 | 3 | 3 | 0 | 4 | 1 | 4 | 4 |
| SSRP1 | 0.79 | 4 | 0 | 0 | 1 | 0 | 0 | 3 | 0 |
| WARS | 0.79 | 6 | 8 | 7 | 4 | 8 | 14 | 11 | 12 |
| ATP5C1 | 0.78 | 1 | 1 | 2 | 0 | 1 | 1 | 3 | 4 |
| CAMK4 | 0.78 | 1 | 0 | 2 | 2 | 1 | 1 | 1 | 0 |
| DIAPH1 | 0.78 | 2 | 2 | 1 | 4 | 1 | 1 | 3 | 1 |
| GAPDH | 0.78 | 3 | 4 | 3 | 4 | 4 | 3 | 6 | 8 |
| NDE1;NDEL1 | 0.78 | 0 | 1 | 0 | 0 | 0 | 2 | 0 | 0 |
| PIP4K2C | 0.78 | 3 | 1 | 5 | 2 | 3 | 3 | 5 | 7 |
| RPL23A | 0.78 | 0 | 1 | 0 | 1 | 0 | 0 | 0 | 1 |
| RPL9 | 0.78 | 6 | 2 | 5 | 6 | 5 | 2 | 5 | 9 |
| SLC25A10 | 0.78 | 1 | 2 | 0 | 0 | 1 | 1 | 3 | 1 |
| AGFG1 | 0.77 | 2 | 2 | 4 | 2 | 4 | 4 | 3 | 6 |
| ARHGDIA | 0.77 | 1 | 0 | 0 | 0 | 1 | 1 | 0 | 0 |

| | | | | | | | | | |
|---|---|---|---|---|---|---|---|---|---|
| ARL1 | 0.77 | 1 | 1 | 1 | 1 | 2 | 2 | 1 | 2 |
| CNOT1 | 0.77 | 1 | 0 | 1 | 0 | 1 | 0 | 2 | 1 |
| DOK1 | 0.77 | 0 | 1 | 1 | 0 | 1 | 1 | 1 | 2 |
| DYNC1I2 | 0.77 | 3 | 4 | 3 | 3 | 5 | 3 | 6 | 8 |
| FLNC | 0.77 | 54 | 57 | 53 | 48 | 53 | 55 | 133 | 112 |
| GFPT1 | 0.77 | 3 | 4 | 4 | 1 | 3 | 3 | 11 | 7 |
| HAUS5 | 0.77 | 1 | 0 | 0 | 0 | 1 | 1 | 0 | 0 |
| HGS | 0.77 | 5 | 4 | 4 | 3 | 4 | 2 | 11 | 10 |
| PSMD12 | 0.77 | 2 | 1 | 1 | 1 | 4 | 0 | 1 | 2 |
| QARS | 0.77 | 3 | 9 | 9 | 8 | 7 | 11 | 12 | 11 |
| RANGAP1 | 0.77 | 14 | 16 | 13 | 21 | 20 | 16 | 26 | 24 |
| SLK | 0.77 | 20 | 25 | 29 | 24 | 29 | 38 | 44 | 51 |
| TBC1D5 | 0.77 | 1 | 1 | 2 | 0 | 1 | 1 | 5 | 2 |
| DVL1;DVL1P1 | 0.76 | 3 | 5 | 4 | 3 | 5 | 5 | 8 | 10 |
| KLC2 | 0.76 | 1 | 2 | 0 | 0 | 1 | 2 | 2 | 0 |
| PLOD1 | 0.76 | 0 | 4 | 2 | 0 | 0 | 1 | 6 | 2 |
| PSMD5 | 0.76 | 0 | 0 | 1 | 0 | 1 | 1 | 0 | 0 |
| RAPH1 | 0.76 | 25 | 18 | 24 | 35 | 32 | 25 | 32 | 41 |
| WIBG | 0.76 | 0 | 1 | 1 | 0 | 0 | 2 | 1 | 1 |
| ABCD3 | 0.75 | 1 | 1 | 1 | 1 | 2 | 1 | 3 | 0 |
| IGBP1 | 0.75 | 2 | 3 | 6 | 4 | 3 | 9 | 4 | 4 |
| INPPL1 | 0.75 | 1 | 2 | 4 | 0 | 2 | 0 | 6 | 5 |
| LARP1B | 0.75 | 0 | 1 | 1 | 1 | 0 | 1 | 0 | 2 |
| MAP4 | 0.75 | 43 | 36 | 47 | 56 | 51 | 61 | 85 | 82 |
| PIP4K2A | 0.75 | 1 | 0 | 1 | 1 | 0 | 0 | 2 | 1 |
| RASSF8;C12orf2 | 0.75 | 1 | 1 | 1 | 1 | 2 | 1 | 3 | 1 |
| ADSL | 0.74 | 1 | 0 | 1 | 0 | 1 | 1 | 2 | 0 |
| CKAP4 | 0.74 | 17 | 17 | 7 | 15 | 23 | 13 | 22 | 27 |
| CYFIP2 | 0.74 | 0 | 1 | 0 | 1 | 1 | 0 | 0 | 0 |
| EIF5A;EIF5A2 | 0.74 | 1 | 2 | 1 | 1 | 2 | 3 | 2 | 3 |
| PSMA2 | 0.74 | 3 | 1 | 1 | 2 | 3 | 3 | 0 | 1 |
| RPL23 | 0.74 | 3 | 3 | 5 | 4 | 4 | 2 | 6 | 11 |
| RUFY3 | 0.74 | 0 | 0 | 1 | 1 | 0 | 1 | 0 | 0 |
| SLC16A3 | 0.74 | 1 | 0 | 1 | 1 | 1 | 1 | 1 | 2 |
| TARS | 0.74 | 0 | 1 | 2 | 1 | 1 | 2 | 2 | 1 |
| TRIO | 0.74 | 0 | 1 | 1 | 0 | 3 | 0 | 0 | 1 |
| ANXA5 | 0.73 | 0 | 0 | 2 | 0 | 1 | 2 | 0 | 0 |
| EIF3C;EIF3CL | 0.73 | 5 | 5 | 2 | 5 | 6 | 4 | 9 | 6 |
| EIF4G3 | 0.73 | 3 | 6 | 6 | 7 | 7 | 6 | 10 | 10 |
| MAP1B | 0.73 | 24 | 26 | 32 | 19 | 28 | 21 | 71 | 60 |
| MARS | 0.73 | 2 | 1 | 5 | 2 | 4 | 4 | 5 | 3 |
| NUDCD3 | 0.73 | 0 | 1 | 0 | 0 | 0 | 0 | 0 | 3 |
| OXSR1 | 0.73 | 1 | 1 | 1 | 2 | 1 | 0 | 3 | 1 |
| PSMD8 | 0.73 | 4 | 6 | 3 | 3 | 5 | 8 | 8 | 9 |
| PXN | 0.73 | 11 | 9 | 15 | 12 | 20 | 20 | 20 | 22 |

| | | | | | | | | | |
|---|---|---|---|---|---|---|---|---|---|
| SYAP1 | 0.73 | 2 | 2 | 2 | 3 | 3 | 3 | 5 | 3 |
| UACA | 0.73 | 0 | 1 | 0 | 0 | 0 | 0 | 0 | 3 |
| CAMK2D | 0.72 | 2 | 2 | 3 | 3 | 5 | 4 | 2 | 3 |
| COPG1 | 0.72 | 4 | 6 | 7 | 9 | 8 | 10 | 5 | 6 |
| SCYL1 | 0.72 | 0 | 2 | 0 | 0 | 0 | 0 | 2 | 2 |
| SNX2 | 0.72 | 1 | 0 | 1 | 1 | 0 | 1 | 2 | 0 |
| AP3M1 | 0.71 | 0 | 2 | 1 | 1 | 2 | 2 | 0 | 0 |
| BCL2L2;BCL2L2-P... | 0.71 | 0 | 0 | 1 | 0 | 0 | 0 | 1 | 2 |
| CTPS1 | 0.71 | 10 | 3 | 7 | 9 | 12 | 9 | 7 | 10 |
| FMNL2 | 0.71 | 1 | 0 | 0 | 0 | 0 | 0 | 1 | 2 |
| MAGED1 | 0.71 | 1 | 0 | 0 | 0 | 0 | 0 | 1 | 2 |
| NAP1L1 | 0.71 | 6 | 15 | 8 | 9 | 17 | 19 | 11 | 13 |
| PKM | 0.71 | 33 | 25 | 33 | 50 | 42 | 52 | 32 | 42 |
| PPP6C | 0.71 | 3 | 2 | 2 | 3 | 3 | 2 | 3 | 8 |
| RARS | 0.71 | 12 | 8 | 15 | 15 | 17 | 16 | 25 | 25 |
| RPS14 | 0.71 | 5 | 3 | 2 | 4 | 6 | 6 | 6 | 5 |
| EIF3F | 0.7 | 2 | 1 | 2 | 2 | 3 | 4 | 3 | 2 |
| PSMD3 | 0.7 | 3 | 4 | 1 | 1 | 3 | 4 | 6 | 6 |
| RABGAP1 | 0.7 | 1 | 1 | 0 | 0 | 0 | 0 | 5 | 0 |
| SHROOM3 | 0.7 | 1 | 1 | 1 | 1 | 2 | 2 | 1 | 3 |
| STAT1 | 0.7 | 14 | 15 | 11 | 13 | 21 | 27 | 25 | 24 |
| SURF4 | 0.7 | 0 | 1 | 3 | 0 | 1 | 2 | 3 | 2 |
| TCEB2 | 0.7 | 0 | 0 | 1 | 0 | 0 | 0 | 2 | 1 |
| ZW10 | 0.7 | 1 | 1 | 0 | 1 | 2 | 1 | 1 | 1 |
| DOCK7 | 0.69 | 10 | 6 | 11 | 11 | 8 | 13 | 25 | 15 |
| GULP1 | 0.69 | 0 | 1 | 0 | 0 | 0 | 1 | 1 | 1 |
| SPC24 | 0.69 | 1 | 0 | 1 | 0 | 0 | 1 | 2 | 2 |
| ARPC5 | 0.68 | 0 | 1 | 1 | 0 | 1 | 1 | 1 | 3 |
| CASP3 | 0.68 | 1 | 2 | 1 | 0 | 1 | 0 | 3 | 6 |
| COMMD8 | 0.68 | 1 | 1 | 0 | 1 | 1 | 1 | 2 | 2 |
| CSE1L | 0.68 | 3 | 0 | 0 | 2 | 0 | 1 | 0 | 1 |
| LMNA | 0.68 | 18 | 8 | 11 | 15 | 12 | 10 | 28 | 28 |
| MYH9 | 0.68 | 816 | 272 | 310 | 677 | 589 | 777 | 74 | 464 |
| NUDC | 0.68 | 9 | 9 | 8 | 11 | 10 | 8 | 17 | 25 |
| PSMB1 | 0.68 | 1 | 1 | 0 | 2 | 1 | 1 | 0 | 0 |
| ARHGAP35 | 0.67 | 5 | 8 | 13 | 9 | 13 | 11 | 17 | 22 |
| ARPC2 | 0.67 | 6 | 3 | 4 | 6 | 4 | 5 | 3 | 15 |
| C1orf198 | 0.67 | 1 | 0 | 0 | 0 | 0 | 1 | 2 | 0 |
| CHMP4B | 0.67 | 1 | 1 | 1 | 3 | 1 | 1 | 2 | 2 |
| CORO1C | 0.67 | 33 | 28 | 25 | 44 | 48 | 53 | 41 | 45 |
| DCP1B | 0.67 | 1 | 2 | 1 | 2 | 3 | 3 | 0 | 1 |
| DHPS | 0.67 | 1 | 1 | 0 | 1 | 1 | 0 | 0 | 3 |
| KANK2 | 0.67 | 13 | 20 | 30 | 30 | 32 | 34 | 46 | 42 |
| MAPRE2 | 0.67 | 4 | 4 | 6 | 4 | 10 | 9 | 4 | 9 |
| NUP214 | 0.67 | 7 | 13 | 11 | 12 | 16 | 11 | 25 | 23 |

| | | | | | | | | | |
|---|---|---|---|---|---|---|---|---|---|
| PABPC1 | 0.67 | 6 | 11 | 5 | 9 | 8 | 10 | 16 | 19 |
| PSMD1 | 0.67 | 1 | 3 | 2 | 2 | 4 | 2 | 6 | 3 |
| RIC8A | 0.67 | 2 | 2 | 2 | 1 | 3 | 5 | 5 | 2 |
| RPL24 | 0.67 | 2 | 4 | 3 | 3 | 4 | 6 | 5 | 9 |
| RUVBL1 | 0.67 | 15 | 16 | 14 | 23 | 26 | 28 | 26 | 31 |
| SHROOM2 | 0.67 | 1 | 0 | 1 | 0 | 0 | 2 | 2 | 1 |
| ZC3HAV1 | 0.67 | 5 | 6 | 7 | 6 | 8 | 10 | 16 | 13 |
| CEP44 | 0.66 | 1 | 1 | 1 | 1 | 1 | 0 | 3 | 4 |
| CEP97 | 0.66 | 1 | 1 | 2 | 0 | 2 | 1 | 4 | 4 |
| CSRP1 | 0.66 | 2 | 2 | 2 | 1 | 1 | 2 | 5 | 8 |
| ECHDC1 | 0.66 | 0 | 0 | 1 | 0 | 1 | 1 | 0 | 1 |
| EPS15 | 0.66 | 0 | 0 | 3 | 0 | 1 | 1 | 1 | 3 |
| FIBP | 0.66 | 0 | 0 | 1 | 0 | 0 | 2 | 0 | 1 |
| MSN | 0.66 | 11 | 12 | 10 | 14 | 21 | 13 | 24 | 26 |
| NUMBL | 0.66 | 1 | 2 | 3 | 3 | 4 | 4 | 3 | 4 |
| RPL14 | 0.66 | 1 | 1 | 1 | 2 | 1 | 2 | 3 | 2 |
| UBAP1 | 0.66 | 0 | 1 | 0 | 0 | 1 | 1 | 1 | 1 |
| YWHAQ | 0.66 | 1 | 1 | 2 | 3 | 1 | 1 | 4 | 2 |
| ALDH1A3 | 0.65 | 2 | 0 | 1 | 0 | 1 | 1 | 3 | 3 |
| EHBP1 | 0.65 | 2 | 6 | 6 | 5 | 4 | 5 | 16 | 7 |
| EIF4H | 0.65 | 0 | 2 | 3 | 0 | 3 | 3 | 2 | 3 |
| FARSB | 0.65 | 2 | 3 | 3 | 5 | 5 | 5 | 2 | 6 |
| FHOD1 | 0.65 | 3 | 2 | 3 | 6 | 3 | 2 | 5 | 6 |
| HECTD1 | 0.65 | 0 | 3 | 1 | 1 | 2 | 1 | 4 | 2 |
| LDHA | 0.65 | 4 | 4 | 7 | 8 | 8 | 11 | 3 | 3 |
| PACSIN2 | 0.65 | 0 | 0 | 1 | 0 | 1 | 1 | 1 | 1 |
| PDLIM7 | 0.65 | 5 | 7 | 4 | 6 | 5 | 5 | 12 | 17 |
| STAM2 | 0.65 | 4 | 3 | 2 | 3 | 4 | 3 | 8 | 9 |
| TMEM263 | 0.65 | 3 | 1 | 3 | 1 | 2 | 1 | 6 | 8 |
| ANK2 | 0.64 | 0 | 1 | 1 | 0 | 1 | 3 | 1 | 1 |
| AP2M1 | 0.64 | 4 | 5 | 3 | 1 | 5 | 2 | 11 | 12 |
| EIF3D | 0.64 | 1 | 1 | 2 | 2 | 3 | 0 | 4 | 3 |
| HSP90AB2P | 0.64 | 0 | 0 | 2 | 0 | 2 | 1 | 1 | 0 |
| MRE11A | 0.64 | 10 | 11 | 9 | 11 | 16 | 12 | 22 | 30 |
| NAP1L4 | 0.64 | 7 | 4 | 3 | 7 | 7 | 11 | 6 | 5 |
| NBAS | 0.64 | 0 | 1 | 0 | 1 | 0 | 1 | 1 | 1 |
| RASA1 | 0.64 | 1 | 0 | 0 | 1 | 1 | 0 | 0 | 1 |
| RPS12 | 0.64 | 2 | 2 | 2 | 4 | 4 | 4 | 3 | 4 |
| SKA3 | 0.64 | 2 | 1 | 1 | 2 | 2 | 3 | 4 | 1 |
| SEPT10 | 0.63 | 2 | 5 | 4 | 5 | 5 | 5 | 10 | 9 |
| NAV1 | 0.63 | 3 | 2 | 2 | 2 | 2 | 2 | 10 | 5 |
| PRKAA1 | 0.63 | 1 | 0 | 1 | 0 | 0 | 0 | 3 | 3 |
| TAGLN | 0.63 | 7 | 5 | 4 | 2 | 5 | 8 | 13 | 16 |
| YAP1 | 0.63 | 4 | 5 | 2 | 4 | 4 | 7 | 10 | 8 |
| BIN1 | 0.62 | 2 | 1 | 3 | 3 | 1 | 2 | 5 | 6 |

| | | | | | | | | | |
|---|---|---|---|---|---|---|---|---|---|
| DAB2 | 0.62 | 4 | 6 | 4 | 6 | 4 | 2 | 12 | 14 |
| DPCD | 0.62 | 0 | 1 | 0 | 1 | 1 | 1 | 0 | 0 |
| ELP4 | 0.62 | 6 | 6 | 5 | 12 | 10 | 7 | 12 | 8 |
| GOLGA3 | 0.62 | 0 | 1 | 1 | 0 | 1 | 0 | 3 | 2 |
| HIST2H3PS2 | 0.62 | 1 | 0 | 0 | 1 | 1 | 1 | 0 | 1 |
| HSP90AA1 | 0.62 | 1 | 2 | 1 | 3 | 3 | 2 | 3 | 3 |
| KIZ | 0.62 | 0 | 1 | 0 | 1 | 0 | 2 | 0 | 0 |
| NACA | 0.62 | 2 | 1 | 2 | 2 | 3 | 5 | 2 | 4 |
| RABL6 | 0.62 | 0 | 1 | 0 | 1 | 1 | 1 | 1 | 1 |
| RBM39 | 0.62 | 0 | 1 | 1 | 2 | 1 | 1 | 2 | 0 |
| STRAP | 0.62 | 6 | 10 | 9 | 16 | 15 | 14 | 13 | 19 |
| BRCC3 | 0.61 | 0 | 0 | 1 | 2 | 0 | 0 | 1 | 0 |
| GEMIN5 | 0.61 | 23 | 24 | 24 | 29 | 53 | 31 | 52 | 55 |
| HSP90B1 | 0.61 | 2 | 0 | 0 | 0 | 1 | 1 | 2 | 2 |
| HSPA9 | 0.61 | 6 | 7 | 10 | 9 | 11 | 10 | 24 | 19 |
| MICALL1 | 0.61 | 0 | 0 | 1 | 1 | 1 | 1 | 0 | 1 |
| MOV10 | 0.61 | 1 | 0 | 1 | 1 | 1 | 3 | 0 | 0 |
| MYL12B;MYL12A | 0.61 | 44 | 8 | 8 | 42 | 21 | 15 | 4 | 14 |
| NUP88 | 0.61 | 0 | 4 | 2 | 1 | 2 | 2 | 6 | 4 |
| PACS1 | 0.61 | 1 | 1 | 2 | 1 | 3 | 4 | 2 | 3 |
| PGRMC2 | 0.61 | 2 | 3 | 2 | 2 | 5 | 4 | 5 | 8 |
| PLEKHA1 | 0.61 | 1 | 0 | 1 | 0 | 0 | 2 | 1 | 3 |
| RPL28 | 0.61 | 0 | 1 | 2 | 0 | 0 | 0 | 3 | 5 |
| RPL35A | 0.61 | 1 | 0 | 0 | 0 | 0 | 0 | 2 | 2 |
| STRN4 | 0.61 | 0 | 0 | 1 | 0 | 0 | 0 | 2 | 2 |
| TPM4 | 0.61 | 4 | 2 | 1 | 5 | 0 | 2 | 0 | 8 |
| VASP | 0.61 | 3 | 4 | 7 | 5 | 4 | 2 | 12 | 15 |
| AP2B1 | 0.6 | 12 | 7 | 11 | 17 | 20 | 12 | 19 | 25 |
| ARHGAP29 | 0.6 | 11 | 17 | 9 | 18 | 16 | 11 | 34 | 30 |
| EFTUD1 | 0.6 | 2 | 1 | 5 | 3 | 4 | 5 | 7 | 6 |
| EPRS | 0.6 | 4 | 13 | 10 | 7 | 14 | 13 | 18 | 28 |
| FAM129B | 0.6 | 5 | 9 | 14 | 13 | 16 | 22 | 17 | 20 |
| FGD6 | 0.6 | 0 | 0 | 1 | 0 | 0 | 0 | 3 | 1 |
| KIAA1671 | 0.6 | 3 | 5 | 4 | 5 | 5 | 4 | 13 | 11 |
| PGAM1 | 0.6 | 2 | 4 | 3 | 3 | 3 | 5 | 8 | 10 |
| PLS3 | 0.6 | 7 | 3 | 7 | 8 | 11 | 13 | 9 | 10 |
| PSMD4 | 0.6 | 2 | 1 | 3 | 2 | 2 | 1 | 7 | 6 |
| RPL11 | 0.6 | 2 | 3 | 2 | 3 | 3 | 2 | 7 | 8 |
| AGO2 | 0.59 | 0 | 2 | 0 | 1 | 1 | 0 | 1 | 3 |
| CSDE1 | 0.59 | 10 | 10 | 11 | 10 | 14 | 6 | 30 | 32 |
| EIF3M | 0.59 | 1 | 2 | 1 | 4 | 3 | 2 | 1 | 2 |
| PGAM5 | 0.59 | 3 | 3 | 2 | 4 | 5 | 5 | 8 | 8 |
| PSMC1 | 0.59 | 2 | 2 | 2 | 2 | 6 | 3 | 5 | 5 |
| SH3BP5L | 0.59 | 4 | 1 | 1 | 1 | 5 | 5 | 1 | 3 |
| SND1 | 0.59 | 1 | 2 | 4 | 3 | 7 | 4 | 4 | 4 |

| | | | | | | | | | |
|---|---|---|---|---|---|---|---|---|---|
| CAST | 0.58 | 9 | 7 | 12 | 12 | 12 | 13 | 28 | 27 |
| EEF1A1;EEF1A1P5 | 0.58 | 9 | 13 | 12 | 23 | 21 | 23 | 17 | 26 |
| KPNB1 | 0.58 | 5 | 6 | 5 | 11 | 10 | 11 | 13 | 8 |
| LIMCH1 | 0.58 | 16 | 22 | 22 | 35 | 36 | 40 | 47 | 57 |
| SEC31A | 0.58 | 1 | 2 | 5 | 3 | 4 | 6 | 3 | 8 |
| VAMP3 | 0.58 | 2 | 0 | 2 | 1 | 2 | 1 | 3 | 5 |
| AK5 | 0.57 | 0 | 0 | 1 | 0 | 1 | 1 | 2 | 1 |
| ARFGAP1 | 0.57 | 1 | 3 | 3 | 1 | 2 | 3 | 6 | 10 |
| ARPC3 | 0.57 | 2 | 1 | 3 | 4 | 3 | 3 | 2 | 8 |
| CACYBP | 0.57 | 2 | 2 | 2 | 4 | 6 | 4 | 2 | 5 |
| CAPZA1 | 0.57 | 15 | 11 | 9 | 22 | 25 | 23 | 14 | 27 |
| CIAPIN1 | 0.57 | 4 | 5 | 2 | 7 | 8 | 5 | 8 | 10 |
| ERC1 | 0.57 | 0 | 6 | 5 | 1 | 3 | 3 | 13 | 2 |
| GIGYF2 | 0.57 | 14 | 13 | 19 | 16 | 20 | 18 | 48 | 48 |
| GPRIN1 | 0.57 | 1 | 4 | 4 | 0 | 1 | 5 | 10 | 7 |
| MAVS | 0.57 | 3 | 4 | 3 | 5 | 7 | 10 | 6 | 6 |
| N4BP1 | 0.57 | 0 | 1 | 2 | 1 | 1 | 3 | 3 | 2 |
| RAB3GAP1 | 0.57 | 2 | 2 | 2 | 4 | 5 | 5 | 4 | 2 |
| TUBA4A | 0.57 | 0 | 1 | 0 | 1 | 0 | 1 | 0 | 2 |
| ABCF2 | 0.56 | 2 | 0 | 0 | 0 | 0 | 0 | 3 | 3 |
| ATP2A2 | 0.56 | 4 | 2 | 3 | 3 | 3 | 4 | 9 | 12 |
| BIN3 | 0.56 | 2 | 1 | 3 | 3 | 1 | 1 | 9 | 4 |
| CDC37 | 0.56 | 0 | 0 | 1 | 0 | 0 | 3 | 0 | 1 |
| CUL2 | 0.56 | 1 | 1 | 1 | 3 | 2 | 3 | 0 | 0 |
| FAU | 0.56 | 1 | 1 | 1 | 3 | 3 | 2 | 0 | 0 |
| HNRNPD | 0.56 | 2 | 2 | 1 | 5 | 4 | 1 | 3 | 1 |
| LRRFIP2 | 0.56 | 1 | 0 | 0 | 1 | 1 | 1 | 0 | 2 |
| NCK1 | 0.56 | 1 | 2 | 2 | 3 | 2 | 1 | 5 | 6 |
| RAB6A | 0.56 | 1 | 0 | 0 | 1 | 1 | 1 | 1 | 2 |
| RPL19 | 0.56 | 1 | 0 | 2 | 2 | 1 | 0 | 2 | 4 |
| SLC30A1 | 0.56 | 0 | 1 | 0 | 1 | 1 | 0 | 2 | 1 |
| SNX4 | 0.56 | 1 | 0 | 0 | 0 | 1 | 2 | 1 | 0 |
| TRIOBP | 0.56 | 3 | 3 | 3 | 3 | 1 | 3 | 14 | 8 |
| UBASH3B | 0.56 | 1 | 1 | 1 | 1 | 2 | 0 | 5 | 3 |
| SEPT8 | 0.55 | 1 | 0 | 3 | 1 | 0 | 2 | 4 | 4 |
| ASAP2 | 0.55 | 0 | 1 | 1 | 1 | 0 | 1 | 3 | 3 |
| ASCC3 | 0.55 | 4 | 6 | 10 | 12 | 13 | 13 | 15 | 20 |
| DCTN2 | 0.55 | 4 | 4 | 3 | 9 | 10 | 5 | 5 | 7 |
| EIF3E | 0.55 | 3 | 6 | 5 | 10 | 8 | 12 | 2 | 2 |
| EZR | 0.55 | 7 | 7 | 8 | 14 | 13 | 15 | 15 | 24 |
| HAUS8 | 0.55 | 0 | 0 | 1 | 1 | 1 | 1 | 2 | 0 |
| INPP5F | 0.55 | 7 | 4 | 6 | 6 | 11 | 6 | 16 | 18 |
| PDLIM1 | 0.55 | 11 | 7 | 12 | 17 | 25 | 21 | 21 | 25 |
| SMARCE1 | 0.55 | 1 | 0 | 0 | 2 | 0 | 0 | 0 | 2 |
| STRIP1;STRIP2 | 0.55 | 1 | 0 | 0 | 1 | 1 | 1 | 2 | 1 |

| | | | | | | | | | |
|---|---|---|---|---|---|---|---|---|---|
| TXNRD1 | 0.55 | 3 | 1 | 1 | 2 | 1 | 1 | 6 | 6 |
| USP47 | 0.55 | 0 | 1 | 1 | 0 | 1 | 2 | 2 | 3 |
| YWHAZ | 0.55 | 2 | 7 | 2 | 6 | 8 | 6 | 9 | 4 |
| ECI2 | 0.54 | 1 | 0 | 0 | 0 | 0 | 0 | 3 | 2 |
| MYH14 | 0.54 | 15 | 2 | 3 | 14 | 8 | 9 | 1 | 2 |
| RPL27A | 0.54 | 1 | 2 | 1 | 4 | 3 | 2 | 3 | 4 |
| RPL5 | 0.54 | 2 | 2 | 1 | 1 | 2 | 6 | 5 | 4 |
| RRM2 | 0.54 | 0 | 1 | 0 | 2 | 0 | 0 | 2 | 0 |
| RUVBL2 | 0.54 | 11 | 10 | 13 | 22 | 27 | 23 | 21 | 30 |
| SLAIN2 | 0.54 | 2 | 1 | 1 | 1 | 2 | 1 | 5 | 6 |
| TECR | 0.54 | 1 | 1 | 1 | 5 | 1 | 2 | 1 | 1 |
| WDR11 | 0.54 | 5 | 9 | 14 | 6 | 15 | 11 | 27 | 28 |
| ZFYVE16 | 0.54 | 1 | 0 | 1 | 1 | 0 | 1 | 4 | 2 |
| CCDC6 | 0.53 | 0 | 0 | 1 | 0 | 0 | 0 | 3 | 2 |
| FAM175B | 0.53 | 0 | 0 | 1 | 0 | 0 | 0 | 3 | 2 |
| GSTO1 | 0.53 | 1 | 1 | 1 | 1 | 0 | 1 | 4 | 6 |
| HMGA2 | 0.53 | 4 | 0 | 0 | 2 | 3 | 0 | 0 | 2 |
| NSFL1C | 0.53 | 1 | 1 | 0 | 1 | 2 | 2 | 3 | 1 |
| PSMA4 | 0.53 | 1 | 0 | 0 | 1 | 1 | 2 | 0 | 0 |
| STAM | 0.53 | 1 | 2 | 1 | 2 | 0 | 2 | 6 | 5 |
| SEPT11 | 0.52 | 7 | 2 | 8 | 5 | 7 | 6 | 16 | 21 |
| AFF4 | 0.52 | 1 | 0 | 0 | 2 | 1 | 0 | 1 | 0 |
| ARFGAP3 | 0.52 | 2 | 1 | 1 | 3 | 3 | 4 | 3 | 5 |
| DIAPH3 | 0.52 | 1 | 0 | 0 | 0 | 1 | 1 | 2 | 2 |
| EEF2K | 0.52 | 2 | 0 | 2 | 2 | 2 | 2 | 3 | 6 |
| EIF4B | 0.52 | 10 | 14 | 13 | 15 | 16 | 17 | 40 | 46 |
| FAM21A;FAM21C | 0.52 | 4 | 2 | 3 | 4 | 8 | 7 | 9 | 7 |
| HN1 | 0.52 | 1 | 1 | 1 | 1 | 3 | 3 | 4 | 4 |
| SH3GL1 | 0.52 | 1 | 1 | 2 | 1 | 1 | 1 | 4 | 9 |
| ATL3 | 0.51 | 2 | 0 | 1 | 2 | 2 | 1 | 4 | 3 |
| COPS3 | 0.51 | 0 | 1 | 0 | 1 | 1 | 1 | 1 | 3 |
| EIF3B | 0.51 | 5 | 3 | 1 | 2 | 8 | 4 | 8 | 8 |
| NUMB | 0.51 | 2 | 4 | 4 | 2 | 4 | 6 | 14 | 10 |
| NUP54 | 0.51 | 1 | 0 | 0 | 1 | 0 | 0 | 3 | 1 |
| PPIP5K2;PPIP5K1 | 0.51 | 0 | 0 | 1 | 1 | 1 | 0 | 2 | 2 |
| RAPGEF6 | 0.51 | 1 | 0 | 2 | 2 | 1 | 0 | 4 | 3 |
| RICTOR | 0.51 | 0 | 0 | 1 | 1 | 1 | 1 | 2 | 2 |
| SKA1 | 0.51 | 0 | 0 | 1 | 1 | 1 | 1 | 2 | 2 |
| SRSF3 | 0.51 | 1 | 0 | 0 | 2 | 2 | 0 | 0 | 0 |
| VCL | 0.51 | 10 | 9 | 14 | 16 | 12 | 18 | 39 | 34 |
| CEP55 | 0.5 | 0 | 2 | 1 | 1 | 2 | 1 | 4 | 4 |
| FAM160B1 | 0.5 | 4 | 1 | 0 | 3 | 3 | 4 | 4 | 2 |
| MYO9B | 0.5 | 0 | 0 | 1 | 0 | 1 | 1 | 3 | 1 |
| ACTR2 | 0.49 | 4 | 3 | 2 | 4 | 6 | 9 | 3 | 12 |
| CDV3 | 0.49 | 1 | 2 | 3 | 1 | 3 | 4 | 7 | 9 |

| | | | | | | | | | |
|---|---|---|---|---|---|---|---|---|---|
| EIF3A | 0.49 | 4 | 6 | 6 | 12 | 8 | 7 | 16 | 17 |
| FLOT2 | 0.49 | 0 | 1 | 0 | 0 | 0 | 0 | 2 | 4 |
| MYOF | 0.49 | 7 | 18 | 15 | 13 | 27 | 29 | 44 | 36 |
| PLEKHA2 | 0.49 | 0 | 1 | 2 | 3 | 2 | 1 | 1 | 4 |
| RACGAP1 | 0.49 | 1 | 0 | 0 | 1 | 1 | 0 | 3 | 0 |
| SH3D19 | 0.49 | 1 | 1 | 1 | 0 | 2 | 2 | 6 | 4 |
| SNX9 | 0.49 | 3 | 2 | 3 | 2 | 6 | 8 | 8 | 10 |
| STMN1 | 0.49 | 3 | 3 | 4 | 10 | 2 | 7 | 8 | 11 |
| WDR45 | 0.49 | 0 | 0 | 1 | 1 | 1 | 1 | 3 | 1 |
| ALDH3A2 | 0.48 | 2 | 2 | 2 | 3 | 4 | 4 | 10 | 7 |
| AVEN | 0.48 | 0 | 1 | 1 | 0 | 1 | 0 | 3 | 5 |
| CD59 | 0.48 | 7 | 2 | 1 | 7 | 8 | 5 | 0 | 8 |
| DBNL | 0.48 | 3 | 5 | 3 | 2 | 7 | 6 | 16 | 12 |
| DDX1 | 0.48 | 3 | 1 | 4 | 2 | 8 | 5 | 8 | 8 |
| EIF3I | 0.48 | 2 | 2 | 2 | 0 | 5 | 3 | 6 | 10 |
| MAP1A | 0.48 | 0 | 0 | 3 | 1 | 1 | 0 | 5 | 2 |
| MTMR6 | 0.48 | 0 | 1 | 1 | 0 | 0 | 1 | 4 | 4 |
| PEAK1 | 0.48 | 3 | 2 | 5 | 5 | 9 | 9 | 10 | 11 |
| PPP2CA;PPP2CB | 0.48 | 1 | 0 | 0 | 2 | 1 | 1 | 1 | 2 |
| RANBP1 | 0.48 | 0 | 0 | 2 | 1 | 1 | 0 | 3 | 3 |
| RPL17;RPL17-C18... | 0.48 | 2 | 2 | 0 | 0 | 2 | 1 | 6 | 5 |
| SCYL2 | 0.48 | 1 | 0 | 0 | 1 | 1 | 2 | 2 | 1 |
| SEC24A | 0.48 | 0 | 1 | 1 | 2 | 2 | 3 | 1 | 0 |
| TRIP4 | 0.48 | 0 | 0 | 2 | 1 | 1 | 0 | 4 | 2 |
| ABLIM3 | 0.47 | 1 | 0 | 1 | 2 | 0 | 2 | 2 | 4 |
| CISD2 | 0.47 | 0 | 0 | 1 | 0 | 1 | 0 | 3 | 2 |
| DCTN3 | 0.47 | 0 | 0 | 1 | 0 | 1 | 0 | 1 | 4 |
| DNM1L | 0.47 | 3 | 4 | 3 | 8 | 9 | 10 | 2 | 6 |
| EIF3K | 0.47 | 0 | 0 | 1 | 0 | 1 | 0 | 2 | 3 |
| GSPT1;GSPT2 | 0.47 | 0 | 1 | 1 | 0 | 2 | 0 | 3 | 4 |
| ILK | 0.47 | 1 | 0 | 0 | 0 | 0 | 1 | 2 | 3 |
| MAP4K5 | 0.47 | 1 | 0 | 1 | 1 | 0 | 2 | 2 | 5 |
| NFKB2 | 0.47 | 1 | 0 | 1 | 2 | 2 | 3 | 2 | 2 |
| PPP1CB | 0.47 | 3 | 4 | 2 | 8 | 9 | 7 | 4 | 6 |
| S100A10 | 0.47 | 2 | 2 | 0 | 2 | 3 | 1 | 5 | 5 |
| SRP68 | 0.47 | 8 | 8 | 8 | 10 | 21 | 13 | 27 | 26 |
| TCHP | 0.47 | 0 | 0 | 1 | 0 | 0 | 0 | 5 | 1 |
| ARCN1 | 0.46 | 3 | 4 | 4 | 4 | 5 | 8 | 15 | 14 |
| CAPZA2 | 0.46 | 2 | 5 | 4 | 6 | 10 | 7 | 7 | 17 |
| LARP1 | 0.46 | 8 | 10 | 9 | 14 | 16 | 19 | 33 | 34 |
| NAA50 | 0.46 | 1 | 0 | 0 | 1 | 3 | 1 | 0 | 0 |
| PHLDB1 | 0.46 | 1 | 0 | 1 | 1 | 2 | 1 | 4 | 3 |
| PSD3 | 0.46 | 3 | 2 | 5 | 11 | 6 | 5 | 10 | 3 |
| ROCK1 | 0.46 | 0 | 1 | 0 | 0 | 1 | 1 | 4 | 1 |
| RPL26;RPL26L1 | 0.46 | 2 | 0 | 1 | 1 | 4 | 3 | 1 | 3 |

| | | | | | | | | | |
|---|---|---|---|---|---|---|---|---|---|
| TJP1 | 0.46 | 14 | 17 | 15 | 22 | 25 | 23 | 58 | 60 |
| AKAP12 | 0.45 | 6 | 9 | 6 | 10 | 8 | 4 | 32 | 25 |
| C14orf166 | 0.45 | 1 | 1 | 1 | 2 | 4 | 4 | 4 | 4 |
| CAMSAP1 | 0.45 | 5 | 7 | 2 | 3 | 6 | 7 | 18 | 19 |
| CCDC88A | 0.45 | 0 | 0 | 1 | 1 | 0 | 0 | 3 | 2 |
| CMTR1 | 0.45 | 1 | 0 | 0 | 2 | 2 | 1 | 0 | 0 |
| PICALM | 0.45 | 3 | 0 | 4 | 4 | 3 | 4 | 7 | 7 |
| RPS16;ZNF90 | 0.45 | 3 | 1 | 1 | 6 | 3 | 1 | 6 | 4 |
| RTN4 | 0.45 | 7 | 3 | 8 | 14 | 16 | 15 | 15 | 14 |
| TNKS1BP1 | 0.45 | 4 | 5 | 7 | 6 | 8 | 9 | 23 | 21 |
| USP14 | 0.45 | 1 | 1 | 4 | 3 | 4 | 5 | 7 | 8 |
| UTRN | 0.45 | 5 | 8 | 8 | 13 | 13 | 16 | 32 | 19 |
| AHCYL1;AHCYL2 | 0.44 | 1 | 0 | 0 | 1 | 2 | 1 | 1 | 3 |
| ARFGAP2 | 0.44 | 1 | 1 | 0 | 3 | 0 | 0 | 2 | 4 |
| CAP2 | 0.44 | 0 | 0 | 1 | 0 | 0 | 0 | 2 | 5 |
| CFL1 | 0.44 | 8 | 4 | 5 | 11 | 15 | 20 | 6 | 8 |
| DDX50 | 0.44 | 0 | 1 | 0 | 0 | 3 | 1 | 2 | 1 |
| EEF1G | 0.44 | 4 | 9 | 8 | 16 | 17 | 22 | 14 | 17 |
| HNRNPH1 | 0.44 | 1 | 4 | 3 | 5 | 9 | 4 | 10 | 6 |
| IPO5 | 0.44 | 1 | 0 | 0 | 3 | 1 | 1 | 0 | 0 |
| MLLT4 | 0.44 | 10 | 14 | 20 | 25 | 34 | 26 | 50 | 53 |
| SYNE1 | 0.44 | 1 | 0 | 0 | 4 | 0 | 0 | 0 | 1 |
| XIAP | 0.44 | 2 | 1 | 0 | 3 | 1 | 1 | 5 | 3 |
| ATP5B | 0.43 | 0 | 1 | 1 | 0 | 0 | 2 | 4 | 4 |
| CAMSAP2 | 0.43 | 1 | 0 | 1 | 1 | 1 | 0 | 5 | 4 |
| HBS1L | 0.43 | 4 | 2 | 4 | 5 | 8 | 14 | 6 | 11 |
| IPO7 | 0.43 | 3 | 3 | 2 | 8 | 5 | 11 | 4 | 5 |
| PPP2R1A | 0.43 | 1 | 2 | 3 | 7 | 9 | 2 | 2 | 1 |
| RAI14 | 0.43 | 14 | 12 | 18 | 23 | 31 | 28 | 64 | 50 |
| RECQL | 0.43 | 0 | 0 | 1 | 0 | 0 | 0 | 4 | 3 |
| VDAC2 | 0.43 | 4 | 4 | 2 | 6 | 3 | 2 | 15 | 14 |
| ACTR3 | 0.42 | 5 | 6 | 5 | 16 | 11 | 16 | 12 | 19 |
| FCHO2 | 0.42 | 1 | 0 | 1 | 2 | 0 | 2 | 5 | 0 |
| MCM7 | 0.42 | 5 | 1 | 0 | 2 | 4 | 4 | 7 | 6 |
| PPP6R1 | 0.42 | 0 | 0 | 1 | 1 | 1 | 2 | 2 | 3 |
| TJP2 | 0.42 | 7 | 8 | 7 | 4 | 10 | 8 | 33 | 36 |
| ABCF1 | 0.41 | 2 | 1 | 1 | 3 | 1 | 5 | 6 | 7 |
| ARHGAP17 | 0.41 | 0 | 0 | 1 | 1 | 0 | 1 | 1 | 5 |
| ARPC1A | 0.41 | 1 | 2 | 1 | 4 | 6 | 4 | 2 | 6 |
| FAM114A2 | 0.41 | 2 | 1 | 0 | 1 | 3 | 2 | 2 | 8 |
| FHL2 | 0.41 | 4 | 2 | 3 | 1 | 3 | 0 | 14 | 19 |
| GOLGA5 | 0.41 | 3 | 3 | 1 | 3 | 5 | 1 | 11 | 11 |
| HLCS | 0.41 | 0 | 1 | 0 | 0 | 0 | 0 | 3 | 5 |
| LMO7 | 0.41 | 22 | 27 | 23 | 51 | 66 | 59 | 97 | 84 |
| PPP1CA;PPP1CB;P... | 0.41 | 16 | 12 | 11 | 35 | 38 | 38 | 22 | 21 |

| | | | | | | | | | |
|---|---|---|---|---|---|---|---|---|---|
| APBB2 | 0.4 | 1 | 1 | 0 | 2 | 4 | 3 | 1 | 1 |
| ARF1;ARF3 | 0.4 | 2 | 0 | 0 | 0 | 3 | 1 | 3 | 3 |
| DCTN4 | 0.4 | 0 | 0 | 1 | 2 | 0 | 0 | 3 | 2 |
| GTF2A1 | 0.4 | 0 | 0 | 1 | 2 | 2 | 2 | 2 | 2 |
| MAP7D1 | 0.4 | 1 | 0 | 1 | 1 | 0 | 2 | 6 | 3 |
| RAB11FIP5 | 0.4 | 3 | 3 | 3 | 4 | 5 | 7 | 16 | 14 |
| SH2D4A | 0.4 | 0 | 0 | 1 | 2 | 2 | 2 | 1 | 2 |
| YTHDF2 | 0.4 | 0 | 1 | 0 | 0 | 0 | 0 | 4 | 4 |
| ATP1A1;ATP1A3 | 0.39 | 2 | 0 | 1 | 3 | 3 | 2 | 6 | 2 |
| EGLN1 | 0.39 | 0 | 1 | 0 | 2 | 1 | 3 | 1 | 2 |
| PPP1R18 | 0.39 | 4 | 2 | 5 | 5 | 15 | 7 | 13 | 13 |
| RBBP4 | 0.39 | 0 | 0 | 1 | 1 | 2 | 2 | 3 | 2 |
| UBE2M | 0.39 | 0 | 1 | 0 | 2 | 0 | 2 | 3 | 1 |
| CUL4B | 0.38 | 5 | 4 | 8 | 17 | 12 | 12 | 19 | 25 |
| EIF3J | 0.38 | 1 | 0 | 0 | 1 | 1 | 1 | 3 | 4 |
| FLII | 0.38 | 19 | 3 | 0 | 9 | 10 | 11 | 2 | 17 |
| NEK1 | 0.38 | 0 | 0 | 1 | 1 | 1 | 1 | 4 | 3 |
| NFKB1 | 0.38 | 0 | 0 | 1 | 2 | 1 | 2 | 3 | 2 |
| PPME1 | 0.38 | 1 | 0 | 0 | 1 | 1 | 1 | 3 | 4 |
| PTPN11 | 0.38 | 1 | 1 | 3 | 2 | 5 | 6 | 9 | 5 |
| FARSA | 0.37 | 1 | 0 | 1 | 1 | 2 | 3 | 5 | 4 |
| LASP1 | 0.37 | 1 | 2 | 3 | 2 | 3 | 6 | 9 | 12 |
| RPL21 | 0.37 | 1 | 1 | 1 | 2 | 0 | 1 | 7 | 8 |
| RPS3A | 0.37 | 2 | 4 | 3 | 9 | 8 | 8 | 13 | 15 |
| SLC3A2 | 0.37 | 2 | 3 | 2 | 5 | 9 | 11 | 9 | 6 |
| CDC23 | 0.36 | 1 | 0 | 0 | 0 | 1 | 1 | 4 | 4 |
| CLCC1 | 0.36 | 0 | 0 | 1 | 2 | 3 | 2 | 0 | 0 |
| DRG1 | 0.36 | 1 | 1 | 1 | 3 | 1 | 3 | 7 | 7 |
| EPHA2 | 0.36 | 2 | 4 | 6 | 8 | 13 | 13 | 16 | 20 |
| HACD3 | 0.36 | 1 | 0 | 0 | 2 | 4 | 1 | 1 | 1 |
| KNTC1 | 0.36 | 0 | 1 | 1 | 2 | 1 | 1 | 8 | 2 |
| MYO18A | 0.36 | 22 | 0 | 0 | 2 | 5 | 14 | 0 | 0 |
| NUP133 | 0.36 | 0 | 0 | 1 | 1 | 2 | 2 | 4 | 1 |
| PAK2 | 0.36 | 1 | 5 | 2 | 4 | 7 | 9 | 10 | 14 |
| PSMC3 | 0.36 | 0 | 1 | 0 | 3 | 0 | 1 | 3 | 2 |
| ACLY | 0.35 | 1 | 3 | 3 | 6 | 6 | 6 | 13 | 12 |
| BABAM1 | 0.35 | 1 | 0 | 0 | 1 | 2 | 3 | 3 | 3 |
| CDC42 | 0.35 | 2 | 0 | 0 | 2 | 5 | 2 | 1 | 2 |
| CTTNBP2NL | 0.35 | 0 | 0 | 1 | 0 | 0 | 2 | 5 | 2 |
| HMGA1 | 0.35 | 6 | 0 | 0 | 4 | 4 | 5 | 3 | 5 |
| IRAK1 | 0.35 | 1 | 0 | 2 | 2 | 7 | 3 | 1 | 4 |
| KEAP1 | 0.35 | 8 | 4 | 4 | 13 | 11 | 21 | 22 | 23 |
| LPXN | 0.35 | 0 | 0 | 1 | 0 | 2 | 1 | 4 | 3 |
| LRRFIP1 | 0.35 | 1 | 2 | 1 | 1 | 3 | 4 | 10 | 8 |
| PIH1D1 | 0.35 | 0 | 0 | 1 | 1 | 2 | 0 | 4 | 3 |

| | | | | | | | | |
|---|---|---|---|---|---|---|---|---|
| PLAT | 0.35 | 0 | 0 | 1 | 1 | 2 | 1 | 4 | 3 |
| RAB3B | 0.35 | 2 | 0 | 0 | 1 | 2 | 0 | 6 | 3 |
| RIPK2 | 0.35 | 1 | 0 | 0 | 2 | 2 | 2 | 4 | 2 |
| ATP6V1A | 0.34 | 1 | 0 | 4 | 5 | 5 | 6 | 5 | 6 |
| ATXN2L | 0.34 | 5 | 4 | 6 | 9 | 14 | 14 | 27 | 26 |
| COBLL1 | 0.34 | 1 | 0 | 0 | 0 | 1 | 2 | 6 | 0 |
| CTNND1 | 0.34 | 4 | 3 | 3 | 8 | 15 | 12 | 16 | 12 |
| G3BP1 | 0.34 | 2 | 1 | 1 | 2 | 4 | 3 | 7 | 12 |
| NUP98 | 0.34 | 0 | 1 | 0 | 0 | 0 | 0 | 5 | 5 |
| PPFIBP1 | 0.34 | 0 | 2 | 1 | 2 | 4 | 5 | 6 | 2 |
| PPP1R12A | 0.34 | 3 | 5 | 4 | 10 | 13 | 4 | 20 | 22 |
| RPL13 | 0.34 | 1 | 0 | 0 | 3 | 1 | 0 | 4 | 1 |
| TARDBP | 0.34 | 0 | 1 | 0 | 1 | 3 | 1 | 1 | 5 |
| YWHAB | 0.34 | 0 | 1 | 0 | 3 | 0 | 2 | 3 | 1 |
| ABCE1 | 0.33 | 0 | 1 | 1 | 1 | 3 | 0 | 7 | 4 |
| CPSF3 | 0.33 | 0 | 1 | 1 | 4 | 4 | 3 | 4 | 4 |
| CSNK2A1;CSNK2A3 | 0.33 | 2 | 0 | 1 | 4 | 6 | 2 | 1 | 4 |
| DDX21 | 0.33 | 3 | 2 | 0 | 2 | 3 | 2 | 13 | 7 |
| DNMT1 | 0.33 | 1 | 0 | 0 | 3 | 1 | 1 | 3 | 3 |
| HNRNPA2B1 | 0.33 | 0 | 3 | 0 | 1 | 1 | 0 | 5 | 8 |
| LIMD1 | 0.33 | 0 | 1 | 0 | 1 | 1 | 1 | 5 | 4 |
| NEDD1 | 0.33 | 1 | 0 | 0 | 0 | 1 | 0 | 6 | 3 |
| RPS26;RPS26P11 | 0.33 | 0 | 1 | 0 | 1 | 2 | 1 | 4 | 4 |
| TEX2 | 0.33 | 0 | 1 | 0 | 2 | 2 | 1 | 5 | 2 |
| TSN | 0.33 | 0 | 0 | 1 | 1 | 3 | 3 | 2 | 3 |
| VDAC3 | 0.33 | 3 | 0 | 0 | 3 | 0 | 0 | 3 | 7 |
| AP3D1 | 0.32 | 3 | 0 | 1 | 1 | 1 | 0 | 10 | 9 |
| DHX29 | 0.32 | 5 | 5 | 6 | 13 | 20 | 19 | 29 | 28 |
| NARS | 0.32 | 0 | 0 | 1 | 1 | 3 | 1 | 4 | 3 |
| NCKAP1 | 0.32 | 1 | 4 | 2 | 9 | 10 | 5 | 10 | 9 |
| PHLDB2 | 0.32 | 0 | 1 | 0 | 1 | 1 | 1 | 8 | 1 |
| SSB | 0.32 | 1 | 0 | 0 | 1 | 3 | 1 | 3 | 4 |
| UFD1L | 0.32 | 1 | 0 | 0 | 0 | 0 | 0 | 5 | 6 |
| YKT6 | 0.32 | 0 | 1 | 2 | 3 | 2 | 5 | 6 | 6 |
| EIF3L | 0.31 | 3 | 1 | 0 | 7 | 6 | 2 | 4 | 3 |
| HNRNPL | 0.31 | 2 | 0 | 1 | 2 | 0 | 1 | 10 | 6 |
| PCNP | 0.31 | 2 | 2 | 1 | 5 | 4 | 6 | 12 | 11 |
| PPP6R3 | 0.31 | 0 | 2 | 0 | 3 | 3 | 3 | 3 | 6 |
| ZC3H15 | 0.31 | 0 | 1 | 0 | 1 | 2 | 2 | 2 | 7 |
| ANKRD17 | 0.3 | 2 | 3 | 1 | 7 | 5 | 6 | 15 | 10 |
| ARHGAP21 | 0.3 | 1 | 1 | 1 | 1 | 3 | 2 | 12 | 7 |
| BUB1B | 0.3 | 0 | 0 | 1 | 2 | 2 | 2 | 6 | 2 |
| ESYT2 | 0.3 | 1 | 1 | 1 | 1 | 5 | 4 | 9 | 8 |
| SERBP1 | 0.3 | 5 | 4 | 5 | 15 | 17 | 16 | 25 | 29 |
| SERPINB2 | 0.3 | 1 | 3 | 6 | 7 | 9 | 12 | 17 | 17 |

| | | | | | | | | | |
|---|---|---|---|---|---|---|---|---|---|
| SMC4 | 0.3 | 3 | 0 | 0 | 0 | 5 | 0 | 6 | 4 |
| DLGAP5 | 0.29 | 1 | 3 | 2 | 4 | 6 | 3 | 17 | 12 |
| DST;MACF1 | 0.29 | 0 | 0 | 1 | 2 | 2 | 2 | 5 | 4 |
| ECD | 0.29 | 8 | 1 | 1 | 10 | 10 | 13 | 7 | 6 |
| PALLD | 0.29 | 3 | 1 | 1 | 4 | 4 | 3 | 14 | 12 |
| RPL10 | 0.29 | 0 | 1 | 1 | 0 | 0 | 1 | 8 | 8 |
| TPR | 0.29 | 10 | 10 | 10 | 14 | 34 | 13 | 68 | 49 |
| VARS | 0.29 | 1 | 2 | 0 | 4 | 5 | 5 | 7 | 8 |
| ALDH18A1 | 0.28 | 0 | 0 | 1 | 0 | 0 | 0 | 7 | 6 |
| DST | 0.28 | 1 | 5 | 5 | 3 | 7 | 7 | 24 | 25 |
| MYO1E | 0.28 | 0 | 1 | 0 | 0 | 1 | 0 | 5 | 7 |
| NR3C1 | 0.28 | 3 | 0 | 1 | 3 | 2 | 1 | 10 | 9 |
| TUFM | 0.28 | 2 | 1 | 2 | 4 | 3 | 3 | 13 | 15 |
| TWF2 | 0.28 | 4 | 2 | 4 | 15 | 18 | 12 | 9 | 13 |
| VAPA | 0.28 | 1 | 2 | 1 | 7 | 4 | 5 | 7 | 12 |
| ASCC2 | 0.27 | 0 | 0 | 1 | 1 | 3 | 3 | 4 | 6 |
| EPB41L2 | 0.27 | 6 | 4 | 10 | 19 | 29 | 27 | 40 | 38 |
| FN1 | 0.27 | 0 | 0 | 2 | 0 | 0 | 0 | 8 | 8 |
| JUP | 0.27 | 3 | 2 | 2 | 9 | 12 | 10 | 9 | 19 |
| PPP1CC | 0.27 | 0 | 0 | 1 | 3 | 4 | 3 | 2 | 3 |
| RANBP2 | 0.27 | 13 | 11 | 15 | 24 | 32 | 28 | 94 | 85 |
| WDR44 | 0.27 | 0 | 1 | 1 | 2 | 3 | 2 | 5 | 10 |
| ARHGEF10 | 0.26 | 1 | 1 | 1 | 7 | 9 | 4 | 4 | 5 |
| COPB2 | 0.26 | 9 | 1 | 0 | 13 | 8 | 6 | 8 | 7 |
| HNRNPA1;HNRNPA1L2 | 0.26 | 2 | 1 | 1 | 0 | 6 | 1 | 13 | 11 |
| HNRNPK | 0.26 | 5 | 3 | 6 | 16 | 25 | 26 | 22 | 19 |
| HNRNPM | 0.26 | 2 | 1 | 0 | 3 | 2 | 2 | 9 | 10 |
| NCAPG | 0.26 | 1 | 0 | 1 | 3 | 1 | 2 | 8 | 7 |
| RDX | 0.26 | 0 | 0 | 3 | 5 | 2 | 2 | 3 | 9 |
| SWAP70 | 0.26 | 1 | 0 | 2 | 5 | 6 | 5 | 8 | 7 |
| EDC4 | 0.25 | 0 | 0 | 2 | 5 | 2 | 3 | 6 | 3 |
| EIF2S3;EIF2S3L | 0.25 | 0 | 1 | 0 | 3 | 3 | 4 | 2 | 6 |
| NPLOC4 | 0.25 | 0 | 0 | 1 | 1 | 1 | 0 | 6 | 7 |
| PEX14 | 0.25 | 0 | 0 | 1 | 3 | 4 | 4 | 5 | 0 |
| SCFD1 | 0.25 | 0 | 1 | 0 | 0 | 3 | 0 | 7 | 4 |
| SYNPO | 0.25 | 0 | 0 | 1 | 2 | 3 | 5 | 4 | 5 |
| TPM1 | 0.25 | 2 | 0 | 0 | 6 | 3 | 4 | 1 | 2 |
| TPM3 | 0.25 | 7 | 1 | 1 | 18 | 7 | 11 | 1 | 7 |
| UBAP2L | 0.25 | 3 | 4 | 3 | 10 | 12 | 12 | 25 | 30 |
| DDX5 | 0.24 | 2 | 1 | 1 | 8 | 9 | 6 | 12 | 11 |
| EPS15L1 | 0.24 | 1 | 2 | 3 | 6 | 9 | 8 | 16 | 18 |
| UGDH | 0.24 | 0 | 1 | 2 | 2 | 6 | 5 | 4 | 12 |
| BASP1 | 0.23 | 1 | 0 | 0 | 1 | 3 | 2 | 6 | 7 |
| CD44 | 0.23 | 3 | 0 | 0 | 6 | 6 | 5 | 1 | 6 |
| LEMD3 | 0.23 | 0 | 1 | 0 | 2 | 1 | 1 | 7 | 7 |

| | | | | | | | | | |
|---|---|---|---|---|---|---|---|---|---|
| ZC3H4 | 0.23 | 0 | 1 | 1 | 3 | 5 | 5 | 9 | 8 |
| ZC3H7A | 0.23 | 1 | 1 | 0 | 2 | 4 | 4 | 11 | 7 |
| ATXN2 | 0.22 | 1 | 0 | 0 | 1 | 2 | 4 | 6 | 6 |
| ESYT1 | 0.22 | 6 | 7 | 4 | 27 | 38 | 29 | 27 | 25 |
| FGD4 | 0.22 | 2 | 0 | 1 | 2 | 5 | 1 | 13 | 8 |
| HNRNPU | 0.22 | 2 | 2 | 2 | 9 | 11 | 6 | 20 | 19 |
| LGALS1 | 0.22 | 1 | 0 | 0 | 1 | 3 | 3 | 5 | 9 |
| LIMA1 | 0.22 | 8 | 8 | 4 | 33 | 32 | 42 | 31 | 45 |
| MYLK | 0.22 | 4 | 5 | 6 | 18 | 22 | 24 | 39 | 45 |
| PRMT7 | 0.22 | 0 | 0 | 1 | 1 | 3 | 1 | 4 | 10 |
| RPAP3 | 0.22 | 1 | 0 | 1 | 1 | 2 | 2 | 7 | 15 |
| TACC1 | 0.22 | 1 | 1 | 1 | 6 | 5 | 4 | 15 | 8 |
| CALM2;CALM1;CALM3 | 0.21 | 1 | 1 | 3 | 13 | 9 | 11 | 2 | 4 |
| GNB2L1 | 0.21 | 0 | 0 | 1 | 1 | 1 | 0 | 8 | 9 |
| LBR | 0.21 | 1 | 0 | 1 | 3 | 6 | 3 | 9 | 8 |
| RANBP3 | 0.21 | 0 | 2 | 4 | 8 | 6 | 8 | 17 | 13 |
| TNRC6B | 0.21 | 1 | 0 | 1 | 2 | 1 | 1 | 12 | 10 |
| COPA | 0.2 | 14 | 2 | 0 | 23 | 18 | 10 | 4 | 7 |
| FLOT1 | 0.2 | 1 | 0 | 0 | 0 | 2 | 0 | 9 | 8 |
| GNB1 | 0.2 | 1 | 0 | 0 | 4 | 3 | 6 | 0 | 6 |
| PTPN14 | 0.2 | 1 | 0 | 0 | 1 | 1 | 1 | 10 | 8 |
| SYNM | 0.2 | 1 | 0 | 1 | 4 | 4 | 4 | 13 | 7 |
| UBAP2 | 0.2 | 0 | 3 | 1 | 2 | 4 | 3 | 15 | 15 |
| DSG2 | 0.19 | 4 | 4 | 3 | 12 | 14 | 15 | 38 | 42 |
| TP53BP2 | 0.19 | 0 | 1 | 0 | 3 | 3 | 1 | 8 | 9 |
| DLG5 | 0.18 | 3 | 4 | 3 | 15 | 17 | 11 | 38 | 36 |
| ERBB2IP | 0.18 | 4 | 4 | 2 | 15 | 19 | 14 | 40 | 30 |
| GNAI2 | 0.18 | 4 | 0 | 0 | 2 | 6 | 1 | 0 | 22 |
| TBPL1 | 0.18 | 1 | 0 | 0 | 4 | 7 | 5 | 7 | 5 |
| TWF1 | 0.18 | 2 | 0 | 2 | 15 | 8 | 8 | 5 | 8 |
| CFL2 | 0.17 | 1 | 0 | 0 | 7 | 5 | 6 | 0 | 0 |
| EFTUD2 | 0.17 | 0 | 1 | 0 | 6 | 5 | 3 | 8 | 1 |
| EGFR | 0.17 | 2 | 0 | 2 | 6 | 9 | 7 | 17 | 14 |
| GNB2 | 0.17 | 1 | 0 | 0 | 4 | 3 | 2 | 1 | 14 |
| SSH1 | 0.17 | 0 | 0 | 1 | 0 | 2 | 4 | 10 | 8 |
| MPRIP | 0.16 | 5 | 1 | 5 | 22 | 29 | 28 | 20 | 38 |
| PLEKHA5 | 0.16 | 1 | 1 | 2 | 6 | 11 | 6 | 24 | 16 |
| RRBP1 | 0.16 | 1 | 1 | 4 | 1 | 2 | 1 | 35 | 27 |
| TMOD3 | 0.16 | 12 | 1 | 2 | 23 | 28 | 31 | 2 | 23 |
| KRT18 | 0.15 | 2 | 1 | 1 | 7 | 13 | 14 | 20 | 22 |
| NONO | 0.15 | 2 | 3 | 1 | 6 | 7 | 9 | 33 | 30 |
| NT5E | 0.15 | 3 | 0 | 0 | 5 | 5 | 10 | 0 | 17 |
| PRRC2C | 0.15 | 2 | 0 | 4 | 5 | 7 | 7 | 29 | 24 |
| CORO2B | 0.14 | 2 | 0 | 0 | 5 | 4 | 10 | 0 | 16 |
| EIF5B | 0.14 | 0 | 0 | 1 | 4 | 4 | 3 | 12 | 10 |

| | | | | | | | | | |
|---|---|---|---|---|---|---|---|---|---|
| LUZP1 | 0.14 | 3 | 2 | 3 | 8 | 19 | 11 | 44 | 40 |
| MYO6 | 0.14 | 3 | 2 | 6 | 23 | 17 | 27 | 0 | 64 |
| ZBTB33 | 0.14 | 0 | 1 | 0 | 2 | 2 | 1 | 15 | 12 |
| ACTC1;ACTA1 | 0.11 | 3 | 0 | 0 | 6 | 7 | 18 | 1 | 18 |
| AKAP2;PALM2-AKA... | 0.11 | 1 | 0 | 0 | 4 | 7 | 3 | 16 | 14 |
| ITGB1 | 0.11 | 0 | 0 | 2 | 4 | 5 | 13 | 15 | 13 |
| MYH10 | 0.11 | 147 | 1 | 3 | 82 | 65 | 95 | 0 | 160 |
| HCFC1 | 0.1 | 1 | 2 | 0 | 12 | 11 | 10 | 25 | 27 |
| PUF60 | 0.1 | 0 | 0 | 1 | 6 | 9 | 4 | 17 | 15 |
| SF3A1 | 0.1 | 0 | 1 | 0 | 1 | 8 | 5 | 17 | 14 |
| WDR1 | 0.1 | 3 | 0 | 0 | 11 | 7 | 23 | 0 | 2 |
| DBN1 | 0.09 | 1 | 3 | 1 | 29 | 23 | 29 | 15 | 39 |
| DLG1 | 0.09 | 0 | 0 | 1 | 11 | 9 | 13 | 13 | 11 |
| NPM1 | 0.09 | 1 | 0 | 0 | 9 | 13 | 9 | 15 | 15 |
| SF3B2 | 0.09 | 0 | 2 | 1 | 6 | 10 | 9 | 29 | 30 |
| ACTB;ACTG1 | 0.08 | 76 | 13 | 19 | 369 | 393 | 479 | 37 | 188 |
| SFPQ | 0.08 | 4 | 0 | 0 | 14 | 20 | 15 | 29 | 27 |
| SMARCA5 | 0.08 | 1 | 0 | 0 | 8 | 7 | 4 | 21 | 20 |
| SPTBN1 | 0.08 | 6 | 5 | 9 | 81 | 86 | 119 | 15 | 129 |
| SF3B1 | 0.07 | 1 | 1 | 0 | 9 | 17 | 14 | 34 | 32 |
| ACTN1 | 0.04 | 1 | 0 | 0 | 17 | 30 | 36 | 14 | 46 |
| ACTN4 | 0.04 | 4 | 3 | 5 | 96 | 112 | 150 | 31 | 87 |
| CHD4 | 0.04 | 0 | 1 | 0 | 16 | 10 | 10 | 38 | 42 |
| MYO1C | 0.04 | 12 | 0 | 0 | 49 | 58 | 33 | 0 | 60 |
| POTEJ | 0.04 | 1 | 0 | 0 | 18 | 40 | 35 | 0 | 0 |
| SPTAN1 | 0.04 | 3 | 3 | 4 | 101 | 90 | 122 | 18 | 154 |
| MKI67 | 0.03 | 4 | 0 | 0 | 24 | 38 | 29 | 74 | 70 |
| ABCC1 | 0 | 0 | 0 | 0 | 0 | 0 | 0 | 0 | 1 |
| ABLIM1 | 0 | 0 | 0 | 0 | 1 | 1 | 2 | 0 | 4 |
| ABR | 0 | 0 | 0 | 0 | 0 | 0 | 0 | 2 | 1 |
| ACAP2 | 0 | 0 | 0 | 0 | 0 | 0 | 0 | 0 | 1 |
| ACBD3 | 0 | 0 | 0 | 0 | 0 | 1 | 1 | 3 | 3 |
| ACBD5 | 0 | 0 | 0 | 0 | 0 | 0 | 0 | 1 | 0 |
| ACIN1 | 0 | 0 | 0 | 0 | 4 | 10 | 9 | 23 | 24 |
| ACOT9 | 0 | 0 | 0 | 0 | 0 | 0 | 0 | 3 | 2 |
| ACSL3 | 0 | 0 | 0 | 0 | 1 | 1 | 0 | 3 | 3 |
| ACTB | 0 | 0 | 0 | 0 | 1 | 2 | 2 | 2 | 3 |
| ACTBL2 | 0 | 0 | 0 | 0 | 1 | 1 | 3 | 0 | 0 |
| ACTR10 | 0 | 0 | 0 | 0 | 0 | 0 | 0 | 0 | 2 |
| ACTR1B | 0 | 0 | 0 | 0 | 0 | 0 | 0 | 3 | 1 |
| ADAR | 0 | 0 | 0 | 0 | 2 | 2 | 4 | 9 | 4 |
| ADD3 | 0 | 0 | 0 | 0 | 0 | 0 | 0 | 1 | 2 |
| ADNP | 0 | 0 | 0 | 0 | 9 | 12 | 6 | 20 | 13 |
| ADRM1 | 0 | 0 | 0 | 0 | 0 | 0 | 0 | 1 | 1 |
| AFAP1 | 0 | 0 | 0 | 0 | 4 | 5 | 8 | 9 | 7 |

| | | | | | | | | | |
|---|---|---|---|---|---|---|---|---|---|
| AFAP1L2 | 0 | 0 | 0 | 0 | 1 | 1 | 0 | 0 | 0 |
| AFTPH | 0 | 0 | 0 | 0 | 0 | 0 | 1 | 1 | 2 |
| AGPAT5 | 0 | 0 | 0 | 0 | 0 | 0 | 0 | 1 | 0 |
| AGPAT9 | 0 | 0 | 0 | 0 | 1 | 2 | 1 | 5 | 1 |
| AGTPBP1 | 0 | 0 | 0 | 0 | 0 | 0 | 0 | 1 | 1 |
| AHCTF1 | 0 | 0 | 0 | 0 | 0 | 0 | 0 | 8 | 4 |
| AHCY | 0 | 0 | 0 | 0 | 0 | 1 | 0 | 0 | 0 |
| AHDC1 | 0 | 0 | 0 | 0 | 0 | 0 | 0 | 1 | 0 |
| AHR | 0 | 0 | 0 | 0 | 0 | 0 | 0 | 0 | 1 |
| AIFM1 | 0 | 0 | 0 | 0 | 0 | 0 | 0 | 3 | 1 |
| AKAP1 | 0 | 0 | 0 | 0 | 1 | 2 | 2 | 3 | 3 |
| AKAP11 | 0 | 0 | 0 | 0 | 1 | 0 | 1 | 2 | 0 |
| AKAP13 | 0 | 0 | 0 | 0 | 0 | 0 | 0 | 1 | 0 |
| AKAP8 | 0 | 0 | 0 | 0 | 1 | 1 | 1 | 1 | 0 |
| ALDOA | 0 | 0 | 0 | 0 | 0 | 0 | 1 | 2 | 5 |
| ALMS1 | 0 | 0 | 0 | 0 | 0 | 0 | 0 | 2 | 0 |
| ANAPC1 | 0 | 0 | 0 | 0 | 1 | 1 | 1 | 2 | 2 |
| ANAPC4 | 0 | 0 | 0 | 0 | 0 | 1 | 1 | 2 | 0 |
| ANK3 | 0 | 0 | 0 | 0 | 0 | 0 | 0 | 1 | 0 |
| ANKFY1 | 0 | 0 | 0 | 0 | 1 | 1 | 1 | 3 | 3 |
| ANKHD1 | 0 | 0 | 0 | 0 | 0 | 0 | 0 | 2 | 1 |
| ANKLE2 | 0 | 0 | 0 | 0 | 0 | 0 | 0 | 5 | 4 |
| ANKRD28 | 0 | 0 | 0 | 0 | 1 | 0 | 1 | 0 | 0 |
| ANKRD50 | 0 | 0 | 0 | 0 | 0 | 0 | 0 | 4 | 1 |
| ANKS1A | 0 | 0 | 0 | 0 | 1 | 2 | 2 | 3 | 3 |
| ANLN | 0 | 0 | 0 | 0 | 5 | 6 | 3 | 16 | 12 |
| ANO10 | 0 | 0 | 0 | 0 | 0 | 0 | 0 | 0 | 1 |
| ANP32E | 0 | 0 | 0 | 0 | 0 | 0 | 1 | 0 | 0 |
| ANXA6 | 0 | 0 | 0 | 0 | 0 | 0 | 0 | 0 | 2 |
| AP1M1 | 0 | 0 | 0 | 0 | 0 | 0 | 0 | 0 | 1 |
| AP2A2 | 0 | 0 | 0 | 0 | 1 | 3 | 2 | 0 | 0 |
| AP4B1 | 0 | 0 | 0 | 0 | 0 | 0 | 0 | 0 | 1 |
| AP4E1 | 0 | 0 | 0 | 0 | 0 | 0 | 0 | 1 | 0 |
| AP4S1 | 0 | 0 | 0 | 0 | 0 | 0 | 0 | 0 | 1 |
| API5 | 0 | 0 | 0 | 0 | 0 | 0 | 0 | 2 | 0 |
| APPL2 | 0 | 0 | 0 | 0 | 0 | 0 | 0 | 1 | 0 |
| ARF5 | 0 | 0 | 0 | 0 | 0 | 0 | 0 | 1 | 0 |
| ARF6 | 0 | 0 | 0 | 0 | 0 | 0 | 0 | 1 | 3 |
| ARHGAP12 | 0 | 0 | 0 | 0 | 0 | 0 | 0 | 1 | 3 |
| ARHGEF12 | 0 | 0 | 0 | 0 | 0 | 0 | 0 | 1 | 0 |
| ARHGEF2 | 0 | 0 | 0 | 0 | 1 | 0 | 1 | 1 | 1 |
| ARHGEF28 | 0 | 0 | 0 | 0 | 0 | 0 | 0 | 2 | 1 |
| ARHGEF7 | 0 | 0 | 0 | 0 | 1 | 1 | 2 | 2 | 1 |
| ARID1A | 0 | 0 | 0 | 0 | 1 | 2 | 1 | 4 | 1 |
| ARL6IP4 | 0 | 0 | 0 | 0 | 0 | 1 | 0 | 0 | 1 |

| | | | | | | | | | |
|---|---|---|---|---|---|---|---|---|---|
| ARL6IP5 | 0 | 0 | 0 | 0 | 0 | 0 | 0 | 3 | 1 |
| ARMC9 | 0 | 0 | 0 | 0 | 1 | 0 | 0 | 0 | 0 |
| ARPC5L | 0 | 0 | 0 | 0 | 0 | 1 | 2 | 0 | 1 |
| ATG2A | 0 | 0 | 0 | 0 | 0 | 0 | 0 | 1 | 0 |
| ATG3 | 0 | 0 | 0 | 0 | 0 | 0 | 0 | 2 | 2 |
| ATP2B1 | 0 | 0 | 0 | 0 | 0 | 1 | 1 | 2 | 2 |
| ATP2C1 | 0 | 0 | 0 | 0 | 0 | 0 | 0 | 0 | 1 |
| ATP5J2 | 0 | 0 | 0 | 0 | 0 | 0 | 0 | 0 | 1 |
| ATP6V1B2 | 0 | 0 | 0 | 0 | 1 | 3 | 0 | 1 | 2 |
| ATP6V1G1 | 0 | 0 | 0 | 0 | 0 | 0 | 0 | 1 | 2 |
| ATRX | 0 | 0 | 0 | 0 | 0 | 0 | 1 | 11 | 8 |
| ATXN10 | 0 | 0 | 0 | 0 | 0 | 0 | 0 | 1 | 1 |
| AUP1 | 0 | 0 | 0 | 0 | 0 | 0 | 0 | 1 | 1 |
| BAP1 | 0 | 0 | 0 | 0 | 0 | 0 | 0 | 1 | 1 |
| BAZ1A | 0 | 0 | 0 | 0 | 0 | 0 | 0 | 3 | 0 |
| BAZ1B | 0 | 0 | 0 | 0 | 2 | 6 | 4 | 24 | 18 |
| BBX | 0 | 0 | 0 | 0 | 0 | 0 | 0 | 7 | 1 |
| BCAP29 | 0 | 0 | 0 | 0 | 0 | 0 | 0 | 1 | 0 |
| BCAP31 | 0 | 0 | 0 | 0 | 0 | 0 | 0 | 0 | 1 |
| BCAR1 | 0 | 0 | 0 | 0 | 0 | 0 | 0 | 0 | 1 |
| BCAR3 | 0 | 0 | 0 | 0 | 0 | 0 | 0 | 1 | 1 |
| BCAS2 | 0 | 0 | 0 | 0 | 0 | 0 | 0 | 1 | 1 |
| BCCIP | 0 | 0 | 0 | 0 | 0 | 0 | 0 | 2 | 0 |
| BCL2L13 | 0 | 0 | 0 | 0 | 1 | 1 | 1 | 1 | 0 |
| BCL9L | 0 | 0 | 0 | 0 | 0 | 0 | 0 | 2 | 2 |
| BCLAF1 | 0 | 0 | 0 | 0 | 7 | 9 | 4 | 17 | 23 |
| BCR | 0 | 0 | 0 | 0 | 0 | 0 | 0 | 1 | 0 |
| BET1 | 0 | 0 | 0 | 0 | 0 | 0 | 0 | 1 | 1 |
| BICC1 | 0 | 0 | 0 | 0 | 0 | 0 | 0 | 0 | 2 |
| BICD2 | 0 | 0 | 0 | 0 | 2 | 0 | 0 | 6 | 4 |
| BLM | 0 | 0 | 0 | 0 | 1 | 2 | 0 | 0 | 1 |
| BLVRA | 0 | 0 | 0 | 0 | 0 | 0 | 0 | 1 | 0 |
| BOD1L1 | 0 | 0 | 0 | 0 | 0 | 0 | 1 | 6 | 4 |
| BRAP | 0 | 0 | 0 | 0 | 1 | 0 | 0 | 10 | 6 |
| BRD4 | 0 | 0 | 0 | 0 | 2 | 4 | 4 | 14 | 15 |
| BRIP1 | 0 | 0 | 0 | 0 | 2 | 0 | 0 | 15 | 4 |
| BRK1 | 0 | 0 | 0 | 0 | 1 | 1 | 1 | 1 | 2 |
| BRMS1L | 0 | 0 | 0 | 0 | 0 | 0 | 0 | 3 | 2 |
| BTF3 | 0 | 0 | 0 | 0 | 0 | 0 | 0 | 0 | 1 |
| BUB3 | 0 | 0 | 0 | 0 | 1 | 2 | 1 | 4 | 3 |
| C10orf88 | 0 | 0 | 0 | 0 | 0 | 0 | 0 | 2 | 0 |
| C11orf58;SMAP | 0 | 0 | 0 | 0 | 0 | 0 | 0 | 4 | 3 |
| C15orf52 | 0 | 0 | 0 | 0 | 0 | 0 | 0 | 2 | 2 |
| C17orf85 | 0 | 0 | 0 | 0 | 0 | 0 | 0 | 1 | 0 |
| C18orf21 | 0 | 0 | 0 | 0 | 0 | 0 | 0 | 0 | 1 |

| | | | | | | | | | |
|---|---|---|---|---|---|---|---|---|---|
| C19orf43 | 0 | 0 | 0 | 0 | 5 | 3 | 5 | 8 | 10 |
| C1QBP | 0 | 0 | 0 | 0 | 4 | 1 | 2 | 7 | 5 |
| C3orf17 | 0 | 0 | 0 | 0 | 0 | 0 | 0 | 1 | 0 |
| C7orf55-LUC7L2;... | 0 | 0 | 0 | 0 | 0 | 0 | 0 | 1 | 0 |
| CACNA2D1 | 0 | 0 | 0 | 0 | 1 | 1 | 0 | 0 | 9 |
| CACTIN | 0 | 0 | 0 | 0 | 0 | 1 | 1 | 1 | 1 |
| CACUL1 | 0 | 0 | 0 | 0 | 0 | 0 | 0 | 1 | 0 |
| CAPN2 | 0 | 0 | 0 | 0 | 0 | 0 | 0 | 0 | 1 |
| CAPNS1 | 0 | 0 | 0 | 0 | 0 | 0 | 0 | 0 | 1 |
| CAPRIN1 | 0 | 0 | 0 | 0 | 2 | 0 | 1 | 2 | 6 |
| CAPZB | 0 | 0 | 0 | 0 | 0 | 0 | 1 | 1 | 1 |
| CARS | 0 | 0 | 0 | 0 | 0 | 0 | 0 | 1 | 0 |
| CASC5 | 0 | 0 | 0 | 0 | 1 | 4 | 0 | 11 | 8 |
| CASK | 0 | 0 | 0 | 0 | 0 | 1 | 0 | 0 | 0 |
| CASP4;CASP12;CASP5 | 0 | 0 | 0 | 0 | 0 | 1 | 1 | 1 | 0 |
| CAT | 0 | 0 | 0 | 0 | 0 | 0 | 0 | 0 | 1 |
| CAV1 | 0 | 0 | 0 | 0 | 0 | 0 | 0 | 3 | 6 |
| CBX3 | 0 | 0 | 0 | 0 | 1 | 0 | 0 | 2 | 2 |
| CC2D1A | 0 | 0 | 0 | 0 | 0 | 1 | 0 | 0 | 0 |
| CCAR1 | 0 | 0 | 0 | 0 | 0 | 1 | 0 | 1 | 1 |
| CCAR2 | 0 | 0 | 0 | 0 | 4 | 3 | 4 | 7 | 8 |
| CCDC115 | 0 | 0 | 0 | 0 | 0 | 1 | 0 | 0 | 1 |
| CCDC43 | 0 | 0 | 0 | 0 | 0 | 1 | 0 | 2 | 1 |
| CCDC47 | 0 | 0 | 0 | 0 | 0 | 1 | 0 | 3 | 0 |
| CCDC55;NSRP1 | 0 | 0 | 0 | 0 | 0 | 0 | 0 | 1 | 2 |
| CCDC85C | 0 | 0 | 0 | 0 | 1 | 0 | 1 | 3 | 1 |
| CCDC94 | 0 | 0 | 0 | 0 | 0 | 0 | 0 | 1 | 0 |
| CCNH | 0 | 0 | 0 | 0 | 0 | 0 | 0 | 0 | 1 |
| CCNK | 0 | 0 | 0 | 0 | 2 | 3 | 1 | 4 | 7 |
| CCNL1 | 0 | 0 | 0 | 0 | 2 | 1 | 2 | 2 | 0 |
| CCNL2 | 0 | 0 | 0 | 0 | 0 | 2 | 0 | 1 | 0 |
| CCNT1 | 0 | 0 | 0 | 0 | 2 | 4 | 6 | 7 | 6 |
| CCNT2 | 0 | 0 | 0 | 0 | 0 | 0 | 0 | 1 | 1 |
| CCS | 0 | 0 | 0 | 0 | 0 | 0 | 1 | 0 | 0 |
| CCSER2 | 0 | 0 | 0 | 0 | 0 | 0 | 0 | 3 | 2 |
| CD151 | 0 | 0 | 0 | 0 | 0 | 0 | 0 | 2 | 1 |
| CD55 | 0 | 0 | 0 | 0 | 0 | 0 | 0 | 0 | 1 |
| CDC123 | 0 | 0 | 0 | 0 | 1 | 0 | 0 | 1 | 0 |
| CDC27 | 0 | 0 | 0 | 0 | 0 | 0 | 1 | 3 | 3 |
| CDC42BPA | 0 | 0 | 0 | 0 | 0 | 0 | 0 | 0 | 2 |
| CDC5L | 0 | 0 | 0 | 0 | 0 | 0 | 0 | 1 | 2 |
| CDC73 | 0 | 0 | 0 | 0 | 0 | 0 | 0 | 1 | 1 |
| CDCA2 | 0 | 0 | 0 | 0 | 0 | 0 | 0 | 3 | 0 |
| CDK11B;CDC2L1;CDK11A | 0 | 0 | 0 | 0 | 5 | 4 | 3 | 8 | 10 |
| CDK12 | 0 | 0 | 0 | 0 | 0 | 1 | 1 | 4 | 2 |

| | | | | | | | | | |
|---|---|---|---|---|---|---|---|---|---|
| CDK13 | 0 | 0 | 0 | 0 | 0 | 0 | 1 | 1 | 0 |
| CDK2;CDK3 | 0 | 0 | 0 | 0 | 1 | 0 | 1 | 0 | 1 |
| CDK7 | 0 | 0 | 0 | 0 | 0 | 0 | 0 | 2 | 0 |
| CDK9 | 0 | 0 | 0 | 0 | 2 | 2 | 0 | 4 | 3 |
| CDKAL1 | 0 | 0 | 0 | 0 | 0 | 0 | 0 | 2 | 1 |
| CDKN2AIP | 0 | 0 | 0 | 0 | 1 | 0 | 1 | 6 | 7 |
| CENPT | 0 | 0 | 0 | 0 | 0 | 0 | 0 | 1 | 0 |
| CEP170B | 0 | 0 | 0 | 0 | 0 | 0 | 1 | 2 | 0 |
| CEP192 | 0 | 0 | 0 | 0 | 0 | 0 | 0 | 4 | 0 |
| CFAP20 | 0 | 0 | 0 | 0 | 1 | 0 | 0 | 1 | 0 |
| CFAP97 | 0 | 0 | 0 | 0 | 0 | 0 | 0 | 1 | 1 |
| CFDP1 | 0 | 0 | 0 | 0 | 1 | 0 | 0 | 2 | 2 |
| CHAMP1 | 0 | 0 | 0 | 0 | 5 | 4 | 5 | 5 | 5 |
| CHD5 | 0 | 0 | 0 | 0 | 0 | 1 | 0 | 1 | 0 |
| CHD8 | 0 | 0 | 0 | 0 | 0 | 1 | 0 | 5 | 3 |
| CHERP | 0 | 0 | 0 | 0 | 5 | 5 | 7 | 8 | 6 |
| CHMP2B | 0 | 0 | 0 | 0 | 0 | 0 | 0 | 2 | 4 |
| CHMP7 | 0 | 0 | 0 | 0 | 1 | 0 | 1 | 0 | 0 |
| CIC | 0 | 0 | 0 | 0 | 0 | 1 | 1 | 3 | 2 |
| CISD1 | 0 | 0 | 0 | 0 | 0 | 0 | 0 | 1 | 0 |
| CIT | 0 | 0 | 0 | 0 | 0 | 0 | 1 | 0 | 1 |
| CIZ1 | 0 | 0 | 0 | 0 | 0 | 0 | 0 | 3 | 2 |
| CLIC4 | 0 | 0 | 0 | 0 | 0 | 0 | 1 | 0 | 0 |
| CLIP1 | 0 | 0 | 0 | 0 | 0 | 0 | 0 | 1 | 0 |
| CLK3 | 0 | 0 | 0 | 0 | 0 | 1 | 0 | 0 | 0 |
| CLMN | 0 | 0 | 0 | 0 | 0 | 0 | 0 | 2 | 0 |
| CNN3 | 0 | 0 | 0 | 0 | 1 | 1 | 2 | 6 | 7 |
| CNOT10 | 0 | 0 | 0 | 0 | 0 | 0 | 0 | 3 | 2 |
| CNOT11 | 0 | 0 | 0 | 0 | 0 | 0 | 0 | 1 | 0 |
| COG1 | 0 | 0 | 0 | 0 | 0 | 0 | 0 | 1 | 1 |
| COG7 | 0 | 0 | 0 | 0 | 0 | 1 | 0 | 1 | 0 |
| COIL | 0 | 0 | 0 | 0 | 0 | 0 | 0 | 2 | 1 |
| COMMD3 | 0 | 0 | 0 | 0 | 1 | 0 | 0 | 0 | 2 |
| COPE | 0 | 0 | 0 | 0 | 2 | 0 | 0 | 0 | 0 |
| COPS2 | 0 | 0 | 0 | 0 | 0 | 0 | 0 | 2 | 3 |
| COPS4 | 0 | 0 | 0 | 0 | 0 | 0 | 0 | 2 | 2 |
| COPS5 | 0 | 0 | 0 | 0 | 0 | 0 | 0 | 0 | 2 |
| COPS6 | 0 | 0 | 0 | 0 | 1 | 0 | 1 | 0 | 3 |
| COPS7A | 0 | 0 | 0 | 0 | 0 | 0 | 0 | 1 | 1 |
| CORO1A | 0 | 0 | 0 | 0 | 0 | 0 | 0 | 1 | 1 |
| CPEB4 | 0 | 0 | 0 | 0 | 1 | 2 | 1 | 2 | 3 |
| CPSF1 | 0 | 0 | 0 | 0 | 0 | 0 | 0 | 3 | 1 |
| CPSF2 | 0 | 0 | 0 | 0 | 0 | 1 | 1 | 1 | 0 |
| CPSF4 | 0 | 0 | 0 | 0 | 0 | 0 | 0 | 3 | 1 |
| CPSF6 | 0 | 0 | 0 | 0 | 6 | 4 | 5 | 7 | 12 |

| | | | | | | | | | |
|---|---|---|---|---|---|---|---|---|---|
| CPSF7 | 0 | 0 | 0 | 0 | 3 | 2 | 4 | 6 | 5 |
| CROCC | 0 | 0 | 0 | 0 | 0 | 0 | 1 | 0 | 0 |
| CSNK1A1 | 0 | 0 | 0 | 0 | 1 | 0 | 0 | 2 | 1 |
| CSNK2A2 | 0 | 0 | 0 | 0 | 1 | 0 | 2 | 0 | 1 |
| CSNK2B | 0 | 0 | 0 | 0 | 2 | 3 | 4 | 2 | 2 |
| CSTF2 | 0 | 0 | 0 | 0 | 0 | 0 | 1 | 3 | 1 |
| CSTF2T | 0 | 0 | 0 | 0 | 0 | 1 | 0 | 3 | 3 |
| CSTF3 | 0 | 0 | 0 | 0 | 0 | 0 | 0 | 0 | 1 |
| CTCF | 0 | 0 | 0 | 0 | 0 | 0 | 0 | 1 | 0 |
| CTIF | 0 | 0 | 0 | 0 | 0 | 1 | 0 | 0 | 0 |
| CTNNA1 | 0 | 0 | 0 | 0 | 0 | 0 | 0 | 1 | 2 |
| CUL1 | 0 | 0 | 0 | 0 | 1 | 0 | 0 | 1 | 0 |
| CUL4A | 0 | 0 | 0 | 0 | 0 | 0 | 0 | 1 | 0 |
| CWC15 | 0 | 0 | 0 | 0 | 0 | 1 | 1 | 5 | 5 |
| CWC22 | 0 | 0 | 0 | 0 | 1 | 0 | 0 | 1 | 1 |
| CWF19L1 | 0 | 0 | 0 | 0 | 0 | 0 | 0 | 1 | 1 |
| CXorf56 | 0 | 0 | 0 | 0 | 0 | 0 | 0 | 4 | 5 |
| DAP | 0 | 0 | 0 | 0 | 0 | 0 | 0 | 0 | 1 |
| DCP1A | 0 | 0 | 0 | 0 | 1 | 2 | 1 | 4 | 2 |
| DDB1 | 0 | 0 | 0 | 0 | 0 | 0 | 2 | 2 | 1 |
| DDOST | 0 | 0 | 0 | 0 | 0 | 3 | 0 | 0 | 0 |
| DDRGK1 | 0 | 0 | 0 | 0 | 0 | 0 | 1 | 1 | 0 |
| DDT;DDTL | 0 | 0 | 0 | 0 | 1 | 0 | 1 | 0 | 0 |
| DDX10 | 0 | 0 | 0 | 0 | 5 | 7 | 5 | 11 | 8 |
| DDX17 | 0 | 0 | 0 | 0 | 0 | 0 | 0 | 0 | 1 |
| DDX19A;DDX19B | 0 | 0 | 0 | 0 | 0 | 0 | 0 | 2 | 3 |
| DDX39B | 0 | 0 | 0 | 0 | 0 | 0 | 0 | 0 | 1 |
| DDX41 | 0 | 0 | 0 | 0 | 0 | 0 | 0 | 1 | 1 |
| DDX42 | 0 | 0 | 0 | 0 | 11 | 14 | 13 | 19 | 18 |
| DDX46 | 0 | 0 | 0 | 0 | 0 | 0 | 0 | 1 | 2 |
| DDX54 | 0 | 0 | 0 | 0 | 0 | 0 | 0 | 1 | 0 |
| DDX58 | 0 | 0 | 0 | 0 | 0 | 0 | 1 | 0 | 0 |
| DDX6 | 0 | 0 | 0 | 0 | 1 | 1 | 2 | 9 | 6 |
| DGKA | 0 | 0 | 0 | 0 | 0 | 0 | 0 | 0 | 1 |
| DHCR7 | 0 | 0 | 0 | 0 | 0 | 2 | 0 | 2 | 2 |
| DHX15 | 0 | 0 | 0 | 0 | 5 | 8 | 6 | 9 | 11 |
| DHX35 | 0 | 0 | 0 | 0 | 0 | 0 | 1 | 2 | 1 |
| DHX36 | 0 | 0 | 0 | 0 | 0 | 0 | 0 | 1 | 1 |
| DHX38 | 0 | 0 | 0 | 0 | 2 | 2 | 0 | 3 | 5 |
| DHX9 | 0 | 0 | 0 | 0 | 2 | 8 | 6 | 15 | 16 |
| DICER1 | 0 | 0 | 0 | 0 | 0 | 0 | 2 | 2 | 3 |
| DIDO1 | 0 | 0 | 0 | 0 | 2 | 3 | 2 | 5 | 5 |
| DIP2A | 0 | 0 | 0 | 0 | 0 | 0 | 1 | 1 | 3 |
| DIP2B | 0 | 0 | 0 | 0 | 1 | 0 | 0 | 0 | 1 |
| DIS3 | 0 | 0 | 0 | 0 | 0 | 0 | 1 | 1 | 2 |

| | | | | | | | | | |
|---|---|---|---|---|---|---|---|---|---|
| DNAAF2 | 0 | 0 | 0 | 0 | 0 | 0 | 0 | 2 | 2 |
| DNAJA1 | 0 | 0 | 0 | 0 | 0 | 1 | 0 | 1 | 1 |
| DNAJA2 | 0 | 0 | 0 | 0 | 0 | 1 | 0 | 2 | 1 |
| DNAJC8 | 0 | 0 | 0 | 0 | 3 | 3 | 1 | 2 | 2 |
| DNAJC9 | 0 | 0 | 0 | 0 | 1 | 0 | 0 | 1 | 1 |
| DNTTIP2 | 0 | 0 | 0 | 0 | 0 | 0 | 0 | 3 | 5 |
| DOCK10 | 0 | 0 | 0 | 0 | 0 | 0 | 0 | 0 | 1 |
| DOCK5 | 0 | 0 | 0 | 0 | 0 | 1 | 0 | 1 | 0 |
| DPY30 | 0 | 0 | 0 | 0 | 0 | 0 | 2 | 0 | 0 |
| DPYSL4 | 0 | 0 | 0 | 0 | 0 | 0 | 0 | 0 | 2 |
| DSP | 0 | 0 | 0 | 0 | 2 | 1 | 0 | 2 | 2 |
| DSTN | 0 | 0 | 0 | 0 | 0 | 1 | 2 | 1 | 0 |
| DTL | 0 | 0 | 0 | 0 | 0 | 0 | 0 | 1 | 0 |
| DYNLT1 | 0 | 0 | 0 | 0 | 0 | 0 | 0 | 1 | 0 |
| EDF1 | 0 | 0 | 0 | 0 | 0 | 0 | 0 | 0 | 1 |
| EEF1B2 | 0 | 0 | 0 | 0 | 0 | 1 | 0 | 6 | 6 |
| EFR3A | 0 | 0 | 0 | 0 | 0 | 1 | 0 | 1 | 0 |
| EHBP1L1 | 0 | 0 | 0 | 0 | 0 | 0 | 0 | 3 | 1 |
| EHD2 | 0 | 0 | 0 | 0 | 0 | 0 | 0 | 1 | 1 |
| EHD4 | 0 | 0 | 0 | 0 | 0 | 0 | 0 | 1 | 0 |
| EHMT1 | 0 | 0 | 0 | 0 | 0 | 0 | 0 | 2 | 0 |
| EIF2AK2 | 0 | 0 | 0 | 0 | 0 | 1 | 0 | 0 | 0 |
| EIF2S1 | 0 | 0 | 0 | 0 | 0 | 1 | 0 | 2 | 2 |
| EIF2S2 | 0 | 0 | 0 | 0 | 0 | 0 | 0 | 1 | 1 |
| EIF3H | 0 | 0 | 0 | 0 | 0 | 0 | 0 | 3 | 3 |
| EIF4A3 | 0 | 0 | 0 | 0 | 1 | 0 | 0 | 0 | 2 |
| EIF4ENIF1 | 0 | 0 | 0 | 0 | 0 | 0 | 1 | 4 | 5 |
| ELAVL1 | 0 | 0 | 0 | 0 | 0 | 0 | 0 | 1 | 1 |
| ELOVL1 | 0 | 0 | 0 | 0 | 0 | 0 | 0 | 1 | 1 |
| ELOVL5 | 0 | 0 | 0 | 0 | 0 | 0 | 0 | 1 | 1 |
| ELP3 | 0 | 0 | 0 | 0 | 1 | 0 | 0 | 0 | 0 |
| EMC1 | 0 | 0 | 0 | 0 | 1 | 1 | 1 | 2 | 0 |
| EMC2 | 0 | 0 | 0 | 0 | 0 | 0 | 0 | 1 | 1 |
| EMD | 0 | 0 | 0 | 0 | 1 | 5 | 1 | 5 | 7 |
| EML4 | 0 | 0 | 0 | 0 | 0 | 0 | 0 | 2 | 0 |
| EMSY;C11orf30 | 0 | 0 | 0 | 0 | 0 | 0 | 0 | 7 | 7 |
| EP400 | 0 | 0 | 0 | 0 | 0 | 0 | 0 | 2 | 2 |
| EPB41 | 0 | 0 | 0 | 0 | 1 | 3 | 2 | 7 | 7 |
| EPB41L1 | 0 | 0 | 0 | 0 | 2 | 1 | 4 | 3 | 9 |
| EPB41L3 | 0 | 0 | 0 | 0 | 0 | 0 | 0 | 1 | 0 |
| EPS8 | 0 | 0 | 0 | 0 | 0 | 0 | 0 | 1 | 1 |
| ERCC6-PGBD3;PGB... | 0 | 0 | 0 | 0 | 0 | 0 | 1 | 0 | 0 |
| ERH | 0 | 0 | 0 | 0 | 1 | 2 | 2 | 1 | 1 |
| ERLIN2 | 0 | 0 | 0 | 0 | 0 | 0 | 1 | 0 | 0 |
| ESF1 | 0 | 0 | 0 | 0 | 0 | 1 | 0 | 7 | 2 |

| | | | | | | | | | |
|---|---|---|---|---|---|---|---|---|---|
| EXOC3 | 0 | 0 | 0 | 0 | 2 | 1 | 2 | 4 | 2 |
| EXOSC1 | 0 | 0 | 0 | 0 | 1 | 0 | 0 | 1 | 0 |
| EXOSC10 | 0 | 0 | 0 | 0 | 0 | 2 | 1 | 11 | 5 |
| EXOSC2 | 0 | 0 | 0 | 0 | 0 | 0 | 0 | 3 | 4 |
| EXOSC3 | 0 | 0 | 0 | 0 | 0 | 0 | 0 | 1 | 1 |
| EXOSC4 | 0 | 0 | 0 | 0 | 1 | 0 | 0 | 2 | 2 |
| EXOSC5 | 0 | 0 | 0 | 0 | 0 | 1 | 0 | 2 | 2 |
| EXOSC6 | 0 | 0 | 0 | 0 | 0 | 0 | 0 | 2 | 1 |
| EXOSC7 | 0 | 0 | 0 | 0 | 0 | 0 | 0 | 2 | 5 |
| EXOSC8 | 0 | 0 | 0 | 0 | 0 | 0 | 0 | 2 | 1 |
| EXOSC9 | 0 | 0 | 0 | 0 | 1 | 1 | 1 | 3 | 3 |
| FAM107B | 0 | 0 | 0 | 0 | 0 | 0 | 0 | 1 | 1 |
| FAM120A | 0 | 0 | 0 | 0 | 0 | 0 | 2 | 8 | 8 |
| FAM171A1 | 0 | 0 | 0 | 0 | 0 | 1 | 0 | 1 | 1 |
| FAM192A;NIP30 | 0 | 0 | 0 | 0 | 0 | 0 | 0 | 5 | 3 |
| FAM208A | 0 | 0 | 0 | 0 | 3 | 7 | 2 | 16 | 15 |
| FAM47E-STBD1 | 0 | 0 | 0 | 0 | 0 | 0 | 0 | 6 | 4 |
| FAM50A | 0 | 0 | 0 | 0 | 0 | 0 | 0 | 1 | 0 |
| FAM98A | 0 | 0 | 0 | 0 | 0 | 0 | 0 | 3 | 1 |
| FBXO30 | 0 | 0 | 0 | 0 | 0 | 0 | 0 | 0 | 1 |
| FEN1 | 0 | 0 | 0 | 0 | 3 | 3 | 1 | 6 | 8 |
| FERMT2 | 0 | 0 | 0 | 0 | 2 | 3 | 2 | 10 | 5 |
| FHL3 | 0 | 0 | 0 | 0 | 0 | 0 | 0 | 3 | 1 |
| FIP1L1 | 0 | 0 | 0 | 0 | 1 | 3 | 2 | 8 | 8 |
| FKBP3 | 0 | 0 | 0 | 0 | 0 | 2 | 2 | 1 | 1 |
| FMNL3 | 0 | 0 | 0 | 0 | 0 | 0 | 0 | 0 | 1 |
| FMR1 | 0 | 0 | 0 | 0 | 0 | 0 | 0 | 1 | 1 |
| FNBP1 | 0 | 0 | 0 | 0 | 0 | 0 | 0 | 1 | 0 |
| FNBP1L | 0 | 0 | 0 | 0 | 1 | 0 | 0 | 0 | 0 |
| FNBP4 | 0 | 0 | 0 | 0 | 0 | 0 | 0 | 0 | 1 |
| FNDC3A | 0 | 0 | 0 | 0 | 2 | 0 | 0 | 3 | 3 |
| FNDC3B | 0 | 0 | 0 | 0 | 0 | 0 | 0 | 0 | 2 |
| FNIP1 | 0 | 0 | 0 | 0 | 0 | 0 | 0 | 1 | 0 |
| FOCAD | 0 | 0 | 0 | 0 | 0 | 1 | 0 | 0 | 1 |
| FOSL1 | 0 | 0 | 0 | 0 | 0 | 0 | 0 | 1 | 0 |
| FOXK1;FOXK2 | 0 | 0 | 0 | 0 | 0 | 0 | 0 | 1 | 0 |
| FRMD6 | 0 | 0 | 0 | 0 | 0 | 0 | 0 | 1 | 0 |
| FSCN1 | 0 | 0 | 0 | 0 | 0 | 0 | 0 | 1 | 0 |
| FTSJ3 | 0 | 0 | 0 | 0 | 0 | 1 | 0 | 5 | 1 |
| FUBP1 | 0 | 0 | 0 | 0 | 1 | 5 | 1 | 4 | 6 |
| FUBP3 | 0 | 0 | 0 | 0 | 0 | 0 | 0 | 1 | 2 |
| FUS | 0 | 0 | 0 | 0 | 1 | 0 | 0 | 3 | 3 |
| FXR1 | 0 | 0 | 0 | 0 | 0 | 0 | 1 | 6 | 3 |
| FXR2 | 0 | 0 | 0 | 0 | 0 | 0 | 0 | 1 | 3 |
| G3BP2 | 0 | 0 | 0 | 0 | 0 | 2 | 1 | 4 | 2 |

| | | | | | | | | | |
|---|---|---|---|---|---|---|---|---|---|
| GABPA | 0 | 0 | 0 | 0 | 0 | 0 | 0 | 1 | 1 |
| GAMT | 0 | 0 | 0 | 0 | 0 | 0 | 0 | 1 | 1 |
| GARS | 0 | 0 | 0 | 0 | 0 | 0 | 0 | 1 | 0 |
| GAS2L3 | 0 | 0 | 0 | 0 | 0 | 0 | 0 | 0 | 1 |
| GATAD2A | 0 | 0 | 0 | 0 | 0 | 0 | 0 | 3 | 1 |
| GATAD2B | 0 | 0 | 0 | 0 | 0 | 1 | 0 | 8 | 3 |
| GGA3 | 0 | 0 | 0 | 0 | 0 | 0 | 0 | 1 | 1 |
| GIT1 | 0 | 0 | 0 | 0 | 0 | 0 | 0 | 0 | 1 |
| GIT2 | 0 | 0 | 0 | 0 | 0 | 2 | 0 | 2 | 0 |
| GLYR1 | 0 | 0 | 0 | 0 | 1 | 2 | 2 | 1 | 1 |
| GMPS | 0 | 0 | 0 | 0 | 3 | 3 | 4 | 2 | 2 |
| GNAI3 | 0 | 0 | 0 | 0 | 0 | 0 | 1 | 0 | 4 |
| GNAS | 0 | 0 | 0 | 0 | 0 | 1 | 0 | 0 | 4 |
| GNB4 | 0 | 0 | 0 | 0 | 0 | 0 | 0 | 0 | 2 |
| GNG12 | 0 | 0 | 0 | 0 | 0 | 0 | 1 | 0 | 3 |
| GNL2 | 0 | 0 | 0 | 0 | 0 | 0 | 0 | 5 | 4 |
| GNL3 | 0 | 0 | 0 | 0 | 0 | 0 | 0 | 1 | 0 |
| GOLGA2 | 0 | 0 | 0 | 0 | 0 | 0 | 0 | 4 | 2 |
| GOLGA4 | 0 | 0 | 0 | 0 | 0 | 0 | 0 | 0 | 1 |
| GOPC | 0 | 0 | 0 | 0 | 0 | 0 | 0 | 0 | 1 |
| GPATCH1 | 0 | 0 | 0 | 0 | 0 | 0 | 1 | 2 | 1 |
| GPATCH8 | 0 | 0 | 0 | 0 | 0 | 0 | 0 | 2 | 0 |
| GPKOW | 0 | 0 | 0 | 0 | 0 | 0 | 0 | 2 | 2 |
| GPRC5A | 0 | 0 | 0 | 0 | 0 | 0 | 0 | 0 | 4 |
| GPS1 | 0 | 0 | 0 | 0 | 0 | 0 | 0 | 5 | 3 |
| GRWD1 | 0 | 0 | 0 | 0 | 0 | 0 | 0 | 1 | 0 |
| GSTCD | 0 | 0 | 0 | 0 | 0 | 0 | 0 | 1 | 0 |
| GTF2A2 | 0 | 0 | 0 | 0 | 0 | 1 | 1 | 2 | 2 |
| GTF2E1 | 0 | 0 | 0 | 0 | 0 | 1 | 2 | 0 | 2 |
| GTF2E2 | 0 | 0 | 0 | 0 | 4 | 3 | 4 | 2 | 1 |
| GTF2F1 | 0 | 0 | 0 | 0 | 2 | 2 | 2 | 11 | 5 |
| GTF2F2 | 0 | 0 | 0 | 0 | 2 | 2 | 4 | 4 | 5 |
| GTF2I | 0 | 0 | 0 | 0 | 1 | 2 | 1 | 10 | 5 |
| GTF3C1 | 0 | 0 | 0 | 0 | 0 | 1 | 0 | 2 | 0 |
| GTF3C3 | 0 | 0 | 0 | 0 | 0 | 0 | 0 | 0 | 1 |
| GTF3C4 | 0 | 0 | 0 | 0 | 0 | 1 | 1 | 0 | 0 |
| GTF3C5 | 0 | 0 | 0 | 0 | 0 | 0 | 0 | 1 | 3 |
| GYG1 | 0 | 0 | 0 | 0 | 0 | 0 | 0 | 1 | 0 |
| HARS | 0 | 0 | 0 | 0 | 0 | 1 | 2 | 0 | 0 |
| HAT1 | 0 | 0 | 0 | 0 | 0 | 0 | 0 | 2 | 4 |
| HAUS1 | 0 | 0 | 0 | 0 | 0 | 0 | 0 | 1 | 0 |
| HDAC1 | 0 | 0 | 0 | 0 | 0 | 0 | 0 | 3 | 3 |
| HDAC2 | 0 | 0 | 0 | 0 | 2 | 1 | 0 | 7 | 7 |
| HDGF | 0 | 0 | 0 | 0 | 1 | 2 | 0 | 3 | 3 |
| HDGFRP2 | 0 | 0 | 0 | 0 | 3 | 5 | 5 | 6 | 8 |

| | | | | | | | | | |
|---|---|---|---|---|---|---|---|---|---|
| HEATR5A | 0 | 0 | 0 | 0 | 1 | 1 | 1 | 0 | 0 |
| HEXIM1 | 0 | 0 | 0 | 0 | 0 | 1 | 0 | 0 | 1 |
| HIGD1A | 0 | 0 | 0 | 0 | 0 | 0 | 0 | 1 | 1 |
| HIVEP1 | 0 | 0 | 0 | 0 | 0 | 0 | 0 | 1 | 2 |
| HK1 | 0 | 0 | 0 | 0 | 0 | 0 | 0 | 2 | 2 |
| HLA-C;HLA-B;HLA-A;HLA-H | 0 | 0 | 0 | 0 | 0 | 0 | 0 | 1 | 1 |
| HMGB1;HMGB1P1 | 0 | 0 | 0 | 0 | 0 | 1 | 0 | 1 | 1 |
| HMGXB4 | 0 | 0 | 0 | 0 | 0 | 0 | 0 | 1 | 1 |
| HNRNPA0 | 0 | 0 | 0 | 0 | 0 | 0 | 0 | 1 | 2 |
| HNRNPA3 | 0 | 0 | 0 | 0 | 0 | 0 | 0 | 1 | 0 |
| HNRNPF | 0 | 0 | 0 | 0 | 1 | 0 | 4 | 3 | 8 |
| HNRNPH3 | 0 | 0 | 0 | 0 | 0 | 0 | 0 | 1 | 0 |
| HP1BP3 | 0 | 0 | 0 | 0 | 9 | 9 | 11 | 15 | 16 |
| HSD17B12 | 0 | 0 | 0 | 0 | 0 | 1 | 2 | 2 | 2 |
| HSPA1L | 0 | 0 | 0 | 0 | 2 | 0 | 0 | 0 | 0 |
| HSPD1 | 0 | 0 | 0 | 0 | 0 | 2 | 0 | 3 | 2 |
| HSPE1-MOB4;MOB4 | 0 | 0 | 0 | 0 | 1 | 0 | 0 | 1 | 1 |
| HTATSF1 | 0 | 0 | 0 | 0 | 1 | 2 | 2 | 1 | 1 |
| HYOU1 | 0 | 0 | 0 | 0 | 0 | 1 | 0 | 0 | 0 |
| IFI16 | 0 | 0 | 0 | 0 | 4 | 6 | 4 | 10 | 11 |
| IFI35 | 0 | 0 | 0 | 0 | 0 | 0 | 1 | 0 | 0 |
| IFIT3 | 0 | 0 | 0 | 0 | 0 | 0 | 0 | 0 | 1 |
| IGF2BP2 | 0 | 0 | 0 | 0 | 0 | 0 | 0 | 4 | 3 |
| IGF2BP3 | 0 | 0 | 0 | 0 | 1 | 2 | 1 | 3 | 1 |
| IK | 0 | 0 | 0 | 0 | 0 | 1 | 0 | 6 | 4 |
| IKBKAP | 0 | 0 | 0 | 0 | 0 | 1 | 0 | 1 | 3 |
| ILKAP;ILKAP3 | 0 | 0 | 0 | 0 | 0 | 0 | 0 | 2 | 2 |
| IMMT | 0 | 0 | 0 | 0 | 1 | 0 | 0 | 0 | 0 |
| IMPDH2 | 0 | 0 | 0 | 0 | 0 | 0 | 0 | 4 | 1 |
| INF2 | 0 | 0 | 0 | 0 | 0 | 1 | 2 | 12 | 3 |
| INTS12 | 0 | 0 | 0 | 0 | 0 | 0 | 0 | 2 | 4 |
| INTS3 | 0 | 0 | 0 | 0 | 1 | 3 | 3 | 1 | 1 |
| INTS9 | 0 | 0 | 0 | 0 | 0 | 0 | 1 | 0 | 0 |
| IPO9 | 0 | 0 | 0 | 0 | 0 | 0 | 0 | 2 | 2 |
| IQGAP1 | 0 | 0 | 0 | 0 | 29 | 46 | 57 | 6 | 28 |
| IRF2BP2 | 0 | 0 | 0 | 0 | 0 | 0 | 0 | 0 | 1 |
| IRF2BPL | 0 | 0 | 0 | 0 | 0 | 0 | 0 | 2 | 5 |
| IST1 | 0 | 0 | 0 | 0 | 1 | 0 | 0 | 3 | 1 |
| ISY1 | 0 | 0 | 0 | 0 | 0 | 2 | 0 | 0 | 0 |
| ITGA2 | 0 | 0 | 0 | 0 | 1 | 1 | 1 | 2 | 0 |
| ITGA3 | 0 | 0 | 0 | 0 | 1 | 4 | 3 | 9 | 4 |
| ITGAV | 0 | 0 | 0 | 0 | 0 | 0 | 2 | 1 | 0 |
| ITGB5 | 0 | 0 | 0 | 0 | 0 | 2 | 1 | 1 | 2 |
| ITSN1 | 0 | 0 | 0 | 0 | 0 | 0 | 0 | 0 | 1 |
| ITSN2 | 0 | 0 | 0 | 0 | 0 | 0 | 1 | 3 | 1 |

| | | | | | | | | | |
|---|---|---|---|---|---|---|---|---|---|
| JMJD1C | 0 | 0 | 0 | 0 | 7 | 10 | 10 | 28 | 18 |
| JUN | 0 | 0 | 0 | 0 | 0 | 0 | 0 | 1 | 0 |
| JUNB | 0 | 0 | 0 | 0 | 1 | 0 | 0 | 1 | 2 |
| KANK1 | 0 | 0 | 0 | 0 | 0 | 0 | 0 | 2 | 2 |
| KATNAL1;KATNA1 | 0 | 0 | 0 | 0 | 0 | 0 | 0 | 1 | 0 |
| KDELR1 | 0 | 0 | 0 | 0 | 0 | 0 | 0 | 0 | 1 |
| KDM3B | 0 | 0 | 0 | 0 | 3 | 6 | 3 | 3 | 7 |
| KHDRBS1 | 0 | 0 | 0 | 0 | 0 | 0 | 0 | 1 | 1 |
| KHSRP | 0 | 0 | 0 | 0 | 2 | 3 | 5 | 10 | 12 |
| KIAA1143 | 0 | 0 | 0 | 0 | 0 | 2 | 0 | 3 | 1 |
| KIAA1244 | 0 | 0 | 0 | 0 | 0 | 0 | 0 | 3 | 0 |
| KIAA1468 | 0 | 0 | 0 | 0 | 1 | 1 | 0 | 0 | 0 |
| KIAA1598 | 0 | 0 | 0 | 0 | 3 | 1 | 5 | 3 | 7 |
| KIF14 | 0 | 0 | 0 | 0 | 0 | 2 | 0 | 2 | 0 |
| KIF22 | 0 | 0 | 0 | 0 | 0 | 0 | 0 | 1 | 0 |
| KIF23 | 0 | 0 | 0 | 0 | 1 | 1 | 0 | 3 | 2 |
| KIF2A | 0 | 0 | 0 | 0 | 0 | 0 | 0 | 3 | 2 |
| KIF2C | 0 | 0 | 0 | 0 | 0 | 0 | 0 | 4 | 1 |
| KIF4A | 0 | 0 | 0 | 0 | 1 | 1 | 1 | 12 | 8 |
| KIFC3 | 0 | 0 | 0 | 0 | 0 | 0 | 0 | 0 | 1 |
| KMT2A | 0 | 0 | 0 | 0 | 2 | 1 | 0 | 3 | 3 |
| KPNA1 | 0 | 0 | 0 | 0 | 0 | 0 | 0 | 1 | 1 |
| KPNA2 | 0 | 0 | 0 | 0 | 1 | 6 | 1 | 3 | 1 |
| KPNA3 | 0 | 0 | 0 | 0 | 2 | 4 | 6 | 1 | 3 |
| KPNA4 | 0 | 0 | 0 | 0 | 5 | 4 | 4 | 3 | 5 |
| KPNA6;KPNA5 | 0 | 0 | 0 | 0 | 1 | 0 | 0 | 2 | 1 |
| KTN1 | 0 | 0 | 0 | 0 | 1 | 1 | 0 | 3 | 0 |
| LARP4 | 0 | 0 | 0 | 0 | 0 | 1 | 0 | 3 | 3 |
| LARP4B | 0 | 0 | 0 | 0 | 0 | 0 | 0 | 0 | 1 |
| LARP6 | 0 | 0 | 0 | 0 | 0 | 0 | 0 | 1 | 1 |
| LENG8 | 0 | 0 | 0 | 0 | 0 | 0 | 0 | 1 | 1 |
| LEO1 | 0 | 0 | 0 | 0 | 1 | 0 | 0 | 1 | 0 |
| LEPROT | 0 | 0 | 0 | 0 | 0 | 0 | 0 | 1 | 1 |
| LGALS8 | 0 | 0 | 0 | 0 | 0 | 0 | 0 | 0 | 1 |
| LIG1 | 0 | 0 | 0 | 0 | 0 | 1 | 0 | 0 | 0 |
| LIG3 | 0 | 0 | 0 | 0 | 0 | 0 | 0 | 0 | 2 |
| LIN37 | 0 | 0 | 0 | 0 | 0 | 1 | 1 | 1 | 0 |
| LIN54 | 0 | 0 | 0 | 0 | 0 | 0 | 0 | 1 | 1 |
| LIN7C;LIN7A | 0 | 0 | 0 | 0 | 1 | 0 | 0 | 0 | 0 |
| LLGL1 | 0 | 0 | 0 | 0 | 0 | 1 | 0 | 1 | 0 |
| LMNB1 | 0 | 0 | 0 | 0 | 1 | 0 | 0 | 1 | 1 |
| LRCH1 | 0 | 0 | 0 | 0 | 0 | 0 | 1 | 2 | 0 |
| LRRC59 | 0 | 0 | 0 | 0 | 0 | 1 | 1 | 0 | 1 |
| LRRC7 | 0 | 0 | 0 | 0 | 0 | 0 | 0 | 0 | 1 |
| LRWD1 | 0 | 0 | 0 | 0 | 6 | 4 | 4 | 11 | 8 |

| | | | | | | | | | |
|---|---|---|---|---|---|---|---|---|---|
| LSG1 | 0 | 0 | 0 | 0 | 2 | 1 | 1 | 5 | 1 |
| LSM12 | 0 | 0 | 0 | 0 | 0 | 0 | 0 | 3 | 3 |
| LSM14B | 0 | 0 | 0 | 0 | 0 | 0 | 0 | 0 | 1 |
| LSM2 | 0 | 0 | 0 | 0 | 1 | 2 | 0 | 1 | 1 |
| LSM4 | 0 | 0 | 0 | 0 | 0 | 0 | 0 | 2 | 2 |
| LXN | 0 | 0 | 0 | 0 | 0 | 0 | 0 | 1 | 1 |
| LYPLAL1 | 0 | 0 | 0 | 0 | 0 | 0 | 2 | 0 | 0 |
| MAP2 | 0 | 0 | 0 | 0 | 0 | 1 | 0 | 2 | 2 |
| MAP3K7 | 0 | 0 | 0 | 0 | 1 | 0 | 0 | 4 | 4 |
| MAP4K4 | 0 | 0 | 0 | 0 | 0 | 0 | 0 | 1 | 1 |
| MAP7D3 | 0 | 0 | 0 | 0 | 1 | 1 | 0 | 21 | 10 |
| MAPRE1 | 0 | 0 | 0 | 0 | 0 | 0 | 1 | 5 | 4 |
| MARK3 | 0 | 0 | 0 | 0 | 0 | 0 | 0 | 2 | 1 |
| MASTL | 0 | 0 | 0 | 0 | 1 | 2 | 3 | 3 | 3 |
| MAT2A | 0 | 0 | 0 | 0 | 0 | 0 | 0 | 0 | 1 |
| MATR3 | 0 | 0 | 0 | 0 | 5 | 9 | 7 | 9 | 8 |
| MBNL1 | 0 | 0 | 0 | 0 | 0 | 0 | 0 | 1 | 1 |
| MBOAT7 | 0 | 0 | 0 | 0 | 1 | 3 | 3 | 1 | 1 |
| MCM3 | 0 | 0 | 0 | 0 | 0 | 0 | 1 | 2 | 2 |
| MCM3AP | 0 | 0 | 0 | 0 | 0 | 0 | 0 | 1 | 0 |
| MCM4 | 0 | 0 | 0 | 0 | 0 | 1 | 1 | 5 | 7 |
| MCM5 | 0 | 0 | 0 | 0 | 0 | 0 | 0 | 2 | 1 |
| MCM6 | 0 | 0 | 0 | 0 | 1 | 4 | 0 | 4 | 4 |
| MCU | 0 | 0 | 0 | 0 | 0 | 0 | 0 | 0 | 1 |
| MDC1 | 0 | 0 | 0 | 0 | 0 | 1 | 0 | 1 | 2 |
| ME1 | 0 | 0 | 0 | 0 | 0 | 0 | 0 | 1 | 1 |
| MED1 | 0 | 0 | 0 | 0 | 1 | 0 | 0 | 8 | 7 |
| MEF2D | 0 | 0 | 0 | 0 | 1 | 2 | 2 | 4 | 4 |
| MEPCE | 0 | 0 | 0 | 0 | 0 | 0 | 0 | 2 | 0 |
| MEST | 0 | 0 | 0 | 0 | 0 | 0 | 0 | 0 | 1 |
| METTL14 | 0 | 0 | 0 | 0 | 0 | 0 | 0 | 1 | 0 |
| METTL3 | 0 | 0 | 0 | 0 | 0 | 0 | 0 | 2 | 0 |
| MFF | 0 | 0 | 0 | 0 | 3 | 0 | 2 | 1 | 3 |
| MICAL2 | 0 | 0 | 0 | 0 | 0 | 0 | 0 | 1 | 0 |
| MICAL3 | 0 | 0 | 0 | 0 | 0 | 0 | 0 | 1 | 0 |
| MINK1 | 0 | 0 | 0 | 0 | 0 | 0 | 0 | 1 | 0 |
| MIOS | 0 | 0 | 0 | 0 | 0 | 0 | 0 | 1 | 0 |
| MLH1 | 0 | 0 | 0 | 0 | 1 | 0 | 2 | 13 | 8 |
| MLLT1 | 0 | 0 | 0 | 0 | 1 | 0 | 0 | 0 | 1 |
| MLLT6 | 0 | 0 | 0 | 0 | 0 | 0 | 0 | 2 | 0 |
| MOB3A | 0 | 0 | 0 | 0 | 0 | 0 | 0 | 0 | 1 |
| MOCS2 | 0 | 0 | 0 | 0 | 0 | 0 | 0 | 0 | 2 |
| MPDZ | 0 | 0 | 0 | 0 | 0 | 0 | 0 | 1 | 2 |
| MPG | 0 | 0 | 0 | 0 | 1 | 0 | 1 | 0 | 0 |
| MPP5 | 0 | 0 | 0 | 0 | 1 | 0 | 0 | 0 | 0 |

| | | | | | | | | | |
|---|---|---|---|---|---|---|---|---|---|
| MPZL1 | 0 | 0 | 0 | 0 | 0 | 0 | 0 | 0 | 1 |
| MSANTD2 | 0 | 0 | 0 | 0 | 4 | 2 | 1 | 10 | 9 |
| MSH2 | 0 | 0 | 0 | 0 | 2 | 2 | 2 | 4 | 2 |
| MSH3 | 0 | 0 | 0 | 0 | 0 | 1 | 0 | 0 | 1 |
| MSRB3 | 0 | 0 | 0 | 0 | 0 | 0 | 1 | 1 | 1 |
| MT-ATP6 | 0 | 0 | 0 | 0 | 0 | 0 | 0 | 0 | 1 |
| MTA1 | 0 | 0 | 0 | 0 | 1 | 1 | 1 | 2 | 3 |
| MTA2 | 0 | 0 | 0 | 0 | 2 | 1 | 1 | 4 | 2 |
| MTDH | 0 | 0 | 0 | 0 | 1 | 2 | 0 | 9 | 10 |
| MTHFD1 | 0 | 0 | 0 | 0 | 0 | 0 | 0 | 0 | 1 |
| MVP | 0 | 0 | 0 | 0 | 1 | 1 | 3 | 2 | 0 |
| MXRA7 | 0 | 0 | 0 | 0 | 0 | 0 | 0 | 2 | 0 |
| MYADM | 0 | 0 | 0 | 0 | 0 | 0 | 0 | 0 | 1 |
| MYBBP1A | 0 | 0 | 0 | 0 | 1 | 0 | 0 | 2 | 0 |
| MYL6B | 0 | 0 | 0 | 0 | 0 | 0 | 1 | 0 | 0 |
| MYO1B | 0 | 0 | 0 | 0 | 17 | 12 | 27 | 0 | 34 |
| MYO5A | 0 | 0 | 0 | 0 | 17 | 12 | 17 | 0 | 33 |
| MYPN | 0 | 0 | 0 | 0 | 0 | 0 | 0 | 1 | 1 |
| NAA15 | 0 | 0 | 0 | 0 | 0 | 1 | 1 | 1 | 1 |
| NAA30 | 0 | 0 | 0 | 0 | 0 | 0 | 0 | 0 | 1 |
| NAA35 | 0 | 0 | 0 | 0 | 0 | 0 | 0 | 1 | 1 |
| NACC1 | 0 | 0 | 0 | 0 | 0 | 2 | 1 | 9 | 8 |
| NANS | 0 | 0 | 0 | 0 | 0 | 2 | 0 | 0 | 0 |
| NASP | 0 | 0 | 0 | 0 | 3 | 8 | 3 | 11 | 15 |
| NAT10 | 0 | 0 | 0 | 0 | 1 | 0 | 0 | 5 | 6 |
| NAV3 | 0 | 0 | 0 | 0 | 0 | 0 | 0 | 3 | 1 |
| NBN | 0 | 0 | 0 | 0 | 0 | 2 | 0 | 6 | 0 |
| NCAPH | 0 | 0 | 0 | 0 | 0 | 1 | 0 | 11 | 11 |
| NCBP1 | 0 | 0 | 0 | 0 | 0 | 0 | 0 | 3 | 3 |
| NCEH1 | 0 | 0 | 0 | 0 | 0 | 0 | 0 | 1 | 0 |
| NCLN | 0 | 0 | 0 | 0 | 0 | 0 | 0 | 0 | 1 |
| NCOA6 | 0 | 0 | 0 | 0 | 0 | 0 | 0 | 1 | 1 |
| NCOA7 | 0 | 0 | 0 | 0 | 0 | 0 | 0 | 5 | 6 |
| NCOR1 | 0 | 0 | 0 | 0 | 0 | 0 | 0 | 2 | 0 |
| NCOR2 | 0 | 0 | 0 | 0 | 3 | 5 | 3 | 4 | 4 |
| NDC80 | 0 | 0 | 0 | 0 | 0 | 0 | 0 | 1 | 0 |
| NDUFA10 | 0 | 0 | 0 | 0 | 1 | 0 | 0 | 1 | 0 |
| NEDD4L | 0 | 0 | 0 | 0 | 0 | 2 | 0 | 2 | 2 |
| NEDD8 | 0 | 0 | 0 | 0 | 0 | 1 | 0 | 0 | 0 |
| NELFA | 0 | 0 | 0 | 0 | 1 | 0 | 0 | 2 | 0 |
| NELFB | 0 | 0 | 0 | 0 | 1 | 1 | 1 | 2 | 2 |
| NELFCD;TH1L | 0 | 0 | 0 | 0 | 1 | 1 | 0 | 1 | 3 |
| NELFE | 0 | 0 | 0 | 0 | 1 | 3 | 1 | 8 | 5 |
| NEXN | 0 | 0 | 0 | 0 | 2 | 6 | 17 | 11 | 16 |
| NF1 | 0 | 0 | 0 | 0 | 0 | 0 | 0 | 0 | 1 |

| | | | | | | | | | |
|---|---|---|---|---|---|---|---|---|---|
| NF2 | 0 | 0 | 0 | 0 | 0 | 1 | 1 | 6 | 5 |
| NFIA | 0 | 0 | 0 | 0 | 0 | 1 | 0 | 2 | 2 |
| NFRKB | 0 | 0 | 0 | 0 | 0 | 0 | 0 | 1 | 0 |
| NHLRC2 | 0 | 0 | 0 | 0 | 2 | 5 | 2 | 3 | 3 |
| NHP2L1 | 0 | 0 | 0 | 0 | 1 | 0 | 0 | 3 | 2 |
| NME1 | 0 | 0 | 0 | 0 | 0 | 0 | 0 | 1 | 1 |
| NMT1 | 0 | 0 | 0 | 0 | 0 | 1 | 2 | 5 | 2 |
| NMT2 | 0 | 0 | 0 | 0 | 0 | 0 | 0 | 0 | 1 |
| NOC2L | 0 | 0 | 0 | 0 | 0 | 0 | 0 | 0 | 1 |
| NOL11 | 0 | 0 | 0 | 0 | 0 | 0 | 0 | 1 | 0 |
| NOL6 | 0 | 0 | 0 | 0 | 0 | 1 | 1 | 0 | 0 |
| NOL8 | 0 | 0 | 0 | 0 | 0 | 0 | 0 | 2 | 0 |
| NOLC1 | 0 | 0 | 0 | 0 | 1 | 0 | 0 | 3 | 3 |
| NOP56 | 0 | 0 | 0 | 0 | 0 | 0 | 0 | 1 | 0 |
| NOTCH2 | 0 | 0 | 0 | 0 | 1 | 2 | 2 | 3 | 2 |
| NPM3 | 0 | 0 | 0 | 0 | 2 | 2 | 3 | 2 | 1 |
| NR2C1 | 0 | 0 | 0 | 0 | 0 | 0 | 0 | 1 | 2 |
| NR2C2 | 0 | 0 | 0 | 0 | 0 | 0 | 0 | 4 | 1 |
| NUCKS1 | 0 | 0 | 0 | 0 | 0 | 0 | 0 | 4 | 7 |
| NUDCD1 | 0 | 0 | 0 | 0 | 0 | 0 | 0 | 4 | 2 |
| NUDCD2 | 0 | 0 | 0 | 0 | 0 | 0 | 0 | 0 | 1 |
| NUDT2 | 0 | 0 | 0 | 0 | 1 | 0 | 0 | 0 | 1 |
| NUDT21 | 0 | 0 | 0 | 0 | 4 | 9 | 9 | 9 | 5 |
| NUFIP2 | 0 | 0 | 0 | 0 | 5 | 7 | 3 | 17 | 20 |
| NUMA1 | 0 | 0 | 0 | 0 | 3 | 11 | 7 | 42 | 34 |
| NUP107 | 0 | 0 | 0 | 0 | 0 | 0 | 0 | 2 | 0 |
| NUP153 | 0 | 0 | 0 | 0 | 7 | 9 | 6 | 36 | 27 |
| NUP205 | 0 | 0 | 0 | 0 | 0 | 0 | 0 | 0 | 2 |
| NUP50 | 0 | 0 | 0 | 0 | 1 | 2 | 1 | 9 | 11 |
| NUP93 | 0 | 0 | 0 | 0 | 1 | 1 | 1 | 2 | 1 |
| NUPL1 | 0 | 0 | 0 | 0 | 0 | 0 | 0 | 1 | 1 |
| OCIAD1 | 0 | 0 | 0 | 0 | 0 | 0 | 0 | 2 | 0 |
| OCRL | 0 | 0 | 0 | 0 | 0 | 0 | 0 | 3 | 1 |
| OGT | 0 | 0 | 0 | 0 | 1 | 2 | 0 | 4 | 1 |
| OPHN1 | 0 | 0 | 0 | 0 | 0 | 0 | 0 | 2 | 1 |
| ORC2 | 0 | 0 | 0 | 0 | 2 | 4 | 5 | 3 | 8 |
| ORC3 | 0 | 0 | 0 | 0 | 1 | 2 | 1 | 0 | 1 |
| OSBPL3 | 0 | 0 | 0 | 0 | 0 | 0 | 0 | 1 | 2 |
| OSBPL8 | 0 | 0 | 0 | 0 | 2 | 6 | 5 | 4 | 0 |
| OTUD4 | 0 | 0 | 0 | 0 | 1 | 0 | 2 | 1 | 1 |
| P4HA1 | 0 | 0 | 0 | 0 | 0 | 2 | 2 | 1 | 3 |
| P4HB | 0 | 0 | 0 | 0 | 0 | 0 | 0 | 2 | 0 |
| PA2G4 | 0 | 0 | 0 | 0 | 0 | 0 | 1 | 5 | 3 |
| PAF1 | 0 | 0 | 0 | 0 | 0 | 1 | 0 | 2 | 1 |
| PAIP1 | 0 | 0 | 0 | 0 | 0 | 1 | 0 | 1 | 1 |

| | | | | | | | | | |
|---|---|---|---|---|---|---|---|---|---|
| PAK4;PAK7 | 0 | 0 | 0 | 0 | 0 | 0 | 0 | 1 | 1 |
| PALM2 | 0 | 0 | 0 | 0 | 0 | 0 | 0 | 1 | 0 |
| PAPOLA | 0 | 0 | 0 | 0 | 12 | 11 | 10 | 20 | 23 |
| PAPSS1 | 0 | 0 | 0 | 0 | 0 | 0 | 0 | 3 | 1 |
| PAPSS2 | 0 | 0 | 0 | 0 | 1 | 1 | 0 | 0 | 2 |
| PARG | 0 | 0 | 0 | 0 | 1 | 0 | 0 | 0 | 1 |
| PARP12 | 0 | 0 | 0 | 0 | 0 | 0 | 0 | 0 | 1 |
| PARP4 | 0 | 0 | 0 | 0 | 2 | 3 | 6 | 1 | 1 |
| PAWR | 0 | 0 | 0 | 0 | 0 | 0 | 0 | 2 | 5 |
| PAXBP1 | 0 | 0 | 0 | 0 | 0 | 0 | 0 | 1 | 1 |
| PCBP3 | 0 | 0 | 0 | 0 | 0 | 0 | 0 | 0 | 1 |
| PCF11 | 0 | 0 | 0 | 0 | 0 | 0 | 0 | 1 | 0 |
| PCM1 | 0 | 0 | 0 | 0 | 0 | 0 | 0 | 9 | 2 |
| PCMT1 | 0 | 0 | 0 | 0 | 0 | 0 | 0 | 2 | 3 |
| PCYT1A | 0 | 0 | 0 | 0 | 1 | 2 | 2 | 5 | 6 |
| PCYT2 | 0 | 0 | 0 | 0 | 0 | 0 | 0 | 1 | 0 |
| PDCD5 | 0 | 0 | 0 | 0 | 0 | 1 | 0 | 3 | 0 |
| PDCD6IP | 0 | 0 | 0 | 0 | 0 | 0 | 0 | 1 | 0 |
| PDCL | 0 | 0 | 0 | 0 | 0 | 0 | 1 | 1 | 0 |
| PDIA3 | 0 | 0 | 0 | 0 | 0 | 0 | 0 | 0 | 2 |
| PDIA6 | 0 | 0 | 0 | 0 | 0 | 0 | 0 | 0 | 1 |
| PDRG1 | 0 | 0 | 0 | 0 | 0 | 0 | 0 | 0 | 1 |
| PDS5A | 0 | 0 | 0 | 0 | 1 | 0 | 0 | 0 | 0 |
| PDS5B | 0 | 0 | 0 | 0 | 1 | 1 | 0 | 3 | 3 |
| PDXDC1 | 0 | 0 | 0 | 0 | 0 | 0 | 1 | 2 | 0 |
| PDZD11 | 0 | 0 | 0 | 0 | 0 | 0 | 1 | 1 | 3 |
| PDZD8 | 0 | 0 | 0 | 0 | 1 | 1 | 3 | 2 | 2 |
| PELO | 0 | 0 | 0 | 0 | 0 | 0 | 1 | 1 | 0 |
| PES1 | 0 | 0 | 0 | 0 | 0 | 0 | 0 | 5 | 3 |
| PFAS | 0 | 0 | 0 | 0 | 0 | 0 | 0 | 2 | 0 |
| PFDN2 | 0 | 0 | 0 | 0 | 0 | 0 | 0 | 1 | 0 |
| PFKL | 0 | 0 | 0 | 0 | 0 | 0 | 1 | 4 | 3 |
| PFKM | 0 | 0 | 0 | 0 | 0 | 0 | 0 | 1 | 0 |
| PFKP | 0 | 0 | 0 | 0 | 0 | 0 | 0 | 4 | 2 |
| PFN2 | 0 | 0 | 0 | 0 | 1 | 0 | 0 | 1 | 1 |
| PHACTR4 | 0 | 0 | 0 | 0 | 0 | 1 | 0 | 7 | 7 |
| PHAX | 0 | 0 | 0 | 0 | 1 | 0 | 1 | 4 | 5 |
| PHC3;PHC2 | 0 | 0 | 0 | 0 | 0 | 0 | 0 | 1 | 0 |
| PHF3 | 0 | 0 | 0 | 0 | 0 | 0 | 0 | 3 | 1 |
| PIK3C2A | 0 | 0 | 0 | 0 | 1 | 0 | 0 | 2 | 2 |
| PIK3R2 | 0 | 0 | 0 | 0 | 0 | 0 | 0 | 1 | 0 |
| PIP4K2B | 0 | 0 | 0 | 0 | 0 | 0 | 0 | 1 | 0 |
| PITPNC1 | 0 | 0 | 0 | 0 | 0 | 0 | 0 | 0 | 4 |
| PKN2 | 0 | 0 | 0 | 0 | 0 | 0 | 0 | 3 | 1 |
| PLAA | 0 | 0 | 0 | 0 | 0 | 0 | 1 | 2 | 1 |

| | | | | | | | | | |
|---|---|---|---|---|---|---|---|---|---|
| PLAUR | 0 | 0 | 0 | 0 | 0 | 0 | 0 | 0 | 1 |
| PLRG1 | 0 | 0 | 0 | 0 | 0 | 0 | 0 | 2 | 0 |
| PML | 0 | 0 | 0 | 0 | 5 | 10 | 12 | 17 | 15 |
| PMS1 | 0 | 0 | 0 | 0 | 1 | 1 | 0 | 4 | 3 |
| PMS2 | 0 | 0 | 0 | 0 | 1 | 0 | 0 | 1 | 1 |
| PNISR | 0 | 0 | 0 | 0 | 2 | 1 | 0 | 4 | 3 |
| PNN | 0 | 0 | 0 | 0 | 6 | 4 | 2 | 14 | 18 |
| POLA1 | 0 | 0 | 0 | 0 | 0 | 0 | 0 | 3 | 0 |
| POLA2 | 0 | 0 | 0 | 0 | 0 | 1 | 3 | 0 | 0 |
| POLD1 | 0 | 0 | 0 | 0 | 2 | 11 | 10 | 6 | 6 |
| POLD3 | 0 | 0 | 0 | 0 | 0 | 1 | 0 | 3 | 2 |
| POLDIP3;PDIP46 | 0 | 0 | 0 | 0 | 0 | 0 | 0 | 2 | 0 |
| POLH | 0 | 0 | 0 | 0 | 0 | 0 | 0 | 1 | 1 |
| POLR2C | 0 | 0 | 0 | 0 | 0 | 1 | 0 | 0 | 0 |
| POM121C;POM121 | 0 | 0 | 0 | 0 | 0 | 0 | 0 | 1 | 1 |
| POP1 | 0 | 0 | 0 | 0 | 4 | 2 | 3 | 7 | 7 |
| POP4 | 0 | 0 | 0 | 0 | 0 | 1 | 1 | 0 | 1 |
| POP5 | 0 | 0 | 0 | 0 | 1 | 0 | 0 | 1 | 1 |
| PPIH | 0 | 0 | 0 | 0 | 0 | 0 | 0 | 1 | 1 |
| PPIL1 | 0 | 0 | 0 | 0 | 0 | 0 | 0 | 1 | 0 |
| PPIL4 | 0 | 0 | 0 | 0 | 1 | 1 | 1 | 0 | 2 |
| PPM1G | 0 | 0 | 0 | 0 | 1 | 0 | 2 | 4 | 4 |
| PPP1R10 | 0 | 0 | 0 | 0 | 15 | 9 | 9 | 16 | 17 |
| PPP1R12C | 0 | 0 | 0 | 0 | 0 | 0 | 0 | 2 | 1 |
| PPP1R2;PPP1R2P3 | 0 | 0 | 0 | 0 | 0 | 0 | 0 | 1 | 1 |
| PPP1R7 | 0 | 0 | 0 | 0 | 0 | 0 | 1 | 1 | 0 |
| PPP1R9B | 0 | 0 | 0 | 0 | 7 | 11 | 5 | 6 | 9 |
| PPP2R2A;PPP2R2D | 0 | 0 | 0 | 0 | 0 | 1 | 1 | 0 | 1 |
| PPP2R5D;PPP2R5C | 0 | 0 | 0 | 0 | 0 | 0 | 1 | 3 | 2 |
| PPP3CA;PPP3CB | 0 | 0 | 0 | 0 | 1 | 0 | 0 | 1 | 0 |
| PPP4C | 0 | 0 | 0 | 0 | 1 | 0 | 0 | 4 | 1 |
| PPP4R1 | 0 | 0 | 0 | 0 | 0 | 0 | 1 | 0 | 0 |
| PPP4R2 | 0 | 0 | 0 | 0 | 0 | 2 | 2 | 4 | 3 |
| PRAF2 | 0 | 0 | 0 | 0 | 0 | 1 | 0 | 1 | 1 |
| PRDX5 | 0 | 0 | 0 | 0 | 0 | 0 | 1 | 1 | 0 |
| PRIM2 | 0 | 0 | 0 | 0 | 0 | 0 | 0 | 1 | 1 |
| PRKAR2A | 0 | 0 | 0 | 0 | 0 | 1 | 1 | 6 | 1 |
| PRMT1 | 0 | 0 | 0 | 0 | 3 | 4 | 2 | 5 | 4 |
| PRMT9 | 0 | 0 | 0 | 0 | 0 | 0 | 1 | 0 | 0 |
| PRNP | 0 | 0 | 0 | 0 | 0 | 0 | 0 | 0 | 1 |
| PRPF19 | 0 | 0 | 0 | 0 | 1 | 1 | 0 | 7 | 8 |
| PRPF3 | 0 | 0 | 0 | 0 | 1 | 3 | 1 | 12 | 10 |
| PRPF31 | 0 | 0 | 0 | 0 | 0 | 0 | 2 | 2 | 3 |
| PRPF4 | 0 | 0 | 0 | 0 | 4 | 7 | 4 | 11 | 9 |
| PRPF40A | 0 | 0 | 0 | 0 | 2 | 0 | 0 | 8 | 7 |

| | | | | | | | | | |
|---|---|---|---|---|---|---|---|---|---|
| PRPF6 | 0 | 0 | 0 | 0 | 0 | 0 | 0 | 0 | 2 |
| PRPF8 | 0 | 0 | 0 | 0 | 2 | 3 | 2 | 16 | 2 |
| PRPS1;PRPS1L1;PRPS2 | 0 | 0 | 0 | 0 | 0 | 0 | 0 | 1 | 0 |
| PRR16 | 0 | 0 | 0 | 0 | 1 | 1 | 1 | 1 | 1 |
| PRR5L | 0 | 0 | 0 | 0 | 0 | 1 | 0 | 0 | 0 |
| PRRC1 | 0 | 0 | 0 | 0 | 0 | 0 | 0 | 1 | 1 |
| PRRC2A | 0 | 0 | 0 | 0 | 0 | 2 | 2 | 6 | 7 |
| PRRC2B | 0 | 0 | 0 | 0 | 1 | 1 | 1 | 1 | 2 |
| PSIP1 | 0 | 0 | 0 | 0 | 0 | 0 | 0 | 1 | 1 |
| PSMA6 | 0 | 0 | 0 | 0 | 0 | 2 | 1 | 0 | 0 |
| PSMA7 | 0 | 0 | 0 | 0 | 0 | 0 | 1 | 1 | 2 |
| PSMB3 | 0 | 0 | 0 | 0 | 1 | 1 | 0 | 0 | 0 |
| PSMD14 | 0 | 0 | 0 | 0 | 0 | 0 | 0 | 1 | 1 |
| PSMD7 | 0 | 0 | 0 | 0 | 1 | 1 | 1 | 0 | 0 |
| PSME1 | 0 | 0 | 0 | 0 | 0 | 0 | 1 | 0 | 0 |
| PSME3 | 0 | 0 | 0 | 0 | 3 | 6 | 4 | 7 | 4 |
| PSPC1 | 0 | 0 | 0 | 0 | 8 | 5 | 7 | 5 | 6 |
| PTBP1 | 0 | 0 | 0 | 0 | 5 | 5 | 3 | 3 | 6 |
| PTMA | 0 | 0 | 0 | 0 | 0 | 0 | 0 | 1 | 4 |
| PTMS | 0 | 0 | 0 | 0 | 0 | 0 | 0 | 0 | 1 |
| PTPN1 | 0 | 0 | 0 | 0 | 1 | 1 | 2 | 7 | 6 |
| PTRF | 0 | 0 | 0 | 0 | 1 | 1 | 0 | 2 | 4 |
| PUM2;PUM1 | 0 | 0 | 0 | 0 | 0 | 0 | 0 | 0 | 2 |
| PVRL2 | 0 | 0 | 0 | 0 | 0 | 0 | 0 | 1 | 1 |
| PVRL3 | 0 | 0 | 0 | 0 | 0 | 0 | 0 | 0 | 1 |
| QRICH1 | 0 | 0 | 0 | 0 | 0 | 1 | 0 | 0 | 1 |
| QSER1 | 0 | 0 | 0 | 0 | 0 | 0 | 0 | 1 | 0 |
| R3HCC1L | 0 | 0 | 0 | 0 | 2 | 1 | 0 | 3 | 2 |
| R3HDM1 | 0 | 0 | 0 | 0 | 0 | 0 | 0 | 1 | 1 |
| RAB11FIP1;RAB11FIP2 | 0 | 0 | 0 | 0 | 0 | 0 | 0 | 1 | 1 |
| RAB13 | 0 | 0 | 0 | 0 | 1 | 1 | 1 | 1 | 3 |
| RAB14 | 0 | 0 | 0 | 0 | 0 | 0 | 0 | 8 | 4 |
| RAB1A | 0 | 0 | 0 | 0 | 0 | 0 | 0 | 2 | 2 |
| RAB1B | 0 | 0 | 0 | 0 | 1 | 4 | 2 | 7 | 9 |
| RAB21 | 0 | 0 | 0 | 0 | 1 | 1 | 0 | 2 | 6 |
| RAB27B;RAB27A | 0 | 0 | 0 | 0 | 0 | 1 | 1 | 0 | 1 |
| RAB32 | 0 | 0 | 0 | 0 | 0 | 1 | 1 | 2 | 0 |
| RAB34 | 0 | 0 | 0 | 0 | 0 | 0 | 0 | 2 | 2 |
| RAB35 | 0 | 0 | 0 | 0 | 0 | 0 | 0 | 0 | 1 |
| RAB5A | 0 | 0 | 0 | 0 | 0 | 0 | 0 | 1 | 1 |
| RAB5C | 0 | 0 | 0 | 0 | 0 | 0 | 0 | 4 | 3 |
| RABGAP1L | 0 | 0 | 0 | 0 | 1 | 0 | 0 | 2 | 2 |
| RABL3 | 0 | 0 | 0 | 0 | 0 | 0 | 0 | 2 | 1 |
| RAC1;RAC2;RAC3 | 0 | 0 | 0 | 0 | 0 | 1 | 1 | 0 | 3 |
| RAD21 | 0 | 0 | 0 | 0 | 0 | 0 | 0 | 1 | 0 |

| | | | | | | | | | |
|---|---|---|---|---|---|---|---|---|---|
| RAE1 | 0 | 0 | 0 | 0 | 1 | 0 | 0 | 3 | 0 |
| RAF1 | 0 | 0 | 0 | 0 | 0 | 0 | 1 | 0 | 1 |
| RAI1 | 0 | 0 | 0 | 0 | 0 | 0 | 0 | 0 | 2 |
| RAP1GDS1 | 0 | 0 | 0 | 0 | 0 | 1 | 0 | 0 | 0 |
| RARS2 | 0 | 0 | 0 | 0 | 1 | 0 | 0 | 1 | 0 |
| RASA3 | 0 | 0 | 0 | 0 | 0 | 1 | 0 | 1 | 3 |
| RASAL2 | 0 | 0 | 0 | 0 | 0 | 1 | 0 | 2 | 3 |
| RAVER1 | 0 | 0 | 0 | 0 | 0 | 0 | 2 | 1 | 1 |
| RB1 | 0 | 0 | 0 | 0 | 2 | 2 | 0 | 11 | 15 |
| RBBP6 | 0 | 0 | 0 | 0 | 1 | 0 | 1 | 5 | 5 |
| RBBP7 | 0 | 0 | 0 | 0 | 5 | 9 | 8 | 9 | 11 |
| RBFOX2;RBFOX1;RBFOX3 | 0 | 0 | 0 | 0 | 0 | 0 | 0 | 0 | 1 |
| RBM10 | 0 | 0 | 0 | 0 | 6 | 6 | 6 | 8 | 11 |
| RBM14;RBM14-RBM... | 0 | 0 | 0 | 0 | 0 | 1 | 1 | 2 | 1 |
| RBM15 | 0 | 0 | 0 | 0 | 0 | 0 | 0 | 0 | 1 |
| RBM17 | 0 | 0 | 0 | 0 | 0 | 3 | 3 | 14 | 5 |
| RBM25 | 0 | 0 | 0 | 0 | 2 | 3 | 0 | 6 | 7 |
| RBM26 | 0 | 0 | 0 | 0 | 3 | 9 | 8 | 13 | 17 |
| RBM27 | 0 | 0 | 0 | 0 | 3 | 2 | 2 | 5 | 7 |
| RBM33 | 0 | 0 | 0 | 0 | 0 | 1 | 2 | 5 | 4 |
| RBM7 | 0 | 0 | 0 | 0 | 1 | 0 | 0 | 1 | 0 |
| RBMX2 | 0 | 0 | 0 | 0 | 0 | 0 | 0 | 1 | 0 |
| RBX1 | 0 | 0 | 0 | 0 | 0 | 0 | 0 | 2 | 2 |
| RC3H2;RC3H1 | 0 | 0 | 0 | 0 | 0 | 0 | 0 | 0 | 1 |
| REEP5 | 0 | 0 | 0 | 0 | 0 | 0 | 0 | 1 | 1 |
| RELL1 | 0 | 0 | 0 | 0 | 1 | 0 | 0 | 1 | 2 |
| REPS1 | 0 | 0 | 0 | 0 | 1 | 1 | 2 | 5 | 5 |
| RER1 | 0 | 0 | 0 | 0 | 1 | 1 | 1 | 1 | 0 |
| RFC4 | 0 | 0 | 0 | 0 | 0 | 0 | 0 | 1 | 2 |
| RFTN1 | 0 | 0 | 0 | 0 | 0 | 0 | 0 | 0 | 1 |
| RFX1 | 0 | 0 | 0 | 0 | 2 | 0 | 0 | 1 | 0 |
| RFX5 | 0 | 0 | 0 | 0 | 0 | 0 | 0 | 1 | 0 |
| RGPD1;RGPD2 | 0 | 0 | 0 | 0 | 1 | 1 | 1 | 0 | 0 |
| RGPD3 | 0 | 0 | 0 | 0 | 1 | 1 | 0 | 1 | 1 |
| RHOA | 0 | 0 | 0 | 0 | 2 | 1 | 0 | 5 | 6 |
| RHOG | 0 | 0 | 0 | 0 | 0 | 0 | 0 | 2 | 1 |
| RIF1 | 0 | 0 | 0 | 0 | 0 | 1 | 0 | 14 | 3 |
| RIN1 | 0 | 0 | 0 | 0 | 1 | 1 | 1 | 0 | 0 |
| RNF20 | 0 | 0 | 0 | 0 | 0 | 0 | 0 | 3 | 1 |
| RNF214 | 0 | 0 | 0 | 0 | 0 | 0 | 0 | 2 | 1 |
| RNF40 | 0 | 0 | 0 | 0 | 0 | 1 | 0 | 0 | 1 |
| RNPS1 | 0 | 0 | 0 | 0 | 0 | 1 | 0 | 5 | 5 |
| ROBO1 | 0 | 0 | 0 | 0 | 0 | 0 | 0 | 0 | 1 |
| RPA1 | 0 | 0 | 0 | 0 | 2 | 1 | 4 | 8 | 12 |
| RPA2 | 0 | 0 | 0 | 0 | 0 | 0 | 0 | 1 | 1 |

| | | | | | | | | | |
|---|---|---|---|---|---|---|---|---|---|
| RPA3 | 0 | 0 | 0 | 0 | 1 | 1 | 2 | 1 | 1 |
| RPL34 | 0 | 0 | 0 | 0 | 0 | 0 | 0 | 1 | 1 |
| RPL36 | 0 | 0 | 0 | 0 | 0 | 0 | 0 | 1 | 0 |
| RPL36A | 0 | 0 | 0 | 0 | 0 | 0 | 0 | 2 | 2 |
| RPL37A | 0 | 0 | 0 | 0 | 0 | 0 | 0 | 1 | 2 |
| RPN1 | 0 | 0 | 0 | 0 | 1 | 1 | 0 | 5 | 1 |
| RPN2 | 0 | 0 | 0 | 0 | 0 | 1 | 1 | 0 | 2 |
| RPP25L | 0 | 0 | 0 | 0 | 0 | 1 | 1 | 0 | 1 |
| RPP30 | 0 | 0 | 0 | 0 | 1 | 4 | 4 | 1 | 1 |
| RPP38 | 0 | 0 | 0 | 0 | 0 | 0 | 0 | 2 | 1 |
| RPP40 | 0 | 0 | 0 | 0 | 0 | 1 | 1 | 1 | 1 |
| RPRD1B | 0 | 0 | 0 | 0 | 1 | 2 | 2 | 0 | 0 |
| RPRD2 | 0 | 0 | 0 | 0 | 0 | 3 | 1 | 8 | 4 |
| RPS21 | 0 | 0 | 0 | 0 | 0 | 0 | 0 | 2 | 1 |
| RPS28 | 0 | 0 | 0 | 0 | 1 | 0 | 0 | 1 | 1 |
| RPS6KC1 | 0 | 0 | 0 | 0 | 0 | 0 | 1 | 1 | 2 |
| RPSA | 0 | 0 | 0 | 0 | 0 | 0 | 0 | 1 | 0 |
| RRAS | 0 | 0 | 0 | 0 | 0 | 0 | 0 | 1 | 1 |
| RRAS2 | 0 | 0 | 0 | 0 | 0 | 0 | 0 | 1 | 2 |
| RRP12 | 0 | 0 | 0 | 0 | 0 | 0 | 0 | 2 | 0 |
| RSRC2 | 0 | 0 | 0 | 0 | 0 | 0 | 0 | 1 | 1 |
| RTCB | 0 | 0 | 0 | 0 | 3 | 3 | 3 | 8 | 7 |
| RTF1 | 0 | 0 | 0 | 0 | 0 | 0 | 0 | 4 | 4 |
| RTN3 | 0 | 0 | 0 | 0 | 1 | 2 | 1 | 7 | 6 |
| S100A16 | 0 | 0 | 0 | 0 | 0 | 0 | 0 | 2 | 1 |
| SAFB2;SAFB | 0 | 0 | 0 | 0 | 0 | 0 | 0 | 4 | 4 |
| SAMD9 | 0 | 0 | 0 | 0 | 0 | 0 | 0 | 2 | 1 |
| SAP130 | 0 | 0 | 0 | 0 | 0 | 1 | 1 | 3 | 4 |
| SAP18 | 0 | 0 | 0 | 0 | 0 | 0 | 0 | 4 | 5 |
| SAP30BP | 0 | 0 | 0 | 0 | 7 | 7 | 7 | 7 | 8 |
| SAR1A;SAR1B | 0 | 0 | 0 | 0 | 0 | 1 | 0 | 0 | 0 |
| SARNP | 0 | 0 | 0 | 0 | 1 | 0 | 0 | 2 | 2 |
| SARS | 0 | 0 | 0 | 0 | 1 | 2 | 1 | 1 | 1 |
| SART1 | 0 | 0 | 0 | 0 | 4 | 4 | 4 | 13 | 10 |
| SART3 | 0 | 0 | 0 | 0 | 4 | 10 | 9 | 14 | 17 |
| SBNO1 | 0 | 0 | 0 | 0 | 0 | 0 | 1 | 3 | 1 |
| SCAF11 | 0 | 0 | 0 | 0 | 1 | 0 | 0 | 1 | 1 |
| SCAF4 | 0 | 0 | 0 | 0 | 1 | 0 | 0 | 3 | 3 |
| SCAMP1 | 0 | 0 | 0 | 0 | 0 | 0 | 0 | 1 | 1 |
| SCAMP3 | 0 | 0 | 0 | 0 | 0 | 0 | 0 | 1 | 0 |
| SCFD2 | 0 | 0 | 0 | 0 | 1 | 2 | 0 | 0 | 0 |
| SCML2 | 0 | 0 | 0 | 0 | 0 | 0 | 0 | 10 | 3 |
| SCRIB | 0 | 0 | 0 | 0 | 2 | 2 | 0 | 1 | 2 |
| SDCCAG3 | 0 | 0 | 0 | 0 | 0 | 0 | 0 | 1 | 1 |
| SEC13 | 0 | 0 | 0 | 0 | 0 | 1 | 2 | 3 | 4 |

| | | | | | | | | | |
|---|---|---|---|---|---|---|---|---|---|
| SEC16A | 0 | 0 | 0 | 0 | 0 | 0 | 0 | 1 | 0 |
| SEC22B | 0 | 0 | 0 | 0 | 0 | 0 | 0 | 2 | 3 |
| SEC23B | 0 | 0 | 0 | 0 | 0 | 2 | 0 | 1 | 3 |
| SEC24D | 0 | 0 | 0 | 0 | 0 | 0 | 1 | 0 | 0 |
| SEC61B | 0 | 0 | 0 | 0 | 0 | 0 | 0 | 1 | 1 |
| SEMA7A | 0 | 0 | 0 | 0 | 0 | 0 | 0 | 0 | 9 |
| SENP3 | 0 | 0 | 0 | 0 | 0 | 0 | 0 | 1 | 1 |
| SENP6 | 0 | 0 | 0 | 0 | 0 | 0 | 1 | 3 | 2 |
| SET;SETSIP | 0 | 0 | 0 | 0 | 4 | 3 | 1 | 4 | 1 |
| SETD2 | 0 | 0 | 0 | 0 | 0 | 0 | 0 | 1 | 0 |
| SF1 | 0 | 0 | 0 | 0 | 3 | 2 | 5 | 10 | 13 |
| SF3A2 | 0 | 0 | 0 | 0 | 0 | 2 | 0 | 5 | 7 |
| SF3A3 | 0 | 0 | 0 | 0 | 1 | 1 | 1 | 6 | 4 |
| SF3B3 | 0 | 0 | 0 | 0 | 6 | 11 | 7 | 22 | 20 |
| SF3B4 | 0 | 0 | 0 | 0 | 1 | 0 | 2 | 6 | 4 |
| SF3B5 | 0 | 0 | 0 | 0 | 0 | 0 | 0 | 1 | 2 |
| SF3B6 | 0 | 0 | 0 | 0 | 0 | 2 | 1 | 6 | 5 |
| SFRP1 | 0 | 0 | 0 | 0 | 0 | 0 | 0 | 0 | 2 |
| SFSWAP | 0 | 0 | 0 | 0 | 0 | 0 | 0 | 0 | 1 |
| SH3BP4 | 0 | 0 | 0 | 0 | 0 | 0 | 0 | 2 | 0 |
| SH3RF1 | 0 | 0 | 0 | 0 | 2 | 1 | 1 | 4 | 4 |
| SHCBP1 | 0 | 0 | 0 | 0 | 0 | 0 | 0 | 1 | 0 |
| SHISA2 | 0 | 0 | 0 | 0 | 0 | 0 | 0 | 1 | 2 |
| SIN3A | 0 | 0 | 0 | 0 | 1 | 1 | 0 | 4 | 1 |
| SIN3B | 0 | 0 | 0 | 0 | 6 | 11 | 9 | 15 | 16 |
| SIPA1L1 | 0 | 0 | 0 | 0 | 0 | 0 | 0 | 0 | 5 |
| SIPA1L3 | 0 | 0 | 0 | 0 | 2 | 2 | 1 | 8 | 4 |
| SIX4 | 0 | 0 | 0 | 0 | 0 | 0 | 0 | 4 | 1 |
| SKA2 | 0 | 0 | 0 | 0 | 0 | 0 | 0 | 1 | 0 |
| SKIV2L2 | 0 | 0 | 0 | 0 | 0 | 1 | 0 | 5 | 1 |
| SKP1 | 0 | 0 | 0 | 0 | 0 | 0 | 0 | 1 | 0 |
| SLC12A4;SLC12A6 | 0 | 0 | 0 | 0 | 0 | 0 | 0 | 1 | 0 |
| SLC25A1 | 0 | 0 | 0 | 0 | 0 | 0 | 0 | 0 | 1 |
| SLC25A13;SLC25A12 | 0 | 0 | 0 | 0 | 0 | 0 | 0 | 0 | 1 |
| SLC35B2 | 0 | 0 | 0 | 0 | 0 | 0 | 0 | 1 | 2 |
| SLC4A1AP | 0 | 0 | 0 | 0 | 0 | 1 | 0 | 2 | 3 |
| SLC4A7 | 0 | 0 | 0 | 0 | 0 | 0 | 0 | 2 | 0 |
| SLC7A5 | 0 | 0 | 0 | 0 | 0 | 0 | 0 | 2 | 2 |
| SLC9A3R2 | 0 | 0 | 0 | 0 | 1 | 1 | 1 | 3 | 2 |
| SLTM | 0 | 0 | 0 | 0 | 0 | 0 | 0 | 1 | 0 |
| SLU7 | 0 | 0 | 0 | 0 | 0 | 0 | 0 | 1 | 1 |
| SMARCA1 | 0 | 0 | 0 | 0 | 0 | 0 | 0 | 2 | 1 |
| SMARCA4;SMARCA2 | 0 | 0 | 0 | 0 | 0 | 0 | 0 | 1 | 0 |
| SMARCC2 | 0 | 0 | 0 | 0 | 0 | 0 | 2 | 0 | 2 |
| SMC1A | 0 | 0 | 0 | 0 | 1 | 1 | 0 | 0 | 0 |

| | | | | | | | | | |
|---|---|---|---|---|---|---|---|---|---|
| SMC2 | 0 | 0 | 0 | 0 | 2 | 2 | 0 | 3 | 2 |
| SMCHD1 | 0 | 0 | 0 | 0 | 1 | 0 | 0 | 2 | 1 |
| SMEK1 | 0 | 0 | 0 | 0 | 1 | 1 | 1 | 6 | 5 |
| SMG9 | 0 | 0 | 0 | 0 | 1 | 0 | 0 | 0 | 0 |
| SMPD4 | 0 | 0 | 0 | 0 | 0 | 1 | 1 | 2 | 0 |
| SMTN | 0 | 0 | 0 | 0 | 0 | 0 | 0 | 1 | 1 |
| SNAP23 | 0 | 0 | 0 | 0 | 0 | 0 | 0 | 0 | 2 |
| SNRNP200 | 0 | 0 | 0 | 0 | 2 | 6 | 3 | 2 | 3 |
| SNRNP70 | 0 | 0 | 0 | 0 | 0 | 1 | 0 | 2 | 3 |
| SNRPA | 0 | 0 | 0 | 0 | 0 | 0 | 0 | 1 | 1 |
| SNRPA1 | 0 | 0 | 0 | 0 | 3 | 6 | 5 | 6 | 8 |
| SNRPB2 | 0 | 0 | 0 | 0 | 2 | 2 | 1 | 8 | 9 |
| SNRPC | 0 | 0 | 0 | 0 | 0 | 0 | 0 | 1 | 1 |
| SNRPD2 | 0 | 0 | 0 | 0 | 0 | 0 | 0 | 2 | 3 |
| SNRPD3 | 0 | 0 | 0 | 0 | 1 | 1 | 0 | 0 | 4 |
| SNRPE | 0 | 0 | 0 | 0 | 0 | 0 | 0 | 1 | 1 |
| SNRPF | 0 | 0 | 0 | 0 | 0 | 0 | 0 | 1 | 1 |
| SNRPG;SNRPGP15 | 0 | 0 | 0 | 0 | 0 | 0 | 0 | 1 | 1 |
| SNRPN;SNRPB | 0 | 0 | 0 | 0 | 0 | 0 | 0 | 1 | 2 |
| SNTB1 | 0 | 0 | 0 | 0 | 1 | 0 | 0 | 2 | 0 |
| SNW1 | 0 | 0 | 0 | 0 | 0 | 4 | 4 | 7 | 5 |
| SNX29 | 0 | 0 | 0 | 0 | 0 | 0 | 0 | 3 | 1 |
| SNX6 | 0 | 0 | 0 | 0 | 1 | 0 | 0 | 1 | 3 |
| SOAT1 | 0 | 0 | 0 | 0 | 0 | 1 | 0 | 2 | 1 |
| SON | 0 | 0 | 0 | 0 | 0 | 0 | 0 | 2 | 3 |
| SOX9;SOX8 | 0 | 0 | 0 | 0 | 0 | 0 | 2 | 3 | 1 |
| SP100 | 0 | 0 | 0 | 0 | 1 | 0 | 0 | 1 | 0 |
| SPACIA2;FAM122B | 0 | 0 | 0 | 0 | 0 | 0 | 1 | 0 | 0 |
| SPAG1 | 0 | 0 | 0 | 0 | 1 | 0 | 1 | 0 | 0 |
| SPAG7 | 0 | 0 | 0 | 0 | 0 | 0 | 0 | 1 | 0 |
| SPATA5 | 0 | 0 | 0 | 0 | 0 | 0 | 0 | 1 | 0 |
| SPATS2L | 0 | 0 | 0 | 0 | 0 | 0 | 0 | 5 | 1 |
| SPCS3 | 0 | 0 | 0 | 0 | 0 | 0 | 0 | 1 | 1 |
| SPDL1 | 0 | 0 | 0 | 0 | 0 | 0 | 1 | 2 | 2 |
| SPECC1 | 0 | 0 | 0 | 0 | 0 | 0 | 1 | 2 | 1 |
| SPECC1L | 0 | 0 | 0 | 0 | 2 | 0 | 1 | 0 | 2 |
| SPEN | 0 | 0 | 0 | 0 | 0 | 0 | 0 | 2 | 0 |
| SRBD1 | 0 | 0 | 0 | 0 | 0 | 0 | 0 | 1 | 0 |
| SREK1 | 0 | 0 | 0 | 0 | 0 | 0 | 0 | 1 | 0 |
| SRGAP2;SRGAP1 | 0 | 0 | 0 | 0 | 0 | 0 | 0 | 1 | 1 |
| SRP54 | 0 | 0 | 0 | 0 | 1 | 0 | 0 | 5 | 2 |
| SRP9 | 0 | 0 | 0 | 0 | 1 | 1 | 0 | 2 | 2 |
| SRPK1 | 0 | 0 | 0 | 0 | 1 | 0 | 0 | 0 | 1 |
| SRPR | 0 | 0 | 0 | 0 | 1 | 0 | 2 | 4 | 5 |
| SRPRB | 0 | 0 | 0 | 0 | 1 | 1 | 1 | 4 | 2 |

| | | | | | | | | | |
|---|---|---|---|---|---|---|---|---|---|
| SRRM1 | 0 | 0 | 0 | 0 | 0 | 0 | 0 | 2 | 0 |
| SRRM2 | 0 | 0 | 0 | 0 | 0 | 0 | 0 | 3 | 7 |
| SRSF11 | 0 | 0 | 0 | 0 | 2 | 7 | 3 | 4 | 6 |
| SRSF6 | 0 | 0 | 0 | 0 | 0 | 0 | 0 | 1 | 1 |
| SRSF7 | 0 | 0 | 0 | 0 | 0 | 0 | 0 | 2 | 2 |
| SSFA2 | 0 | 0 | 0 | 0 | 0 | 1 | 0 | 19 | 5 |
| SSH3 | 0 | 0 | 0 | 0 | 0 | 1 | 1 | 0 | 1 |
| SSR3 | 0 | 0 | 0 | 0 | 1 | 0 | 0 | 0 | 1 |
| STAG1 | 0 | 0 | 0 | 0 | 0 | 0 | 0 | 1 | 1 |
| STAU1 | 0 | 0 | 0 | 0 | 0 | 0 | 0 | 2 | 0 |
| STEAP3 | 0 | 0 | 0 | 0 | 0 | 2 | 3 | 2 | 2 |
| STIM1 | 0 | 0 | 0 | 0 | 1 | 1 | 1 | 4 | 5 |
| STIM2 | 0 | 0 | 0 | 0 | 1 | 0 | 0 | 1 | 1 |
| STIP1 | 0 | 0 | 0 | 0 | 0 | 0 | 0 | 1 | 2 |
| STK24 | 0 | 0 | 0 | 0 | 1 | 1 | 5 | 4 | 3 |
| STK3;STK4 | 0 | 0 | 0 | 0 | 0 | 0 | 0 | 1 | 0 |
| STK33 | 0 | 0 | 0 | 0 | 0 | 0 | 0 | 2 | 1 |
| STMN2 | 0 | 0 | 0 | 0 | 0 | 0 | 0 | 1 | 1 |
| STRN | 0 | 0 | 0 | 0 | 0 | 0 | 0 | 1 | 1 |
| STRN3 | 0 | 0 | 0 | 0 | 0 | 0 | 1 | 1 | 0 |
| STT3A | 0 | 0 | 0 | 0 | 0 | 0 | 1 | 2 | 1 |
| STXBP4 | 0 | 0 | 0 | 0 | 2 | 0 | 0 | 0 | 0 |
| STXBP5 | 0 | 0 | 0 | 0 | 0 | 0 | 0 | 1 | 0 |
| SUB1 | 0 | 0 | 0 | 0 | 2 | 0 | 2 | 0 | 0 |
| SUGP1 | 0 | 0 | 0 | 0 | 4 | 5 | 1 | 12 | 6 |
| SUGP2 | 0 | 0 | 0 | 0 | 1 | 2 | 1 | 7 | 5 |
| SUPT16H | 0 | 0 | 0 | 0 | 0 | 0 | 0 | 0 | 2 |
| SVIL | 0 | 0 | 0 | 0 | 0 | 4 | 4 | 21 | 19 |
| SYDE1 | 0 | 0 | 0 | 0 | 1 | 1 | 1 | 1 | 0 |
| SYNRG | 0 | 0 | 0 | 0 | 0 | 0 | 0 | 4 | 6 |
| TAF15 | 0 | 0 | 0 | 0 | 0 | 0 | 1 | 1 | 2 |
| TAF6 | 0 | 0 | 0 | 0 | 1 | 0 | 0 | 2 | 1 |
| TAF7 | 0 | 0 | 0 | 0 | 0 | 0 | 2 | 2 | 0 |
| TAF9B;TAF9 | 0 | 0 | 0 | 0 | 0 | 0 | 0 | 1 | 1 |
| TALDO1 | 0 | 0 | 0 | 0 | 3 | 1 | 4 | 3 | 5 |
| TANGO6 | 0 | 0 | 0 | 0 | 2 | 0 | 0 | 1 | 4 |
| TAOK3;TAOK1;TAOK2 | 0 | 0 | 0 | 0 | 0 | 0 | 0 | 1 | 2 |
| TBC1D10B | 0 | 0 | 0 | 0 | 0 | 0 | 0 | 1 | 1 |
| TBC1D2 | 0 | 0 | 0 | 0 | 2 | 2 | 3 | 2 | 2 |
| TBC1D24 | 0 | 0 | 0 | 0 | 0 | 0 | 0 | 1 | 0 |
| TBC1D4 | 0 | 0 | 0 | 0 | 0 | 0 | 0 | 0 | 1 |
| TBCB | 0 | 0 | 0 | 0 | 0 | 0 | 0 | 3 | 1 |
| TBCE | 0 | 0 | 0 | 0 | 2 | 1 | 1 | 0 | 0 |
| TBCEL | 0 | 0 | 0 | 0 | 0 | 0 | 0 | 0 | 1 |
| TCEB1 | 0 | 0 | 0 | 0 | 1 | 2 | 2 | 5 | 1 |

| | | | | | | | | | |
|---|---|---|---|---|---|---|---|---|---|
| TCEB3 | 0 | 0 | 0 | 0 | 1 | 2 | 2 | 6 | 4 |
| TCERG1 | 0 | 0 | 0 | 0 | 4 | 5 | 6 | 11 | 7 |
| TCOF1 | 0 | 0 | 0 | 0 | 3 | 6 | 1 | 22 | 20 |
| TDRD3 | 0 | 0 | 0 | 0 | 0 | 0 | 0 | 1 | 3 |
| TES | 0 | 0 | 0 | 0 | 0 | 0 | 0 | 1 | 2 |
| TEX10 | 0 | 0 | 0 | 0 | 0 | 0 | 0 | 1 | 0 |
| TEX264 | 0 | 0 | 0 | 0 | 0 | 0 | 0 | 1 | 2 |
| TFE3;FFAR4 | 0 | 0 | 0 | 0 | 0 | 0 | 0 | 2 | 0 |
| TFG | 0 | 0 | 0 | 0 | 0 | 0 | 0 | 1 | 3 |
| TFIP11 | 0 | 0 | 0 | 0 | 0 | 0 | 0 | 1 | 2 |
| TGFB1I1 | 0 | 0 | 0 | 0 | 1 | 0 | 1 | 0 | 1 |
| THOC1 | 0 | 0 | 0 | 0 | 0 | 0 | 0 | 3 | 1 |
| THOC2 | 0 | 0 | 0 | 0 | 2 | 4 | 4 | 7 | 6 |
| THOC3 | 0 | 0 | 0 | 0 | 0 | 0 | 0 | 0 | 1 |
| THOC5 | 0 | 0 | 0 | 0 | 0 | 0 | 0 | 2 | 0 |
| THOC6 | 0 | 0 | 0 | 0 | 0 | 0 | 0 | 1 | 0 |
| THOC7;NIF3L1BP1 | 0 | 0 | 0 | 0 | 0 | 0 | 0 | 1 | 1 |
| THRAP3 | 0 | 0 | 0 | 0 | 7 | 6 | 8 | 24 | 22 |
| THY1 | 0 | 0 | 0 | 0 | 0 | 0 | 0 | 0 | 3 |
| TIMM50 | 0 | 0 | 0 | 0 | 0 | 0 | 0 | 0 | 1 |
| TKT | 0 | 0 | 0 | 0 | 0 | 0 | 0 | 3 | 2 |
| TLE3 | 0 | 0 | 0 | 0 | 0 | 0 | 0 | 2 | 2 |
| TM9SF3 | 0 | 0 | 0 | 0 | 0 | 1 | 0 | 0 | 0 |
| TMEM199 | 0 | 0 | 0 | 0 | 0 | 0 | 0 | 1 | 1 |
| TMEM256-PLSCR3 | 0 | 0 | 0 | 0 | 0 | 1 | 0 | 0 | 0 |
| TMEM57 | 0 | 0 | 0 | 0 | 0 | 0 | 0 | 1 | 2 |
| TMF1 | 0 | 0 | 0 | 0 | 0 | 0 | 0 | 2 | 0 |
| TMPO | 0 | 0 | 0 | 0 | 8 | 6 | 5 | 13 | 11 |
| TNFRSF11A | 0 | 0 | 0 | 0 | 0 | 0 | 0 | 1 | 0 |
| TNIK | 0 | 0 | 0 | 0 | 1 | 1 | 1 | 0 | 1 |
| TNIP1 | 0 | 0 | 0 | 0 | 3 | 0 | 3 | 4 | 3 |
| TNRC6A | 0 | 0 | 0 | 0 | 1 | 0 | 0 | 6 | 4 |
| TNS1 | 0 | 0 | 0 | 0 | 0 | 0 | 0 | 2 | 0 |
| TOLLIP | 0 | 0 | 0 | 0 | 0 | 0 | 1 | 0 | 0 |
| TOMM22 | 0 | 0 | 0 | 0 | 0 | 0 | 0 | 1 | 1 |
| TOMM34 | 0 | 0 | 0 | 0 | 0 | 0 | 0 | 1 | 0 |
| TOMM40 | 0 | 0 | 0 | 0 | 0 | 0 | 0 | 1 | 0 |
| TOP2B | 0 | 0 | 0 | 0 | 0 | 0 | 0 | 1 | 0 |
| TOP3B | 0 | 0 | 0 | 0 | 0 | 0 | 0 | 1 | 1 |
| TOR1AIP1 | 0 | 0 | 0 | 0 | 6 | 1 | 4 | 9 | 10 |
| TOR1AIP2 | 0 | 0 | 0 | 0 | 0 | 0 | 0 | 1 | 3 |
| TOX4 | 0 | 0 | 0 | 0 | 9 | 16 | 11 | 14 | 14 |
| TP53BP1 | 0 | 0 | 0 | 0 | 2 | 6 | 4 | 19 | 12 |
| TPP2 | 0 | 0 | 0 | 0 | 0 | 2 | 0 | 1 | 1 |
| TPX2 | 0 | 0 | 0 | 0 | 5 | 3 | 7 | 11 | 14 |

| | | | | | | | | | |
|---|---|---|---|---|---|---|---|---|---|
| TRA2B | 0 | 0 | 0 | 0 | 0 | 0 | 0 | 0 | 2 |
| TRAPPC5 | 0 | 0 | 0 | 0 | 0 | 0 | 2 | 0 | 0 |
| TRIM23 | 0 | 0 | 0 | 0 | 0 | 0 | 0 | 0 | 1 |
| TRIM28 | 0 | 0 | 0 | 0 | 1 | 7 | 5 | 14 | 14 |
| TRIM38 | 0 | 0 | 0 | 0 | 0 | 0 | 0 | 0 | 1 |
| TRIM56 | 0 | 0 | 0 | 0 | 0 | 0 | 0 | 2 | 2 |
| TRIP10 | 0 | 0 | 0 | 0 | 2 | 1 | 0 | 0 | 0 |
| TRIP11 | 0 | 0 | 0 | 0 | 0 | 0 | 0 | 1 | 0 |
| TRIP12 | 0 | 0 | 0 | 0 | 1 | 0 | 0 | 5 | 1 |
| TRMT112 | 0 | 0 | 0 | 0 | 0 | 1 | 0 | 1 | 0 |
| TRMT5 | 0 | 0 | 0 | 0 | 0 | 0 | 0 | 0 | 2 |
| TRMT6 | 0 | 0 | 0 | 0 | 0 | 0 | 1 | 2 | 2 |
| TRMT61A | 0 | 0 | 0 | 0 | 0 | 0 | 0 | 1 | 1 |
| TRPC5 | 0 | 0 | 0 | 0 | 0 | 0 | 0 | 0 | 2 |
| TSC1 | 0 | 0 | 0 | 0 | 0 | 0 | 0 | 1 | 1 |
| TSNAX;DISC1 | 0 | 0 | 0 | 0 | 0 | 0 | 0 | 1 | 1 |
| TTN | 0 | 0 | 0 | 0 | 0 | 2 | 0 | 0 | 0 |
| TUBB2A | 0 | 0 | 0 | 0 | 0 | 0 | 0 | 1 | 1 |
| TUBGCP2 | 0 | 0 | 0 | 0 | 1 | 1 | 1 | 1 | 0 |
| TXLNG | 0 | 0 | 0 | 0 | 0 | 0 | 0 | 1 | 0 |
| U2AF1;U2AF1L4 | 0 | 0 | 0 | 0 | 0 | 0 | 0 | 1 | 0 |
| U2AF2 | 0 | 0 | 0 | 0 | 1 | 0 | 0 | 1 | 0 |
| U2SURP | 0 | 0 | 0 | 0 | 16 | 12 | 16 | 24 | 17 |
| UBA1 | 0 | 0 | 0 | 0 | 2 | 0 | 1 | 4 | 2 |
| UBE2J1 | 0 | 0 | 0 | 0 | 1 | 0 | 0 | 1 | 1 |
| UBFD1 | 0 | 0 | 0 | 0 | 0 | 0 | 0 | 1 | 1 |
| UBIAD1 | 0 | 0 | 0 | 0 | 0 | 2 | 1 | 0 | 1 |
| UBTD1;UBTD2 | 0 | 0 | 0 | 0 | 0 | 0 | 0 | 0 | 1 |
| UBXN4 | 0 | 0 | 0 | 0 | 0 | 0 | 0 | 1 | 2 |
| UBXN7 | 0 | 0 | 0 | 0 | 0 | 0 | 0 | 0 | 1 |
| UFL1 | 0 | 0 | 0 | 0 | 0 | 0 | 0 | 3 | 1 |
| UHRF1BP1L | 0 | 0 | 0 | 0 | 0 | 0 | 1 | 0 | 1 |
| UIMC1 | 0 | 0 | 0 | 0 | 0 | 0 | 0 | 2 | 1 |
| UPF1 | 0 | 0 | 0 | 0 | 0 | 0 | 0 | 0 | 1 |
| USP10 | 0 | 0 | 0 | 0 | 1 | 0 | 0 | 4 | 6 |
| USP16 | 0 | 0 | 0 | 0 | 0 | 1 | 0 | 3 | 0 |
| USP24 | 0 | 0 | 0 | 0 | 0 | 0 | 1 | 1 | 0 |
| USP28 | 0 | 0 | 0 | 0 | 0 | 1 | 0 | 1 | 0 |
| USP7 | 0 | 0 | 0 | 0 | 3 | 5 | 4 | 9 | 9 |
| USP8 | 0 | 0 | 0 | 0 | 0 | 0 | 0 | 1 | 0 |
| UTP14A;UTP14C | 0 | 0 | 0 | 0 | 2 | 0 | 0 | 0 | 0 |
| VAMP5 | 0 | 0 | 0 | 0 | 0 | 0 | 0 | 1 | 2 |
| VANGL1 | 0 | 0 | 0 | 0 | 0 | 0 | 0 | 5 | 2 |
| VAPB | 0 | 0 | 0 | 0 | 0 | 5 | 4 | 8 | 5 |
| VCP | 0 | 0 | 0 | 0 | 0 | 2 | 1 | 5 | 3 |

| | | | | | | | | | |
|---|---|---|---|---|---|---|---|---|---|
| VPS25 | 0 | 0 | 0 | 0 | 0 | 1 | 1 | 0 | 0 |
| VPS28 | 0 | 0 | 0 | 0 | 1 | 0 | 0 | 1 | 0 |
| VPS37B | 0 | 0 | 0 | 0 | 0 | 0 | 0 | 1 | 0 |
| VPS45 | 0 | 0 | 0 | 0 | 0 | 1 | 0 | 0 | 0 |
| VPS4A;VPS4B;FIGNL1 | 0 | 0 | 0 | 0 | 0 | 0 | 1 | 0 | 0 |
| VPS51 | 0 | 0 | 0 | 0 | 0 | 0 | 0 | 1 | 0 |
| VPS72 | 0 | 0 | 0 | 0 | 0 | 0 | 0 | 1 | 1 |
| VRK2 | 0 | 0 | 0 | 0 | 0 | 1 | 1 | 3 | 3 |
| WAPAL | 0 | 0 | 0 | 0 | 11 | 11 | 11 | 16 | 14 |
| WASL | 0 | 0 | 0 | 0 | 0 | 0 | 1 | 0 | 0 |
| WBP11 | 0 | 0 | 0 | 0 | 0 | 0 | 2 | 0 | 0 |
| WDFY1 | 0 | 0 | 0 | 0 | 0 | 0 | 0 | 3 | 1 |
| WDHD1 | 0 | 0 | 0 | 0 | 4 | 5 | 2 | 12 | 14 |
| WDR33 | 0 | 0 | 0 | 0 | 0 | 0 | 0 | 2 | 1 |
| WDR43 | 0 | 0 | 0 | 0 | 0 | 0 | 1 | 3 | 0 |
| WDR70 | 0 | 0 | 0 | 0 | 0 | 2 | 0 | 5 | 7 |
| WDR82 | 0 | 0 | 0 | 0 | 1 | 2 | 1 | 5 | 5 |
| WHSC1 | 0 | 0 | 0 | 0 | 0 | 0 | 0 | 1 | 0 |
| WHSC1L1 | 0 | 0 | 0 | 0 | 0 | 0 | 0 | 0 | 1 |
| WIZ | 0 | 0 | 0 | 0 | 0 | 0 | 0 | 1 | 1 |
| WNK1 | 0 | 0 | 0 | 0 | 0 | 0 | 0 | 0 | 1 |
| WRNIP1 | 0 | 0 | 0 | 0 | 0 | 0 | 0 | 1 | 0 |
| WTAP | 0 | 0 | 0 | 0 | 0 | 0 | 0 | 0 | 1 |
| XAB2 | 0 | 0 | 0 | 0 | 1 | 0 | 0 | 5 | 7 |
| XRCC1 | 0 | 0 | 0 | 0 | 2 | 1 | 1 | 4 | 4 |
| XRCC4 | 0 | 0 | 0 | 0 | 0 | 0 | 0 | 1 | 1 |
| XRN2 | 0 | 0 | 0 | 0 | 2 | 3 | 5 | 9 | 3 |
| YEATS2 | 0 | 0 | 0 | 0 | 0 | 0 | 0 | 7 | 3 |
| YES1 | 0 | 0 | 0 | 0 | 0 | 0 | 0 | 0 | 4 |
| YLPM1 | 0 | 0 | 0 | 0 | 5 | 5 | 8 | 18 | 28 |
| YTHDF1 | 0 | 0 | 0 | 0 | 0 | 0 | 0 | 2 | 0 |
| YTHDF3 | 0 | 0 | 0 | 0 | 0 | 0 | 0 | 2 | 2 |
| ZBTB21 | 0 | 0 | 0 | 0 | 0 | 0 | 0 | 1 | 0 |
| ZBTB9 | 0 | 0 | 0 | 0 | 0 | 0 | 0 | 0 | 1 |
| ZC2HC1A | 0 | 0 | 0 | 0 | 0 | 0 | 0 | 3 | 2 |
| ZC3H11A | 0 | 0 | 0 | 0 | 3 | 2 | 1 | 8 | 6 |
| ZC3H14 | 0 | 0 | 0 | 0 | 0 | 0 | 0 | 9 | 4 |
| ZC3H18 | 0 | 0 | 0 | 0 | 0 | 0 | 0 | 2 | 2 |
| ZC3HC1 | 0 | 0 | 0 | 0 | 0 | 0 | 0 | 0 | 2 |
| ZCCHC6 | 0 | 0 | 0 | 0 | 0 | 1 | 0 | 2 | 2 |
| ZDHHC5 | 0 | 0 | 0 | 0 | 0 | 0 | 1 | 2 | 3 |
| ZFP91 | 0 | 0 | 0 | 0 | 1 | 2 | 1 | 0 | 0 |
| ZFR | 0 | 0 | 0 | 0 | 0 | 0 | 0 | 5 | 3 |
| ZFYVE9 | 0 | 0 | 0 | 0 | 0 | 2 | 0 | 1 | 0 |
| ZHX3 | 0 | 0 | 0 | 0 | 4 | 1 | 3 | 13 | 12 |

| | | | | | | | | | |
|---|---|---|---|---|---|---|---|---|---|
| ZMYM1 | 0 | 0 | 0 | 0 | 0 | 0 | 0 | 1 | 1 |
| ZMYND8 | 0 | 0 | 0 | 0 | 0 | 0 | 1 | 12 | 12 |
| ZNF131 | 0 | 0 | 0 | 0 | 0 | 0 | 0 | 1 | 2 |
| ZNF148 | 0 | 0 | 0 | 0 | 1 | 1 | 1 | 10 | 6 |
| ZNF207 | 0 | 0 | 0 | 0 | 0 | 1 | 0 | 1 | 0 |
| ZNF280C | 0 | 0 | 0 | 0 | 0 | 0 | 0 | 2 | 1 |
| ZNF281 | 0 | 0 | 0 | 0 | 1 | 2 | 0 | 5 | 4 |
| ZNF318 | 0 | 0 | 0 | 0 | 0 | 0 | 0 | 10 | 6 |
| ZNF451 | 0 | 0 | 0 | 0 | 0 | 0 | 0 | 1 | 3 |
| ZNF512 | 0 | 0 | 0 | 0 | 0 | 0 | 0 | 2 | 2 |
| ZNF592 | 0 | 0 | 0 | 0 | 0 | 0 | 0 | 1 | 1 |
| ZNF598 | 0 | 0 | 0 | 0 | 0 | 0 | 0 | 1 | 1 |
| ZNF609 | 0 | 0 | 0 | 0 | 0 | 0 | 0 | 1 | 1 |
| ZNF638 | 0 | 0 | 0 | 0 | 1 | 1 | 0 | 11 | 4 |
| ZRANB2 | 0 | 0 | 0 | 0 | 0 | 0 | 0 | 2 | 3 |

OPTN-BirA* RPE BioID

| | | Spectral counts | | | | | | |
|---|---|---|---|---|---|---|---|---|
| | | OPTN-BirA* | | BirA* | | | | |
| Gene name | Fold change (FC-B) | A | B | A | B | C | D | E |
| OPTN | 52.28 | 181 | 164 | 2 | 1 | 2 | 4 | 1 |
| MTMR9 | 6.27 | 8 | 5 | 0 | 0 | 0 | 1 | 0 |
| SPATA2 | 5.91 | 7 | 5 | 0 | 0 | 0 | 1 | 0 |
| MTMR6 | 5.08 | 20 | 12 | 0 | 0 | 1 | 4 | 4 |
| KANK1 | 4.11 | 8 | 6 | 0 | 0 | 0 | 2 | 2 |
| CYLD | 4.09 | 29 | 25 | 4 | 5 | 3 | 9 | 8 |
| GNG12 | 3.84 | 6 | 7 | 0 | 0 | 1 | 0 | 3 |
| SPATA2L | 3.77 | 3 | 2 | 0 | 0 | 0 | 0 | 0 |
| IFI35 | 3.64 | 4 | 3 | 0 | 0 | 1 | 0 | 0 |
| DYNLL2 | 3.39 | 3 | 3 | 0 | 0 | 0 | 1 | 0 |
| SEMA7A | 3.31 | 17 | 5 | 0 | 0 | 0 | 0 | 9 |
| CAMK2G | 3.27 | 2 | 2 | 0 | 0 | 0 | 0 | 0 |
| ERC1 | 3.2 | 20 | 18 | 1 | 3 | 3 | 13 | 2 |
| FLOT2 | 3.18 | 9 | 5 | 0 | 0 | 0 | 2 | 4 |
| UACA | 3.18 | 6 | 3 | 0 | 0 | 0 | 0 | 3 |
| TBC1D2 | 3.1 | 9 | 9 | 2 | 2 | 3 | 2 | 2 |
| CDC16 | 2.9 | 3 | 2 | 0 | 0 | 0 | 1 | 0 |
| PSMA5 | 2.78 | 3 | 2 | 0 | 1 | 0 | 0 | 0 |
| GOLGA3 | 2.71 | 11 | 3 | 0 | 1 | 0 | 3 | 2 |
| SH3RF1 | 2.69 | 8 | 10 | 2 | 1 | 1 | 4 | 4 |
| NMI | 2.68 | 1 | 2 | 0 | 0 | 0 | 0 | 0 |
| FLOT1 | 2.66 | 18 | 12 | 0 | 2 | 0 | 9 | 8 |
| TBC1D17 | 2.62 | 4 | 3 | 0 | 0 | 0 | 2 | 1 |
| MME | 2.61 | 2 | 1 | 0 | 0 | 0 | 0 | 0 |

| | | | | | | | | |
|---|---|---|---|---|---|---|---|---|
| RAB18 | 2.61 | 2 | 1 | 0 | 0 | 0 | 0 | 0 |
| CCDC88A | 2.58 | 8 | 4 | 1 | 0 | 0 | 3 | 2 |
| LGALS8 | 2.56 | 2 | 2 | 0 | 0 | 0 | 0 | 1 |
| RPL10A | 2.54 | 4 | 3 | 0 | 1 | 0 | 1 | 1 |
| BCR | 2.52 | 2 | 2 | 0 | 0 | 0 | 1 | 0 |
| DBT | 2.52 | 2 | 2 | 0 | 0 | 0 | 1 | 0 |
| HAUS1 | 2.52 | 2 | 2 | 0 | 0 | 0 | 1 | 0 |
| PHLDB2 | 2.44 | 11 | 6 | 1 | 1 | 1 | 8 | 1 |
| BIRC6 | 2.42 | 5 | 0 | 0 | 0 | 0 | 0 | 0 |
| AIMP1 | 2.4 | 10 | 8 | 3 | 2 | 2 | 2 | 6 |
| DST | 2.33 | 43 | 29 | 3 | 7 | 7 | 24 | 25 |
| FMNL2 | 2.3 | 7 | 1 | 0 | 0 | 0 | 1 | 2 |
| TANC1 | 2.28 | 22 | 21 | 5 | 6 | 7 | 10 | 17 |
| RPL7 | 2.25 | 14 | 10 | 2 | 3 | 4 | 6 | 8 |
| TMEM263 | 2.25 | 10 | 11 | 1 | 2 | 1 | 6 | 8 |
| MAP3K7 | 2.24 | 7 | 6 | 1 | 0 | 0 | 4 | 4 |
| MKL2 | 2.22 | 38 | 33 | 7 | 7 | 4 | 30 | 20 |
| DNAJB1 | 2.17 | 17 | 16 | 9 | 6 | 6 | 7 | 7 |
| NT5E | 2.17 | 27 | 16 | 5 | 5 | 10 | 0 | 17 |
| RRAS2 | 2.15 | 6 | 1 | 0 | 0 | 0 | 1 | 2 |
| CEP85 | 2.14 | 1 | 1 | 0 | 0 | 0 | 0 | 0 |
| CYB5R3 | 2.14 | 1 | 1 | 0 | 0 | 0 | 0 | 0 |
| KIF1B;KIF1A | 2.14 | 1 | 1 | 0 | 0 | 0 | 0 | 0 |
| RPL35 | 2.14 | 1 | 1 | 0 | 0 | 0 | 0 | 0 |
| SFN | 2.14 | 1 | 1 | 0 | 0 | 0 | 0 | 0 |
| LRRFIP2 | 2.12 | 3 | 4 | 1 | 1 | 1 | 0 | 2 |
| BCAR1 | 2.1 | 1 | 2 | 0 | 0 | 0 | 0 | 1 |
| CEP192 | 2.1 | 5 | 2 | 0 | 0 | 0 | 4 | 0 |
| NEDD1 | 2.1 | 6 | 7 | 0 | 1 | 0 | 6 | 3 |
| GNAI2 | 2.09 | 24 | 13 | 2 | 6 | 1 | 0 | 22 |
| LDHB | 2.09 | 5 | 2 | 0 | 1 | 0 | 1 | 2 |
| RPL34 | 2.08 | 2 | 2 | 0 | 0 | 0 | 1 | 1 |
| RPL36A | 2.05 | 3 | 3 | 0 | 0 | 0 | 2 | 2 |
| GIT2 | 2.04 | 2 | 5 | 0 | 2 | 0 | 2 | 0 |
| HAUS4 | 2.04 | 2 | 2 | 0 | 0 | 0 | 2 | 0 |
| ZBTB33 | 2.03 | 18 | 15 | 2 | 2 | 1 | 15 | 12 |
| ACTB | 2.02 | 6 | 4 | 1 | 2 | 2 | 2 | 3 |
| PPIA | 2.02 | 4 | 4 | 1 | 2 | 0 | 2 | 1 |
| SEC16A | 2.01 | 2 | 1 | 0 | 0 | 0 | 1 | 0 |
| VPS13C | 2.01 | 2 | 1 | 0 | 0 | 0 | 1 | 0 |
| BCAR3 | 1.98 | 1 | 3 | 0 | 0 | 0 | 1 | 1 |
| KIF11 | 1.98 | 3 | 0 | 0 | 0 | 0 | 0 | 0 |
| LYZ | 1.98 | 3 | 0 | 0 | 0 | 0 | 0 | 0 |
| ROCK2 | 1.98 | 8 | 5 | 2 | 3 | 2 | 4 | 2 |
| RPL37 | 1.98 | 3 | 0 | 0 | 0 | 0 | 0 | 0 |

| | | | | | | | | |
|---|---|---|---|---|---|---|---|---|
| CASC5 | 1.96 | 9 | 18 | 1 | 4 | 0 | 11 | 8 |
| KIAA1671 | 1.96 | 17 | 16 | 5 | 5 | 4 | 13 | 11 |
| TTK | 1.96 | 3 | 2 | 0 | 0 | 1 | 1 | 1 |
| ATP5C1 | 1.95 | 6 | 4 | 0 | 1 | 1 | 3 | 4 |
| GAPVD1 | 1.95 | 16 | 13 | 5 | 6 | 7 | 11 | 6 |
| RPL18 | 1.95 | 4 | 3 | 0 | 0 | 1 | 3 | 1 |
| RPL4 | 1.95 | 22 | 17 | 4 | 5 | 5 | 18 | 13 |
| CAMK2D | 1.94 | 9 | 8 | 3 | 5 | 4 | 2 | 3 |
| CCP110 | 1.93 | 2 | 1 | 0 | 0 | 1 | 0 | 0 |
| FLII | 1.93 | 34 | 14 | 9 | 10 | 11 | 2 | 17 |
| LGALS1 | 1.92 | 11 | 8 | 1 | 3 | 3 | 5 | 9 |
| CEP44 | 1.91 | 6 | 4 | 1 | 1 | 0 | 3 | 4 |
| DYNLL1 | 1.91 | 11 | 14 | 6 | 8 | 4 | 5 | 5 |
| GNB2 | 1.91 | 16 | 9 | 4 | 3 | 2 | 1 | 14 |
| YWHAH | 1.91 | 6 | 3 | 0 | 0 | 0 | 5 | 2 |
| EXOC4 | 1.89 | 7 | 9 | 4 | 1 | 2 | 5 | 4 |
| PLEC | 1.89 | 108 | 65 | 17 | 27 | 30 | 65 | 72 |
| PABPC4 | 1.88 | 3 | 2 | 0 | 1 | 1 | 1 | 1 |
| RPL12 | 1.87 | 6 | 5 | 0 | 0 | 0 | 4 | 6 |
| SQSTM1 | 1.87 | 8 | 6 | 0 | 4 | 1 | 4 | 4 |
| ARPC3 | 1.86 | 10 | 8 | 4 | 3 | 3 | 2 | 8 |
| RPL28 | 1.85 | 5 | 4 | 0 | 0 | 0 | 3 | 5 |
| ERLIN1 | 1.83 | 4 | 3 | 1 | 2 | 0 | 1 | 2 |
| MYO6 | 1.82 | 80 | 45 | 23 | 17 | 27 | 0 | 64 |
| CAMSAP1 | 1.81 | 26 | 18 | 3 | 6 | 7 | 18 | 19 |
| PTPN23 | 1.81 | 7 | 9 | 3 | 3 | 5 | 5 | 4 |
| PCCA | 1.79 | 53 | 51 | 20 | 28 | 34 | 35 | 30 |
| GIGYF2 | 1.78 | 59 | 52 | 16 | 20 | 18 | 48 | 48 |
| OCRL | 1.76 | 2 | 3 | 0 | 0 | 0 | 3 | 1 |
| RPL13A | 1.76 | 7 | 4 | 1 | 0 | 1 | 3 | 6 |
| RPL7A | 1.76 | 9 | 9 | 0 | 3 | 3 | 6 | 9 |
| YWHAG | 1.76 | 6 | 6 | 2 | 3 | 0 | 5 | 3 |
| TBC1D5 | 1.75 | 9 | 2 | 0 | 1 | 1 | 5 | 2 |
| ARPC5 | 1.74 | 3 | 3 | 0 | 1 | 1 | 1 | 3 |
| DYNC1I2 | 1.73 | 10 | 9 | 3 | 5 | 3 | 6 | 8 |
| GAS2L3 | 1.73 | 4 | 0 | 0 | 0 | 0 | 0 | 1 |
| RPL3 | 1.73 | 12 | 8 | 1 | 3 | 0 | 10 | 7 |
| ARHGAP29 | 1.72 | 48 | 34 | 18 | 16 | 11 | 34 | 30 |
| MYH10 | 1.72 | 206 | 142 | 82 | 65 | 95 | 0 | 160 |
| TBC1D15 | 1.72 | 24 | 20 | 10 | 13 | 13 | 16 | 10 |
| ABCG2 | 1.71 | 2 | 0 | 0 | 0 | 0 | 0 | 0 |
| ACSL4 | 1.71 | 2 | 0 | 0 | 0 | 0 | 0 | 0 |
| ASCC1 | 1.71 | 2 | 0 | 0 | 0 | 0 | 0 | 0 |
| CD109 | 1.71 | 2 | 0 | 0 | 0 | 0 | 0 | 0 |
| CKAP5 | 1.71 | 41 | 35 | 16 | 19 | 19 | 34 | 28 |

| | | | | | | | | |
|---|---|---|---|---|---|---|---|---|
| LRRC49 | 1.71 | 5 | 6 | 2 | 3 | 3 | 1 | 4 |
| NECAP2 | 1.71 | 2 | 0 | 0 | 0 | 0 | 0 | 0 |
| RAP2B;RAP2A | 1.71 | 2 | 0 | 0 | 0 | 0 | 0 | 0 |
| RBM12 | 1.71 | 2 | 0 | 0 | 0 | 0 | 0 | 0 |
| RGS20 | 1.71 | 2 | 0 | 0 | 0 | 0 | 0 | 0 |
| RHOC | 1.71 | 2 | 0 | 0 | 0 | 0 | 0 | 0 |
| TTF2 | 1.71 | 2 | 0 | 0 | 0 | 0 | 0 | 0 |
| TRIP11 | 1.7 | 4 | 0 | 0 | 0 | 0 | 1 | 0 |
| RABGAP1 | 1.69 | 4 | 2 | 0 | 0 | 0 | 5 | 0 |
| TNRC6B | 1.69 | 16 | 8 | 2 | 1 | 1 | 12 | 10 |
| VDAC3 | 1.69 | 9 | 5 | 3 | 0 | 0 | 3 | 7 |
| ACACB | 1.68 | 1 | 1 | 0 | 0 | 0 | 0 | 1 |
| DYNC1LI1 | 1.68 | 1 | 1 | 0 | 0 | 0 | 0 | 1 |
| GNB2L1 | 1.68 | 9 | 8 | 1 | 1 | 0 | 8 | 9 |
| KIAA1217 | 1.68 | 5 | 6 | 0 | 2 | 0 | 4 | 5 |
| LRRC7 | 1.68 | 1 | 1 | 0 | 0 | 0 | 0 | 1 |
| MYO5A | 1.68 | 42 | 27 | 17 | 12 | 17 | 0 | 33 |
| SMG7 | 1.68 | 4 | 1 | 0 | 1 | 1 | 1 | 1 |
| TRIM23 | 1.68 | 1 | 1 | 0 | 0 | 0 | 0 | 1 |
| MCCC1 | 1.67 | 41 | 43 | 22 | 20 | 26 | 32 | 33 |
| TBK1 | 1.67 | 4 | 4 | 0 | 3 | 1 | 3 | 1 |
| CORO2B | 1.66 | 18 | 13 | 5 | 4 | 10 | 0 | 16 |
| HSPA6;HSPA7 | 1.66 | 2 | 1 | 0 | 0 | 0 | 1 | 1 |
| IFT74 | 1.66 | 2 | 2 | 1 | 0 | 0 | 0 | 2 |
| PSMD14 | 1.66 | 2 | 1 | 0 | 0 | 0 | 1 | 1 |
| RAB1B | 1.66 | 11 | 8 | 1 | 4 | 2 | 7 | 9 |
| VBP1 | 1.66 | 2 | 1 | 0 | 0 | 0 | 1 | 1 |
| CCDC85C | 1.65 | 3 | 3 | 1 | 0 | 1 | 3 | 1 |
| MAP7D3 | 1.65 | 16 | 13 | 1 | 1 | 0 | 21 | 10 |
| ANK3 | 1.64 | 1 | 1 | 0 | 0 | 0 | 1 | 0 |
| ARF5 | 1.64 | 1 | 1 | 0 | 0 | 0 | 1 | 0 |
| CEP55 | 1.64 | 4 | 6 | 1 | 2 | 1 | 4 | 4 |
| PDCD6IP | 1.64 | 1 | 1 | 0 | 0 | 0 | 1 | 0 |
| PSMB6 | 1.64 | 1 | 1 | 0 | 0 | 0 | 1 | 0 |
| RIPK1 | 1.64 | 1 | 1 | 0 | 0 | 0 | 1 | 0 |
| RPL36 | 1.64 | 1 | 1 | 0 | 0 | 0 | 1 | 0 |
| MAGED1 | 1.63 | 3 | 1 | 0 | 0 | 0 | 1 | 2 |
| RPL37A | 1.63 | 3 | 1 | 0 | 0 | 0 | 1 | 2 |
| SHC1 | 1.63 | 3 | 2 | 0 | 1 | 1 | 2 | 1 |
| PHLDB1 | 1.62 | 5 | 4 | 1 | 2 | 1 | 4 | 3 |
| TUBA1C | 1.62 | 3 | 3 | 0 | 0 | 0 | 3 | 3 |
| MCCC2 | 1.61 | 46 | 32 | 19 | 22 | 26 | 31 | 30 |
| RAB32 | 1.6 | 5 | 1 | 0 | 1 | 1 | 2 | 0 |
| SDCBP | 1.6 | 2 | 3 | 1 | 1 | 0 | 2 | 1 |
| C11orf49 | 1.59 | 4 | 3 | 1 | 3 | 0 | 2 | 1 |

| | | | | | | | |
|---|---|---|---|---|---|---|---|
| CDC23 | 1.59 | 6 | 3 | 0 | 1 | 1 | 4 | 4 |
| DIAPH3 | 1.59 | 6 | 1 | 0 | 1 | 1 | 2 | 2 |
| EIF4E2 | 1.58 | 18 | 13 | 4 | 8 | 5 | 13 | 14 |
| MYL6B | 1.58 | 1 | 1 | 0 | 0 | 1 | 0 | 0 |
| PCCB | 1.58 | 35 | 34 | 15 | 25 | 20 | 28 | 25 |
| PSMA6 | 1.58 | 2 | 2 | 0 | 2 | 1 | 0 | 0 |
| RPL27 | 1.58 | 3 | 2 | 1 | 1 | 1 | 2 | 0 |
| PPP1R37 | 1.57 | 4 | 3 | 1 | 1 | 0 | 2 | 4 |
| RPL10 | 1.57 | 9 | 6 | 0 | 0 | 1 | 8 | 8 |
| PITPNC1 | 1.56 | 2 | 2 | 0 | 0 | 0 | 0 | 4 |
| AHNAK2 | 1.55 | 282 | 236 | 121 | 151 | 151 | 223 | 242 |
| CD55 | 1.55 | 3 | 0 | 0 | 0 | 0 | 0 | 1 |
| FLNC | 1.55 | 128 | 123 | 48 | 53 | 55 | 133 | 112 |
| MTHFD1 | 1.55 | 3 | 0 | 0 | 0 | 0 | 0 | 1 |
| OSBPL11 | 1.55 | 7 | 5 | 2 | 2 | 2 | 4 | 7 |
| SEC23B | 1.55 | 3 | 3 | 0 | 2 | 0 | 1 | 3 |
| HN1L | 1.54 | 10 | 5 | 2 | 3 | 4 | 6 | 6 |
| XRCC6 | 1.54 | 3 | 1 | 0 | 0 | 1 | 2 | 0 |
| ZC2HC1A | 1.54 | 3 | 2 | 0 | 0 | 0 | 3 | 2 |
| CAPZA2 | 1.52 | 15 | 15 | 6 | 10 | 7 | 7 | 17 |
| HGS | 1.52 | 11 | 10 | 3 | 4 | 2 | 11 | 10 |
| UBE2N;UBE2NL | 1.52 | 0 | 1 | 0 | 0 | 0 | 0 | 0 |
| FHL3 | 1.5 | 2 | 2 | 0 | 0 | 0 | 3 | 1 |
| MOB2 | 1.5 | 3 | 4 | 1 | 1 | 3 | 1 | 3 |
| RAB14 | 1.48 | 5 | 5 | 0 | 0 | 0 | 8 | 4 |
| RPL18A | 1.48 | 9 | 6 | 3 | 3 | 2 | 7 | 7 |
| RAB5C | 1.46 | 3 | 3 | 0 | 0 | 0 | 4 | 3 |
| EIF4G2 | 1.45 | 25 | 20 | 12 | 12 | 18 | 19 | 17 |
| CEP170 | 1.44 | 97 | 76 | 33 | 49 | 49 | 88 | 85 |
| PDAP1 | 1.44 | 12 | 10 | 3 | 2 | 4 | 11 | 13 |
| SH3D19 | 1.44 | 5 | 5 | 0 | 2 | 2 | 6 | 4 |
| VCPIP1 | 1.44 | 38 | 34 | 23 | 26 | 25 | 28 | 32 |
| DVL3 | 1.43 | 10 | 9 | 4 | 6 | 9 | 5 | 7 |
| HNRNPC | 1.43 | 6 | 3 | 1 | 3 | 0 | 3 | 4 |
| RPL17;RPL17-C18... | 1.43 | 7 | 4 | 0 | 2 | 1 | 6 | 5 |
| RPS27 | 1.43 | 4 | 3 | 1 | 1 | 0 | 4 | 3 |
| RPS27A;UBB;UBC;UBA52 | 1.43 | 14 | 12 | 1 | 3 | 2 | 15 | 16 |
| CHMP2B | 1.41 | 2 | 3 | 0 | 0 | 0 | 2 | 4 |
| DOCK7 | 1.41 | 25 | 18 | 11 | 8 | 13 | 25 | 15 |
| EIF4ENIF1 | 1.41 | 4 | 4 | 0 | 0 | 1 | 4 | 5 |
| ENO1 | 1.41 | 17 | 19 | 8 | 10 | 9 | 17 | 20 |
| AASDHPPT | 1.4 | 1 | 0 | 0 | 0 | 0 | 0 | 0 |
| ANAPC5 | 1.4 | 1 | 0 | 0 | 0 | 0 | 0 | 0 |
| APEH | 1.4 | 1 | 0 | 0 | 0 | 0 | 0 | 0 |
| ARHGEF18 | 1.4 | 1 | 0 | 0 | 0 | 0 | 0 | 0 |

| | | | | | | | | |
|---|---|---|---|---|---|---|---|---|
| ASPH | 1.4 | 1 | 0 | 0 | 0 | 0 | 0 | 0 |
| BPIFA1 | 1.4 | 1 | 0 | 0 | 0 | 0 | 0 | 0 |
| C12orf75;OCC1 | 1.4 | 1 | 0 | 0 | 0 | 0 | 0 | 0 |
| COPZ2 | 1.4 | 1 | 0 | 0 | 0 | 0 | 0 | 0 |
| DBN1 | 1.4 | 49 | 35 | 29 | 23 | 29 | 15 | 39 |
| DHFRL1;DHFR | 1.4 | 1 | 0 | 0 | 0 | 0 | 0 | 0 |
| DVL1;DVL1P1 | 1.4 | 10 | 8 | 3 | 5 | 5 | 8 | 10 |
| ETFA | 1.4 | 1 | 0 | 0 | 0 | 0 | 0 | 0 |
| FBL | 1.4 | 1 | 0 | 0 | 0 | 0 | 0 | 0 |
| FLRT2 | 1.4 | 1 | 0 | 0 | 0 | 0 | 0 | 0 |
| IRS1 | 1.4 | 1 | 0 | 0 | 0 | 0 | 0 | 0 |
| LMCD1 | 1.4 | 1 | 0 | 0 | 0 | 0 | 0 | 0 |
| LTF | 1.4 | 1 | 0 | 0 | 0 | 0 | 0 | 0 |
| MIA3 | 1.4 | 1 | 0 | 0 | 0 | 0 | 0 | 0 |
| NHSL1 | 1.4 | 1 | 0 | 0 | 0 | 0 | 0 | 0 |
| NPLOC4 | 1.4 | 6 | 5 | 1 | 1 | 0 | 6 | 7 |
| PARP9 | 1.4 | 1 | 0 | 0 | 0 | 0 | 0 | 0 |
| PLK1 | 1.4 | 1 | 0 | 0 | 0 | 0 | 0 | 0 |
| PPA1 | 1.4 | 1 | 0 | 0 | 0 | 0 | 0 | 0 |
| PSMG2 | 1.4 | 1 | 0 | 0 | 0 | 0 | 0 | 0 |
| RP2 | 1.4 | 1 | 0 | 0 | 0 | 0 | 0 | 0 |
| RPL22L1 | 1.4 | 1 | 0 | 0 | 0 | 0 | 0 | 0 |
| RPL35A | 1.4 | 3 | 1 | 0 | 0 | 0 | 2 | 2 |
| RPL39P5;RPL39 | 1.4 | 1 | 0 | 0 | 0 | 0 | 0 | 0 |
| SECISBP2 | 1.4 | 1 | 0 | 0 | 0 | 0 | 0 | 0 |
| SH3GLB2 | 1.4 | 1 | 0 | 0 | 0 | 0 | 0 | 0 |
| SLC25A4 | 1.4 | 1 | 0 | 0 | 0 | 0 | 0 | 0 |
| SLC35E1 | 1.4 | 1 | 0 | 0 | 0 | 0 | 0 | 0 |
| TMEM194A | 1.4 | 1 | 0 | 0 | 0 | 0 | 0 | 0 |
| TNPO1 | 1.4 | 1 | 0 | 0 | 0 | 0 | 0 | 0 |
| TUBB8 | 1.4 | 3 | 3 | 1 | 0 | 0 | 3 | 3 |
| ZG16B | 1.4 | 1 | 0 | 0 | 0 | 0 | 0 | 0 |
| ALDH1A3 | 1.39 | 2 | 4 | 0 | 1 | 1 | 3 | 3 |
| FHL2 | 1.39 | 16 | 11 | 1 | 3 | 0 | 14 | 19 |
| SNX29 | 1.39 | 3 | 1 | 0 | 0 | 0 | 3 | 1 |
| USP14 | 1.39 | 10 | 6 | 3 | 4 | 5 | 7 | 8 |
| ABCF2 | 1.38 | 3 | 2 | 0 | 0 | 0 | 3 | 3 |
| ALDH18A1 | 1.38 | 6 | 4 | 0 | 0 | 0 | 7 | 6 |
| CDC42BPA | 1.38 | 1 | 1 | 0 | 0 | 0 | 0 | 2 |
| EIF4G3 | 1.38 | 12 | 10 | 7 | 7 | 6 | 10 | 10 |
| HIST2H3A;HIST2H... | 1.38 | 4 | 4 | 0 | 2 | 1 | 4 | 4 |
| PCBP2 | 1.38 | 8 | 7 | 5 | 4 | 3 | 6 | 7 |
| SPAG9 | 1.38 | 19 | 17 | 7 | 10 | 7 | 19 | 19 |
| TRA2B | 1.38 | 1 | 1 | 0 | 0 | 0 | 0 | 2 |
| CSDE1 | 1.37 | 34 | 23 | 10 | 14 | 6 | 30 | 32 |

| | | | | | | | | |
|---|---|---|---|---|---|---|---|---|
| GNAI3 | 1.37 | 4 | 1 | 0 | 0 | 1 | 0 | 4 |
| RPL15 | 1.37 | 11 | 7 | 1 | 5 | 1 | 9 | 9 |
| SH3GL1 | 1.37 | 7 | 4 | 1 | 1 | 1 | 4 | 9 |
| TJP2 | 1.37 | 33 | 25 | 4 | 10 | 8 | 33 | 36 |
| BET1 | 1.36 | 1 | 1 | 0 | 0 | 0 | 1 | 1 |
| HAUS7 | 1.36 | 1 | 1 | 0 | 0 | 0 | 1 | 1 |
| HIST2H2AA3;HIST2H2AC | 1.36 | 1 | 1 | 0 | 0 | 0 | 1 | 1 |
| ME1 | 1.36 | 1 | 1 | 0 | 0 | 0 | 1 | 1 |
| NDUFA4 | 1.36 | 1 | 1 | 0 | 0 | 0 | 1 | 1 |
| NME1 | 1.36 | 1 | 1 | 0 | 0 | 0 | 1 | 1 |
| RAB11FIP1;RAB11FIP2 | 1.36 | 1 | 1 | 0 | 0 | 0 | 1 | 1 |
| STMN2 | 1.36 | 1 | 1 | 0 | 0 | 0 | 1 | 1 |
| ELP6 | 1.35 | 10 | 7 | 5 | 7 | 6 | 6 | 6 |
| PC | 1.35 | 82 | 72 | 47 | 59 | 66 | 62 | 63 |
| TUBB3 | 1.35 | 6 | 3 | 1 | 0 | 0 | 5 | 5 |
| AZI2 | 1.34 | 1 | 1 | 0 | 0 | 0 | 2 | 0 |
| HLCS | 1.34 | 3 | 3 | 0 | 0 | 0 | 3 | 5 |
| HSPA1B;HSPA1A | 1.34 | 28 | 24 | 19 | 21 | 14 | 20 | 19 |
| MYADM | 1.34 | 2 | 0 | 0 | 0 | 0 | 0 | 1 |
| PLAUR | 1.34 | 2 | 0 | 0 | 0 | 0 | 0 | 1 |
| PRNP | 1.34 | 2 | 0 | 0 | 0 | 0 | 0 | 1 |
| PSMA3 | 1.34 | 2 | 0 | 0 | 0 | 0 | 0 | 1 |
| RAB35 | 1.34 | 2 | 0 | 0 | 0 | 0 | 0 | 1 |
| RPLP0;RPLP0P6 | 1.34 | 6 | 4 | 3 | 2 | 1 | 5 | 4 |
| TBC1D4 | 1.34 | 2 | 0 | 0 | 0 | 0 | 0 | 1 |
| TNS1 | 1.34 | 1 | 1 | 0 | 0 | 0 | 2 | 0 |
| WNK1 | 1.34 | 2 | 0 | 0 | 0 | 0 | 0 | 1 |
| CRKL | 1.33 | 9 | 8 | 5 | 5 | 5 | 7 | 10 |
| DVL2 | 1.33 | 11 | 8 | 3 | 4 | 2 | 10 | 12 |
| FLNB | 1.33 | 157 | 118 | 86 | 103 | 111 | 125 | 142 |
| INPP5F | 1.33 | 18 | 15 | 6 | 11 | 6 | 16 | 18 |
| RAC1;RAC2;RAC3 | 1.33 | 4 | 1 | 0 | 1 | 1 | 0 | 3 |
| RUFY1 | 1.33 | 3 | 3 | 1 | 2 | 2 | 3 | 1 |
| USP15 | 1.33 | 30 | 35 | 20 | 27 | 25 | 26 | 32 |
| BAG2 | 1.32 | 2 | 0 | 0 | 0 | 0 | 1 | 0 |
| PPP2R2A;PPP2R2D | 1.32 | 2 | 1 | 0 | 1 | 1 | 0 | 1 |
| S100A10 | 1.32 | 6 | 4 | 2 | 3 | 1 | 5 | 5 |
| BABAM1 | 1.31 | 3 | 4 | 1 | 2 | 3 | 3 | 3 |
| FOCAD | 1.31 | 1 | 1 | 0 | 1 | 0 | 0 | 1 |
| RBM14;RBM14-RBM... | 1.31 | 3 | 1 | 0 | 1 | 1 | 2 | 1 |
| RPL38 | 1.31 | 1 | 1 | 0 | 1 | 0 | 0 | 1 |
| SLC25A3 | 1.31 | 7 | 5 | 3 | 4 | 4 | 5 | 7 |
| ARPC1B | 1.3 | 14 | 10 | 5 | 9 | 11 | 5 | 11 |
| FN1 | 1.3 | 14 | 2 | 0 | 0 | 0 | 8 | 8 |
| PELO | 1.3 | 1 | 1 | 0 | 0 | 1 | 1 | 0 |

| | | | | | | | | |
|---|---|---|---|---|---|---|---|---|
| SH3KBP1 | 1.3 | 34 | 27 | 18 | 21 | 16 | 34 | 30 |
| ASAP1 | 1.29 | 7 | 6 | 2 | 6 | 4 | 4 | 7 |
| DNAJC13 | 1.29 | 2 | 2 | 1 | 0 | 0 | 0 | 4 |
| MYO1B | 1.29 | 36 | 25 | 17 | 12 | 27 | 0 | 34 |
| LASP1 | 1.28 | 11 | 8 | 2 | 3 | 6 | 9 | 12 |
| PCBP1 | 1.28 | 15 | 11 | 6 | 7 | 4 | 14 | 16 |
| RTN3 | 1.28 | 7 | 4 | 1 | 2 | 1 | 7 | 6 |
| STK38 | 1.28 | 14 | 12 | 7 | 9 | 9 | 12 | 16 |
| VPS28 | 1.28 | 4 | 0 | 1 | 0 | 0 | 1 | 0 |
| XRN1 | 1.28 | 52 | 54 | 36 | 40 | 45 | 52 | 53 |
| ANAPC1 | 1.27 | 2 | 2 | 1 | 1 | 1 | 2 | 2 |
| PGAM5 | 1.27 | 11 | 5 | 4 | 5 | 5 | 8 | 8 |
| PSMB5 | 1.27 | 2 | 0 | 0 | 0 | 1 | 0 | 0 |
| SFRP1 | 1.27 | 3 | 0 | 0 | 0 | 0 | 0 | 2 |
| SKA1 | 1.27 | 2 | 2 | 1 | 1 | 1 | 2 | 2 |
| ACACA | 1.26 | 160 | 149 | 102 | 123 | 146 | 108 | 116 |
| COL12A1 | 1.26 | 2 | 0 | 0 | 1 | 0 | 0 | 0 |
| HAUS6 | 1.26 | 12 | 3 | 2 | 5 | 1 | 7 | 5 |
| HSPA9 | 1.26 | 19 | 18 | 9 | 11 | 10 | 24 | 19 |
| SSH1 | 1.26 | 14 | 4 | 0 | 2 | 4 | 10 | 8 |
| TPGS1 | 1.26 | 4 | 4 | 3 | 4 | 2 | 2 | 1 |
| USO1 | 1.26 | 15 | 6 | 5 | 7 | 8 | 6 | 11 |
| CAP1 | 1.25 | 28 | 16 | 16 | 18 | 16 | 19 | 19 |
| TNRC6A | 1.25 | 5 | 3 | 1 | 0 | 0 | 6 | 4 |
| TUFM | 1.25 | 12 | 10 | 4 | 3 | 3 | 13 | 15 |
| CBL | 1.24 | 4 | 3 | 0 | 2 | 1 | 2 | 6 |
| CRK | 1.24 | 10 | 10 | 2 | 9 | 7 | 10 | 10 |
| ERCC6L | 1.24 | 7 | 7 | 2 | 2 | 3 | 9 | 9 |
| LMNA | 1.24 | 28 | 22 | 15 | 12 | 10 | 28 | 28 |
| CAPZA1 | 1.23 | 33 | 25 | 22 | 25 | 23 | 14 | 27 |
| EIF3F | 1.23 | 5 | 3 | 2 | 3 | 4 | 3 | 2 |
| FAM175B | 1.23 | 3 | 1 | 0 | 0 | 0 | 3 | 2 |
| MPRIP | 1.23 | 39 | 29 | 22 | 29 | 28 | 20 | 38 |
| ANKRD50 | 1.22 | 3 | 1 | 0 | 0 | 0 | 4 | 1 |
| DAB2 | 1.22 | 11 | 11 | 6 | 4 | 2 | 12 | 14 |
| EIF4G1 | 1.22 | 64 | 52 | 42 | 50 | 45 | 59 | 55 |
| HIST1H2BI;HIST1H2BN;HIST1H2BL | 1.22 | 7 | 8 | 4 | 7 | 5 | 7 | 8 |
| KIAA1524 | 1.22 | 12 | 12 | 10 | 7 | 9 | 12 | 9 |
| LIMD1 | 1.22 | 4 | 3 | 1 | 1 | 1 | 5 | 4 |
| MYO1C | 1.22 | 78 | 51 | 49 | 58 | 33 | 0 | 60 |
| NFKB2 | 1.22 | 3 | 3 | 2 | 2 | 3 | 2 | 2 |
| RPL30 | 1.22 | 6 | 4 | 3 | 4 | 0 | 2 | 6 |
| RPS10 | 1.22 | 2 | 1 | 1 | 1 | 1 | 1 | 0 |
| SEC23IP | 1.22 | 11 | 7 | 6 | 6 | 7 | 10 | 8 |
| YWHAQ | 1.22 | 4 | 3 | 3 | 1 | 1 | 4 | 2 |

| | | | | | | | | |
|---|---|---|---|---|---|---|---|---|
| NRBF2 | 1.21 | 6 | 5 | 4 | 1 | 5 | 5 | 4 |
| THY1 | 1.21 | 4 | 0 | 0 | 0 | 0 | 0 | 3 |
| TOMM22 | 1.21 | 0 | 2 | 0 | 0 | 0 | 1 | 1 |
| ARPC1A | 1.2 | 9 | 4 | 4 | 6 | 4 | 2 | 6 |
| CYFIP2 | 1.2 | 1 | 1 | 1 | 1 | 0 | 0 | 0 |
| HSP90B1 | 1.2 | 3 | 1 | 0 | 1 | 1 | 2 | 2 |
| OSBPL9 | 1.2 | 1 | 1 | 1 | 0 | 1 | 0 | 0 |
| SLK | 1.2 | 52 | 36 | 24 | 29 | 38 | 44 | 51 |
| DLST | 1.19 | 3 | 2 | 2 | 2 | 1 | 2 | 2 |
| EIF3G | 1.19 | 3 | 4 | 2 | 2 | 1 | 4 | 4 |
| GIT1 | 1.19 | 0 | 1 | 0 | 0 | 0 | 0 | 1 |
| GOLGA4 | 1.19 | 0 | 1 | 0 | 0 | 0 | 0 | 1 |
| MAT2A | 1.19 | 0 | 1 | 0 | 0 | 0 | 0 | 1 |
| PFKP | 1.19 | 2 | 2 | 0 | 0 | 0 | 4 | 2 |
| PLEKHA5 | 1.19 | 21 | 13 | 6 | 11 | 6 | 24 | 16 |
| RPL11 | 1.19 | 9 | 4 | 3 | 3 | 2 | 7 | 8 |
| RPL27A | 1.19 | 6 | 3 | 4 | 3 | 2 | 3 | 4 |
| RPS8 | 1.19 | 9 | 9 | 4 | 9 | 6 | 8 | 10 |
| TAF15 | 1.19 | 2 | 1 | 0 | 0 | 1 | 1 | 2 |
| ACTC1;ACTA1 | 1.18 | 21 | 10 | 6 | 7 | 18 | 1 | 18 |
| EIF4B | 1.18 | 42 | 25 | 15 | 16 | 17 | 40 | 46 |
| PPP1R13L | 1.18 | 6 | 7 | 6 | 3 | 7 | 3 | 4 |
| RPS5 | 1.18 | 3 | 3 | 0 | 2 | 2 | 3 | 4 |
| USP47 | 1.18 | 5 | 1 | 0 | 1 | 2 | 2 | 3 |
| APPL2 | 1.17 | 0 | 1 | 0 | 0 | 0 | 1 | 0 |
| BLVRA | 1.17 | 0 | 1 | 0 | 0 | 0 | 1 | 0 |
| DLGAP5 | 1.17 | 11 | 11 | 4 | 6 | 3 | 17 | 12 |
| DYNLT1 | 1.17 | 0 | 1 | 0 | 0 | 0 | 1 | 0 |
| FNBP1 | 1.17 | 0 | 1 | 0 | 0 | 0 | 1 | 0 |
| PIP4K2A | 1.17 | 1 | 2 | 1 | 0 | 0 | 2 | 1 |
| RHEB | 1.17 | 2 | 3 | 2 | 2 | 2 | 1 | 1 |
| RPL21 | 1.17 | 6 | 5 | 2 | 0 | 1 | 7 | 8 |
| STT3A | 1.17 | 2 | 1 | 0 | 0 | 1 | 2 | 1 |
| STXBP5 | 1.17 | 0 | 1 | 0 | 0 | 0 | 1 | 0 |
| VDAC2 | 1.17 | 13 | 10 | 6 | 3 | 2 | 15 | 14 |
| ZFYVE1 | 1.17 | 0 | 1 | 0 | 0 | 0 | 1 | 0 |
| BCL2L2;BCL2L2-P... | 1.16 | 1 | 1 | 0 | 0 | 0 | 1 | 2 |
| HSPD1 | 1.16 | 5 | 1 | 0 | 2 | 0 | 3 | 2 |
| KIAA1244 | 1.16 | 4 | 0 | 0 | 0 | 0 | 3 | 0 |
| PSMC1 | 1.16 | 7 | 4 | 2 | 6 | 3 | 5 | 5 |
| SNX1 | 1.16 | 10 | 11 | 10 | 8 | 6 | 10 | 10 |
| TAB2 | 1.16 | 3 | 1 | 0 | 1 | 1 | 3 | 1 |
| ANKRD17 | 1.15 | 11 | 10 | 7 | 5 | 6 | 15 | 10 |
| ARFGAP3 | 1.15 | 6 | 3 | 3 | 3 | 4 | 3 | 5 |
| BIN3 | 1.15 | 7 | 4 | 3 | 1 | 1 | 9 | 4 |

| | | | | | | | | |
|---|---|---|---|---|---|---|---|---|
| CDC2;CDK1 | 1.15 | 3 | 1 | 0 | 1 | 2 | 0 | 2 |
| DYNC1H1 | 1.15 | 51 | 23 | 31 | 35 | 23 | 23 | 25 |
| GAPDH | 1.15 | 7 | 5 | 4 | 4 | 3 | 6 | 8 |
| NUP88 | 1.15 | 7 | 2 | 1 | 2 | 2 | 6 | 4 |
| RPS16;ZNF90 | 1.15 | 8 | 4 | 6 | 3 | 1 | 6 | 4 |
| SCYL2 | 1.15 | 1 | 3 | 1 | 1 | 2 | 2 | 1 |
| TRIM25 | 1.15 | 15 | 9 | 7 | 10 | 11 | 13 | 13 |
| AAK1 | 1.14 | 19 | 16 | 13 | 14 | 14 | 16 | 24 |
| ABLIM1 | 1.14 | 5 | 1 | 1 | 1 | 2 | 0 | 4 |
| EFHD2 | 1.14 | 21 | 15 | 15 | 13 | 17 | 17 | 19 |
| PICALM | 1.14 | 7 | 5 | 4 | 3 | 4 | 7 | 7 |
| SIPA1L3 | 1.14 | 4 | 5 | 2 | 2 | 1 | 8 | 4 |
| SEPT2 | 1.13 | 18 | 15 | 11 | 16 | 12 | 18 | 19 |
| BAG3 | 1.13 | 20 | 9 | 6 | 9 | 5 | 18 | 17 |
| CTTN | 1.13 | 62 | 44 | 29 | 50 | 49 | 52 | 63 |
| S100A6 | 1.13 | 5 | 1 | 2 | 2 | 2 | 1 | 1 |
| SNAP29 | 1.13 | 3 | 4 | 2 | 2 | 3 | 5 | 3 |
| VCL | 1.13 | 27 | 29 | 16 | 12 | 18 | 39 | 34 |
| ARPC5L | 1.12 | 2 | 1 | 0 | 1 | 2 | 0 | 1 |
| MAP1B | 1.12 | 49 | 46 | 19 | 28 | 21 | 71 | 60 |
| PFKL | 1.12 | 3 | 2 | 0 | 0 | 1 | 4 | 3 |
| RAB6A | 1.12 | 2 | 1 | 1 | 1 | 1 | 1 | 2 |
| RPL13 | 1.12 | 2 | 4 | 3 | 1 | 0 | 4 | 1 |
| TMEM33 | 1.12 | 2 | 1 | 1 | 1 | 1 | 1 | 2 |
| ACTR2 | 1.11 | 12 | 6 | 4 | 6 | 9 | 3 | 12 |
| ARF1;ARF3 | 1.11 | 4 | 2 | 0 | 3 | 1 | 3 | 3 |
| ARHGEF7 | 1.11 | 3 | 1 | 1 | 1 | 2 | 2 | 1 |
| CAPZB | 1.11 | 1 | 1 | 0 | 0 | 1 | 1 | 1 |
| CCT4 | 1.11 | 23 | 21 | 20 | 19 | 20 | 23 | 19 |
| CLINT1 | 1.11 | 14 | 13 | 11 | 12 | 13 | 14 | 16 |
| DSTN | 1.11 | 2 | 1 | 0 | 1 | 2 | 1 | 0 |
| EIF3H | 1.11 | 3 | 1 | 0 | 0 | 0 | 3 | 3 |
| EIF4H | 1.11 | 3 | 3 | 0 | 3 | 3 | 2 | 3 |
| GNB4 | 1.11 | 2 | 0 | 0 | 0 | 0 | 0 | 2 |
| PAIP1 | 1.11 | 1 | 1 | 0 | 1 | 0 | 1 | 1 |
| RPL23 | 1.11 | 7 | 6 | 4 | 4 | 2 | 6 | 11 |
| ABCC1 | 1.1 | 1 | 0 | 0 | 0 | 0 | 0 | 1 |
| ACAP2 | 1.1 | 1 | 0 | 0 | 0 | 0 | 0 | 1 |
| AHR | 1.1 | 1 | 0 | 0 | 0 | 0 | 0 | 1 |
| AP4S1 | 1.1 | 1 | 0 | 0 | 0 | 0 | 0 | 1 |
| ARFGAP1 | 1.1 | 10 | 3 | 1 | 2 | 3 | 6 | 10 |
| CAPN2 | 1.1 | 1 | 0 | 0 | 0 | 0 | 0 | 1 |
| CAPNS1 | 1.1 | 1 | 0 | 0 | 0 | 0 | 0 | 1 |
| DOCK10 | 1.1 | 1 | 0 | 0 | 0 | 0 | 0 | 1 |
| FSIP2 | 1.1 | 1 | 0 | 0 | 0 | 0 | 0 | 1 |

| | | | | | | | |
|---|---|---|---|---|---|---|---|
| LARP4B | 1.1 | 1 | 0 | 0 | 0 | 0 | 0 | 1 |
| LRCH1 | 1.1 | 1 | 1 | 0 | 0 | 1 | 2 | 0 |
| PAWR | 1.1 | 2 | 2 | 0 | 0 | 0 | 2 | 5 |
| PCBP3 | 1.1 | 1 | 0 | 0 | 0 | 0 | 0 | 1 |
| RBM15 | 1.1 | 1 | 0 | 0 | 0 | 0 | 0 | 1 |
| RPS26;RPS26P11 | 1.1 | 3 | 3 | 1 | 2 | 1 | 4 | 4 |
| TIMM50 | 1.1 | 1 | 0 | 0 | 0 | 0 | 0 | 1 |
| TRAPPC3 | 1.1 | 1 | 0 | 0 | 0 | 0 | 0 | 1 |
| UBTD1;UBTD2 | 1.1 | 1 | 0 | 0 | 0 | 0 | 0 | 1 |
| UPF1 | 1.1 | 1 | 0 | 0 | 0 | 0 | 0 | 1 |
| VDAC1 | 1.1 | 4 | 5 | 4 | 4 | 4 | 1 | 4 |
| SEPT11 | 1.09 | 16 | 10 | 5 | 7 | 6 | 16 | 21 |
| AP3S1 | 1.09 | 2 | 0 | 0 | 0 | 0 | 1 | 1 |
| ARFIP1 | 1.09 | 2 | 0 | 0 | 0 | 0 | 1 | 1 |
| CCDC6 | 1.09 | 1 | 2 | 0 | 0 | 0 | 3 | 2 |
| KLC1 | 1.09 | 19 | 14 | 14 | 17 | 14 | 16 | 13 |
| MYPN | 1.09 | 2 | 0 | 0 | 0 | 0 | 1 | 1 |
| NFKB1 | 1.09 | 3 | 2 | 2 | 1 | 2 | 3 | 2 |
| RAB11FIP5 | 1.09 | 13 | 9 | 4 | 5 | 7 | 16 | 14 |
| RAPGEF6 | 1.09 | 4 | 2 | 2 | 1 | 0 | 4 | 3 |
| RPS14 | 1.09 | 7 | 5 | 4 | 6 | 6 | 6 | 5 |
| SPTBN1 | 1.09 | 152 | 76 | 81 | 86 | 119 | 15 | 129 |
| TJP1 | 1.09 | 53 | 33 | 22 | 25 | 23 | 58 | 60 |
| AGFG1 | 1.08 | 3 | 6 | 2 | 4 | 4 | 3 | 6 |
| AP4E1 | 1.08 | 1 | 0 | 0 | 0 | 0 | 1 | 0 |
| CCT8 | 1.08 | 46 | 39 | 34 | 40 | 45 | 44 | 48 |
| CLIP1 | 1.08 | 1 | 0 | 0 | 0 | 0 | 1 | 0 |
| CUL4A | 1.08 | 1 | 0 | 0 | 0 | 0 | 1 | 0 |
| FRMD6 | 1.08 | 1 | 0 | 0 | 0 | 0 | 1 | 0 |
| GNB1 | 1.08 | 7 | 4 | 4 | 3 | 6 | 0 | 6 |
| HDLBP | 1.08 | 33 | 28 | 18 | 28 | 29 | 29 | 42 |
| HNRNPA3 | 1.08 | 1 | 0 | 0 | 0 | 0 | 1 | 0 |
| KATNAL1;KATNA1 | 1.08 | 1 | 0 | 0 | 0 | 0 | 1 | 0 |
| KIF5B | 1.08 | 42 | 35 | 38 | 33 | 33 | 37 | 44 |
| MAP4 | 1.08 | 78 | 60 | 56 | 51 | 61 | 85 | 82 |
| NDC80 | 1.08 | 1 | 0 | 0 | 0 | 0 | 1 | 0 |
| PABPC1 | 1.08 | 16 | 11 | 9 | 8 | 10 | 16 | 19 |
| PALLD | 1.08 | 13 | 6 | 4 | 4 | 3 | 14 | 12 |
| PALM2 | 1.08 | 1 | 0 | 0 | 0 | 0 | 1 | 0 |
| PDLIM7 | 1.08 | 12 | 9 | 6 | 5 | 5 | 12 | 17 |
| PFKM | 1.08 | 1 | 0 | 0 | 0 | 0 | 1 | 0 |
| PIP4K2B | 1.08 | 1 | 0 | 0 | 0 | 0 | 1 | 0 |
| PRPS1;PRPS1L1;PRPS2 | 1.08 | 1 | 0 | 0 | 0 | 0 | 1 | 0 |
| PSMA4 | 1.08 | 1 | 2 | 1 | 1 | 2 | 0 | 0 |
| RPS24 | 1.08 | 3 | 3 | 1 | 2 | 3 | 4 | 3 |

| | | | | | | | | |
|---|---|---|---|---|---|---|---|---|
| SNRPD2 | 1.08 | 2 | 1 | 0 | 0 | 0 | 2 | 3 |
| TOMM40 | 1.08 | 1 | 0 | 0 | 0 | 0 | 1 | 0 |
| TOP2B | 1.08 | 1 | 0 | 0 | 0 | 0 | 1 | 0 |
| VIPAS39 | 1.08 | 2 | 1 | 0 | 1 | 3 | 0 | 0 |
| ALMS1 | 1.07 | 2 | 0 | 0 | 0 | 0 | 2 | 0 |
| HSP90AB1 | 1.07 | 17 | 18 | 15 | 16 | 8 | 20 | 22 |
| LPP | 1.07 | 9 | 3 | 4 | 2 | 4 | 8 | 4 |
| RPL31 | 1.07 | 2 | 3 | 2 | 1 | 1 | 3 | 3 |
| RPL9 | 1.07 | 6 | 7 | 6 | 5 | 2 | 5 | 9 |
| RPS23 | 1.07 | 3 | 4 | 1 | 3 | 2 | 3 | 6 |
| RPS28 | 1.07 | 1 | 1 | 1 | 0 | 0 | 1 | 1 |
| SEC61A1 | 1.07 | 4 | 3 | 2 | 3 | 4 | 3 | 4 |
| STAM2 | 1.07 | 6 | 6 | 3 | 4 | 3 | 8 | 9 |
| STAU1 | 1.07 | 2 | 0 | 0 | 0 | 0 | 2 | 0 |
| CAMSAP2 | 1.06 | 7 | 1 | 1 | 1 | 0 | 5 | 4 |
| CAV1 | 1.06 | 3 | 2 | 0 | 0 | 0 | 3 | 6 |
| ERBB2IP | 1.06 | 27 | 24 | 15 | 19 | 14 | 40 | 30 |
| GNAS | 1.06 | 2 | 1 | 0 | 1 | 0 | 0 | 4 |
| KEAP1 | 1.06 | 26 | 14 | 13 | 11 | 21 | 22 | 23 |
| NAV1 | 1.06 | 4 | 6 | 2 | 2 | 2 | 10 | 5 |
| OSTC | 1.06 | 1 | 1 | 0 | 1 | 1 | 1 | 1 |
| SYNJ2 | 1.06 | 43 | 37 | 35 | 37 | 39 | 47 | 40 |
| TRIP4 | 1.06 | 4 | 1 | 1 | 1 | 0 | 4 | 2 |
| UBASH3B | 1.06 | 2 | 4 | 1 | 2 | 0 | 5 | 3 |
| DDX3X;DDX3Y | 1.05 | 20 | 15 | 9 | 15 | 20 | 21 | 18 |
| DTX3L | 1.05 | 2 | 0 | 0 | 0 | 1 | 0 | 1 |
| GEMIN5 | 1.05 | 51 | 42 | 29 | 53 | 31 | 52 | 55 |
| GPRC5A | 1.05 | 4 | 0 | 0 | 0 | 0 | 0 | 4 |
| KIAA1033 | 1.05 | 2 | 0 | 0 | 0 | 1 | 0 | 1 |
| AP3B1 | 1.04 | 8 | 5 | 6 | 7 | 4 | 6 | 6 |
| CCDC50 | 1.04 | 2 | 1 | 0 | 0 | 1 | 2 | 2 |
| CCT3 | 1.04 | 22 | 17 | 14 | 15 | 24 | 22 | 22 |
| CROCC | 1.04 | 1 | 0 | 0 | 0 | 1 | 0 | 0 |
| DCTN1 | 1.04 | 12 | 6 | 4 | 4 | 4 | 15 | 11 |
| ROCK1 | 1.04 | 3 | 1 | 0 | 1 | 1 | 4 | 1 |
| SPC24 | 1.04 | 2 | 1 | 0 | 0 | 1 | 2 | 2 |
| SURF4 | 1.04 | 2 | 2 | 0 | 1 | 2 | 3 | 2 |
| VPS4A;VPS4B;FIGNL1 | 1.04 | 1 | 0 | 0 | 0 | 1 | 0 | 0 |
| WASL | 1.04 | 1 | 0 | 0 | 0 | 1 | 0 | 0 |
| COPB1 | 1.03 | 10 | 13 | 6 | 11 | 10 | 14 | 16 |
| CTIF | 1.03 | 1 | 0 | 0 | 1 | 0 | 0 | 0 |
| DSG2 | 1.03 | 29 | 23 | 12 | 14 | 15 | 38 | 42 |
| EIF4A1 | 1.03 | 12 | 9 | 6 | 11 | 3 | 13 | 13 |
| H2AFY | 1.03 | 1 | 0 | 0 | 1 | 0 | 0 | 0 |
| PATL1 | 1.03 | 1 | 0 | 0 | 1 | 0 | 0 | 0 |

| | | | | | | | | |
|---|---|---|---|---|---|---|---|---|
| RAB21 | 1.03 | 3 | 2 | 1 | 1 | 0 | 2 | 6 |
| RELA | 1.03 | 3 | 2 | 2 | 1 | 2 | 2 | 4 |
| SFXN3 | 1.03 | 2 | 2 | 0 | 2 | 1 | 3 | 2 |
| TM9SF3 | 1.03 | 1 | 0 | 0 | 1 | 0 | 0 | 0 |
| TNKS1BP1 | 1.03 | 15 | 14 | 6 | 8 | 9 | 23 | 21 |
| FAM91A1 | 1.02 | 11 | 7 | 5 | 7 | 7 | 13 | 12 |
| INPPL1 | 1.02 | 6 | 2 | 0 | 2 | 0 | 6 | 5 |
| NCL | 1.02 | 14 | 10 | 7 | 14 | 6 | 14 | 14 |
| RAPH1 | 1.02 | 35 | 33 | 35 | 32 | 25 | 32 | 41 |
| RPL24 | 1.02 | 6 | 5 | 3 | 4 | 6 | 5 | 9 |
| RPL5 | 1.02 | 6 | 3 | 1 | 2 | 6 | 5 | 4 |
| RPS6 | 1.02 | 10 | 10 | 7 | 11 | 11 | 11 | 11 |
| RPS9 | 1.02 | 9 | 8 | 7 | 10 | 8 | 6 | 8 |
| ABLIM3 | 1.01 | 5 | 1 | 2 | 0 | 2 | 2 | 4 |
| ACTR3 | 1.01 | 19 | 13 | 16 | 11 | 16 | 12 | 19 |
| ARF6 | 1.01 | 1 | 1 | 0 | 0 | 0 | 1 | 3 |
| ARPC4 | 1.01 | 9 | 3 | 3 | 7 | 5 | 3 | 6 |
| ASCC2 | 1.01 | 3 | 4 | 1 | 3 | 3 | 4 | 6 |
| DIP2B | 1.01 | 2 | 0 | 1 | 0 | 0 | 0 | 1 |
| ELP4 | 1.01 | 11 | 11 | 12 | 10 | 7 | 12 | 8 |
| UBAP2 | 1.01 | 9 | 9 | 2 | 4 | 3 | 15 | 15 |
| WDR11 | 1.01 | 20 | 18 | 6 | 15 | 11 | 27 | 28 |
| ACTN1 | 1 | 44 | 23 | 17 | 30 | 36 | 14 | 46 |
| EEF1D | 1 | 10 | 8 | 8 | 11 | 7 | 9 | 10 |
| HIST2H3PS2 | 1 | 1 | 1 | 1 | 1 | 1 | 0 | 1 |
| HSPA8 | 1 | 64 | 52 | 63 | 53 | 55 | 54 | 58 |
| NUP214 | 1 | 23 | 13 | 12 | 16 | 11 | 25 | 23 |
| RPS12 | 1 | 4 | 4 | 4 | 4 | 4 | 3 | 4 |
| SSR4 | 1 | 3 | 1 | 1 | 2 | 2 | 2 | 2 |
| TAB1 | 1 | 3 | 4 | 4 | 4 | 0 | 2 | 3 |
| TUBGCP2 | 1 | 1 | 1 | 1 | 1 | 1 | 1 | 0 |
| UBE2O | 1 | 17 | 6 | 9 | 10 | 9 | 11 | 15 |
| WIPI2 | 1 | 10 | 9 | 6 | 6 | 11 | 13 | 11 |
| C15orf52 | 0.99 | 1 | 1 | 0 | 0 | 0 | 2 | 2 |
| COPS4 | 0.99 | 1 | 1 | 0 | 0 | 0 | 2 | 2 |
| DNMBP | 0.99 | 40 | 37 | 35 | 38 | 44 | 41 | 41 |
| FNBP1L | 0.99 | 1 | 0 | 1 | 0 | 0 | 0 | 0 |
| IMMT | 0.99 | 1 | 0 | 1 | 0 | 0 | 0 | 0 |
| SPG20 | 0.99 | 11 | 10 | 7 | 3 | 12 | 16 | 10 |
| STAM | 0.99 | 3 | 4 | 2 | 0 | 2 | 6 | 5 |
| STRN4 | 0.99 | 1 | 1 | 0 | 0 | 0 | 2 | 2 |
| TXNDC5 | 0.99 | 1 | 0 | 1 | 0 | 0 | 0 | 0 |
| ACTR1B | 0.98 | 1 | 1 | 0 | 0 | 0 | 3 | 1 |
| ATP5A1 | 0.98 | 18 | 15 | 16 | 12 | 20 | 16 | 19 |
| CD59 | 0.98 | 8 | 6 | 7 | 8 | 5 | 0 | 8 |

| | | | | | | | | |
|---|---|---|---|---|---|---|---|---|
| KLC2 | 0.98 | 2 | 1 | 0 | 1 | 2 | 2 | 0 |
| MVP | 0.98 | 3 | 1 | 1 | 1 | 3 | 2 | 0 |
| PDLIM4 | 0.98 | 16 | 10 | 7 | 14 | 12 | 18 | 15 |
| PHB | 0.98 | 5 | 6 | 4 | 8 | 2 | 6 | 5 |
| PSD3 | 0.98 | 12 | 6 | 11 | 6 | 5 | 10 | 3 |
| RHOA | 0.98 | 4 | 3 | 2 | 1 | 0 | 5 | 6 |
| RPS15A | 0.98 | 6 | 6 | 3 | 8 | 7 | 5 | 6 |
| STRAP | 0.98 | 19 | 12 | 16 | 15 | 14 | 13 | 19 |
| TBCB | 0.98 | 1 | 1 | 0 | 0 | 0 | 3 | 1 |
| ELAVL1 | 0.97 | 0 | 1 | 0 | 0 | 0 | 1 | 1 |
| FAM101B | 0.97 | 0 | 1 | 0 | 0 | 0 | 1 | 1 |
| FAM107B | 0.97 | 0 | 1 | 0 | 0 | 0 | 1 | 1 |
| HNRNPR | 0.97 | 0 | 1 | 0 | 0 | 0 | 1 | 1 |
| MAP1A | 0.97 | 4 | 1 | 1 | 1 | 0 | 5 | 2 |
| PPP1R12A | 0.97 | 17 | 12 | 10 | 13 | 4 | 20 | 22 |
| PSMA7 | 0.97 | 1 | 1 | 0 | 0 | 1 | 1 | 2 |
| RAB3B | 0.97 | 4 | 2 | 1 | 2 | 0 | 6 | 3 |
| SRSF6 | 0.97 | 0 | 1 | 0 | 0 | 0 | 1 | 1 |
| UNC45A | 0.97 | 14 | 10 | 11 | 8 | 7 | 16 | 16 |
| UTS2 | 0.97 | 0 | 1 | 0 | 0 | 0 | 1 | 1 |
| YWHAE | 0.97 | 8 | 10 | 8 | 6 | 11 | 11 | 10 |
| SEPT7 | 0.96 | 18 | 17 | 16 | 19 | 20 | 23 | 23 |
| SEPT9 | 0.96 | 23 | 29 | 23 | 26 | 30 | 34 | 35 |
| CTPS1 | 0.96 | 9 | 10 | 9 | 12 | 9 | 7 | 10 |
| DBNL | 0.96 | 9 | 9 | 2 | 7 | 6 | 16 | 12 |
| PSMC2 | 0.96 | 5 | 4 | 2 | 5 | 5 | 6 | 3 |
| RPS7 | 0.96 | 6 | 6 | 6 | 7 | 6 | 4 | 4 |
| SMC1A | 0.96 | 2 | 0 | 1 | 1 | 0 | 0 | 0 |
| TGM2 | 0.96 | 9 | 8 | 8 | 6 | 13 | 7 | 7 |
| TRIP6 | 0.96 | 1 | 1 | 0 | 1 | 0 | 2 | 1 |
| AIMP2 | 0.95 | 2 | 3 | 1 | 2 | 1 | 4 | 4 |
| ATL3 | 0.95 | 5 | 1 | 2 | 2 | 1 | 4 | 3 |
| DCTN3 | 0.95 | 2 | 1 | 0 | 1 | 0 | 1 | 4 |
| LARS | 0.95 | 2 | 1 | 0 | 0 | 0 | 4 | 2 |
| SPTAN1 | 0.95 | 166 | 69 | 101 | 90 | 122 | 18 | 154 |
| TANK | 0.95 | 1 | 1 | 0 | 0 | 1 | 3 | 0 |
| TMF1 | 0.95 | 0 | 1 | 0 | 0 | 0 | 2 | 0 |
| UBE2M | 0.95 | 2 | 2 | 2 | 0 | 2 | 3 | 1 |
| CD44 | 0.94 | 7 | 4 | 6 | 6 | 5 | 1 | 6 |
| CDV3 | 0.94 | 6 | 4 | 1 | 3 | 4 | 7 | 9 |
| COPZ1 | 0.94 | 1 | 1 | 0 | 0 | 2 | 1 | 1 |
| EIF3K | 0.94 | 2 | 1 | 0 | 1 | 0 | 2 | 3 |
| EPRS | 0.94 | 25 | 9 | 7 | 14 | 13 | 18 | 28 |
| IGF2R | 0.94 | 12 | 13 | 4 | 3 | 6 | 26 | 19 |
| RASAL2 | 0.94 | 6 | 0 | 0 | 1 | 0 | 2 | 3 |

| | | | | | | | | |
|---|---|---|---|---|---|---|---|---|
| SERPINB2 | 0.94 | 12 | 11 | 7 | 9 | 12 | 17 | 17 |
| SNRNP70 | 0.94 | 2 | 1 | 0 | 1 | 0 | 2 | 3 |
| TCP1 | 0.94 | 27 | 25 | 22 | 33 | 29 | 20 | 22 |
| TXNDC9 | 0.94 | 2 | 3 | 3 | 3 | 2 | 2 | 2 |
| YES1 | 0.94 | 3 | 0 | 0 | 0 | 0 | 0 | 4 |
| ABCE1 | 0.93 | 4 | 3 | 1 | 3 | 0 | 7 | 4 |
| CAST | 0.93 | 21 | 14 | 12 | 12 | 13 | 28 | 27 |
| DDX39A | 0.93 | 5 | 3 | 1 | 3 | 1 | 6 | 7 |
| EIF3D | 0.93 | 2 | 3 | 2 | 3 | 0 | 4 | 3 |
| HIVEP1 | 0.93 | 2 | 0 | 0 | 0 | 0 | 1 | 2 |
| HNRNPA0 | 0.93 | 2 | 0 | 0 | 0 | 0 | 1 | 2 |
| PPP2R5D;PPP2R5C | 0.93 | 2 | 1 | 0 | 0 | 1 | 3 | 2 |
| SLC25A11 | 0.93 | 5 | 2 | 4 | 3 | 2 | 4 | 3 |
| STIP1 | 0.93 | 2 | 0 | 0 | 0 | 0 | 1 | 2 |
| SVIL | 0.93 | 16 | 7 | 0 | 4 | 4 | 21 | 19 |
| XRCC5 | 0.93 | 3 | 1 | 1 | 3 | 1 | 2 | 2 |
| ZYX | 0.93 | 23 | 20 | 19 | 24 | 27 | 23 | 25 |
| ATP6V1E1 | 0.92 | 0 | 1 | 0 | 1 | 0 | 1 | 0 |
| HIST1H2AD | 0.92 | 2 | 1 | 1 | 2 | 2 | 1 | 0 |
| HSD17B12 | 0.92 | 2 | 1 | 0 | 1 | 2 | 2 | 2 |
| KANK2 | 0.92 | 36 | 27 | 30 | 32 | 34 | 46 | 42 |
| KRT18 | 0.92 | 21 | 9 | 7 | 13 | 14 | 20 | 22 |
| MSTO1 | 0.92 | 0 | 1 | 0 | 0 | 1 | 1 | 0 |
| NUDC | 0.92 | 14 | 13 | 11 | 10 | 8 | 17 | 25 |
| PLEKHA1 | 0.92 | 2 | 1 | 0 | 0 | 2 | 1 | 3 |
| RPS2 | 0.92 | 10 | 9 | 15 | 7 | 7 | 9 | 10 |
| SKA3 | 0.92 | 3 | 2 | 2 | 2 | 3 | 4 | 1 |
| SYNM | 0.92 | 9 | 4 | 4 | 4 | 4 | 13 | 7 |
| TBC1D2B | 0.92 | 1 | 2 | 0 | 0 | 3 | 2 | 1 |
| TP53BP2 | 0.92 | 8 | 3 | 3 | 3 | 1 | 8 | 9 |
| SEPT10 | 0.91 | 9 | 4 | 5 | 5 | 5 | 10 | 9 |
| ABR | 0.91 | 2 | 0 | 0 | 0 | 0 | 2 | 1 |
| ALDOA | 0.91 | 3 | 1 | 0 | 0 | 1 | 2 | 5 |
| ARPC2 | 0.91 | 9 | 5 | 6 | 4 | 5 | 3 | 15 |
| CLIC1 | 0.91 | 3 | 4 | 5 | 2 | 3 | 4 | 4 |
| CNN3 | 0.91 | 3 | 4 | 1 | 1 | 2 | 6 | 7 |
| CSRP1 | 0.91 | 4 | 3 | 1 | 1 | 2 | 5 | 8 |
| DCTN2 | 0.91 | 11 | 5 | 9 | 10 | 5 | 5 | 7 |
| GFPT1 | 0.91 | 5 | 5 | 1 | 3 | 3 | 11 | 7 |
| LARP1 | 0.91 | 24 | 18 | 14 | 16 | 19 | 33 | 34 |
| NCAPH | 0.91 | 7 | 4 | 0 | 1 | 0 | 11 | 11 |
| PLOD1 | 0.91 | 4 | 1 | 0 | 0 | 1 | 6 | 2 |
| PPP1R12C | 0.91 | 2 | 0 | 0 | 0 | 0 | 2 | 1 |
| RNF214 | 0.91 | 2 | 0 | 0 | 0 | 0 | 2 | 1 |
| RPL14 | 0.91 | 1 | 3 | 2 | 1 | 2 | 3 | 2 |

| | | | | | | | | |
|---|---|---|---|---|---|---|---|---|
| ACTR10 | 0.9 | 1 | 0 | 0 | 0 | 0 | 0 | 2 |
| ANXA6 | 0.9 | 1 | 0 | 0 | 0 | 0 | 0 | 2 |
| DPYSL4 | 0.9 | 1 | 0 | 0 | 0 | 0 | 0 | 2 |
| HIST1H1B | 0.9 | 3 | 3 | 3 | 4 | 3 | 4 | 3 |
| HSP90AB2P | 0.9 | 1 | 1 | 0 | 2 | 1 | 1 | 0 |
| MOCS2 | 0.9 | 1 | 0 | 0 | 0 | 0 | 0 | 2 |
| PLS1 | 0.9 | 1 | 0 | 0 | 0 | 0 | 0 | 2 |
| RPS11 | 0.9 | 12 | 7 | 8 | 14 | 7 | 11 | 10 |
| TMOD3 | 0.9 | 28 | 21 | 23 | 28 | 31 | 2 | 23 |
| ADRM1 | 0.89 | 1 | 0 | 0 | 0 | 0 | 1 | 1 |
| AGTPBP1 | 0.89 | 1 | 0 | 0 | 0 | 0 | 1 | 1 |
| AUP1 | 0.89 | 1 | 0 | 0 | 0 | 0 | 1 | 1 |
| CD2AP | 0.89 | 41 | 30 | 44 | 38 | 36 | 36 | 39 |
| CORO1A | 0.89 | 1 | 0 | 0 | 0 | 0 | 1 | 1 |
| EIF2S2 | 0.89 | 1 | 0 | 0 | 0 | 0 | 1 | 1 |
| EIF3I | 0.89 | 6 | 4 | 0 | 5 | 3 | 6 | 10 |
| ELOVL5 | 0.89 | 1 | 0 | 0 | 0 | 0 | 1 | 1 |
| EPS8 | 0.89 | 1 | 0 | 0 | 0 | 0 | 1 | 1 |
| GGA3 | 0.89 | 1 | 0 | 0 | 0 | 0 | 1 | 1 |
| HAUS3 | 0.89 | 0 | 1 | 0 | 2 | 0 | 0 | 0 |
| LARP6 | 0.89 | 1 | 0 | 0 | 0 | 0 | 1 | 1 |
| LIMS1 | 0.89 | 1 | 0 | 0 | 0 | 0 | 1 | 1 |
| MAP4K4 | 0.89 | 1 | 0 | 0 | 0 | 0 | 1 | 1 |
| MAP4K5 | 0.89 | 2 | 2 | 1 | 0 | 2 | 2 | 5 |
| MMP14 | 0.89 | 1 | 0 | 0 | 0 | 0 | 1 | 1 |
| PAK4;PAK7 | 0.89 | 1 | 0 | 0 | 0 | 0 | 1 | 1 |
| PHB2 | 0.89 | 9 | 5 | 7 | 9 | 7 | 8 | 8 |
| RANBP2 | 0.89 | 56 | 44 | 24 | 32 | 28 | 94 | 85 |
| REEP5 | 0.89 | 1 | 0 | 0 | 0 | 0 | 1 | 1 |
| RRAS | 0.89 | 1 | 0 | 0 | 0 | 0 | 1 | 1 |
| SEC61B | 0.89 | 1 | 0 | 0 | 0 | 0 | 1 | 1 |
| SMC3 | 0.89 | 1 | 0 | 0 | 0 | 0 | 1 | 1 |
| SNRPE | 0.89 | 1 | 0 | 0 | 0 | 0 | 1 | 1 |
| SNRPF | 0.89 | 1 | 0 | 0 | 0 | 0 | 1 | 1 |
| SNRPG;SNRPGP15 | 0.89 | 1 | 0 | 0 | 0 | 0 | 1 | 1 |
| SRGAP2;SRGAP1 | 0.89 | 1 | 0 | 0 | 0 | 0 | 1 | 1 |
| TSC1 | 0.89 | 1 | 0 | 0 | 0 | 0 | 1 | 1 |
| TUBB2A | 0.89 | 1 | 0 | 0 | 0 | 0 | 1 | 1 |
| TUBB4B | 0.89 | 42 | 40 | 46 | 46 | 47 | 47 | 47 |
| YIPF5 | 0.89 | 1 | 0 | 0 | 0 | 0 | 1 | 1 |
| C10orf88 | 0.88 | 1 | 0 | 0 | 0 | 0 | 2 | 0 |
| C14orf166 | 0.88 | 3 | 3 | 2 | 4 | 4 | 4 | 4 |
| EPS15L1 | 0.88 | 10 | 10 | 6 | 9 | 8 | 16 | 18 |
| HIST2H2BE;HIST1H2BB;HIST1H2BO | 0.88 | 2 | 2 | 1 | 3 | 3 | 2 | 1 |
| IARS | 0.88 | 8 | 5 | 3 | 7 | 8 | 9 | 10 |

| | | | | | | | | |
|---|---|---|---|---|---|---|---|---|
| NFKBIE | 0.88 | 1 | 0 | 0 | 0 | 0 | 2 | 0 |
| PPP3CA;PPP3CB | 0.88 | 0 | 1 | 1 | 0 | 0 | 1 | 0 |
| RPL22 | 0.88 | 3 | 1 | 0 | 2 | 4 | 1 | 1 |
| SEC13 | 0.88 | 2 | 2 | 0 | 1 | 2 | 3 | 4 |
| CPSF3 | 0.87 | 5 | 2 | 4 | 4 | 3 | 4 | 4 |
| CTTNBP2NL | 0.87 | 2 | 2 | 0 | 0 | 2 | 5 | 2 |
| DFNA5 | 0.87 | 10 | 8 | 10 | 11 | 10 | 12 | 12 |
| EIF3A | 0.87 | 20 | 6 | 12 | 8 | 7 | 16 | 17 |
| HECTD1 | 0.87 | 3 | 1 | 1 | 2 | 1 | 4 | 2 |
| HIST1H1C;HIST1H1D | 0.87 | 13 | 6 | 12 | 8 | 5 | 11 | 12 |
| HN1 | 0.87 | 2 | 3 | 1 | 3 | 3 | 4 | 4 |
| HSP90AA1 | 0.87 | 5 | 1 | 3 | 3 | 2 | 3 | 3 |
| NEK1 | 0.87 | 3 | 1 | 1 | 1 | 1 | 4 | 3 |
| PRDX1 | 0.87 | 10 | 6 | 6 | 6 | 10 | 8 | 15 |
| PRDX6 | 0.87 | 6 | 3 | 2 | 4 | 5 | 2 | 9 |
| PRRC2C | 0.87 | 14 | 13 | 5 | 7 | 7 | 29 | 24 |
| PSMD2 | 0.87 | 10 | 10 | 10 | 10 | 6 | 15 | 16 |
| RARS | 0.87 | 13 | 19 | 15 | 17 | 16 | 25 | 25 |
| SLAIN2 | 0.87 | 4 | 2 | 1 | 2 | 1 | 5 | 6 |
| TXNRD1 | 0.87 | 5 | 2 | 2 | 1 | 1 | 6 | 6 |
| AHNAK | 0.86 | 686 | 559 | 604 | 776 | 792 | 579 | 597 |
| DDT;DDTL | 0.86 | 0 | 1 | 1 | 0 | 1 | 0 | 0 |
| EEF1E1 | 0.86 | 1 | 1 | 1 | 2 | 1 | 0 | 1 |
| GOLGA5 | 0.86 | 9 | 4 | 3 | 5 | 1 | 11 | 11 |
| LYPLA2 | 0.86 | 5 | 4 | 4 | 4 | 5 | 8 | 7 |
| MSH3 | 0.86 | 1 | 0 | 0 | 1 | 0 | 0 | 1 |
| PDLIM5 | 0.86 | 39 | 36 | 36 | 44 | 53 | 41 | 46 |
| PIK3C3 | 0.86 | 3 | 0 | 1 | 0 | 0 | 2 | 1 |
| PIK3R4 | 0.86 | 1 | 0 | 0 | 1 | 0 | 0 | 1 |
| RAF1 | 0.86 | 1 | 0 | 0 | 0 | 1 | 0 | 1 |
| RAI14 | 0.86 | 43 | 26 | 23 | 31 | 28 | 64 | 50 |
| RAPGEF2 | 0.86 | 1 | 0 | 0 | 1 | 0 | 0 | 1 |
| RPL8 | 0.86 | 2 | 0 | 0 | 1 | 1 | 0 | 1 |
| SDC1 | 0.86 | 0 | 1 | 1 | 1 | 0 | 0 | 0 |
| SLC16A1 | 0.86 | 1 | 0 | 0 | 0 | 1 | 0 | 1 |
| UBR4 | 0.86 | 1 | 0 | 0 | 1 | 0 | 0 | 1 |
| VASP | 0.86 | 9 | 6 | 5 | 4 | 2 | 12 | 15 |
| EEF1A1;EEF1A1P5 | 0.85 | 21 | 17 | 23 | 21 | 23 | 17 | 26 |
| EHBP1 | 0.85 | 7 | 6 | 5 | 4 | 5 | 16 | 7 |
| GTF2A2 | 0.85 | 1 | 1 | 0 | 1 | 1 | 2 | 2 |
| KIF13B | 0.85 | 1 | 0 | 0 | 0 | 1 | 1 | 0 |
| NR3C1 | 0.85 | 8 | 3 | 3 | 2 | 1 | 10 | 9 |
| PDCL | 0.85 | 1 | 0 | 0 | 0 | 1 | 1 | 0 |
| PYGB | 0.85 | 8 | 8 | 6 | 10 | 13 | 8 | 10 |
| RPL32 | 0.85 | 1 | 2 | 2 | 2 | 0 | 2 | 1 |

| | | | | | | | | |
|---|---|---|---|---|---|---|---|---|
| SERPINH1 | 0.85 | 9 | 3 | 4 | 6 | 5 | 4 | 12 |
| UBAP2L | 0.85 | 20 | 11 | 10 | 12 | 12 | 25 | 30 |
| ACLY | 0.84 | 10 | 5 | 6 | 6 | 6 | 13 | 12 |
| DLG5 | 0.84 | 22 | 19 | 15 | 17 | 11 | 38 | 36 |
| EPS15 | 0.84 | 1 | 1 | 0 | 1 | 1 | 1 | 3 |
| FAM114A1 | 0.84 | 13 | 9 | 11 | 10 | 12 | 16 | 19 |
| STMN1 | 0.84 | 10 | 5 | 10 | 2 | 7 | 8 | 11 |
| YAP1 | 0.84 | 5 | 6 | 4 | 4 | 7 | 10 | 8 |
| ARHGAP17 | 0.83 | 2 | 1 | 1 | 0 | 1 | 1 | 5 |
| ASAP2 | 0.83 | 2 | 1 | 1 | 0 | 1 | 3 | 3 |
| CACNA2D1 | 0.83 | 12 | 0 | 1 | 1 | 0 | 0 | 9 |
| CAD | 0.83 | 5 | 2 | 3 | 5 | 4 | 2 | 5 |
| GORASP2 | 0.83 | 3 | 3 | 2 | 1 | 4 | 5 | 5 |
| KARS | 0.83 | 6 | 2 | 3 | 5 | 4 | 6 | 3 |
| MRE11A | 0.83 | 22 | 10 | 11 | 16 | 12 | 22 | 30 |
| RANBP1 | 0.83 | 2 | 1 | 1 | 1 | 0 | 3 | 3 |
| RPL23A | 0.83 | 1 | 0 | 1 | 0 | 0 | 0 | 1 |
| SLC25A6 | 0.83 | 17 | 15 | 15 | 24 | 20 | 11 | 14 |
| SSR3 | 0.83 | 1 | 0 | 1 | 0 | 0 | 0 | 1 |
| ASCC3 | 0.82 | 15 | 8 | 12 | 13 | 13 | 15 | 20 |
| CKAP4 | 0.82 | 18 | 14 | 15 | 23 | 13 | 22 | 27 |
| COPS3 | 0.82 | 1 | 1 | 1 | 1 | 1 | 1 | 3 |
| DYNC1LI2 | 0.82 | 4 | 2 | 1 | 2 | 2 | 5 | 7 |
| FXR1 | 0.82 | 2 | 2 | 0 | 0 | 1 | 6 | 3 |
| MYLK | 0.82 | 36 | 15 | 18 | 22 | 24 | 39 | 45 |
| RAN | 0.82 | 13 | 7 | 14 | 9 | 12 | 10 | 11 |
| RPS3 | 0.82 | 15 | 13 | 14 | 21 | 17 | 13 | 18 |
| RUVBL2 | 0.82 | 24 | 16 | 22 | 27 | 23 | 21 | 30 |
| TAGLN | 0.82 | 9 | 7 | 2 | 5 | 8 | 13 | 16 |
| TTN | 0.82 | 1 | 0 | 0 | 2 | 0 | 0 | 0 |
| ARCN1 | 0.81 | 12 | 5 | 4 | 5 | 8 | 15 | 14 |
| ARHGEF28 | 0.81 | 0 | 1 | 0 | 0 | 0 | 2 | 1 |
| ARPIN | 0.81 | 3 | 3 | 3 | 3 | 2 | 5 | 6 |
| CEP97 | 0.81 | 2 | 2 | 0 | 2 | 1 | 4 | 4 |
| HSPB1 | 0.81 | 7 | 3 | 5 | 6 | 7 | 4 | 4 |
| LRBA | 0.81 | 17 | 20 | 17 | 22 | 21 | 36 | 23 |
| NDUFA10 | 0.81 | 1 | 0 | 1 | 0 | 0 | 1 | 0 |
| PAICS | 0.81 | 25 | 19 | 24 | 25 | 33 | 22 | 27 |
| PSMC3 | 0.81 | 3 | 1 | 3 | 0 | 1 | 3 | 2 |
| RHOG | 0.81 | 0 | 1 | 0 | 0 | 0 | 2 | 1 |
| SEC24B | 0.81 | 2 | 2 | 2 | 2 | 3 | 3 | 4 |
| STAT1 | 0.81 | 19 | 16 | 13 | 21 | 27 | 25 | 24 |
| TLN1 | 0.81 | 159 | 137 | 155 | 184 | 212 | 153 | 165 |
| TUBB6 | 0.81 | 10 | 5 | 8 | 11 | 8 | 7 | 9 |
| UFD1L | 0.81 | 4 | 1 | 0 | 0 | 0 | 5 | 6 |

| | | | | | | | | |
|---|---|---|---|---|---|---|---|---|
| CORO1C | 0.8 | 42 | 35 | 44 | 48 | 53 | 41 | 45 |
| DPYSL3 | 0.8 | 25 | 19 | 24 | 25 | 34 | 24 | 25 |
| FHOD1 | 0.8 | 5 | 3 | 6 | 3 | 2 | 5 | 6 |
| IMPDH2 | 0.8 | 3 | 0 | 0 | 0 | 0 | 4 | 1 |
| KIF2C | 0.8 | 3 | 0 | 0 | 0 | 0 | 4 | 1 |
| LIMCH1 | 0.8 | 38 | 26 | 35 | 36 | 40 | 47 | 57 |
| LRCH3 | 0.8 | 2 | 0 | 1 | 1 | 1 | 1 | 0 |
| MSN | 0.8 | 18 | 13 | 14 | 21 | 13 | 24 | 26 |
| RIN1 | 0.8 | 2 | 0 | 1 | 1 | 1 | 0 | 0 |
| UGDH | 0.8 | 8 | 3 | 2 | 6 | 5 | 4 | 12 |
| WDR45 | 0.8 | 1 | 1 | 1 | 1 | 1 | 3 | 1 |
| ABI1 | 0.79 | 4 | 1 | 1 | 1 | 2 | 4 | 5 |
| ANKRD28 | 0.79 | 1 | 0 | 1 | 0 | 1 | 0 | 0 |
| BICD2 | 0.79 | 3 | 2 | 2 | 0 | 0 | 6 | 4 |
| DDX1 | 0.79 | 5 | 5 | 2 | 8 | 5 | 8 | 8 |
| DDX5 | 0.79 | 8 | 6 | 8 | 9 | 6 | 12 | 11 |
| EEF1B2 | 0.79 | 2 | 3 | 0 | 1 | 0 | 6 | 6 |
| EHBP1L1 | 0.79 | 2 | 0 | 0 | 0 | 0 | 3 | 1 |
| MLLT4 | 0.79 | 31 | 27 | 25 | 34 | 26 | 50 | 53 |
| NME1-NME2;NME2 | 0.79 | 7 | 3 | 4 | 6 | 8 | 6 | 7 |
| PSMB3 | 0.79 | 1 | 0 | 1 | 1 | 0 | 0 | 0 |
| RPS4X | 0.79 | 4 | 3 | 4 | 5 | 5 | 1 | 1 |
| TNS3 | 0.79 | 14 | 12 | 11 | 18 | 16 | 22 | 21 |
| C1orf198 | 0.78 | 0 | 1 | 0 | 0 | 1 | 2 | 0 |
| NEK9 | 0.78 | 23 | 18 | 22 | 30 | 22 | 27 | 36 |
| PPP2CA;PPP2CB | 0.78 | 1 | 1 | 2 | 1 | 1 | 1 | 2 |
| RPS3A | 0.78 | 9 | 7 | 9 | 8 | 8 | 13 | 15 |
| UGP2 | 0.78 | 1 | 1 | 2 | 1 | 1 | 1 | 2 |
| CCDC43 | 0.77 | 2 | 0 | 0 | 1 | 0 | 2 | 1 |
| CCT2 | 0.77 | 24 | 16 | 19 | 30 | 29 | 18 | 23 |
| CCT5 | 0.77 | 18 | 14 | 19 | 22 | 22 | 21 | 16 |
| CLTC | 0.77 | 25 | 11 | 16 | 27 | 24 | 13 | 13 |
| CNN2 | 0.77 | 7 | 5 | 1 | 8 | 9 | 10 | 10 |
| FAM160B1 | 0.77 | 3 | 2 | 3 | 3 | 4 | 4 | 2 |
| JUP | 0.77 | 13 | 6 | 9 | 12 | 10 | 9 | 19 |
| NUDCD3 | 0.77 | 1 | 0 | 0 | 0 | 0 | 0 | 3 |
| SPECC1 | 0.77 | 2 | 0 | 0 | 0 | 1 | 2 | 1 |
| AGO2 | 0.76 | 3 | 0 | 1 | 1 | 0 | 1 | 3 |
| ATP6V1G1 | 0.76 | 1 | 0 | 0 | 0 | 0 | 1 | 2 |
| CISD2 | 0.76 | 1 | 1 | 0 | 1 | 0 | 3 | 2 |
| COPE | 0.76 | 1 | 0 | 2 | 0 | 0 | 0 | 0 |
| CTNNA1 | 0.76 | 1 | 0 | 0 | 0 | 0 | 1 | 2 |
| DDX6 | 0.76 | 5 | 2 | 1 | 1 | 2 | 9 | 6 |
| EIF5 | 0.76 | 11 | 15 | 16 | 18 | 16 | 19 | 23 |
| HSPE1-MOB4;MOB4 | 0.76 | 0 | 1 | 1 | 0 | 0 | 1 | 1 |

| | | | | | | | | |
|---|---|---|---|---|---|---|---|---|
| MPDZ | 0.76 | 1 | 0 | 0 | 0 | 0 | 1 | 2 |
| MYH9 | 0.76 | 585 | 459 | 677 | 589 | 777 | 74 | 464 |
| PFN2 | 0.76 | 0 | 1 | 1 | 0 | 0 | 1 | 1 |
| PSMC5 | 0.76 | 2 | 1 | 1 | 1 | 0 | 4 | 3 |
| QARS | 0.76 | 10 | 5 | 8 | 7 | 11 | 12 | 11 |
| SLC2A1 | 0.76 | 4 | 5 | 7 | 4 | 6 | 5 | 7 |
| STX5 | 0.76 | 2 | 1 | 0 | 2 | 0 | 3 | 3 |
| CSNK1A1 | 0.75 | 2 | 0 | 1 | 0 | 0 | 2 | 1 |
| EXOSC8 | 0.75 | 1 | 0 | 0 | 0 | 0 | 2 | 1 |
| GART | 0.75 | 10 | 3 | 9 | 5 | 7 | 8 | 9 |
| IFIT5 | 0.75 | 15 | 10 | 12 | 20 | 19 | 14 | 12 |
| NEDD4L | 0.75 | 1 | 1 | 0 | 2 | 0 | 2 | 2 |
| OPHN1 | 0.75 | 1 | 0 | 0 | 0 | 0 | 2 | 1 |
| RUVBL1 | 0.75 | 21 | 17 | 23 | 26 | 28 | 26 | 31 |
| S100A16 | 0.75 | 1 | 0 | 0 | 0 | 0 | 2 | 1 |
| SAMD9 | 0.75 | 1 | 0 | 0 | 0 | 0 | 2 | 1 |
| SRP54 | 0.75 | 2 | 1 | 1 | 0 | 0 | 5 | 2 |
| STAT3 | 0.75 | 8 | 7 | 8 | 9 | 10 | 11 | 17 |
| STK33 | 0.75 | 1 | 0 | 0 | 0 | 0 | 2 | 1 |
| TPP2 | 0.75 | 2 | 0 | 0 | 2 | 0 | 1 | 1 |
| TRIOBP | 0.75 | 6 | 4 | 3 | 1 | 3 | 14 | 8 |
| VCP | 0.75 | 3 | 1 | 0 | 2 | 1 | 5 | 3 |
| BRE | 0.74 | 2 | 0 | 0 | 1 | 1 | 2 | 1 |
| EIF2A | 0.74 | 9 | 11 | 13 | 11 | 16 | 15 | 14 |
| FGD4 | 0.74 | 6 | 4 | 2 | 5 | 1 | 13 | 8 |
| FKBP15 | 0.74 | 4 | 0 | 1 | 1 | 0 | 4 | 1 |
| KIAA1598 | 0.74 | 3 | 3 | 3 | 1 | 5 | 3 | 7 |
| KPNB1 | 0.74 | 9 | 7 | 11 | 10 | 11 | 13 | 8 |
| LUZP1 | 0.74 | 25 | 16 | 8 | 19 | 11 | 44 | 40 |
| NUP62 | 0.74 | 2 | 1 | 3 | 0 | 1 | 3 | 1 |
| RPL19 | 0.74 | 2 | 1 | 2 | 1 | 0 | 2 | 4 |
| TEX2 | 0.74 | 3 | 1 | 2 | 2 | 1 | 5 | 2 |
| TPD52L2 | 0.74 | 3 | 2 | 3 | 4 | 4 | 2 | 2 |
| TUBA4A | 0.74 | 2 | 0 | 1 | 0 | 1 | 0 | 2 |
| ACTR1A | 0.73 | 7 | 8 | 13 | 7 | 9 | 10 | 12 |
| EIF3B | 0.73 | 7 | 3 | 2 | 8 | 4 | 8 | 8 |
| EIF3C;EIF3CL | 0.73 | 6 | 3 | 5 | 6 | 4 | 9 | 6 |
| EZR | 0.73 | 14 | 9 | 14 | 13 | 15 | 15 | 24 |
| GULP1 | 0.73 | 1 | 0 | 0 | 0 | 1 | 1 | 1 |
| LRRFIP1 | 0.73 | 4 | 4 | 1 | 3 | 4 | 10 | 8 |
| LSM12 | 0.73 | 3 | 0 | 0 | 0 | 0 | 3 | 3 |
| PXN | 0.73 | 15 | 11 | 12 | 20 | 20 | 20 | 22 |
| RANBP3 | 0.73 | 7 | 8 | 8 | 6 | 8 | 17 | 13 |
| RBMX;RBMXL1 | 0.73 | 1 | 0 | 0 | 1 | 0 | 1 | 1 |
| RGPD3 | 0.73 | 0 | 1 | 1 | 1 | 0 | 1 | 1 |

| | | | | | | | | |
|---|---|---|---|---|---|---|---|---|
| RPL29 | 0.73 | 1 | 1 | 2 | 2 | 1 | 1 | 1 |
| TPM1 | 0.73 | 3 | 3 | 6 | 3 | 4 | 1 | 2 |
| WIPI1 | 0.73 | 3 | 3 | 2 | 4 | 6 | 5 | 3 |
| XIAP | 0.73 | 4 | 1 | 3 | 1 | 1 | 5 | 3 |
| AP3D1 | 0.72 | 3 | 4 | 1 | 1 | 0 | 10 | 9 |
| ATP2A2 | 0.72 | 5 | 4 | 3 | 3 | 4 | 9 | 12 |
| CEP170B | 0.72 | 1 | 0 | 0 | 0 | 1 | 2 | 0 |
| HNRNPM | 0.72 | 4 | 4 | 3 | 2 | 2 | 9 | 10 |
| REPS1 | 0.72 | 2 | 2 | 1 | 1 | 2 | 5 | 5 |
| SSFA2 | 0.72 | 5 | 4 | 0 | 1 | 0 | 19 | 5 |
| SWAP70 | 0.72 | 4 | 4 | 5 | 6 | 5 | 8 | 7 |
| TNIP1 | 0.72 | 2 | 2 | 3 | 0 | 3 | 4 | 3 |
| DDX19A;DDX19B | 0.71 | 2 | 0 | 0 | 0 | 0 | 2 | 3 |
| ECHDC1 | 0.71 | 1 | 0 | 0 | 1 | 1 | 0 | 1 |
| EIF4A3 | 0.71 | 1 | 0 | 1 | 0 | 0 | 0 | 2 |
| EXOSC2 | 0.71 | 1 | 1 | 0 | 0 | 0 | 3 | 4 |
| HSPA5 | 0.71 | 41 | 30 | 36 | 53 | 63 | 34 | 43 |
| PCM1 | 0.71 | 3 | 1 | 0 | 0 | 0 | 9 | 2 |
| PCMT1 | 0.71 | 2 | 0 | 0 | 0 | 0 | 2 | 3 |
| RAB34 | 0.71 | 0 | 1 | 0 | 0 | 0 | 2 | 2 |
| RECQL | 0.71 | 1 | 1 | 0 | 0 | 0 | 4 | 3 |
| SEC22B | 0.71 | 2 | 0 | 0 | 0 | 0 | 2 | 3 |
| ACOT9 | 0.7 | 2 | 0 | 0 | 0 | 0 | 3 | 2 |
| CDC27 | 0.7 | 1 | 1 | 0 | 0 | 1 | 3 | 3 |
| CRYBG3 | 0.7 | 1 | 0 | 1 | 0 | 0 | 1 | 1 |
| EEF1G | 0.7 | 13 | 12 | 16 | 17 | 22 | 14 | 17 |
| EXOC3 | 0.7 | 2 | 1 | 2 | 1 | 2 | 4 | 2 |
| FLNA | 0.7 | 542 | 561 | 626 | 852 | 915 | 559 | 480 |
| GSPT1;GSPT2 | 0.7 | 2 | 1 | 0 | 2 | 0 | 3 | 4 |
| HNRNPH1 | 0.7 | 11 | 2 | 5 | 9 | 4 | 10 | 6 |
| HSPH1 | 0.7 | 9 | 6 | 5 | 8 | 9 | 15 | 17 |
| ILF2 | 0.7 | 1 | 0 | 0 | 1 | 1 | 1 | 0 |
| KIF2A | 0.7 | 2 | 0 | 0 | 0 | 0 | 3 | 2 |
| LMNB1 | 0.7 | 1 | 0 | 1 | 0 | 0 | 1 | 1 |
| NCK1 | 0.7 | 3 | 2 | 3 | 2 | 1 | 5 | 6 |
| PTPN14 | 0.7 | 5 | 2 | 1 | 1 | 1 | 10 | 8 |
| RASSF8;C12orf2 | 0.7 | 1 | 1 | 1 | 2 | 1 | 3 | 1 |
| SH3BP5L | 0.7 | 3 | 2 | 1 | 5 | 5 | 1 | 3 |
| SLC25A5 | 0.7 | 7 | 5 | 7 | 10 | 10 | 7 | 5 |
| TKT | 0.7 | 2 | 0 | 0 | 0 | 0 | 3 | 2 |
| UBE2J1 | 0.7 | 1 | 0 | 1 | 0 | 0 | 1 | 1 |
| WDR44 | 0.7 | 3 | 3 | 2 | 3 | 2 | 5 | 10 |
| AP2B1 | 0.69 | 16 | 10 | 17 | 20 | 12 | 19 | 25 |
| AP2S1 | 0.69 | 3 | 0 | 1 | 1 | 1 | 2 | 3 |
| BRAP | 0.69 | 4 | 2 | 1 | 0 | 0 | 10 | 6 |

| | | | | | | | | |
|---|---|---|---|---|---|---|---|---|
| CSE1L | 0.69 | 2 | 0 | 2 | 0 | 1 | 0 | 1 |
| PDZD11 | 0.69 | 2 | 0 | 0 | 0 | 1 | 1 | 3 |
| PIP4K2C | 0.69 | 3 | 2 | 2 | 3 | 3 | 5 | 7 |
| PPP6C | 0.69 | 2 | 3 | 3 | 3 | 2 | 3 | 8 |
| SND1 | 0.69 | 7 | 1 | 3 | 7 | 4 | 4 | 4 |
| TBC1D8 | 0.69 | 2 | 0 | 1 | 1 | 2 | 0 | 0 |
| TWF1 | 0.69 | 10 | 5 | 15 | 8 | 8 | 5 | 8 |
| CNOT1 | 0.68 | 0 | 1 | 0 | 1 | 0 | 2 | 1 |
| DNAJA2 | 0.68 | 0 | 1 | 0 | 1 | 0 | 2 | 1 |
| EIF2S1 | 0.68 | 2 | 0 | 0 | 1 | 0 | 2 | 2 |
| LIMA1 | 0.68 | 35 | 17 | 33 | 32 | 42 | 31 | 45 |
| LPCAT2 | 0.68 | 1 | 0 | 1 | 0 | 1 | 0 | 1 |
| MCM7 | 0.68 | 4 | 2 | 2 | 4 | 4 | 7 | 6 |
| PTRF | 0.68 | 1 | 1 | 1 | 1 | 0 | 2 | 4 |
| TGFB1I1 | 0.68 | 1 | 0 | 1 | 0 | 1 | 0 | 1 |
| TIPRL | 0.68 | 9 | 5 | 10 | 8 | 11 | 9 | 13 |
| TUBB | 0.68 | 8 | 10 | 11 | 16 | 14 | 8 | 10 |
| SEPT8 | 0.67 | 1 | 2 | 1 | 0 | 2 | 4 | 4 |
| ACTBL2 | 0.67 | 0 | 2 | 1 | 1 | 3 | 0 | 0 |
| ARHGEF5 | 0.67 | 1 | 0 | 1 | 1 | 0 | 1 | 1 |
| CASP3 | 0.67 | 2 | 1 | 0 | 1 | 0 | 3 | 6 |
| DRG1 | 0.67 | 4 | 2 | 3 | 1 | 3 | 7 | 7 |
| HIST1H4A;HIST1H... | 0.67 | 10 | 8 | 10 | 17 | 15 | 8 | 9 |
| PPP1R18 | 0.67 | 9 | 6 | 5 | 15 | 7 | 13 | 13 |
| PSMA2 | 0.67 | 1 | 2 | 2 | 3 | 3 | 0 | 1 |
| SMC4 | 0.67 | 2 | 3 | 0 | 5 | 0 | 6 | 4 |
| SNX6 | 0.67 | 2 | 0 | 1 | 0 | 0 | 1 | 3 |
| SRP68 | 0.67 | 19 | 9 | 10 | 21 | 13 | 27 | 26 |
| TARDBP | 0.67 | 2 | 1 | 1 | 3 | 1 | 1 | 5 |
| TUBA1A;TUBA3C | 0.67 | 3 | 4 | 5 | 4 | 8 | 3 | 5 |
| ARHGAP21 | 0.66 | 6 | 2 | 1 | 3 | 2 | 12 | 7 |
| CAP2 | 0.66 | 3 | 0 | 0 | 0 | 0 | 2 | 5 |
| CPNE3;CPNE8;CPNE2 | 0.66 | 1 | 0 | 1 | 1 | 1 | 0 | 1 |
| EEF2 | 0.66 | 20 | 13 | 26 | 21 | 28 | 21 | 19 |
| GCN1L1 | 0.66 | 36 | 9 | 24 | 34 | 26 | 21 | 19 |
| GTF2A1 | 0.66 | 1 | 1 | 2 | 2 | 2 | 2 | 2 |
| KIF13A | 0.66 | 1 | 3 | 3 | 4 | 3 | 2 | 2 |
| MAPKAP1 | 0.66 | 1 | 0 | 1 | 1 | 1 | 0 | 0 |
| MAPRE1 | 0.66 | 2 | 1 | 0 | 0 | 1 | 5 | 4 |
| MICALL1 | 0.66 | 1 | 0 | 1 | 1 | 1 | 0 | 1 |
| NUFIP2 | 0.66 | 9 | 6 | 5 | 7 | 3 | 17 | 20 |
| PEA15 | 0.66 | 1 | 0 | 1 | 1 | 1 | 0 | 0 |
| RAB13 | 0.66 | 2 | 0 | 1 | 1 | 1 | 1 | 3 |
| RAE1 | 0.66 | 0 | 1 | 1 | 0 | 0 | 3 | 0 |
| RER1 | 0.66 | 1 | 0 | 1 | 1 | 1 | 1 | 0 |

| | | | | | | | | |
|---|---|---|---|---|---|---|---|---|
| TDRD3 | 0.66 | 1 | 0 | 0 | 0 | 0 | 1 | 3 |
| TFG | 0.66 | 1 | 0 | 0 | 0 | 0 | 1 | 3 |
| TOR1AIP2 | 0.66 | 1 | 0 | 0 | 0 | 0 | 1 | 3 |
| TPR | 0.66 | 31 | 23 | 14 | 34 | 13 | 68 | 49 |
| ARHGAP35 | 0.65 | 11 | 7 | 9 | 13 | 11 | 17 | 22 |
| ARL6IP5 | 0.65 | 1 | 0 | 0 | 0 | 0 | 3 | 1 |
| ATG2B | 0.65 | 11 | 5 | 13 | 5 | 6 | 14 | 14 |
| ATG3 | 0.65 | 1 | 0 | 0 | 0 | 0 | 2 | 2 |
| ATXN2L | 0.65 | 9 | 14 | 9 | 14 | 14 | 27 | 26 |
| BRCC3 | 0.65 | 1 | 0 | 2 | 0 | 0 | 1 | 0 |
| DNAAF2 | 0.65 | 1 | 0 | 0 | 0 | 0 | 2 | 2 |
| FAM21A;FAM21C | 0.65 | 6 | 3 | 4 | 8 | 7 | 9 | 7 |
| GEN1 | 0.65 | 1 | 0 | 0 | 0 | 0 | 3 | 1 |
| IPO9 | 0.65 | 1 | 0 | 0 | 0 | 0 | 2 | 2 |
| PHGDH | 0.65 | 1 | 2 | 2 | 1 | 4 | 3 | 1 |
| RAB1A | 0.65 | 1 | 0 | 0 | 0 | 0 | 2 | 2 |
| SCYL1 | 0.65 | 1 | 0 | 0 | 0 | 0 | 2 | 2 |
| SLC7A5 | 0.65 | 1 | 0 | 0 | 0 | 0 | 2 | 2 |
| TRIM56 | 0.65 | 1 | 0 | 0 | 0 | 0 | 2 | 2 |
| TSG101 | 0.65 | 1 | 1 | 2 | 1 | 1 | 1 | 4 |
| UFL1 | 0.65 | 1 | 0 | 0 | 0 | 0 | 3 | 1 |
| WDFY1 | 0.65 | 1 | 0 | 0 | 0 | 0 | 3 | 1 |
| YTHDF3 | 0.65 | 1 | 0 | 0 | 0 | 0 | 2 | 2 |
| AFTPH | 0.64 | 1 | 0 | 0 | 0 | 1 | 1 | 2 |
| AKAP11 | 0.64 | 0 | 1 | 1 | 0 | 1 | 2 | 0 |
| CCT6A | 0.64 | 15 | 13 | 19 | 26 | 22 | 17 | 16 |
| CFL1 | 0.64 | 11 | 8 | 11 | 15 | 20 | 6 | 8 |
| DHX29 | 0.64 | 19 | 9 | 13 | 20 | 19 | 29 | 28 |
| ENAH | 0.64 | 1 | 0 | 0 | 0 | 1 | 1 | 2 |
| MAPRE2 | 0.64 | 6 | 4 | 4 | 10 | 9 | 4 | 9 |
| PDLIM1 | 0.64 | 13 | 13 | 17 | 25 | 21 | 21 | 25 |
| PSMD3 | 0.64 | 3 | 2 | 1 | 3 | 4 | 6 | 6 |
| PSMD8 | 0.64 | 4 | 4 | 3 | 5 | 8 | 8 | 9 |
| RPL6 | 0.64 | 4 | 3 | 4 | 8 | 6 | 4 | 4 |
| YWHAZ | 0.64 | 6 | 3 | 6 | 8 | 6 | 9 | 4 |
| ANXA1 | 0.63 | 10 | 9 | 14 | 16 | 17 | 12 | 11 |
| CCDC25 | 0.63 | 1 | 0 | 0 | 1 | 0 | 2 | 1 |
| PEAK1 | 0.63 | 8 | 3 | 5 | 9 | 9 | 10 | 11 |
| PLAA | 0.63 | 1 | 0 | 0 | 0 | 1 | 2 | 1 |
| PSMD12 | 0.63 | 1 | 1 | 1 | 4 | 0 | 1 | 2 |
| WARS | 0.63 | 9 | 4 | 4 | 8 | 14 | 11 | 12 |
| AP2M1 | 0.62 | 6 | 3 | 1 | 5 | 2 | 11 | 12 |
| CCSER2 | 0.62 | 0 | 1 | 0 | 0 | 0 | 3 | 2 |
| CHMP4B | 0.62 | 1 | 1 | 3 | 1 | 1 | 2 | 2 |
| CNOT10 | 0.62 | 0 | 1 | 0 | 0 | 0 | 3 | 2 |

| | | | | | | | | |
|---|---|---|---|---|---|---|---|---|
| COPG2 | 0.62 | 32 | 27 | 46 | 45 | 52 | 48 | 56 |
| DOK1 | 0.62 | 1 | 0 | 0 | 1 | 1 | 1 | 2 |
| GAK | 0.62 | 3 | 1 | 3 | 2 | 3 | 5 | 1 |
| GOLGA2 | 0.62 | 2 | 0 | 0 | 0 | 0 | 4 | 2 |
| ILK | 0.62 | 2 | 0 | 0 | 0 | 1 | 2 | 3 |
| PGAM1 | 0.62 | 4 | 3 | 3 | 3 | 5 | 8 | 10 |
| PKM | 0.62 | 31 | 28 | 50 | 42 | 52 | 32 | 42 |
| RPN2 | 0.62 | 1 | 0 | 0 | 1 | 1 | 0 | 2 |
| RRM2 | 0.62 | 0 | 1 | 2 | 0 | 0 | 2 | 0 |
| S100A11 | 0.62 | 1 | 1 | 2 | 2 | 1 | 3 | 2 |
| SEC23A | 0.62 | 4 | 5 | 5 | 7 | 11 | 5 | 9 |
| SMC2 | 0.62 | 1 | 1 | 2 | 2 | 0 | 3 | 2 |
| SYNJ1 | 0.62 | 2 | 2 | 1 | 1 | 4 | 6 | 4 |
| TAGLN2 | 0.62 | 16 | 10 | 17 | 19 | 28 | 13 | 17 |
| WIBG | 0.62 | 1 | 0 | 0 | 0 | 2 | 1 | 1 |
| ANXA2;ANXA2P2 | 0.61 | 39 | 36 | 57 | 63 | 67 | 36 | 41 |
| ARF4 | 0.61 | 8 | 5 | 11 | 10 | 12 | 6 | 7 |
| BUB1B | 0.61 | 1 | 2 | 2 | 2 | 2 | 6 | 2 |
| CORO1B | 0.61 | 37 | 30 | 55 | 53 | 59 | 31 | 32 |
| HIST3H2A;HIST1H2AB | 0.61 | 5 | 6 | 8 | 11 | 10 | 6 | 6 |
| HNRNPF | 0.61 | 4 | 1 | 1 | 0 | 4 | 3 | 8 |
| KIF14 | 0.61 | 1 | 0 | 0 | 2 | 0 | 2 | 0 |
| LMO7 | 0.61 | 46 | 36 | 51 | 66 | 59 | 97 | 84 |
| MYOF | 0.61 | 24 | 13 | 13 | 27 | 29 | 44 | 36 |
| PTK2 | 0.61 | 4 | 1 | 4 | 4 | 2 | 4 | 5 |
| PTPN12 | 0.61 | 11 | 7 | 13 | 11 | 11 | 20 | 20 |
| RANGAP1 | 0.61 | 14 | 11 | 21 | 20 | 16 | 26 | 24 |
| RPL26;RPL26L1 | 0.61 | 2 | 1 | 1 | 4 | 3 | 1 | 3 |
| SRPR | 0.61 | 2 | 1 | 1 | 0 | 2 | 4 | 5 |
| VIM | 0.61 | 83 | 69 | 111 | 139 | 123 | 85 | 87 |
| ACTN4 | 0.6 | 90 | 55 | 96 | 112 | 150 | 31 | 87 |
| AP2A1 | 0.6 | 10 | 8 | 15 | 19 | 6 | 15 | 13 |
| DCTN4 | 0.6 | 3 | 0 | 2 | 0 | 0 | 3 | 2 |
| PAPSS2 | 0.6 | 1 | 0 | 1 | 1 | 0 | 0 | 2 |
| PRRC2B | 0.6 | 1 | 0 | 1 | 1 | 1 | 1 | 2 |
| SLC16A3 | 0.6 | 1 | 0 | 1 | 1 | 1 | 1 | 2 |
| SNX24 | 0.6 | 0 | 1 | 2 | 1 | 1 | 1 | 1 |
| SNX4 | 0.6 | 1 | 0 | 0 | 1 | 2 | 1 | 0 |
| SNX9 | 0.6 | 5 | 3 | 2 | 6 | 8 | 8 | 10 |
| SRPRB | 0.6 | 0 | 2 | 1 | 1 | 1 | 4 | 2 |
| TRIO | 0.6 | 1 | 0 | 0 | 3 | 0 | 0 | 1 |
| YTHDF2 | 0.6 | 3 | 0 | 0 | 0 | 0 | 4 | 4 |
| COMMD8 | 0.59 | 0 | 1 | 1 | 1 | 1 | 2 | 2 |
| EFTUD1 | 0.59 | 2 | 3 | 3 | 4 | 5 | 7 | 6 |
| ELP5 | 0.59 | 4 | 2 | 6 | 5 | 5 | 4 | 5 |

| | | | | | | | | |
|---|---|---|---|---|---|---|---|---|
| EXOSC4 | 0.59 | 0 | 1 | 1 | 0 | 0 | 2 | 2 |
| NF2 | 0.59 | 1 | 2 | 0 | 1 | 1 | 6 | 5 |
| NUP93 | 0.59 | 1 | 0 | 1 | 1 | 1 | 2 | 1 |
| PLIN3 | 0.59 | 24 | 18 | 31 | 34 | 44 | 26 | 28 |
| PSMD13 | 0.59 | 1 | 1 | 2 | 3 | 2 | 1 | 0 |
| SF3A2 | 0.59 | 3 | 1 | 0 | 2 | 0 | 5 | 7 |
| SNX2 | 0.59 | 1 | 0 | 1 | 0 | 1 | 2 | 0 |
| STRIP1;STRIP2 | 0.59 | 1 | 0 | 1 | 1 | 1 | 2 | 1 |
| ACTB;ACTG1 | 0.58 | 263 | 217 | 369 | 393 | 479 | 37 | 188 |
| ARL1 | 0.58 | 2 | 0 | 1 | 2 | 2 | 1 | 2 |
| COPS2 | 0.58 | 1 | 0 | 0 | 0 | 0 | 2 | 3 |
| DARS | 0.58 | 2 | 1 | 5 | 2 | 2 | 2 | 2 |
| NCKAP1 | 0.58 | 8 | 3 | 9 | 10 | 5 | 10 | 9 |
| NHLRC2 | 0.58 | 2 | 1 | 2 | 5 | 2 | 3 | 3 |
| RPS13 | 0.58 | 2 | 0 | 1 | 2 | 1 | 0 | 3 |
| SH2D4A | 0.58 | 0 | 2 | 2 | 2 | 2 | 1 | 2 |
| ZC3H7A | 0.58 | 3 | 3 | 2 | 4 | 4 | 11 | 7 |
| ATP6V1A | 0.57 | 6 | 1 | 5 | 5 | 6 | 5 | 6 |
| DCP1A | 0.57 | 3 | 0 | 1 | 2 | 1 | 4 | 2 |
| LBR | 0.57 | 5 | 2 | 3 | 6 | 3 | 9 | 8 |
| N4BP1 | 0.57 | 3 | 0 | 1 | 1 | 3 | 3 | 2 |
| PIH1D1 | 0.57 | 1 | 1 | 1 | 2 | 0 | 4 | 3 |
| RASA3 | 0.57 | 1 | 0 | 0 | 1 | 0 | 1 | 3 |
| RPLP2 | 0.57 | 1 | 2 | 5 | 2 | 2 | 3 | 2 |
| RPS20 | 0.57 | 3 | 2 | 4 | 5 | 6 | 2 | 4 |
| UTRN | 0.57 | 11 | 9 | 13 | 13 | 16 | 32 | 19 |
| AHSA1 | 0.56 | 3 | 2 | 4 | 6 | 2 | 4 | 7 |
| ARFGAP2 | 0.56 | 4 | 0 | 3 | 0 | 0 | 2 | 4 |
| ATP2B1 | 0.56 | 1 | 0 | 0 | 1 | 1 | 2 | 2 |
| ATP6V1B2 | 0.56 | 2 | 0 | 1 | 3 | 0 | 1 | 2 |
| BIN1 | 0.56 | 3 | 1 | 3 | 1 | 2 | 5 | 6 |
| COPB2 | 0.56 | 6 | 4 | 13 | 8 | 6 | 8 | 7 |
| IGBP1 | 0.56 | 6 | 1 | 4 | 3 | 9 | 4 | 4 |
| MAP7D1 | 0.56 | 5 | 0 | 1 | 0 | 2 | 6 | 3 |
| MGEA5 | 0.56 | 5 | 3 | 6 | 9 | 8 | 4 | 8 |
| MYL6 | 0.56 | 7 | 13 | 20 | 15 | 18 | 5 | 10 |
| PPP1CC | 0.56 | 1 | 2 | 3 | 4 | 3 | 2 | 3 |
| RRBP1 | 0.56 | 13 | 6 | 1 | 2 | 1 | 35 | 27 |
| SNCA | 0.56 | 4 | 2 | 7 | 5 | 5 | 3 | 6 |
| SYNRG | 0.56 | 1 | 1 | 0 | 0 | 0 | 4 | 6 |
| TPM4 | 0.56 | 4 | 1 | 5 | 0 | 2 | 0 | 8 |
| ZCCHC6 | 0.56 | 1 | 0 | 0 | 1 | 0 | 2 | 2 |
| ZW10 | 0.56 | 1 | 0 | 1 | 2 | 1 | 1 | 1 |
| ATP5B | 0.55 | 4 | 0 | 0 | 0 | 2 | 4 | 4 |
| EIF5A;EIF5A2 | 0.55 | 3 | 0 | 1 | 2 | 3 | 2 | 3 |

| | | | | | | | | |
|---|---|---|---|---|---|---|---|---|
| GBF1 | 0.55 | 0 | 2 | 1 | 3 | 1 | 2 | 3 |
| PPP1CB | 0.55 | 4 | 4 | 8 | 9 | 7 | 4 | 6 |
| RABGAP1L | 0.55 | 1 | 0 | 1 | 0 | 0 | 2 | 2 |
| SEC31A | 0.55 | 5 | 1 | 3 | 4 | 6 | 3 | 8 |
| SRP9 | 0.55 | 1 | 0 | 1 | 1 | 0 | 2 | 2 |
| SYAP1 | 0.55 | 1 | 2 | 3 | 3 | 3 | 5 | 3 |
| TCHP | 0.55 | 0 | 1 | 0 | 0 | 0 | 5 | 1 |
| AP3M1 | 0.54 | 0 | 1 | 1 | 2 | 2 | 0 | 0 |
| AVEN | 0.54 | 3 | 0 | 0 | 1 | 0 | 3 | 5 |
| CYFIP1 | 0.54 | 10 | 5 | 12 | 15 | 15 | 12 | 16 |
| DHPS | 0.54 | 1 | 0 | 1 | 1 | 0 | 0 | 3 |
| DST;MACF1 | 0.54 | 5 | 0 | 2 | 2 | 2 | 5 | 4 |
| EXOSC9 | 0.54 | 2 | 0 | 1 | 1 | 1 | 3 | 3 |
| KNTC1 | 0.54 | 2 | 1 | 2 | 1 | 1 | 8 | 2 |
| MAGED2 | 0.54 | 2 | 0 | 1 | 1 | 1 | 4 | 2 |
| MYO9B | 0.54 | 1 | 0 | 0 | 1 | 1 | 3 | 1 |
| NUMB | 0.54 | 5 | 3 | 2 | 4 | 6 | 14 | 10 |
| PLEKHA2 | 0.54 | 4 | 0 | 3 | 2 | 1 | 1 | 4 |
| SLC25A10 | 0.54 | 1 | 0 | 0 | 1 | 1 | 3 | 1 |
| TACC1 | 0.54 | 4 | 4 | 6 | 5 | 4 | 15 | 8 |
| TPM3 | 0.54 | 8 | 5 | 18 | 7 | 11 | 1 | 7 |
| TRIP12 | 0.54 | 2 | 0 | 1 | 0 | 0 | 5 | 1 |
| TSN | 0.54 | 1 | 1 | 1 | 3 | 3 | 2 | 3 |
| TWF2 | 0.54 | 10 | 6 | 15 | 18 | 12 | 9 | 13 |
| ZC3HAV1 | 0.54 | 9 | 3 | 6 | 8 | 10 | 16 | 13 |
| AKAP12 | 0.53 | 14 | 6 | 10 | 8 | 4 | 32 | 25 |
| ATXN2 | 0.53 | 1 | 3 | 1 | 2 | 4 | 6 | 6 |
| EIF4E | 0.53 | 2 | 5 | 6 | 7 | 8 | 7 | 4 |
| IRAK1 | 0.53 | 3 | 1 | 2 | 7 | 3 | 1 | 4 |
| TTC28 | 0.53 | 1 | 0 | 0 | 1 | 2 | 2 | 0 |
| VARS | 0.53 | 3 | 2 | 4 | 5 | 5 | 7 | 8 |
| ANK2 | 0.52 | 1 | 0 | 0 | 1 | 3 | 1 | 1 |
| DENND4C | 0.52 | 6 | 3 | 8 | 8 | 7 | 12 | 12 |
| DPYSL2 | 0.52 | 42 | 42 | 75 | 78 | 91 | 42 | 42 |
| EPB41L2 | 0.52 | 18 | 12 | 19 | 29 | 27 | 40 | 38 |
| HNRNPK | 0.52 | 17 | 7 | 16 | 25 | 26 | 22 | 19 |
| PPP6R3 | 0.52 | 2 | 1 | 3 | 3 | 3 | 3 | 6 |
| PRKAA1 | 0.52 | 1 | 0 | 0 | 0 | 0 | 3 | 3 |
| RPN1 | 0.52 | 2 | 0 | 1 | 1 | 0 | 5 | 1 |
| ATP1A1;ATP1A3 | 0.51 | 2 | 1 | 3 | 3 | 2 | 6 | 2 |
| EPHA2 | 0.51 | 8 | 5 | 8 | 13 | 13 | 16 | 20 |
| MTAP | 0.51 | 1 | 0 | 1 | 2 | 1 | 2 | 1 |
| NUMBL | 0.51 | 2 | 1 | 3 | 4 | 4 | 3 | 4 |
| RIPK2 | 0.51 | 0 | 2 | 2 | 2 | 2 | 4 | 2 |
| SPATS2L | 0.51 | 1 | 0 | 0 | 0 | 0 | 5 | 1 |

| | | | | | | | | |
|---|---|---|---|---|---|---|---|---|
| STEAP3 | 0.51 | 2 | 0 | 0 | 2 | 3 | 2 | 2 |
| TACC2 | 0.51 | 0 | 2 | 1 | 1 | 2 | 5 | 2 |
| TUBA1B | 0.51 | 36 | 39 | 68 | 76 | 78 | 49 | 53 |
| YBX1 | 0.51 | 4 | 1 | 2 | 6 | 2 | 7 | 5 |
| ZDHHC5 | 0.51 | 1 | 0 | 0 | 0 | 1 | 2 | 3 |
| ARHGAP18 | 0.5 | 8 | 7 | 15 | 15 | 18 | 15 | 13 |
| FASN | 0.5 | 157 | 148 | 300 | 307 | 314 | 170 | 177 |
| LDHA | 0.5 | 4 | 4 | 8 | 8 | 11 | 3 | 3 |
| NCAPG | 0.5 | 2 | 2 | 3 | 1 | 2 | 8 | 7 |
| PHACTR4 | 0.5 | 1 | 2 | 0 | 1 | 0 | 7 | 7 |
| SCFD1 | 0.5 | 2 | 1 | 0 | 3 | 0 | 7 | 4 |
| SMAP1 | 0.5 | 1 | 2 | 2 | 6 | 4 | 2 | 1 |
| TACC3 | 0.5 | 1 | 1 | 4 | 1 | 2 | 3 | 1 |
| TOP2A | 0.5 | 1 | 0 | 0 | 0 | 0 | 6 | 0 |
| ABCF3 | 0.49 | 1 | 0 | 1 | 1 | 0 | 3 | 2 |
| CALM2;CALM1;CALM3 | 0.49 | 5 | 5 | 13 | 9 | 11 | 2 | 4 |
| DHCR7 | 0.49 | 1 | 0 | 0 | 2 | 0 | 2 | 2 |
| EEF2K | 0.49 | 1 | 1 | 2 | 2 | 2 | 3 | 6 |
| HIST1H1E | 0.49 | 1 | 1 | 2 | 4 | 3 | 2 | 1 |
| PAK2 | 0.49 | 5 | 3 | 4 | 7 | 9 | 10 | 14 |
| RIC8A | 0.49 | 6 | 0 | 1 | 3 | 5 | 5 | 2 |
| RTN4 | 0.49 | 8 | 6 | 14 | 16 | 15 | 15 | 14 |
| SERBP1 | 0.49 | 10 | 8 | 15 | 17 | 16 | 25 | 29 |
| SNRPD3 | 0.49 | 1 | 0 | 1 | 1 | 0 | 0 | 4 |
| DNM1L | 0.48 | 7 | 2 | 8 | 9 | 10 | 2 | 6 |
| ETF1 | 0.48 | 8 | 0 | 3 | 3 | 5 | 4 | 8 |
| FAM120A | 0.48 | 3 | 1 | 0 | 0 | 2 | 8 | 8 |
| G3BP1 | 0.48 | 3 | 2 | 2 | 4 | 3 | 7 | 12 |
| IQGAP1 | 0.48 | 31 | 13 | 29 | 46 | 57 | 6 | 28 |
| MAVS | 0.48 | 3 | 3 | 5 | 7 | 10 | 6 | 6 |
| PDZD8 | 0.48 | 0 | 1 | 1 | 1 | 3 | 2 | 2 |
| RAB3GAP1 | 0.48 | 3 | 1 | 4 | 5 | 5 | 4 | 2 |
| SUGT1 | 0.48 | 3 | 1 | 5 | 4 | 5 | 4 | 5 |
| ZFYVE16 | 0.48 | 0 | 1 | 1 | 0 | 1 | 4 | 2 |
| DSP | 0.47 | 1 | 0 | 2 | 1 | 0 | 2 | 2 |
| EXOSC7 | 0.47 | 1 | 0 | 0 | 0 | 0 | 2 | 5 |
| NAP1L1 | 0.47 | 7 | 6 | 9 | 17 | 19 | 11 | 13 |
| PSMD4 | 0.47 | 2 | 1 | 2 | 2 | 1 | 7 | 6 |
| SEC24A | 0.47 | 2 | 0 | 2 | 2 | 3 | 1 | 0 |
| SF1 | 0.47 | 3 | 3 | 3 | 2 | 5 | 10 | 13 |
| SYNE1 | 0.47 | 1 | 0 | 4 | 0 | 0 | 0 | 1 |
| CDC42 | 0.46 | 3 | 0 | 2 | 5 | 2 | 1 | 2 |
| CIAPIN1 | 0.46 | 3 | 3 | 7 | 8 | 5 | 8 | 10 |
| CSTF2T | 0.46 | 1 | 0 | 0 | 1 | 0 | 3 | 3 |
| IGF2BP2 | 0.46 | 1 | 0 | 0 | 0 | 0 | 4 | 3 |

| | | | | | | | | |
|---|---|---|---|---|---|---|---|---|
| LPXN | 0.46 | 2 | 0 | 0 | 2 | 1 | 4 | 3 |
| ACSL3 | 0.45 | 1 | 0 | 1 | 1 | 0 | 3 | 3 |
| CCT7 | 0.45 | 15 | 8 | 25 | 28 | 24 | 14 | 13 |
| DICER1 | 0.45 | 1 | 0 | 0 | 0 | 2 | 2 | 3 |
| EIF3J | 0.45 | 0 | 1 | 1 | 1 | 1 | 3 | 4 |
| MYO1E | 0.45 | 1 | 1 | 0 | 1 | 0 | 5 | 7 |
| FAM47E-STBD1 | 0.44 | 2 | 0 | 0 | 0 | 0 | 6 | 4 |
| NUP98 | 0.44 | 2 | 0 | 0 | 0 | 0 | 5 | 5 |
| PLS3 | 0.44 | 8 | 2 | 8 | 11 | 13 | 9 | 10 |
| PMS1 | 0.44 | 0 | 1 | 1 | 1 | 0 | 4 | 3 |
| RAB3GAP2 | 0.44 | 3 | 1 | 3 | 6 | 6 | 3 | 6 |
| SF3B6 | 0.44 | 1 | 1 | 0 | 2 | 1 | 6 | 5 |
| CXorf56 | 0.43 | 0 | 1 | 0 | 0 | 0 | 4 | 5 |
| DDX21 | 0.43 | 4 | 1 | 2 | 3 | 2 | 13 | 7 |
| DIAPH1 | 0.43 | 2 | 0 | 4 | 1 | 1 | 3 | 1 |
| ESYT2 | 0.43 | 2 | 2 | 1 | 5 | 4 | 9 | 8 |
| HNRNPU | 0.43 | 9 | 3 | 9 | 11 | 6 | 20 | 19 |
| KMT2A | 0.43 | 0 | 1 | 2 | 1 | 0 | 3 | 3 |
| MYL12B;MYL12A | 0.43 | 10 | 12 | 42 | 21 | 15 | 4 | 14 |
| SYNCRIP | 0.43 | 2 | 1 | 4 | 7 | 1 | 1 | 3 |
| TMEM165 | 0.43 | 1 | 0 | 2 | 2 | 2 | 1 | 0 |
| KIAA0196 | 0.42 | 2 | 0 | 3 | 2 | 3 | 1 | 1 |
| NOTCH2 | 0.42 | 1 | 0 | 1 | 2 | 2 | 3 | 2 |
| PSMC6 | 0.42 | 2 | 0 | 2 | 3 | 0 | 4 | 3 |
| PSMD1 | 0.42 | 1 | 1 | 2 | 4 | 2 | 6 | 3 |
| RPAP3 | 0.42 | 4 | 1 | 1 | 2 | 2 | 7 | 15 |
| ADD1 | 0.41 | 5 | 2 | 8 | 6 | 10 | 11 | 10 |
| ANKS1A | 0.41 | 1 | 0 | 1 | 2 | 2 | 3 | 3 |
| BUB3 | 0.41 | 0 | 1 | 1 | 2 | 1 | 4 | 3 |
| EGLN1 | 0.41 | 1 | 0 | 2 | 1 | 3 | 1 | 2 |
| FERMT2 | 0.41 | 2 | 1 | 2 | 3 | 2 | 10 | 5 |
| G3BP2 | 0.41 | 1 | 0 | 0 | 2 | 1 | 4 | 2 |
| NCOA7 | 0.41 | 2 | 0 | 0 | 0 | 0 | 5 | 6 |
| PPME1 | 0.41 | 1 | 0 | 1 | 1 | 1 | 3 | 4 |
| PRKDC | 0.41 | 16 | 4 | 16 | 30 | 18 | 16 | 22 |
| STK24 | 0.41 | 3 | 0 | 1 | 1 | 5 | 4 | 3 |
| EIF2S3;EIF2S3L | 0.4 | 1 | 1 | 3 | 3 | 4 | 2 | 6 |
| FAM114A2 | 0.4 | 3 | 0 | 1 | 3 | 2 | 2 | 8 |
| USP10 | 0.4 | 2 | 0 | 1 | 0 | 0 | 4 | 6 |
| VAPA | 0.4 | 3 | 2 | 7 | 4 | 5 | 7 | 12 |
| VPS33B | 0.4 | 1 | 0 | 1 | 3 | 3 | 1 | 0 |
| ANKLE2 | 0.39 | 1 | 0 | 0 | 0 | 0 | 5 | 4 |
| EHD1 | 0.39 | 1 | 0 | 2 | 3 | 2 | 2 | 1 |
| FAM129B | 0.39 | 6 | 6 | 13 | 16 | 22 | 17 | 20 |
| HACD3 | 0.39 | 1 | 0 | 2 | 4 | 1 | 1 | 1 |

| | | | | | | | |
|---|---|---|---|---|---|---|---|
| NONO | 0.39 | 9 | 5 | 6 | 7 | 9 | 33 | 30 |
| PNN | 0.39 | 4 | 3 | 6 | 4 | 2 | 14 | 18 |
| TCEB1 | 0.39 | 0 | 1 | 1 | 2 | 2 | 5 | 1 |
| WASF2 | 0.39 | 3 | 1 | 5 | 7 | 6 | 3 | 1 |
| AKAP2;PALM2-AKA... | 0.38 | 5 | 2 | 4 | 7 | 3 | 16 | 14 |
| PA2G4 | 0.38 | 1 | 0 | 0 | 0 | 1 | 5 | 3 |
| PLAT | 0.38 | 1 | 0 | 1 | 2 | 1 | 4 | 3 |
| VAMP3 | 0.38 | 0 | 1 | 1 | 2 | 1 | 3 | 5 |
| CHMP5 | 0.37 | 0 | 2 | 3 | 3 | 5 | 4 | 1 |
| HCFC1 | 0.37 | 6 | 6 | 12 | 11 | 10 | 25 | 27 |
| NAP1L4 | 0.37 | 3 | 2 | 7 | 7 | 11 | 6 | 5 |
| STAT2 | 0.37 | 1 | 3 | 5 | 8 | 7 | 6 | 3 |
| TOX4 | 0.37 | 4 | 4 | 9 | 16 | 11 | 14 | 14 |
| ALDH3A2 | 0.36 | 1 | 2 | 3 | 4 | 4 | 10 | 7 |
| EDC4 | 0.36 | 1 | 1 | 5 | 2 | 3 | 6 | 3 |
| HSPA4 | 0.36 | 1 | 1 | 3 | 4 | 5 | 5 | 6 |
| NEXN | 0.36 | 7 | 2 | 2 | 6 | 17 | 11 | 16 |
| TBPL1 | 0.36 | 2 | 1 | 4 | 7 | 5 | 7 | 5 |
| YEATS2 | 0.36 | 1 | 0 | 0 | 0 | 0 | 7 | 3 |
| YWHAB | 0.36 | 1 | 0 | 3 | 0 | 2 | 3 | 1 |
| ARHGEF10 | 0.35 | 3 | 1 | 7 | 9 | 4 | 4 | 5 |
| DCP1B | 0.35 | 1 | 0 | 2 | 3 | 3 | 0 | 1 |
| DNMT1 | 0.35 | 1 | 0 | 3 | 1 | 1 | 3 | 3 |
| INF2 | 0.35 | 1 | 1 | 0 | 1 | 2 | 12 | 3 |
| NRBP1 | 0.35 | 1 | 0 | 2 | 3 | 3 | 1 | 1 |
| SLC1A5 | 0.35 | 1 | 0 | 3 | 2 | 2 | 3 | 3 |
| STIM1 | 0.35 | 1 | 0 | 1 | 1 | 1 | 4 | 5 |
| ANLN | 0.34 | 3 | 2 | 5 | 6 | 3 | 16 | 12 |
| CAPRIN1 | 0.34 | 1 | 0 | 2 | 0 | 1 | 2 | 6 |
| JMJD1C | 0.34 | 5 | 4 | 7 | 10 | 10 | 28 | 18 |
| NARS | 0.34 | 1 | 0 | 1 | 3 | 1 | 4 | 3 |
| NSF | 0.34 | 2 | 0 | 2 | 5 | 3 | 4 | 4 |
| RTCB | 0.34 | 4 | 0 | 3 | 3 | 3 | 8 | 7 |
| SNRNP200 | 0.34 | 2 | 0 | 2 | 6 | 3 | 2 | 3 |
| TECR | 0.34 | 1 | 0 | 5 | 1 | 2 | 1 | 1 |
| ESYT1 | 0.33 | 15 | 6 | 27 | 38 | 29 | 27 | 25 |
| MARCKS | 0.33 | 3 | 0 | 0 | 2 | 1 | 7 | 8 |
| NT5C2 | 0.33 | 2 | 0 | 4 | 3 | 4 | 2 | 1 |
| PACS1 | 0.33 | 1 | 0 | 1 | 3 | 4 | 2 | 3 |
| PGRMC2 | 0.33 | 1 | 1 | 2 | 5 | 4 | 5 | 8 |
| PPP1CA;PPP1CB;P... | 0.33 | 13 | 10 | 35 | 38 | 38 | 22 | 21 |
| RBBP7 | 0.33 | 2 | 2 | 5 | 9 | 8 | 9 | 11 |
| APBB2 | 0.32 | 1 | 0 | 2 | 4 | 3 | 1 | 1 |
| HBS1L | 0.32 | 3 | 2 | 5 | 8 | 14 | 6 | 11 |
| PLA2G4A | 0.32 | 4 | 1 | 6 | 9 | 9 | 9 | 12 |

| | | | | | | | | |
|---|---|---|---|---|---|---|---|---|
| PRRC2A | 0.32 | 2 | 0 | 0 | 2 | 2 | 6 | 7 |
| SF3A3 | 0.32 | 1 | 0 | 1 | 1 | 1 | 6 | 4 |
| SNRPB2 | 0.32 | 1 | 1 | 2 | 2 | 1 | 8 | 9 |
| ZC3H15 | 0.32 | 1 | 0 | 1 | 2 | 2 | 2 | 7 |
| KPNA2 | 0.31 | 1 | 0 | 1 | 6 | 1 | 3 | 1 |
| NBEA | 0.31 | 0 | 1 | 1 | 2 | 8 | 0 | 0 |
| SF3B4 | 0.31 | 1 | 0 | 1 | 0 | 2 | 6 | 4 |
| SAP30BP | 0.3 | 1 | 2 | 7 | 7 | 7 | 7 | 8 |
| EIF5B | 0.29 | 6 | 0 | 4 | 4 | 3 | 12 | 10 |
| FARSB | 0.29 | 0 | 2 | 5 | 5 | 5 | 2 | 6 |
| HDAC2 | 0.29 | 2 | 0 | 2 | 1 | 0 | 7 | 7 |
| MCM4 | 0.29 | 1 | 0 | 0 | 1 | 1 | 5 | 7 |
| MYH14 | 0.29 | 3 | 2 | 14 | 8 | 9 | 1 | 2 |
| PEX14 | 0.29 | 0 | 1 | 3 | 4 | 4 | 5 | 0 |
| TMPO | 0.29 | 2 | 2 | 8 | 6 | 5 | 13 | 11 |
| ZC3H14 | 0.29 | 1 | 0 | 0 | 0 | 0 | 9 | 4 |
| OSBPL8 | 0.28 | 2 | 0 | 2 | 6 | 5 | 4 | 0 |
| PRMT7 | 0.28 | 2 | 0 | 1 | 3 | 1 | 4 | 10 |
| TALDO1 | 0.28 | 1 | 0 | 3 | 1 | 4 | 3 | 5 |
| ARHGAP1 | 0.27 | 1 | 0 | 0 | 2 | 3 | 4 | 7 |
| CACYBP | 0.27 | 2 | 0 | 4 | 6 | 4 | 2 | 5 |
| CAND1 | 0.27 | 1 | 0 | 4 | 3 | 4 | 1 | 1 |
| CTNND1 | 0.27 | 4 | 2 | 8 | 15 | 12 | 16 | 12 |
| PPP2R1A | 0.27 | 1 | 1 | 7 | 9 | 2 | 2 | 1 |
| PRMT1 | 0.27 | 1 | 0 | 3 | 4 | 2 | 5 | 4 |
| CUL4B | 0.26 | 5 | 3 | 17 | 12 | 12 | 19 | 25 |
| EGFR | 0.26 | 4 | 1 | 6 | 9 | 7 | 17 | 14 |
| KPNA3 | 0.26 | 1 | 0 | 2 | 4 | 6 | 1 | 3 |
| MARS | 0.26 | 1 | 0 | 2 | 4 | 4 | 5 | 3 |
| RDX | 0.26 | 2 | 0 | 5 | 2 | 2 | 3 | 9 |
| SYNPO | 0.26 | 1 | 0 | 2 | 3 | 5 | 4 | 5 |
| CPSF7 | 0.25 | 1 | 0 | 3 | 2 | 4 | 6 | 5 |
| ECD | 0.25 | 4 | 1 | 10 | 10 | 13 | 7 | 6 |
| EPB41 | 0.25 | 0 | 1 | 1 | 3 | 2 | 7 | 7 |
| BASP1 | 0.24 | 1 | 0 | 1 | 3 | 2 | 6 | 7 |
| COPA | 0.24 | 6 | 2 | 23 | 18 | 10 | 4 | 7 |
| EPB41L1 | 0.24 | 1 | 0 | 2 | 1 | 4 | 3 | 9 |
| MSANTD2 | 0.24 | 3 | 0 | 4 | 2 | 1 | 10 | 9 |
| RBM25 | 0.24 | 1 | 0 | 2 | 3 | 0 | 6 | 7 |
| C1QBP | 0.23 | 1 | 0 | 4 | 1 | 2 | 7 | 5 |
| EIF3L | 0.23 | 2 | 0 | 7 | 6 | 2 | 4 | 3 |
| ITGA3 | 0.23 | 1 | 0 | 1 | 4 | 3 | 9 | 4 |
| NUDT5 | 0.23 | 0 | 1 | 1 | 7 | 5 | 5 | 2 |
| POP1 | 0.23 | 0 | 1 | 4 | 2 | 3 | 7 | 7 |
| PPP1R9B | 0.23 | 5 | 0 | 7 | 11 | 5 | 6 | 9 |

| | | | | | | | | |
|---|---|---|---|---|---|---|---|---|
| PRPF40A | 0.23 | 1 | 0 | 2 | 0 | 0 | 8 | 7 |
| G6PD | 0.22 | 5 | 0 | 7 | 11 | 9 | 8 | 7 |
| GPRIN1 | 0.22 | 2 | 0 | 0 | 1 | 5 | 10 | 7 |
| PRPF8 | 0.22 | 2 | 0 | 2 | 3 | 2 | 16 | 2 |
| XRN2 | 0.22 | 1 | 0 | 2 | 3 | 5 | 9 | 3 |
| DLG1 | 0.21 | 3 | 1 | 11 | 9 | 13 | 13 | 11 |
| NACC1 | 0.21 | 1 | 0 | 0 | 2 | 1 | 9 | 8 |
| NPM1 | 0.21 | 3 | 1 | 9 | 13 | 9 | 15 | 15 |
| PTBP1 | 0.21 | 1 | 0 | 5 | 5 | 3 | 3 | 6 |
| PTPN11 | 0.2 | 0 | 1 | 2 | 5 | 6 | 9 | 5 |
| KIAA0368;ECM29 | 0.19 | 4 | 1 | 10 | 19 | 15 | 1 | 0 |
| KIF4A | 0.19 | 1 | 0 | 1 | 1 | 1 | 12 | 8 |
| WDR1 | 0.19 | 3 | 1 | 11 | 7 | 23 | 0 | 2 |
| CFL2 | 0.18 | 1 | 0 | 7 | 5 | 6 | 0 | 0 |
| PCNP | 0.18 | 2 | 0 | 5 | 4 | 6 | 12 | 11 |
| SF3B2 | 0.18 | 3 | 2 | 6 | 10 | 9 | 29 | 30 |
| USP7 | 0.18 | 1 | 0 | 3 | 5 | 4 | 9 | 9 |
| AFAP1 | 0.17 | 1 | 0 | 4 | 5 | 8 | 9 | 7 |
| C19orf43 | 0.17 | 1 | 0 | 5 | 3 | 5 | 8 | 10 |
| CDK11B;CDC2L1;CDK11A | 0.17 | 1 | 0 | 5 | 4 | 3 | 8 | 10 |
| HNRNPA1;HNRNPA1L2 | 0.17 | 2 | 0 | 0 | 6 | 1 | 13 | 11 |
| EIF3E | 0.16 | 3 | 0 | 10 | 8 | 12 | 2 | 2 |
| ITGB1 | 0.16 | 4 | 0 | 4 | 5 | 13 | 15 | 13 |
| KHSRP | 0.16 | 1 | 0 | 2 | 3 | 5 | 10 | 12 |
| PSPC1 | 0.16 | 1 | 0 | 8 | 5 | 7 | 5 | 6 |
| PUF60 | 0.16 | 1 | 1 | 6 | 9 | 4 | 17 | 15 |
| SFPQ | 0.16 | 3 | 2 | 14 | 20 | 15 | 29 | 27 |
| SLC3A2 | 0.16 | 2 | 0 | 5 | 9 | 11 | 9 | 6 |
| COPG1 | 0.15 | 2 | 0 | 9 | 8 | 10 | 5 | 6 |
| IPO7 | 0.15 | 0 | 1 | 8 | 5 | 11 | 4 | 5 |
| RB1 | 0.15 | 1 | 0 | 2 | 2 | 0 | 11 | 15 |
| TP53BP1 | 0.14 | 2 | 0 | 2 | 6 | 4 | 19 | 12 |
| BRD4 | 0.13 | 1 | 0 | 2 | 4 | 4 | 14 | 15 |
| DHX9 | 0.13 | 2 | 0 | 2 | 8 | 6 | 15 | 16 |
| SF3A1 | 0.12 | 0 | 1 | 1 | 8 | 5 | 17 | 14 |
| TRIM28 | 0.12 | 1 | 0 | 1 | 7 | 5 | 14 | 14 |
| FAM208A | 0.11 | 1 | 0 | 3 | 7 | 2 | 16 | 15 |
| SIN3B | 0.1 | 1 | 0 | 6 | 11 | 9 | 15 | 16 |
| NUMA1 | 0.05 | 1 | 0 | 3 | 11 | 7 | 42 | 34 |
| POTEJ | 0.04 | 0 | 1 | 18 | 40 | 35 | 0 | 0 |
| ABCD3 | 0 | 0 | 0 | 1 | 2 | 1 | 3 | 0 |
| ABCF1 | 0 | 0 | 0 | 3 | 1 | 5 | 6 | 7 |
| ACBD3 | 0 | 0 | 0 | 0 | 1 | 1 | 3 | 3 |
| ACBD5 | 0 | 0 | 0 | 0 | 0 | 0 | 1 | 0 |
| ACIN1 | 0 | 0 | 0 | 4 | 10 | 9 | 23 | 24 |

| | | | | | | | | |
|---|---|---|---|---|---|---|---|---|
| ADAR | 0 | 0 | 0 | 2 | 2 | 4 | 9 | 4 |
| ADD3 | 0 | 0 | 0 | 0 | 0 | 0 | 1 | 2 |
| ADNP | 0 | 0 | 0 | 9 | 12 | 6 | 20 | 13 |
| ADSL | 0 | 0 | 0 | 0 | 1 | 1 | 2 | 0 |
| AFAP1L2 | 0 | 0 | 0 | 1 | 1 | 0 | 0 | 0 |
| AFF4 | 0 | 0 | 0 | 2 | 1 | 0 | 1 | 0 |
| AGPAT5 | 0 | 0 | 0 | 0 | 0 | 0 | 1 | 0 |
| AGPAT9 | 0 | 0 | 0 | 1 | 2 | 1 | 5 | 1 |
| AHCTF1 | 0 | 0 | 0 | 0 | 0 | 0 | 8 | 4 |
| AHCY | 0 | 0 | 0 | 0 | 1 | 0 | 0 | 0 |
| AHCYL1;AHCYL2 | 0 | 0 | 0 | 1 | 2 | 1 | 1 | 3 |
| AHDC1 | 0 | 0 | 0 | 0 | 0 | 0 | 1 | 0 |
| AIFM1 | 0 | 0 | 0 | 0 | 0 | 0 | 3 | 1 |
| AK5 | 0 | 0 | 0 | 0 | 1 | 1 | 2 | 1 |
| AKAP1 | 0 | 0 | 0 | 1 | 2 | 2 | 3 | 3 |
| AKAP13 | 0 | 0 | 0 | 0 | 0 | 0 | 1 | 0 |
| AKAP8 | 0 | 0 | 0 | 1 | 1 | 1 | 1 | 0 |
| ALG13 | 0 | 0 | 0 | 1 | 2 | 1 | 1 | 1 |
| AMPD2 | 0 | 0 | 0 | 1 | 1 | 1 | 0 | 2 |
| ANAPC4 | 0 | 0 | 0 | 0 | 1 | 1 | 2 | 0 |
| ANKFY1 | 0 | 0 | 0 | 1 | 1 | 1 | 3 | 3 |
| ANKHD1 | 0 | 0 | 0 | 0 | 0 | 0 | 2 | 1 |
| ANKRD52 | 0 | 0 | 0 | 1 | 1 | 1 | 1 | 0 |
| ANO10 | 0 | 0 | 0 | 0 | 0 | 0 | 0 | 1 |
| ANP32E | 0 | 0 | 0 | 0 | 0 | 1 | 0 | 0 |
| ANXA5 | 0 | 0 | 0 | 0 | 1 | 2 | 0 | 0 |
| AP1M1 | 0 | 0 | 0 | 0 | 0 | 0 | 0 | 1 |
| AP2A2 | 0 | 0 | 0 | 1 | 3 | 2 | 0 | 0 |
| AP4B1 | 0 | 0 | 0 | 0 | 0 | 0 | 0 | 1 |
| API5 | 0 | 0 | 0 | 0 | 0 | 0 | 2 | 0 |
| ARHGAP12 | 0 | 0 | 0 | 0 | 0 | 0 | 1 | 3 |
| ARHGDIA | 0 | 0 | 0 | 0 | 1 | 1 | 0 | 0 |
| ARHGEF12 | 0 | 0 | 0 | 0 | 0 | 0 | 1 | 0 |
| ARHGEF2 | 0 | 0 | 0 | 1 | 0 | 1 | 1 | 1 |
| ARID1A | 0 | 0 | 0 | 1 | 2 | 1 | 4 | 1 |
| ARL3 | 0 | 0 | 0 | 0 | 0 | 0 | 1 | 0 |
| ARL6IP4 | 0 | 0 | 0 | 0 | 1 | 0 | 0 | 1 |
| ARMC9 | 0 | 0 | 0 | 1 | 0 | 0 | 0 | 0 |
| ASPSCR1 | 0 | 0 | 0 | 0 | 2 | 2 | 1 | 0 |
| ATG2A | 0 | 0 | 0 | 0 | 0 | 0 | 1 | 0 |
| ATP2C1 | 0 | 0 | 0 | 0 | 0 | 0 | 0 | 1 |
| ATP5J2 | 0 | 0 | 0 | 0 | 0 | 0 | 0 | 1 |
| ATRX | 0 | 0 | 0 | 0 | 0 | 1 | 11 | 8 |
| ATXN10 | 0 | 0 | 0 | 0 | 0 | 0 | 1 | 1 |
| BAP1 | 0 | 0 | 0 | 0 | 0 | 0 | 1 | 1 |

| | | | | | | | | |
|---|---|---|---|---|---|---|---|---|
| BAZ1A | 0 | 0 | 0 | 0 | 0 | 0 | 3 | 0 |
| BAZ1B | 0 | 0 | 0 | 2 | 6 | 4 | 24 | 18 |
| BBX | 0 | 0 | 0 | 0 | 0 | 0 | 7 | 1 |
| BCAP29 | 0 | 0 | 0 | 0 | 0 | 0 | 1 | 0 |
| BCAP31 | 0 | 0 | 0 | 0 | 0 | 0 | 0 | 1 |
| BCAS2 | 0 | 0 | 0 | 0 | 0 | 0 | 1 | 1 |
| BCCIP | 0 | 0 | 0 | 0 | 0 | 0 | 2 | 0 |
| BCL2L13 | 0 | 0 | 0 | 1 | 1 | 1 | 1 | 0 |
| BCL9L | 0 | 0 | 0 | 0 | 0 | 0 | 2 | 2 |
| BCLAF1 | 0 | 0 | 0 | 7 | 9 | 4 | 17 | 23 |
| BICC1 | 0 | 0 | 0 | 0 | 0 | 0 | 0 | 2 |
| BLM | 0 | 0 | 0 | 1 | 2 | 0 | 0 | 1 |
| BOD1L1 | 0 | 0 | 0 | 0 | 0 | 1 | 6 | 4 |
| BRIP1 | 0 | 0 | 0 | 2 | 0 | 0 | 15 | 4 |
| BRK1 | 0 | 0 | 0 | 1 | 1 | 1 | 1 | 2 |
| BRMS1L | 0 | 0 | 0 | 0 | 0 | 0 | 3 | 2 |
| BTF3 | 0 | 0 | 0 | 0 | 0 | 0 | 0 | 1 |
| C11orf58;SMAP | 0 | 0 | 0 | 0 | 0 | 0 | 4 | 3 |
| C17orf85 | 0 | 0 | 0 | 0 | 0 | 0 | 1 | 0 |
| C18orf21 | 0 | 0 | 0 | 0 | 0 | 0 | 0 | 1 |
| C3orf17 | 0 | 0 | 0 | 0 | 0 | 0 | 1 | 0 |
| C7orf55-LUC7L2;... | 0 | 0 | 0 | 0 | 0 | 0 | 1 | 0 |
| CACTIN | 0 | 0 | 0 | 0 | 1 | 1 | 1 | 1 |
| CACUL1 | 0 | 0 | 0 | 0 | 0 | 0 | 1 | 0 |
| CALD1 | 0 | 0 | 0 | 1 | 5 | 2 | 0 | 0 |
| CAMK4 | 0 | 0 | 0 | 2 | 1 | 1 | 1 | 0 |
| CARS | 0 | 0 | 0 | 0 | 0 | 0 | 1 | 0 |
| CASK | 0 | 0 | 0 | 0 | 1 | 0 | 0 | 0 |
| CASP4;CASP12;CASP5 | 0 | 0 | 0 | 0 | 1 | 1 | 1 | 0 |
| CAT | 0 | 0 | 0 | 0 | 0 | 0 | 0 | 1 |
| CBX3 | 0 | 0 | 0 | 1 | 0 | 0 | 2 | 2 |
| CBX5 | 0 | 0 | 0 | 0 | 0 | 0 | 2 | 0 |
| CC2D1A | 0 | 0 | 0 | 0 | 1 | 0 | 0 | 0 |
| CCAR1 | 0 | 0 | 0 | 0 | 1 | 0 | 1 | 1 |
| CCAR2 | 0 | 0 | 0 | 4 | 3 | 4 | 7 | 8 |
| CCDC115 | 0 | 0 | 0 | 0 | 1 | 0 | 0 | 1 |
| CCDC47 | 0 | 0 | 0 | 0 | 1 | 0 | 3 | 0 |
| CCDC55;NSRP1 | 0 | 0 | 0 | 0 | 0 | 0 | 1 | 2 |
| CCDC94 | 0 | 0 | 0 | 0 | 0 | 0 | 1 | 0 |
| CCNH | 0 | 0 | 0 | 0 | 0 | 0 | 0 | 1 |
| CCNK | 0 | 0 | 0 | 2 | 3 | 1 | 4 | 7 |
| CCNL1 | 0 | 0 | 0 | 2 | 1 | 2 | 2 | 0 |
| CCNL2 | 0 | 0 | 0 | 0 | 2 | 0 | 1 | 0 |
| CCNT1 | 0 | 0 | 0 | 2 | 4 | 6 | 7 | 6 |
| CCNT2 | 0 | 0 | 0 | 0 | 0 | 0 | 1 | 1 |

| | | | | | | | | |
|---|---|---|---|---|---|---|---|---|
| CCS | 0 | 0 | 0 | 0 | 0 | 1 | 0 | 0 |
| CD151 | 0 | 0 | 0 | 0 | 0 | 0 | 2 | 1 |
| CDC123 | 0 | 0 | 0 | 1 | 0 | 0 | 1 | 0 |
| CDC37 | 0 | 0 | 0 | 0 | 0 | 3 | 0 | 1 |
| CDC5L | 0 | 0 | 0 | 0 | 0 | 0 | 1 | 2 |
| CDC73 | 0 | 0 | 0 | 0 | 0 | 0 | 1 | 1 |
| CDCA2 | 0 | 0 | 0 | 0 | 0 | 0 | 3 | 0 |
| CDK12 | 0 | 0 | 0 | 0 | 1 | 1 | 4 | 2 |
| CDK13 | 0 | 0 | 0 | 0 | 0 | 1 | 1 | 0 |
| CDK2;CDK3 | 0 | 0 | 0 | 1 | 0 | 1 | 0 | 1 |
| CDK7 | 0 | 0 | 0 | 0 | 0 | 0 | 2 | 0 |
| CDK9 | 0 | 0 | 0 | 2 | 2 | 0 | 4 | 3 |
| CDKAL1 | 0 | 0 | 0 | 0 | 0 | 0 | 2 | 1 |
| CDKN2AIP | 0 | 0 | 0 | 1 | 0 | 1 | 6 | 7 |
| CDR2L | 0 | 0 | 0 | 0 | 0 | 1 | 0 | 0 |
| CENPC | 0 | 0 | 0 | 0 | 0 | 0 | 2 | 0 |
| CENPT | 0 | 0 | 0 | 0 | 0 | 0 | 1 | 0 |
| CFAP20 | 0 | 0 | 0 | 1 | 0 | 0 | 1 | 0 |
| CFAP97 | 0 | 0 | 0 | 0 | 0 | 0 | 1 | 1 |
| CFDP1 | 0 | 0 | 0 | 1 | 0 | 0 | 2 | 2 |
| CHAMP1 | 0 | 0 | 0 | 5 | 4 | 5 | 5 | 5 |
| CHD4 | 0 | 0 | 0 | 16 | 10 | 10 | 38 | 42 |
| CHD5 | 0 | 0 | 0 | 0 | 1 | 0 | 1 | 0 |
| CHD8 | 0 | 0 | 0 | 0 | 1 | 0 | 5 | 3 |
| CHERP | 0 | 0 | 0 | 5 | 5 | 7 | 8 | 6 |
| CHMP7 | 0 | 0 | 0 | 1 | 0 | 1 | 0 | 0 |
| CIC | 0 | 0 | 0 | 0 | 1 | 1 | 3 | 2 |
| CISD1 | 0 | 0 | 0 | 0 | 0 | 0 | 1 | 0 |
| CIT | 0 | 0 | 0 | 0 | 0 | 1 | 0 | 1 |
| CIZ1 | 0 | 0 | 0 | 0 | 0 | 0 | 3 | 2 |
| CLCC1 | 0 | 0 | 0 | 2 | 3 | 2 | 0 | 0 |
| CLIC4 | 0 | 0 | 0 | 0 | 0 | 1 | 0 | 0 |
| CLK3 | 0 | 0 | 0 | 0 | 1 | 0 | 0 | 0 |
| CLMN | 0 | 0 | 0 | 0 | 0 | 0 | 2 | 0 |
| CLNS1A | 0 | 0 | 0 | 0 | 1 | 0 | 1 | 2 |
| CMTR1 | 0 | 0 | 0 | 2 | 2 | 1 | 0 | 0 |
| CNGB3 | 0 | 0 | 0 | 0 | 0 | 1 | 0 | 0 |
| CNOT11 | 0 | 0 | 0 | 0 | 0 | 0 | 1 | 0 |
| COBLL1 | 0 | 0 | 0 | 0 | 1 | 2 | 6 | 0 |
| COG1 | 0 | 0 | 0 | 0 | 0 | 0 | 1 | 1 |
| COG7 | 0 | 0 | 0 | 0 | 1 | 0 | 1 | 0 |
| COIL | 0 | 0 | 0 | 0 | 0 | 0 | 2 | 1 |
| COMMD3 | 0 | 0 | 0 | 1 | 0 | 0 | 0 | 2 |
| COPS5 | 0 | 0 | 0 | 0 | 0 | 0 | 0 | 2 |
| COPS6 | 0 | 0 | 0 | 1 | 0 | 1 | 0 | 3 |

| | | | | | | | | |
|---|---|---|---|---|---|---|---|---|
| COPS7A | 0 | 0 | 0 | 0 | 0 | 0 | 1 | 1 |
| CPEB4 | 0 | 0 | 0 | 1 | 2 | 1 | 2 | 3 |
| CPSF1 | 0 | 0 | 0 | 0 | 0 | 0 | 3 | 1 |
| CPSF2 | 0 | 0 | 0 | 0 | 1 | 1 | 1 | 0 |
| CPSF4 | 0 | 0 | 0 | 0 | 0 | 0 | 3 | 1 |
| CPSF6 | 0 | 0 | 0 | 6 | 4 | 5 | 7 | 12 |
| CSNK2A1;CSNK2A3 | 0 | 0 | 0 | 4 | 6 | 2 | 1 | 4 |
| CSNK2A2 | 0 | 0 | 0 | 1 | 0 | 2 | 0 | 1 |
| CSNK2B | 0 | 0 | 0 | 2 | 3 | 4 | 2 | 2 |
| CSTF2 | 0 | 0 | 0 | 0 | 0 | 1 | 3 | 1 |
| CSTF3 | 0 | 0 | 0 | 0 | 0 | 0 | 0 | 1 |
| CTCF | 0 | 0 | 0 | 0 | 0 | 0 | 1 | 0 |
| CTSC | 0 | 0 | 0 | 0 | 1 | 0 | 0 | 0 |
| CUL1 | 0 | 0 | 0 | 1 | 0 | 0 | 1 | 0 |
| CUL2 | 0 | 0 | 0 | 3 | 2 | 3 | 0 | 0 |
| CWC15 | 0 | 0 | 0 | 0 | 1 | 1 | 5 | 5 |
| CWC22 | 0 | 0 | 0 | 1 | 0 | 0 | 1 | 1 |
| CWF19L1 | 0 | 0 | 0 | 0 | 0 | 0 | 1 | 1 |
| DAP | 0 | 0 | 0 | 0 | 0 | 0 | 0 | 1 |
| DDB1 | 0 | 0 | 0 | 0 | 0 | 2 | 2 | 1 |
| DDOST | 0 | 0 | 0 | 0 | 3 | 0 | 0 | 0 |
| DDRGK1 | 0 | 0 | 0 | 0 | 0 | 1 | 1 | 0 |
| DDX10 | 0 | 0 | 0 | 5 | 7 | 5 | 11 | 8 |
| DDX17 | 0 | 0 | 0 | 0 | 0 | 0 | 0 | 1 |
| DDX39B | 0 | 0 | 0 | 0 | 0 | 0 | 0 | 1 |
| DDX41 | 0 | 0 | 0 | 0 | 0 | 0 | 1 | 1 |
| DDX42 | 0 | 0 | 0 | 11 | 14 | 13 | 19 | 18 |
| DDX46 | 0 | 0 | 0 | 0 | 0 | 0 | 1 | 2 |
| DDX50 | 0 | 0 | 0 | 0 | 3 | 1 | 2 | 1 |
| DDX54 | 0 | 0 | 0 | 0 | 0 | 0 | 1 | 0 |
| DDX58 | 0 | 0 | 0 | 0 | 0 | 1 | 0 | 0 |
| DERL1 | 0 | 0 | 0 | 0 | 0 | 0 | 1 | 0 |
| DGKA | 0 | 0 | 0 | 0 | 0 | 0 | 0 | 1 |
| DHX15 | 0 | 0 | 0 | 5 | 8 | 6 | 9 | 11 |
| DHX35 | 0 | 0 | 0 | 0 | 0 | 1 | 2 | 1 |
| DHX36 | 0 | 0 | 0 | 0 | 0 | 0 | 1 | 1 |
| DHX38 | 0 | 0 | 0 | 2 | 2 | 0 | 3 | 5 |
| DIABLO | 0 | 0 | 0 | 0 | 1 | 0 | 0 | 0 |
| DIDO1 | 0 | 0 | 0 | 2 | 3 | 2 | 5 | 5 |
| DIP2A | 0 | 0 | 0 | 0 | 0 | 1 | 1 | 3 |
| DIS3 | 0 | 0 | 0 | 0 | 0 | 1 | 1 | 2 |
| DNAJA1 | 0 | 0 | 0 | 0 | 1 | 0 | 1 | 1 |
| DNAJC8 | 0 | 0 | 0 | 3 | 3 | 1 | 2 | 2 |
| DNAJC9 | 0 | 0 | 0 | 1 | 0 | 0 | 1 | 1 |
| DNTTIP2 | 0 | 0 | 0 | 0 | 0 | 0 | 3 | 5 |

| | | | | | | | | |
|---|---|---|---|---|---|---|---|---|
| DOCK5 | 0 | 0 | 0 | 0 | 1 | 0 | 1 | 0 |
| DPCD | 0 | 0 | 0 | 1 | 1 | 1 | 0 | 0 |
| DPM1 | 0 | 0 | 0 | 0 | 1 | 2 | 0 | 1 |
| DPY30 | 0 | 0 | 0 | 0 | 0 | 2 | 0 | 0 |
| DTD1 | 0 | 0 | 0 | 0 | 0 | 0 | 2 | 0 |
| DTL | 0 | 0 | 0 | 0 | 0 | 0 | 1 | 0 |
| ECI2 | 0 | 0 | 0 | 0 | 0 | 0 | 3 | 2 |
| EDF1 | 0 | 0 | 0 | 0 | 0 | 0 | 0 | 1 |
| EFR3A | 0 | 0 | 0 | 0 | 1 | 0 | 1 | 0 |
| EFTUD2 | 0 | 0 | 0 | 6 | 5 | 3 | 8 | 1 |
| EHD2 | 0 | 0 | 0 | 0 | 0 | 0 | 1 | 1 |
| EHD4 | 0 | 0 | 0 | 0 | 0 | 0 | 1 | 0 |
| EHMT1 | 0 | 0 | 0 | 0 | 0 | 0 | 2 | 0 |
| EIF2AK2 | 0 | 0 | 0 | 0 | 1 | 0 | 0 | 0 |
| EIF3M | 0 | 0 | 0 | 4 | 3 | 2 | 1 | 2 |
| ELOVL1 | 0 | 0 | 0 | 0 | 0 | 0 | 1 | 1 |
| ELP3 | 0 | 0 | 0 | 1 | 0 | 0 | 0 | 0 |
| EMC1 | 0 | 0 | 0 | 1 | 1 | 1 | 2 | 0 |
| EMC2 | 0 | 0 | 0 | 0 | 0 | 0 | 1 | 1 |
| EMD | 0 | 0 | 0 | 1 | 5 | 1 | 5 | 7 |
| EML4 | 0 | 0 | 0 | 0 | 0 | 0 | 2 | 0 |
| EMSY;C11orf30 | 0 | 0 | 0 | 0 | 0 | 0 | 7 | 7 |
| ENSA | 0 | 0 | 0 | 1 | 2 | 3 | 0 | 0 |
| EP400 | 0 | 0 | 0 | 0 | 0 | 0 | 2 | 2 |
| EPB41L3 | 0 | 0 | 0 | 0 | 0 | 0 | 1 | 0 |
| EPS8L2 | 0 | 0 | 0 | 1 | 0 | 1 | 1 | 0 |
| ERCC6-PGBD3;PGB... | 0 | 0 | 0 | 0 | 0 | 1 | 0 | 0 |
| ERH | 0 | 0 | 0 | 1 | 2 | 2 | 1 | 1 |
| ERLIN2 | 0 | 0 | 0 | 0 | 0 | 1 | 0 | 0 |
| ESF1 | 0 | 0 | 0 | 0 | 1 | 0 | 7 | 2 |
| EXOSC1 | 0 | 0 | 0 | 1 | 0 | 0 | 1 | 0 |
| EXOSC10 | 0 | 0 | 0 | 0 | 2 | 1 | 11 | 5 |
| EXOSC3 | 0 | 0 | 0 | 0 | 0 | 0 | 1 | 1 |
| EXOSC5 | 0 | 0 | 0 | 0 | 1 | 0 | 2 | 2 |
| EXOSC6 | 0 | 0 | 0 | 0 | 0 | 0 | 2 | 1 |
| FAM171A1 | 0 | 0 | 0 | 0 | 1 | 0 | 1 | 1 |
| FAM192A;NIP30 | 0 | 0 | 0 | 0 | 0 | 0 | 5 | 3 |
| FAM50A | 0 | 0 | 0 | 0 | 0 | 0 | 1 | 0 |
| FAM98A | 0 | 0 | 0 | 0 | 0 | 0 | 3 | 1 |
| FARSA | 0 | 0 | 0 | 1 | 2 | 3 | 5 | 4 |
| FAU | 0 | 0 | 0 | 3 | 3 | 2 | 0 | 0 |
| FBXO30 | 0 | 0 | 0 | 0 | 0 | 0 | 0 | 1 |
| FCHO2 | 0 | 0 | 0 | 2 | 0 | 2 | 5 | 0 |
| FEN1 | 0 | 0 | 0 | 3 | 3 | 1 | 6 | 8 |
| FGD6 | 0 | 0 | 0 | 0 | 0 | 0 | 3 | 1 |

| | | | | | | | | |
|---|---|---|---|---|---|---|---|---|
| FIBP | 0 | 0 | 0 | 0 | 0 | 2 | 0 | 1 |
| FIP1L1 | 0 | 0 | 0 | 1 | 3 | 2 | 8 | 8 |
| FKBP3 | 0 | 0 | 0 | 0 | 2 | 2 | 1 | 1 |
| FMNL3 | 0 | 0 | 0 | 0 | 0 | 0 | 0 | 1 |
| FMR1 | 0 | 0 | 0 | 0 | 0 | 0 | 1 | 1 |
| FNBP4 | 0 | 0 | 0 | 0 | 0 | 0 | 0 | 1 |
| FNDC3A | 0 | 0 | 0 | 2 | 0 | 0 | 3 | 3 |
| FNDC3B | 0 | 0 | 0 | 0 | 0 | 0 | 0 | 2 |
| FNIP1 | 0 | 0 | 0 | 0 | 0 | 0 | 1 | 0 |
| FOSL1 | 0 | 0 | 0 | 0 | 0 | 0 | 1 | 0 |
| FOXK1;FOXK2 | 0 | 0 | 0 | 0 | 0 | 0 | 1 | 0 |
| FRYL | 0 | 0 | 0 | 1 | 2 | 1 | 1 | 0 |
| FSCN1 | 0 | 0 | 0 | 0 | 0 | 0 | 1 | 0 |
| FTSJ3 | 0 | 0 | 0 | 0 | 1 | 0 | 5 | 1 |
| FUBP1 | 0 | 0 | 0 | 1 | 5 | 1 | 4 | 6 |
| FUBP3 | 0 | 0 | 0 | 0 | 0 | 0 | 1 | 2 |
| FUS | 0 | 0 | 0 | 1 | 0 | 0 | 3 | 3 |
| FXR2 | 0 | 0 | 0 | 0 | 0 | 0 | 1 | 3 |
| GABPA | 0 | 0 | 0 | 0 | 0 | 0 | 1 | 1 |
| GAMT | 0 | 0 | 0 | 0 | 0 | 0 | 1 | 1 |
| GANAB | 0 | 0 | 0 | 0 | 2 | 0 | 0 | 1 |
| GARS | 0 | 0 | 0 | 0 | 0 | 0 | 1 | 0 |
| GATAD2A | 0 | 0 | 0 | 0 | 0 | 0 | 3 | 1 |
| GATAD2B | 0 | 0 | 0 | 0 | 1 | 0 | 8 | 3 |
| GLYR1 | 0 | 0 | 0 | 1 | 2 | 2 | 1 | 1 |
| GMPS | 0 | 0 | 0 | 3 | 3 | 4 | 2 | 2 |
| GNL2 | 0 | 0 | 0 | 0 | 0 | 0 | 5 | 4 |
| GNL3 | 0 | 0 | 0 | 0 | 0 | 0 | 1 | 0 |
| GOPC | 0 | 0 | 0 | 0 | 0 | 0 | 0 | 1 |
| GPATCH1 | 0 | 0 | 0 | 0 | 0 | 1 | 2 | 1 |
| GPATCH8 | 0 | 0 | 0 | 0 | 0 | 0 | 2 | 0 |
| GPHN | 0 | 0 | 0 | 0 | 0 | 0 | 1 | 0 |
| GPKOW | 0 | 0 | 0 | 0 | 0 | 0 | 2 | 2 |
| GPS1 | 0 | 0 | 0 | 0 | 0 | 0 | 5 | 3 |
| GRWD1 | 0 | 0 | 0 | 0 | 0 | 0 | 1 | 0 |
| GSTCD | 0 | 0 | 0 | 0 | 0 | 0 | 1 | 0 |
| GSTO1 | 0 | 0 | 0 | 1 | 0 | 1 | 4 | 6 |
| GTF2E1 | 0 | 0 | 0 | 0 | 1 | 2 | 0 | 2 |
| GTF2E2 | 0 | 0 | 0 | 4 | 3 | 4 | 2 | 1 |
| GTF2F1 | 0 | 0 | 0 | 2 | 2 | 2 | 11 | 5 |
| GTF2F2 | 0 | 0 | 0 | 2 | 2 | 4 | 4 | 5 |
| GTF2I | 0 | 0 | 0 | 1 | 2 | 1 | 10 | 5 |
| GTF3C1 | 0 | 0 | 0 | 0 | 1 | 0 | 2 | 0 |
| GTF3C3 | 0 | 0 | 0 | 0 | 0 | 0 | 0 | 1 |
| GTF3C4 | 0 | 0 | 0 | 0 | 1 | 1 | 0 | 0 |

| | | | | | | | | |
|---|---|---|---|---|---|---|---|---|
| GTF3C5 | 0 | 0 | 0 | 0 | 0 | 0 | 1 | 3 |
| GTPBP1 | 0 | 0 | 0 | 0 | 1 | 0 | 0 | 0 |
| GYG1 | 0 | 0 | 0 | 0 | 0 | 0 | 1 | 0 |
| H2AFV;H2AFZ | 0 | 0 | 0 | 1 | 1 | 1 | 1 | 2 |
| H3F3B;H3F3A;H3F3C | 0 | 0 | 0 | 1 | 0 | 0 | 0 | 0 |
| HARS | 0 | 0 | 0 | 0 | 1 | 2 | 0 | 0 |
| HAT1 | 0 | 0 | 0 | 0 | 0 | 0 | 2 | 4 |
| HAUS5 | 0 | 0 | 0 | 0 | 1 | 1 | 0 | 0 |
| HAUS8 | 0 | 0 | 0 | 1 | 1 | 1 | 2 | 0 |
| HDAC1 | 0 | 0 | 0 | 0 | 0 | 0 | 3 | 3 |
| HDGF | 0 | 0 | 0 | 1 | 2 | 0 | 3 | 3 |
| HDGFRP2 | 0 | 0 | 0 | 3 | 5 | 5 | 6 | 8 |
| HEATR5A | 0 | 0 | 0 | 1 | 1 | 1 | 0 | 0 |
| HEXIM1 | 0 | 0 | 0 | 0 | 1 | 0 | 0 | 1 |
| HIGD1A | 0 | 0 | 0 | 0 | 0 | 0 | 1 | 1 |
| HK1 | 0 | 0 | 0 | 0 | 0 | 0 | 2 | 2 |
| HLA-C;HLA-B;HLA-A;HLA-H | 0 | 0 | 0 | 0 | 0 | 0 | 1 | 1 |
| HMGA1 | 0 | 0 | 0 | 4 | 4 | 5 | 3 | 5 |
| HMGA2 | 0 | 0 | 0 | 2 | 3 | 0 | 0 | 2 |
| HMGB1;HMGB1P1 | 0 | 0 | 0 | 0 | 1 | 0 | 1 | 1 |
| HMGN2 | 0 | 0 | 0 | 0 | 0 | 0 | 0 | 1 |
| HMGXB4 | 0 | 0 | 0 | 0 | 0 | 0 | 1 | 1 |
| HNRNPA2B1 | 0 | 0 | 0 | 1 | 1 | 0 | 5 | 8 |
| HNRNPAB | 0 | 0 | 0 | 1 | 3 | 1 | 0 | 1 |
| HNRNPD | 0 | 0 | 0 | 5 | 4 | 1 | 3 | 1 |
| HNRNPH3 | 0 | 0 | 0 | 0 | 0 | 0 | 1 | 0 |
| HNRNPL | 0 | 0 | 0 | 2 | 0 | 1 | 10 | 6 |
| HNRNPUL1 | 0 | 0 | 0 | 3 | 2 | 1 | 2 | 1 |
| HOOK3 | 0 | 0 | 0 | 0 | 0 | 1 | 0 | 0 |
| HP1BP3 | 0 | 0 | 0 | 9 | 9 | 11 | 15 | 16 |
| HSPA1L | 0 | 0 | 0 | 2 | 0 | 0 | 0 | 0 |
| HTATSF1 | 0 | 0 | 0 | 1 | 2 | 2 | 1 | 1 |
| HUWE1 | 0 | 0 | 0 | 0 | 1 | 2 | 0 | 0 |
| HYOU1 | 0 | 0 | 0 | 0 | 1 | 0 | 0 | 0 |
| IDH1 | 0 | 0 | 0 | 3 | 1 | 1 | 1 | 1 |
| IFI16 | 0 | 0 | 0 | 4 | 6 | 4 | 10 | 11 |
| IFIT3 | 0 | 0 | 0 | 0 | 0 | 0 | 0 | 1 |
| IGF2BP3 | 0 | 0 | 0 | 1 | 2 | 1 | 3 | 1 |
| IK | 0 | 0 | 0 | 0 | 1 | 0 | 6 | 4 |
| IKBKAP | 0 | 0 | 0 | 0 | 1 | 0 | 1 | 3 |
| ILF3 | 0 | 0 | 0 | 0 | 0 | 0 | 2 | 1 |
| ILKAP;ILKAP3 | 0 | 0 | 0 | 0 | 0 | 0 | 2 | 2 |
| INTS12 | 0 | 0 | 0 | 0 | 0 | 0 | 2 | 4 |
| INTS3 | 0 | 0 | 0 | 1 | 3 | 3 | 1 | 1 |
| INTS9 | 0 | 0 | 0 | 0 | 0 | 1 | 0 | 0 |

| | | | | | | | | |
|---|---|---|---|---|---|---|---|---|
| IPO5 | 0 | 0 | 0 | 3 | 1 | 1 | 0 | 0 |
| IRF2BP2 | 0 | 0 | 0 | 0 | 0 | 0 | 0 | 1 |
| IRF2BPL | 0 | 0 | 0 | 0 | 0 | 0 | 2 | 5 |
| IRS2 | 0 | 0 | 0 | 0 | 0 | 0 | 0 | 1 |
| IST1 | 0 | 0 | 0 | 1 | 0 | 0 | 3 | 1 |
| ISY1 | 0 | 0 | 0 | 0 | 2 | 0 | 0 | 0 |
| ITGA2 | 0 | 0 | 0 | 1 | 1 | 1 | 2 | 0 |
| ITGAV | 0 | 0 | 0 | 0 | 0 | 2 | 1 | 0 |
| ITGB5 | 0 | 0 | 0 | 0 | 2 | 1 | 1 | 2 |
| ITGB6 | 0 | 0 | 0 | 1 | 1 | 1 | 0 | 1 |
| ITPR3 | 0 | 0 | 0 | 0 | 1 | 0 | 1 | 0 |
| ITSN1 | 0 | 0 | 0 | 0 | 0 | 0 | 0 | 1 |
| ITSN2 | 0 | 0 | 0 | 0 | 0 | 1 | 3 | 1 |
| JUN | 0 | 0 | 0 | 0 | 0 | 0 | 1 | 0 |
| JUNB | 0 | 0 | 0 | 1 | 0 | 0 | 1 | 2 |
| KDELR1 | 0 | 0 | 0 | 0 | 0 | 0 | 0 | 1 |
| KDM3B | 0 | 0 | 0 | 3 | 6 | 3 | 3 | 7 |
| KHDRBS1 | 0 | 0 | 0 | 0 | 0 | 0 | 1 | 1 |
| KIAA1143 | 0 | 0 | 0 | 0 | 2 | 0 | 3 | 1 |
| KIAA1468 | 0 | 0 | 0 | 1 | 1 | 0 | 0 | 0 |
| KIDINS220 | 0 | 0 | 0 | 0 | 0 | 0 | 2 | 0 |
| KIF22 | 0 | 0 | 0 | 0 | 0 | 0 | 1 | 0 |
| KIF23 | 0 | 0 | 0 | 1 | 1 | 0 | 3 | 2 |
| KIFC3 | 0 | 0 | 0 | 0 | 0 | 0 | 0 | 1 |
| KIZ | 0 | 0 | 0 | 1 | 0 | 2 | 0 | 0 |
| KPNA1 | 0 | 0 | 0 | 0 | 0 | 0 | 1 | 1 |
| KPNA4 | 0 | 0 | 0 | 5 | 4 | 4 | 3 | 5 |
| KPNA6;KPNA5 | 0 | 0 | 0 | 1 | 0 | 0 | 2 | 1 |
| KTN1 | 0 | 0 | 0 | 1 | 1 | 0 | 3 | 0 |
| LARP1B | 0 | 0 | 0 | 1 | 0 | 1 | 0 | 2 |
| LARP4 | 0 | 0 | 0 | 0 | 1 | 0 | 3 | 3 |
| LEMD3 | 0 | 0 | 0 | 2 | 1 | 1 | 7 | 7 |
| LENG8 | 0 | 0 | 0 | 0 | 0 | 0 | 1 | 1 |
| LEO1 | 0 | 0 | 0 | 1 | 0 | 0 | 1 | 0 |
| LEPROT | 0 | 0 | 0 | 0 | 0 | 0 | 1 | 1 |
| LIG1 | 0 | 0 | 0 | 0 | 1 | 0 | 0 | 0 |
| LIG3 | 0 | 0 | 0 | 0 | 0 | 0 | 0 | 2 |
| LIN37 | 0 | 0 | 0 | 0 | 1 | 1 | 1 | 0 |
| LIN54 | 0 | 0 | 0 | 0 | 0 | 0 | 1 | 1 |
| LIN7C;LIN7A | 0 | 0 | 0 | 1 | 0 | 0 | 0 | 0 |
| LLGL1 | 0 | 0 | 0 | 0 | 1 | 0 | 1 | 0 |
| LMAN1 | 0 | 0 | 0 | 0 | 0 | 0 | 1 | 0 |
| LOXL2 | 0 | 0 | 0 | 0 | 1 | 0 | 0 | 0 |
| LRP1 | 0 | 0 | 0 | 0 | 1 | 0 | 0 | 0 |
| LRRC59 | 0 | 0 | 0 | 0 | 1 | 1 | 0 | 1 |

| | | | | | | | | |
|---|---|---|---|---|---|---|---|---|
| LRWD1 | 0 | 0 | 0 | 6 | 4 | 4 | 11 | 8 |
| LSG1 | 0 | 0 | 0 | 2 | 1 | 1 | 5 | 1 |
| LSM14B | 0 | 0 | 0 | 0 | 0 | 0 | 0 | 1 |
| LSM2 | 0 | 0 | 0 | 1 | 2 | 0 | 1 | 1 |
| LSM4 | 0 | 0 | 0 | 0 | 0 | 0 | 2 | 2 |
| LXN | 0 | 0 | 0 | 0 | 0 | 0 | 1 | 1 |
| LYPLAL1 | 0 | 0 | 0 | 0 | 0 | 2 | 0 | 0 |
| MAP2 | 0 | 0 | 0 | 0 | 1 | 0 | 2 | 2 |
| MARK3 | 0 | 0 | 0 | 0 | 0 | 0 | 2 | 1 |
| MAST4 | 0 | 0 | 0 | 0 | 0 | 1 | 0 | 0 |
| MASTL | 0 | 0 | 0 | 1 | 2 | 3 | 3 | 3 |
| MATR3 | 0 | 0 | 0 | 5 | 9 | 7 | 9 | 8 |
| MB21D2 | 0 | 0 | 0 | 0 | 0 | 0 | 0 | 1 |
| MBNL1 | 0 | 0 | 0 | 0 | 0 | 0 | 1 | 1 |
| MBOAT7 | 0 | 0 | 0 | 1 | 3 | 3 | 1 | 1 |
| MCM3 | 0 | 0 | 0 | 0 | 0 | 1 | 2 | 2 |
| MCM3AP | 0 | 0 | 0 | 0 | 0 | 0 | 1 | 0 |
| MCM5 | 0 | 0 | 0 | 0 | 0 | 0 | 2 | 1 |
| MCM6 | 0 | 0 | 0 | 1 | 4 | 0 | 4 | 4 |
| MCU | 0 | 0 | 0 | 0 | 0 | 0 | 0 | 1 |
| MDC1 | 0 | 0 | 0 | 0 | 1 | 0 | 1 | 2 |
| MED1 | 0 | 0 | 0 | 1 | 0 | 0 | 8 | 7 |
| MEF2D | 0 | 0 | 0 | 1 | 2 | 2 | 4 | 4 |
| MEPCE | 0 | 0 | 0 | 0 | 0 | 0 | 2 | 0 |
| MEST | 0 | 0 | 0 | 0 | 0 | 0 | 0 | 1 |
| METTL14 | 0 | 0 | 0 | 0 | 0 | 0 | 1 | 0 |
| METTL3 | 0 | 0 | 0 | 0 | 0 | 0 | 2 | 0 |
| MFF | 0 | 0 | 0 | 3 | 0 | 2 | 1 | 3 |
| MFSD10 | 0 | 0 | 0 | 1 | 0 | 0 | 0 | 0 |
| MICAL2 | 0 | 0 | 0 | 0 | 0 | 0 | 1 | 0 |
| MICAL3 | 0 | 0 | 0 | 0 | 0 | 0 | 1 | 0 |
| MINK1 | 0 | 0 | 0 | 0 | 0 | 0 | 1 | 0 |
| MIOS | 0 | 0 | 0 | 0 | 0 | 0 | 1 | 0 |
| MKI67 | 0 | 0 | 0 | 24 | 38 | 29 | 74 | 70 |
| MLH1 | 0 | 0 | 0 | 1 | 0 | 2 | 13 | 8 |
| MLLT1 | 0 | 0 | 0 | 1 | 0 | 0 | 0 | 1 |
| MLLT6 | 0 | 0 | 0 | 0 | 0 | 0 | 2 | 0 |
| MOB3A | 0 | 0 | 0 | 0 | 0 | 0 | 0 | 1 |
| MOV10 | 0 | 0 | 0 | 1 | 1 | 3 | 0 | 0 |
| MPG | 0 | 0 | 0 | 1 | 0 | 1 | 0 | 0 |
| MPP5 | 0 | 0 | 0 | 1 | 0 | 0 | 0 | 0 |
| MPZL1 | 0 | 0 | 0 | 0 | 0 | 0 | 0 | 1 |
| MRPL4 | 0 | 0 | 0 | 1 | 0 | 0 | 0 | 0 |
| MSH2 | 0 | 0 | 0 | 2 | 2 | 2 | 4 | 2 |
| MSH6 | 0 | 0 | 0 | 2 | 1 | 1 | 0 | 0 |

| | | | | | | | | |
|---|---|---|---|---|---|---|---|---|
| MSRB3 | 0 | 0 | 0 | 0 | 0 | 1 | 1 | 1 |
| MT-ATP6 | 0 | 0 | 0 | 0 | 0 | 0 | 0 | 1 |
| MTA1 | 0 | 0 | 0 | 1 | 1 | 1 | 2 | 3 |
| MTA2 | 0 | 0 | 0 | 2 | 1 | 1 | 4 | 2 |
| MTDH | 0 | 0 | 0 | 1 | 2 | 0 | 9 | 10 |
| MXRA7 | 0 | 0 | 0 | 0 | 0 | 0 | 2 | 0 |
| MYBBP1A | 0 | 0 | 0 | 1 | 0 | 0 | 2 | 0 |
| MYL9 | 0 | 0 | 0 | 2 | 3 | 0 | 0 | 0 |
| MYO18A | 0 | 0 | 0 | 2 | 5 | 14 | 0 | 0 |
| NAA15 | 0 | 0 | 0 | 0 | 1 | 1 | 1 | 1 |
| NAA30 | 0 | 0 | 0 | 0 | 0 | 0 | 0 | 1 |
| NAA35 | 0 | 0 | 0 | 0 | 0 | 0 | 1 | 1 |
| NAA50 | 0 | 0 | 0 | 1 | 3 | 1 | 0 | 0 |
| NACA | 0 | 0 | 0 | 2 | 3 | 5 | 2 | 4 |
| NAMPT | 0 | 0 | 0 | 0 | 0 | 1 | 1 | 0 |
| NANS | 0 | 0 | 0 | 0 | 2 | 0 | 0 | 0 |
| NASP | 0 | 0 | 0 | 3 | 8 | 3 | 11 | 15 |
| NAT10 | 0 | 0 | 0 | 1 | 0 | 0 | 5 | 6 |
| NAV3 | 0 | 0 | 0 | 0 | 0 | 0 | 3 | 1 |
| NBAS | 0 | 0 | 0 | 1 | 0 | 1 | 1 | 1 |
| NBN | 0 | 0 | 0 | 0 | 2 | 0 | 6 | 0 |
| NCBP1 | 0 | 0 | 0 | 0 | 0 | 0 | 3 | 3 |
| NCEH1 | 0 | 0 | 0 | 0 | 0 | 0 | 1 | 0 |
| NCLN | 0 | 0 | 0 | 0 | 0 | 0 | 0 | 1 |
| NCOA6 | 0 | 0 | 0 | 0 | 0 | 0 | 1 | 1 |
| NCOR1 | 0 | 0 | 0 | 0 | 0 | 0 | 2 | 0 |
| NCOR2 | 0 | 0 | 0 | 3 | 5 | 3 | 4 | 4 |
| NDE1;NDEL1 | 0 | 0 | 0 | 0 | 0 | 2 | 0 | 0 |
| NEDD8 | 0 | 0 | 0 | 0 | 1 | 0 | 0 | 0 |
| NEK7 | 0 | 0 | 0 | 0 | 0 | 0 | 1 | 0 |
| NELFA | 0 | 0 | 0 | 1 | 0 | 0 | 2 | 0 |
| NELFB | 0 | 0 | 0 | 1 | 1 | 1 | 2 | 2 |
| NELFCD;TH1L | 0 | 0 | 0 | 1 | 1 | 0 | 1 | 3 |
| NELFE | 0 | 0 | 0 | 1 | 3 | 1 | 8 | 5 |
| NF1 | 0 | 0 | 0 | 0 | 0 | 0 | 0 | 1 |
| NFIA | 0 | 0 | 0 | 0 | 1 | 0 | 2 | 2 |
| NFRKB | 0 | 0 | 0 | 0 | 0 | 0 | 1 | 0 |
| NHP2L1 | 0 | 0 | 0 | 1 | 0 | 0 | 3 | 2 |
| NME7 | 0 | 0 | 0 | 1 | 0 | 0 | 0 | 0 |
| NMT1 | 0 | 0 | 0 | 0 | 1 | 2 | 5 | 2 |
| NMT2 | 0 | 0 | 0 | 0 | 0 | 0 | 0 | 1 |
| NOC2L | 0 | 0 | 0 | 0 | 0 | 0 | 0 | 1 |
| NOL11 | 0 | 0 | 0 | 0 | 0 | 0 | 1 | 0 |
| NOL6 | 0 | 0 | 0 | 0 | 1 | 1 | 0 | 0 |
| NOL8 | 0 | 0 | 0 | 0 | 0 | 0 | 2 | 0 |

| | | | | | | | | |
|---|---|---|---|---|---|---|---|---|
| NOLC1 | 0 | 0 | 0 | 1 | 0 | 0 | 3 | 3 |
| NOP56 | 0 | 0 | 0 | 0 | 0 | 0 | 1 | 0 |
| NPM3 | 0 | 0 | 0 | 2 | 2 | 3 | 2 | 1 |
| NR2C1 | 0 | 0 | 0 | 0 | 0 | 0 | 1 | 2 |
| NR2C2 | 0 | 0 | 0 | 0 | 0 | 0 | 4 | 1 |
| NSFL1C | 0 | 0 | 0 | 1 | 2 | 2 | 3 | 1 |
| NUCKS1 | 0 | 0 | 0 | 0 | 0 | 0 | 4 | 7 |
| NUDCD1 | 0 | 0 | 0 | 0 | 0 | 0 | 4 | 2 |
| NUDCD2 | 0 | 0 | 0 | 0 | 0 | 0 | 0 | 1 |
| NUDT2 | 0 | 0 | 0 | 1 | 0 | 0 | 0 | 1 |
| NUDT21 | 0 | 0 | 0 | 4 | 9 | 9 | 9 | 5 |
| NUP107 | 0 | 0 | 0 | 0 | 0 | 0 | 2 | 0 |
| NUP133 | 0 | 0 | 0 | 1 | 2 | 2 | 4 | 1 |
| NUP153 | 0 | 0 | 0 | 7 | 9 | 6 | 36 | 27 |
| NUP205 | 0 | 0 | 0 | 0 | 0 | 0 | 0 | 2 |
| NUP50 | 0 | 0 | 0 | 1 | 2 | 1 | 9 | 11 |
| NUP54 | 0 | 0 | 0 | 1 | 0 | 0 | 3 | 1 |
| NUPL1 | 0 | 0 | 0 | 0 | 0 | 0 | 1 | 1 |
| OCIAD1 | 0 | 0 | 0 | 0 | 0 | 0 | 2 | 0 |
| OGDH | 0 | 0 | 0 | 0 | 1 | 1 | 0 | 0 |
| OGT | 0 | 0 | 0 | 1 | 2 | 0 | 4 | 1 |
| ORC2 | 0 | 0 | 0 | 2 | 4 | 5 | 3 | 8 |
| ORC3 | 0 | 0 | 0 | 1 | 2 | 1 | 0 | 1 |
| OSBPL3 | 0 | 0 | 0 | 0 | 0 | 0 | 1 | 2 |
| OTUD4 | 0 | 0 | 0 | 1 | 0 | 2 | 1 | 1 |
| OXSR1 | 0 | 0 | 0 | 2 | 1 | 0 | 3 | 1 |
| P4HA1 | 0 | 0 | 0 | 0 | 2 | 2 | 1 | 3 |
| P4HB | 0 | 0 | 0 | 0 | 0 | 0 | 2 | 0 |
| PACSIN2 | 0 | 0 | 0 | 0 | 1 | 1 | 1 | 1 |
| PAF1 | 0 | 0 | 0 | 0 | 1 | 0 | 2 | 1 |
| PAPOLA | 0 | 0 | 0 | 12 | 11 | 10 | 20 | 23 |
| PAPSS1 | 0 | 0 | 0 | 0 | 0 | 0 | 3 | 1 |
| PARG | 0 | 0 | 0 | 1 | 0 | 0 | 0 | 1 |
| PARP1 | 0 | 0 | 0 | 0 | 0 | 0 | 1 | 1 |
| PARP12 | 0 | 0 | 0 | 0 | 0 | 0 | 0 | 1 |
| PARP4 | 0 | 0 | 0 | 2 | 3 | 6 | 1 | 1 |
| PARVA | 0 | 0 | 0 | 2 | 3 | 3 | 1 | 2 |
| PASK | 0 | 0 | 0 | 0 | 1 | 0 | 0 | 0 |
| PAXBP1 | 0 | 0 | 0 | 0 | 0 | 0 | 1 | 1 |
| PCF11 | 0 | 0 | 0 | 0 | 0 | 0 | 1 | 0 |
| PCYT1A | 0 | 0 | 0 | 1 | 2 | 2 | 5 | 6 |
| PCYT2 | 0 | 0 | 0 | 0 | 0 | 0 | 1 | 0 |
| PDCD5 | 0 | 0 | 0 | 0 | 1 | 0 | 3 | 0 |
| PDIA3 | 0 | 0 | 0 | 0 | 0 | 0 | 0 | 2 |
| PDIA6 | 0 | 0 | 0 | 0 | 0 | 0 | 0 | 1 |

| | | | | | | | | |
|---|---|---|---|---|---|---|---|---|
| PDRG1 | 0 | 0 | 0 | 0 | 0 | 0 | 0 | 1 |
| PDS5A | 0 | 0 | 0 | 1 | 0 | 0 | 0 | 0 |
| PDS5B | 0 | 0 | 0 | 1 | 1 | 0 | 3 | 3 |
| PDXDC1 | 0 | 0 | 0 | 0 | 0 | 1 | 2 | 0 |
| PES1 | 0 | 0 | 0 | 0 | 0 | 0 | 5 | 3 |
| PFAS | 0 | 0 | 0 | 0 | 0 | 0 | 2 | 0 |
| PFDN2 | 0 | 0 | 0 | 0 | 0 | 0 | 1 | 0 |
| PHAX | 0 | 0 | 0 | 1 | 0 | 1 | 4 | 5 |
| PHC3;PHC2 | 0 | 0 | 0 | 0 | 0 | 0 | 1 | 0 |
| PHF3 | 0 | 0 | 0 | 0 | 0 | 0 | 3 | 1 |
| PIK3C2A | 0 | 0 | 0 | 1 | 0 | 0 | 2 | 2 |
| PIK3R2 | 0 | 0 | 0 | 0 | 0 | 0 | 1 | 0 |
| PKN2 | 0 | 0 | 0 | 0 | 0 | 0 | 3 | 1 |
| PLOD2 | 0 | 0 | 0 | 0 | 0 | 0 | 1 | 0 |
| PLRG1 | 0 | 0 | 0 | 0 | 0 | 0 | 2 | 0 |
| PML | 0 | 0 | 0 | 5 | 10 | 12 | 17 | 15 |
| PMS2 | 0 | 0 | 0 | 1 | 0 | 0 | 1 | 1 |
| PNISR | 0 | 0 | 0 | 2 | 1 | 0 | 4 | 3 |
| POLA1 | 0 | 0 | 0 | 0 | 0 | 0 | 3 | 0 |
| POLA2 | 0 | 0 | 0 | 0 | 1 | 3 | 0 | 0 |
| POLD1 | 0 | 0 | 0 | 2 | 11 | 10 | 6 | 6 |
| POLD3 | 0 | 0 | 0 | 0 | 1 | 0 | 3 | 2 |
| POLDIP3;PDIP46 | 0 | 0 | 0 | 0 | 0 | 0 | 2 | 0 |
| POLH | 0 | 0 | 0 | 0 | 0 | 0 | 1 | 1 |
| POLR2C | 0 | 0 | 0 | 0 | 1 | 0 | 0 | 0 |
| POLR2H | 0 | 0 | 0 | 1 | 1 | 0 | 0 | 0 |
| POM121C;POM121 | 0 | 0 | 0 | 0 | 0 | 0 | 1 | 1 |
| POP4 | 0 | 0 | 0 | 0 | 1 | 1 | 0 | 1 |
| POP5 | 0 | 0 | 0 | 1 | 0 | 0 | 1 | 1 |
| PPFIBP1 | 0 | 0 | 0 | 2 | 4 | 5 | 6 | 2 |
| PPIH | 0 | 0 | 0 | 0 | 0 | 0 | 1 | 1 |
| PPIL1 | 0 | 0 | 0 | 0 | 0 | 0 | 1 | 0 |
| PPIL4 | 0 | 0 | 0 | 1 | 1 | 1 | 0 | 2 |
| PPIP5K2;PPIP5K1 | 0 | 0 | 0 | 1 | 1 | 0 | 2 | 2 |
| PPM1G | 0 | 0 | 0 | 1 | 0 | 2 | 4 | 4 |
| PPP1R10 | 0 | 0 | 0 | 15 | 9 | 9 | 16 | 17 |
| PPP1R2;PPP1R2P3 | 0 | 0 | 0 | 0 | 0 | 0 | 1 | 1 |
| PPP1R7 | 0 | 0 | 0 | 0 | 0 | 1 | 1 | 0 |
| PPP4C | 0 | 0 | 0 | 1 | 0 | 0 | 4 | 1 |
| PPP4R1 | 0 | 0 | 0 | 0 | 0 | 1 | 0 | 0 |
| PPP4R2 | 0 | 0 | 0 | 0 | 2 | 2 | 4 | 3 |
| PPP6R1 | 0 | 0 | 0 | 1 | 1 | 2 | 2 | 3 |
| PRAF2 | 0 | 0 | 0 | 0 | 1 | 0 | 1 | 1 |
| PRDX2 | 0 | 0 | 0 | 2 | 2 | 2 | 2 | 3 |
| PRDX5 | 0 | 0 | 0 | 0 | 0 | 1 | 1 | 0 |

| | | | | | | | | |
|---|---|---|---|---|---|---|---|---|
| PRIM2 | 0 | 0 | 0 | 0 | 0 | 0 | 1 | 1 |
| PRKAB1 | 0 | 0 | 0 | 0 | 1 | 0 | 0 | 0 |
| PRKAR2A | 0 | 0 | 0 | 0 | 1 | 1 | 6 | 1 |
| PRMT9 | 0 | 0 | 0 | 0 | 0 | 1 | 0 | 0 |
| PRPF19 | 0 | 0 | 0 | 1 | 1 | 0 | 7 | 8 |
| PRPF3 | 0 | 0 | 0 | 1 | 3 | 1 | 12 | 10 |
| PRPF31 | 0 | 0 | 0 | 0 | 0 | 2 | 2 | 3 |
| PRPF4 | 0 | 0 | 0 | 4 | 7 | 4 | 11 | 9 |
| PRPF6 | 0 | 0 | 0 | 0 | 0 | 0 | 0 | 2 |
| PRR16 | 0 | 0 | 0 | 1 | 1 | 1 | 1 | 1 |
| PRR5L | 0 | 0 | 0 | 0 | 1 | 0 | 0 | 0 |
| PRRC1 | 0 | 0 | 0 | 0 | 0 | 0 | 1 | 1 |
| PSIP1 | 0 | 0 | 0 | 0 | 0 | 0 | 1 | 1 |
| PSMA1 | 0 | 0 | 0 | 0 | 1 | 1 | 1 | 1 |
| PSMB1 | 0 | 0 | 0 | 2 | 1 | 1 | 0 | 0 |
| PSMB2 | 0 | 0 | 0 | 0 | 0 | 0 | 1 | 0 |
| PSMC4 | 0 | 0 | 0 | 1 | 4 | 5 | 2 | 2 |
| PSMD11 | 0 | 0 | 0 | 1 | 4 | 4 | 0 | 1 |
| PSMD5 | 0 | 0 | 0 | 0 | 1 | 1 | 0 | 0 |
| PSMD6 | 0 | 0 | 0 | 0 | 2 | 1 | 0 | 0 |
| PSMD7 | 0 | 0 | 0 | 1 | 1 | 1 | 0 | 0 |
| PSME1 | 0 | 0 | 0 | 0 | 0 | 1 | 0 | 0 |
| PSME2 | 0 | 0 | 0 | 2 | 3 | 3 | 1 | 3 |
| PSME3 | 0 | 0 | 0 | 3 | 6 | 4 | 7 | 4 |
| PTMA | 0 | 0 | 0 | 0 | 0 | 0 | 1 | 4 |
| PTMS | 0 | 0 | 0 | 0 | 0 | 0 | 0 | 1 |
| PTPN1 | 0 | 0 | 0 | 1 | 1 | 2 | 7 | 6 |
| PUM2;PUM1 | 0 | 0 | 0 | 0 | 0 | 0 | 0 | 2 |
| PVRL2 | 0 | 0 | 0 | 0 | 0 | 0 | 1 | 1 |
| PVRL3 | 0 | 0 | 0 | 0 | 0 | 0 | 0 | 1 |
| QRICH1 | 0 | 0 | 0 | 0 | 1 | 0 | 0 | 1 |
| QSER1 | 0 | 0 | 0 | 0 | 0 | 0 | 1 | 0 |
| R3HCC1L | 0 | 0 | 0 | 2 | 1 | 0 | 3 | 2 |
| R3HDM1 | 0 | 0 | 0 | 0 | 0 | 0 | 1 | 1 |
| RAB27B;RAB27A | 0 | 0 | 0 | 0 | 1 | 1 | 0 | 1 |
| RAB5A | 0 | 0 | 0 | 0 | 0 | 0 | 1 | 1 |
| RABL3 | 0 | 0 | 0 | 0 | 0 | 0 | 2 | 1 |
| RABL6 | 0 | 0 | 0 | 1 | 1 | 1 | 1 | 1 |
| RACGAP1 | 0 | 0 | 0 | 1 | 1 | 0 | 3 | 0 |
| RAD21 | 0 | 0 | 0 | 0 | 0 | 0 | 1 | 0 |
| RAD50 | 0 | 0 | 0 | 1 | 2 | 1 | 1 | 2 |
| RAI1 | 0 | 0 | 0 | 0 | 0 | 0 | 0 | 2 |
| RAP1GDS1 | 0 | 0 | 0 | 0 | 1 | 0 | 0 | 0 |
| RARS2 | 0 | 0 | 0 | 1 | 0 | 0 | 1 | 0 |
| RASA1 | 0 | 0 | 0 | 1 | 1 | 0 | 0 | 1 |

| | | | | | | | | |
|---|---|---|---|---|---|---|---|---|
| RAVER1 | 0 | 0 | 0 | 0 | 0 | 2 | 1 | 1 |
| RBBP4 | 0 | 0 | 0 | 1 | 2 | 2 | 3 | 2 |
| RBBP6 | 0 | 0 | 0 | 1 | 0 | 1 | 5 | 5 |
| RBFOX2;RBFOX1;RBFOX3 | 0 | 0 | 0 | 0 | 0 | 0 | 0 | 1 |
| RBM10 | 0 | 0 | 0 | 6 | 6 | 6 | 8 | 11 |
| RBM17 | 0 | 0 | 0 | 0 | 3 | 3 | 14 | 5 |
| RBM26 | 0 | 0 | 0 | 3 | 9 | 8 | 13 | 17 |
| RBM27 | 0 | 0 | 0 | 3 | 2 | 2 | 5 | 7 |
| RBM33 | 0 | 0 | 0 | 0 | 1 | 2 | 5 | 4 |
| RBM39 | 0 | 0 | 0 | 2 | 1 | 1 | 2 | 0 |
| RBM7 | 0 | 0 | 0 | 1 | 0 | 0 | 1 | 0 |
| RBMX2 | 0 | 0 | 0 | 0 | 0 | 0 | 1 | 0 |
| RBX1 | 0 | 0 | 0 | 0 | 0 | 0 | 2 | 2 |
| RC3H2;RC3H1 | 0 | 0 | 0 | 0 | 0 | 0 | 0 | 1 |
| RELL1 | 0 | 0 | 0 | 1 | 0 | 0 | 1 | 2 |
| RFC4 | 0 | 0 | 0 | 0 | 0 | 0 | 1 | 2 |
| RFTN1 | 0 | 0 | 0 | 0 | 0 | 0 | 0 | 1 |
| RFX1 | 0 | 0 | 0 | 2 | 0 | 0 | 1 | 0 |
| RFX5 | 0 | 0 | 0 | 0 | 0 | 0 | 1 | 0 |
| RGPD1;RGPD2 | 0 | 0 | 0 | 1 | 1 | 1 | 0 | 0 |
| RICTOR | 0 | 0 | 0 | 1 | 1 | 1 | 2 | 2 |
| RIF1 | 0 | 0 | 0 | 0 | 1 | 0 | 14 | 3 |
| RNF20 | 0 | 0 | 0 | 0 | 0 | 0 | 3 | 1 |
| RNF213 | 0 | 0 | 0 | 0 | 0 | 0 | 1 | 1 |
| RNF40 | 0 | 0 | 0 | 0 | 1 | 0 | 0 | 1 |
| RNPS1 | 0 | 0 | 0 | 0 | 1 | 0 | 5 | 5 |
| ROBO1 | 0 | 0 | 0 | 0 | 0 | 0 | 0 | 1 |
| RPA1 | 0 | 0 | 0 | 2 | 1 | 4 | 8 | 12 |
| RPA2 | 0 | 0 | 0 | 0 | 0 | 0 | 1 | 1 |
| RPA3 | 0 | 0 | 0 | 1 | 1 | 2 | 1 | 1 |
| RPLP1 | 0 | 0 | 0 | 1 | 3 | 1 | 1 | 0 |
| RPP25L | 0 | 0 | 0 | 0 | 1 | 1 | 0 | 1 |
| RPP30 | 0 | 0 | 0 | 1 | 4 | 4 | 1 | 1 |
| RPP38 | 0 | 0 | 0 | 0 | 0 | 0 | 2 | 1 |
| RPP40 | 0 | 0 | 0 | 0 | 1 | 1 | 1 | 1 |
| RPRD1B | 0 | 0 | 0 | 1 | 2 | 2 | 0 | 0 |
| RPRD2 | 0 | 0 | 0 | 0 | 3 | 1 | 8 | 4 |
| RPS21 | 0 | 0 | 0 | 0 | 0 | 0 | 2 | 1 |
| RPS27L;RPS27 | 0 | 0 | 0 | 0 | 5 | 0 | 0 | 0 |
| RPS6KC1 | 0 | 0 | 0 | 0 | 0 | 1 | 1 | 2 |
| RPSA | 0 | 0 | 0 | 0 | 0 | 0 | 1 | 0 |
| RRP12 | 0 | 0 | 0 | 0 | 0 | 0 | 2 | 0 |
| RSL1D1 | 0 | 0 | 0 | 1 | 2 | 0 | 0 | 0 |
| RSRC2 | 0 | 0 | 0 | 0 | 0 | 0 | 1 | 1 |
| RTF1 | 0 | 0 | 0 | 0 | 0 | 0 | 4 | 4 |

| | | | | | | | | |
|---|---|---|---|---|---|---|---|---|
| RUFY3 | 0 | 0 | 0 | 1 | 0 | 1 | 0 | 0 |
| SAFB2;SAFB | 0 | 0 | 0 | 0 | 0 | 0 | 4 | 4 |
| SAP130 | 0 | 0 | 0 | 0 | 1 | 1 | 3 | 4 |
| SAP18 | 0 | 0 | 0 | 0 | 0 | 0 | 4 | 5 |
| SAR1A;SAR1B | 0 | 0 | 0 | 0 | 1 | 0 | 0 | 0 |
| SARNP | 0 | 0 | 0 | 1 | 0 | 0 | 2 | 2 |
| SARS | 0 | 0 | 0 | 1 | 2 | 1 | 1 | 1 |
| SART1 | 0 | 0 | 0 | 4 | 4 | 4 | 13 | 10 |
| SART3 | 0 | 0 | 0 | 4 | 10 | 9 | 14 | 17 |
| SBNO1 | 0 | 0 | 0 | 0 | 0 | 1 | 3 | 1 |
| SCAF11 | 0 | 0 | 0 | 1 | 0 | 0 | 1 | 1 |
| SCAF4 | 0 | 0 | 0 | 1 | 0 | 0 | 3 | 3 |
| SCAMP1 | 0 | 0 | 0 | 0 | 0 | 0 | 1 | 1 |
| SCAMP3 | 0 | 0 | 0 | 0 | 0 | 0 | 1 | 0 |
| SCFD2 | 0 | 0 | 0 | 1 | 2 | 0 | 0 | 0 |
| SCML2 | 0 | 0 | 0 | 0 | 0 | 0 | 10 | 3 |
| SCRIB | 0 | 0 | 0 | 2 | 2 | 0 | 1 | 2 |
| SCYL3 | 0 | 0 | 0 | 1 | 1 | 2 | 1 | 1 |
| SDCCAG3 | 0 | 0 | 0 | 0 | 0 | 0 | 1 | 1 |
| SEC24D | 0 | 0 | 0 | 0 | 0 | 1 | 0 | 0 |
| SENP3 | 0 | 0 | 0 | 0 | 0 | 0 | 1 | 1 |
| SENP6 | 0 | 0 | 0 | 0 | 0 | 1 | 3 | 2 |
| SET;SETSIP | 0 | 0 | 0 | 4 | 3 | 1 | 4 | 1 |
| SETD2 | 0 | 0 | 0 | 0 | 0 | 0 | 1 | 0 |
| SF3B1 | 0 | 0 | 0 | 9 | 17 | 14 | 34 | 32 |
| SF3B3 | 0 | 0 | 0 | 6 | 11 | 7 | 22 | 20 |
| SF3B5 | 0 | 0 | 0 | 0 | 0 | 0 | 1 | 2 |
| SFSWAP | 0 | 0 | 0 | 0 | 0 | 0 | 0 | 1 |
| SH3BP4 | 0 | 0 | 0 | 0 | 0 | 0 | 2 | 0 |
| SH3PXD2A;SH3PXD2B | 0 | 0 | 0 | 0 | 0 | 0 | 1 | 0 |
| SHCBP1 | 0 | 0 | 0 | 0 | 0 | 0 | 1 | 0 |
| SHISA2 | 0 | 0 | 0 | 0 | 0 | 0 | 1 | 2 |
| SHROOM2 | 0 | 0 | 0 | 0 | 0 | 2 | 2 | 1 |
| SHROOM3 | 0 | 0 | 0 | 1 | 2 | 2 | 1 | 3 |
| SIN3A | 0 | 0 | 0 | 1 | 1 | 0 | 4 | 1 |
| SIPA1L1 | 0 | 0 | 0 | 0 | 0 | 0 | 0 | 5 |
| SIX4 | 0 | 0 | 0 | 0 | 0 | 0 | 4 | 1 |
| SKA2 | 0 | 0 | 0 | 0 | 0 | 0 | 1 | 0 |
| SKIV2L | 0 | 0 | 0 | 1 | 1 | 1 | 0 | 0 |
| SKIV2L2 | 0 | 0 | 0 | 0 | 1 | 0 | 5 | 1 |
| SKP1 | 0 | 0 | 0 | 0 | 0 | 0 | 1 | 0 |
| SLC12A4;SLC12A6 | 0 | 0 | 0 | 0 | 0 | 0 | 1 | 0 |
| SLC25A1 | 0 | 0 | 0 | 0 | 0 | 0 | 0 | 1 |
| SLC25A13;SLC25A12 | 0 | 0 | 0 | 0 | 0 | 0 | 0 | 1 |
| SLC30A1 | 0 | 0 | 0 | 1 | 1 | 0 | 2 | 1 |

| | | | | | | | | |
|---|---|---|---|---|---|---|---|---|
| SLC35B2 | 0 | 0 | 0 | 0 | 0 | 0 | 1 | 2 |
| SLC4A1AP | 0 | 0 | 0 | 0 | 1 | 0 | 2 | 3 |
| SLC4A7 | 0 | 0 | 0 | 0 | 0 | 0 | 2 | 0 |
| SLC9A3R2 | 0 | 0 | 0 | 1 | 1 | 1 | 3 | 2 |
| SLTM | 0 | 0 | 0 | 0 | 0 | 0 | 1 | 0 |
| SLU7 | 0 | 0 | 0 | 0 | 0 | 0 | 1 | 1 |
| SMARCA1 | 0 | 0 | 0 | 0 | 0 | 0 | 2 | 1 |
| SMARCA4;SMARCA2 | 0 | 0 | 0 | 0 | 0 | 0 | 1 | 0 |
| SMARCA5 | 0 | 0 | 0 | 8 | 7 | 4 | 21 | 20 |
| SMARCC2 | 0 | 0 | 0 | 0 | 0 | 2 | 0 | 2 |
| SMARCE1 | 0 | 0 | 0 | 2 | 0 | 0 | 0 | 2 |
| SMCHD1 | 0 | 0 | 0 | 1 | 0 | 0 | 2 | 1 |
| SMEK1 | 0 | 0 | 0 | 1 | 1 | 1 | 6 | 5 |
| SMG8 | 0 | 0 | 0 | 0 | 0 | 0 | 1 | 0 |
| SMG9 | 0 | 0 | 0 | 1 | 0 | 0 | 0 | 0 |
| SMPD4 | 0 | 0 | 0 | 0 | 1 | 1 | 2 | 0 |
| SMTN | 0 | 0 | 0 | 0 | 0 | 0 | 1 | 1 |
| SNAP23 | 0 | 0 | 0 | 0 | 0 | 0 | 0 | 2 |
| SNAPIN | 0 | 0 | 0 | 0 | 1 | 0 | 0 | 0 |
| SNRPA | 0 | 0 | 0 | 0 | 0 | 0 | 1 | 1 |
| SNRPA1 | 0 | 0 | 0 | 3 | 6 | 5 | 6 | 8 |
| SNRPC | 0 | 0 | 0 | 0 | 0 | 0 | 1 | 1 |
| SNRPD1 | 0 | 0 | 0 | 0 | 0 | 0 | 1 | 0 |
| SNRPN;SNRPB | 0 | 0 | 0 | 0 | 0 | 0 | 1 | 2 |
| SNTB1 | 0 | 0 | 0 | 1 | 0 | 0 | 2 | 0 |
| SNTB2 | 0 | 0 | 0 | 1 | 1 | 2 | 1 | 2 |
| SNW1 | 0 | 0 | 0 | 0 | 4 | 4 | 7 | 5 |
| SOAT1 | 0 | 0 | 0 | 0 | 1 | 0 | 2 | 1 |
| SON | 0 | 0 | 0 | 0 | 0 | 0 | 2 | 3 |
| SOX9;SOX8 | 0 | 0 | 0 | 0 | 0 | 2 | 3 | 1 |
| SP100 | 0 | 0 | 0 | 1 | 0 | 0 | 1 | 0 |
| SPACIA2;FAM122B | 0 | 0 | 0 | 0 | 0 | 1 | 0 | 0 |
| SPAG1 | 0 | 0 | 0 | 1 | 0 | 1 | 0 | 0 |
| SPAG7 | 0 | 0 | 0 | 0 | 0 | 0 | 1 | 0 |
| SPATA5 | 0 | 0 | 0 | 0 | 0 | 0 | 1 | 0 |
| SPCS3 | 0 | 0 | 0 | 0 | 0 | 0 | 1 | 1 |
| SPDL1 | 0 | 0 | 0 | 0 | 0 | 1 | 2 | 2 |
| SPECC1L | 0 | 0 | 0 | 2 | 0 | 1 | 0 | 2 |
| SPEN | 0 | 0 | 0 | 0 | 0 | 0 | 2 | 0 |
| SRBD1 | 0 | 0 | 0 | 0 | 0 | 0 | 1 | 0 |
| SREK1 | 0 | 0 | 0 | 0 | 0 | 0 | 1 | 0 |
| SRP14 | 0 | 0 | 0 | 0 | 2 | 1 | 1 | 0 |
| SRP72 | 0 | 0 | 0 | 3 | 5 | 5 | 1 | 1 |
| SRPK1 | 0 | 0 | 0 | 1 | 0 | 0 | 0 | 1 |
| SRRM1 | 0 | 0 | 0 | 0 | 0 | 0 | 2 | 0 |

| | | | | | | | | |
|---|---|---|---|---|---|---|---|---|
| SRRM2 | 0 | 0 | 0 | 0 | 0 | 0 | 3 | 7 |
| SRSF11 | 0 | 0 | 0 | 2 | 7 | 3 | 4 | 6 |
| SRSF3 | 0 | 0 | 0 | 2 | 2 | 0 | 0 | 0 |
| SRSF7 | 0 | 0 | 0 | 0 | 0 | 0 | 2 | 2 |
| SSB | 0 | 0 | 0 | 1 | 3 | 1 | 3 | 4 |
| SSH3 | 0 | 0 | 0 | 0 | 1 | 1 | 0 | 1 |
| SSRP1 | 0 | 0 | 0 | 1 | 0 | 0 | 3 | 0 |
| STAG1 | 0 | 0 | 0 | 0 | 0 | 0 | 1 | 1 |
| STAMBP | 0 | 0 | 0 | 0 | 0 | 0 | 1 | 0 |
| STAT6 | 0 | 0 | 0 | 1 | 1 | 1 | 1 | 0 |
| STIM2 | 0 | 0 | 0 | 1 | 0 | 0 | 1 | 1 |
| STK3;STK4 | 0 | 0 | 0 | 0 | 0 | 0 | 1 | 0 |
| STK39 | 0 | 0 | 0 | 1 | 0 | 2 | 2 | 1 |
| STRN | 0 | 0 | 0 | 0 | 0 | 0 | 1 | 1 |
| STRN3 | 0 | 0 | 0 | 0 | 0 | 1 | 1 | 0 |
| STXBP4 | 0 | 0 | 0 | 2 | 0 | 0 | 0 | 0 |
| SUB1 | 0 | 0 | 0 | 2 | 0 | 2 | 0 | 0 |
| SUGP1 | 0 | 0 | 0 | 4 | 5 | 1 | 12 | 6 |
| SUGP2 | 0 | 0 | 0 | 1 | 2 | 1 | 7 | 5 |
| SUMO1 | 0 | 0 | 0 | 0 | 2 | 0 | 0 | 0 |
| SUPT16H | 0 | 0 | 0 | 0 | 0 | 0 | 0 | 2 |
| SYDE1 | 0 | 0 | 0 | 1 | 1 | 1 | 1 | 0 |
| TAF6 | 0 | 0 | 0 | 1 | 0 | 0 | 2 | 1 |
| TAF7 | 0 | 0 | 0 | 0 | 0 | 2 | 2 | 0 |
| TAF9B;TAF9 | 0 | 0 | 0 | 0 | 0 | 0 | 1 | 1 |
| TANGO6 | 0 | 0 | 0 | 2 | 0 | 0 | 1 | 4 |
| TAOK3;TAOK1;TAOK2 | 0 | 0 | 0 | 0 | 0 | 0 | 1 | 2 |
| TARS | 0 | 0 | 0 | 1 | 1 | 2 | 2 | 1 |
| TBC1D10B | 0 | 0 | 0 | 0 | 0 | 0 | 1 | 1 |
| TBC1D24 | 0 | 0 | 0 | 0 | 0 | 0 | 1 | 0 |
| TBCE | 0 | 0 | 0 | 2 | 1 | 1 | 0 | 0 |
| TBCEL | 0 | 0 | 0 | 0 | 0 | 0 | 0 | 1 |
| TCEB2 | 0 | 0 | 0 | 0 | 0 | 0 | 2 | 1 |
| TCEB3 | 0 | 0 | 0 | 1 | 2 | 2 | 6 | 4 |
| TCERG1 | 0 | 0 | 0 | 4 | 5 | 6 | 11 | 7 |
| TCOF1 | 0 | 0 | 0 | 3 | 6 | 1 | 22 | 20 |
| TES | 0 | 0 | 0 | 0 | 0 | 0 | 1 | 2 |
| TEX10 | 0 | 0 | 0 | 0 | 0 | 0 | 1 | 0 |
| TEX264 | 0 | 0 | 0 | 0 | 0 | 0 | 1 | 2 |
| TFE3;FFAR4 | 0 | 0 | 0 | 0 | 0 | 0 | 2 | 0 |
| TFIP11 | 0 | 0 | 0 | 0 | 0 | 0 | 1 | 2 |
| THOC1 | 0 | 0 | 0 | 0 | 0 | 0 | 3 | 1 |
| THOC2 | 0 | 0 | 0 | 2 | 4 | 4 | 7 | 6 |
| THOC3 | 0 | 0 | 0 | 0 | 0 | 0 | 0 | 1 |
| THOC5 | 0 | 0 | 0 | 0 | 0 | 0 | 2 | 0 |

| | | | | | | | | |
|---|---|---|---|---|---|---|---|---|
| THOC6 | 0 | 0 | 0 | 0 | 0 | 0 | 1 | 0 |
| THOC7;NIF3L1BP1 | 0 | 0 | 0 | 0 | 0 | 0 | 1 | 1 |
| THRAP3 | 0 | 0 | 0 | 7 | 6 | 8 | 24 | 22 |
| TLE3 | 0 | 0 | 0 | 0 | 0 | 0 | 2 | 2 |
| TLN2 | 0 | 0 | 0 | 0 | 0 | 0 | 0 | 1 |
| TMEM199 | 0 | 0 | 0 | 0 | 0 | 0 | 1 | 1 |
| TMEM256-PLSCR3 | 0 | 0 | 0 | 0 | 1 | 0 | 0 | 0 |
| TMEM57 | 0 | 0 | 0 | 0 | 0 | 0 | 1 | 2 |
| TMSB10 | 0 | 0 | 0 | 0 | 3 | 1 | 0 | 2 |
| TNFRSF11A | 0 | 0 | 0 | 0 | 0 | 0 | 1 | 0 |
| TNIK | 0 | 0 | 0 | 1 | 1 | 1 | 0 | 1 |
| TOLLIP | 0 | 0 | 0 | 0 | 0 | 1 | 0 | 0 |
| TOM1L2 | 0 | 0 | 0 | 0 | 0 | 1 | 1 | 0 |
| TOMM34 | 0 | 0 | 0 | 0 | 0 | 0 | 1 | 0 |
| TOP3B | 0 | 0 | 0 | 0 | 0 | 0 | 1 | 1 |
| TOR1AIP1 | 0 | 0 | 0 | 6 | 1 | 4 | 9 | 10 |
| TPX2 | 0 | 0 | 0 | 5 | 3 | 7 | 11 | 14 |
| TRAPPC5 | 0 | 0 | 0 | 0 | 0 | 2 | 0 | 0 |
| TRIM38 | 0 | 0 | 0 | 0 | 0 | 0 | 0 | 1 |
| TRIP10 | 0 | 0 | 0 | 2 | 1 | 0 | 0 | 0 |
| TRMT112 | 0 | 0 | 0 | 0 | 1 | 0 | 1 | 0 |
| TRMT5 | 0 | 0 | 0 | 0 | 0 | 0 | 0 | 2 |
| TRMT6 | 0 | 0 | 0 | 0 | 0 | 1 | 2 | 2 |
| TRMT61A | 0 | 0 | 0 | 0 | 0 | 0 | 1 | 1 |
| TRPC5 | 0 | 0 | 0 | 0 | 0 | 0 | 0 | 2 |
| TSNAX;DISC1 | 0 | 0 | 0 | 0 | 0 | 0 | 1 | 1 |
| TTC1 | 0 | 0 | 0 | 1 | 0 | 0 | 0 | 1 |
| TUBG1;TUBG2 | 0 | 0 | 0 | 0 | 1 | 1 | 0 | 0 |
| TXLNG | 0 | 0 | 0 | 0 | 0 | 0 | 1 | 0 |
| TXNL1 | 0 | 0 | 0 | 0 | 0 | 1 | 1 | 1 |
| U2AF1;U2AF1L4 | 0 | 0 | 0 | 0 | 0 | 0 | 1 | 0 |
| U2AF2 | 0 | 0 | 0 | 1 | 0 | 0 | 1 | 0 |
| U2SURP | 0 | 0 | 0 | 16 | 12 | 16 | 24 | 17 |
| UBA1 | 0 | 0 | 0 | 2 | 0 | 1 | 4 | 2 |
| UBAP1 | 0 | 0 | 0 | 0 | 1 | 1 | 1 | 1 |
| UBFD1 | 0 | 0 | 0 | 0 | 0 | 0 | 1 | 1 |
| UBIAD1 | 0 | 0 | 0 | 0 | 2 | 1 | 0 | 1 |
| UBTF | 0 | 0 | 0 | 0 | 0 | 0 | 1 | 0 |
| UBXN4 | 0 | 0 | 0 | 0 | 0 | 0 | 1 | 2 |
| UBXN7 | 0 | 0 | 0 | 0 | 0 | 0 | 0 | 1 |
| UHRF1BP1L | 0 | 0 | 0 | 0 | 0 | 1 | 0 | 1 |
| UIMC1 | 0 | 0 | 0 | 0 | 0 | 0 | 2 | 1 |
| USP16 | 0 | 0 | 0 | 0 | 1 | 0 | 3 | 0 |
| USP24 | 0 | 0 | 0 | 0 | 0 | 1 | 1 | 0 |
| USP28 | 0 | 0 | 0 | 0 | 1 | 0 | 1 | 0 |

| | | | | | | | | |
|---|---|---|---|---|---|---|---|---|
| USP8 | 0 | 0 | 0 | 0 | 0 | 0 | 1 | 0 |
| UTP14A;UTP14C | 0 | 0 | 0 | 2 | 0 | 0 | 0 | 0 |
| VAMP5 | 0 | 0 | 0 | 0 | 0 | 0 | 1 | 2 |
| VANGL1 | 0 | 0 | 0 | 0 | 0 | 0 | 5 | 2 |
| VAPB | 0 | 0 | 0 | 0 | 5 | 4 | 8 | 5 |
| VKORC1 | 0 | 0 | 0 | 0 | 0 | 0 | 0 | 1 |
| VPS25 | 0 | 0 | 0 | 0 | 1 | 1 | 0 | 0 |
| VPS37B | 0 | 0 | 0 | 0 | 0 | 0 | 1 | 0 |
| VPS45 | 0 | 0 | 0 | 0 | 1 | 0 | 0 | 0 |
| VPS51 | 0 | 0 | 0 | 0 | 0 | 0 | 1 | 0 |
| VPS72 | 0 | 0 | 0 | 0 | 0 | 0 | 1 | 1 |
| VRK2 | 0 | 0 | 0 | 0 | 1 | 1 | 3 | 3 |
| WAPAL | 0 | 0 | 0 | 11 | 11 | 11 | 16 | 14 |
| WASH6P | 0 | 0 | 0 | 0 | 0 | 0 | 1 | 0 |
| WBP11 | 0 | 0 | 0 | 0 | 0 | 2 | 0 | 0 |
| WDHD1 | 0 | 0 | 0 | 4 | 5 | 2 | 12 | 14 |
| WDR33 | 0 | 0 | 0 | 0 | 0 | 0 | 2 | 1 |
| WDR43 | 0 | 0 | 0 | 0 | 0 | 1 | 3 | 0 |
| WDR70 | 0 | 0 | 0 | 0 | 2 | 0 | 5 | 7 |
| WDR82 | 0 | 0 | 0 | 1 | 2 | 1 | 5 | 5 |
| WHSC1 | 0 | 0 | 0 | 0 | 0 | 0 | 1 | 0 |
| WHSC1L1 | 0 | 0 | 0 | 0 | 0 | 0 | 0 | 1 |
| WIZ | 0 | 0 | 0 | 0 | 0 | 0 | 1 | 1 |
| WRNIP1 | 0 | 0 | 0 | 0 | 0 | 0 | 1 | 0 |
| WTAP | 0 | 0 | 0 | 0 | 0 | 0 | 0 | 1 |
| XAB2 | 0 | 0 | 0 | 1 | 0 | 0 | 5 | 7 |
| XPOT | 0 | 0 | 0 | 0 | 0 | 0 | 1 | 0 |
| XRCC1 | 0 | 0 | 0 | 2 | 1 | 1 | 4 | 4 |
| XRCC4 | 0 | 0 | 0 | 0 | 0 | 0 | 1 | 1 |
| YKT6 | 0 | 0 | 0 | 3 | 2 | 5 | 6 | 6 |
| YLPM1 | 0 | 0 | 0 | 5 | 5 | 8 | 18 | 28 |
| YTHDF1 | 0 | 0 | 0 | 0 | 0 | 0 | 2 | 0 |
| ZBTB21 | 0 | 0 | 0 | 0 | 0 | 0 | 1 | 0 |
| ZBTB9 | 0 | 0 | 0 | 0 | 0 | 0 | 0 | 1 |
| ZC3H11A | 0 | 0 | 0 | 3 | 2 | 1 | 8 | 6 |
| ZC3H18 | 0 | 0 | 0 | 0 | 0 | 0 | 2 | 2 |
| ZC3H4 | 0 | 0 | 0 | 3 | 5 | 5 | 9 | 8 |
| ZC3HC1 | 0 | 0 | 0 | 0 | 0 | 0 | 0 | 2 |
| ZFP91 | 0 | 0 | 0 | 1 | 2 | 1 | 0 | 0 |
| ZFR | 0 | 0 | 0 | 0 | 0 | 0 | 5 | 3 |
| ZFYVE9 | 0 | 0 | 0 | 0 | 2 | 0 | 1 | 0 |
| ZHX3 | 0 | 0 | 0 | 4 | 1 | 3 | 13 | 12 |
| ZMYM1 | 0 | 0 | 0 | 0 | 0 | 0 | 1 | 1 |
| ZMYND8 | 0 | 0 | 0 | 0 | 0 | 1 | 12 | 12 |
| ZNF131 | 0 | 0 | 0 | 0 | 0 | 0 | 1 | 2 |

| | | | | | | | | |
|---|---|---|---|---|---|---|---|---|
| ZNF148 | 0 | 0 | 0 | 1 | 1 | 1 | 10 | 6 |
| ZNF207 | 0 | 0 | 0 | 0 | 1 | 0 | 1 | 0 |
| ZNF280C | 0 | 0 | 0 | 0 | 0 | 0 | 2 | 1 |
| ZNF281 | 0 | 0 | 0 | 1 | 2 | 0 | 5 | 4 |
| ZNF318 | 0 | 0 | 0 | 0 | 0 | 0 | 10 | 6 |
| ZNF451 | 0 | 0 | 0 | 0 | 0 | 0 | 1 | 3 |
| ZNF512 | 0 | 0 | 0 | 0 | 0 | 0 | 2 | 2 |
| ZNF592 | 0 | 0 | 0 | 0 | 0 | 0 | 1 | 1 |
| ZNF598 | 0 | 0 | 0 | 0 | 0 | 0 | 1 | 1 |
| ZNF609 | 0 | 0 | 0 | 0 | 0 | 0 | 1 | 1 |
| ZNF638 | 0 | 0 | 0 | 1 | 1 | 0 | 11 | 4 |
| ZRANB2 | 0 | 0 | 0 | 0 | 0 | 0 | 2 | 3 |
| ZWILCH | 0 | 0 | 0 | 0 | 0 | 1 | 0 | 0 |
