## Supplementary material for "OPTN translocates to an ATG9A-positive compartment to regulate innate immune signalling and cytokine secretion": Figure Legends for Supplemental Figures

**SI Figure Legends**

**Figure S1 –** **RPE cells show a robust TLR3/RIG-I response.**

(A) CXCL8 secretion from RPE cells stimulated with the indicated ligands. Bars depict mean of ≥n=4 independent experiments ±SEM. Statistical significance was calculated by one-way ANOVA and a Bonferroni post-hoc test. *** = p<0.001. (B) Volcano plot of fold change versus adjusted p value from SILAC secretome experiments (n=3). Red points are p value significant (p<0.05) in the poly(I:C) stimulated versus unstimulated condition. Significantly enriched and other notable proteins are labelled. (C) Poly(I:C)-induced CXCL8 secretion (i), IL6 secretion (ii), NF-κB-luciferase (iii), IFNα/β secretion (iv) and IFN-β mRNA expression in RPE cells over the time courses indicated. Bars depict mean of n=4 (i, ii & iv) or n=3 (iii & v) experiments ±SEM. (E) Immunoblot analysis of lysates from RPE cells stimulated with poly(I:C) for the indicated times and probed with p-p65(Ser536), p-IRF3 and vinculin (loading control) antibodies.

**Figure S2 – Composition of OPTN foci**.

(A) Confocal microscope images of RPE cells stably expressing GFP-OPTN (green) and treated with 2’,3’-cGAMP, LPS and Pam3CSK4 for 24 hours. Cells were stained with Hoechst to label DNA (blue). Scale bar, 20 µm. (B) Confocal microscope images of RPE cells stably expressing GFP-OPTN (green) and treated with poly(I:C) for 24 hours. Cells were immunostained with antibodies (red) against EEA1 (i), LAMP1 (ii), LC3 (iii) and CIMPR (iv) and Hoechst was used to visual DNA (blue). Scale bars, 20 µm. Graphs depict pixel intensity in green (OPTN) and red (EEA1, LAMP1, LC3 and CIMPR) channels along line profiles highlighted in image. (C) Confocal microscope images of RPE cells stably expressing GFP-OPTN (green) and treated with poly(I:C) for 24 hours. Cells were immunostained with a MYO6 antibody (red) and Hoechst was used to visual DNA (blue). Scale bar, 20 µm. Graph depicts Pearson’s correlation coefficient calculated for GFP-OPTN versus MYO6 after treatment with poly(I:C) for 0 and 24 hours. Bars represent the mean of n=3 independent experiments ±SEM. Cells were quantified from ≥20 randomly selected fields of view (1 cell/image). Statistical significance was calculated using a two-sample t-test. ** = p<0.01. (D) Confocal microscope images of RPE cells stably expressing GFP-OPTN (green) and treated with mock (upper panels) or MYO6 (lower panels) siRNA were stimulated with vehicle (left column) or poly(I:C) (right column). DNA was labelled with Hoechst (blue). Scale bar, 20 µm.

**Figure S3 – TLR3 knockdown perturbs foci formation.**

(A) Confocal microscope images of RPE cells stably expressing mCherry-OPTN (red) and TLR3-CFP treated with poly(I:C) for 0 hours or 24 hours. Cells were immunostained with a GFP antibody to detect TLR3-CFP (green) and Hoechst to label DNA (blue). Scale bars, 20 µm. (B) Graph depicting relative TLR3 mRNA expression in RPE dCas9-KRAB cells expressing a non-targeting sgRNA (GAL4) or a sgRNA targeting TLR3. Bar represents the mean from experiments with two different qPCR primers ±SEM. (C) Widefield microscope images of RPE dCas9-KRAB cells stably expressing GFP-OPTN and GAL4 or TLR3 sgRNAs. Cells were treated for 0 hours or 24 hours with poly(I:C) and stained with Hoechst to label DNA (blue). Scale bar, 20 µm. (D) Percentage of GAL4 or TLR3 sgRNA-expressing GFP-OPTN cells containing foci after poly(I:C) treatment. Cells were manually counted from ≥5 randomly selected fields of view across n=3 independent experiments ±SEM. Statistical significance was determined by repeated measures ANOVA and Bonferroni post-hoc test. ** = p<0.01.

**Figure S4 – TBK1 inhibition perturbs foci formation.**

(A) Left, Immunoblot analysis of lysates from RPE cells stimulated with poly(I:C) for the indicated times and probed with p-TBK1 and vinculin (loading control) antibodies. Right, graph depicting gel band density analysis for p-TBK1. Points represent mean of n=3 experiments and error bars indicate SEM. (B) Confocal microscope images of RPE cells stably expressing GFP-OPTN (green) and treated with vehicle (top row) or poly(I:C) for 24 hours (bottom row). Cells were simultaneously treated with DMSO or BX795 for 16 hours (added after 8 hours) or 24 hours (added after 0 hours). DNA was visualised with Hoechst (blue). Scale bar, 20 µm. (C) Relative foci counts/GFP-OPTN cell after treatment with poly(I:C) for 24 hours combined with BX795 addition after the indicated times. Points represent mean of n=3 independent experiments ±SEM.

**Figure S5 – Characterisation of OPTN BioID cells.**

(A) Confocal microscope images of RPE cells stably expressing myc-BirA*-OPTN (upper panels) and OPTN-BirA*-HA (lower panels). Cells were immunostained with an anti-myc antibody (green), biotin was visualised with fluorescently-labelled streptavidin (red) and DNA with Hoechst (blue). Scale bar, 20 µm. (B) Immunoblot analysis of lysates from myc-BirA*-OPTN or OPTN-BirA*-HA RPE cells probed with myc or HA antibodies respectively.

**Figure S6 – OPTN foci are ubiquitinated but don’t require HOIP activity.**

(A) Confocal microscope images of RPE cells stably expressing GFP-OPTN (green) and RFP-UBAN (top) and RFP-UBAN F312A (bottom; red) and stimulated with poly(I:C) for 24 hours. Scale bar, 20 µm. (B) Confocal microscope images of RPE cells stably expressing GFP-OPTN (green) and treated with mock or HOIP siRNA were stimulated with vehicle or poly(I:C). DNA was labelled with Hoechst (blue). Scale bar, 20 µm. Mock-treated condition same as shown in Figure S1. (C) Confocal microscope images of RPE cells stably expressing GFP-OPTN (green) and transiently transfected with HA-Ub K63. Cells were treated with poly(I:C) for 24 hours and immunostained with a HA antibody (red) and Hoechst to label DNA (blue). Scale bar, 20 µm.

**Figure S7 – OPTN mutants also modulate RIG-I-induced cytokine secretion.**

(A-B) CXCL8 (A) and IL6 (B) secretion from RPE cells expressing GFP-OPTN wild-type, E50K and E478G and stimulated with poly(I:C) as indicated. Graphs depicts mean of n=3 independent experiments ±SEM. Statistical significance was calculated by one-way ANOVA and a Bonferroni post-hoc test. * = p<0.05.
