## Supplemental Table 1 for "OPTN translocates to an ATG9A-positive compartment to regulate innate immune signalling and cytokine secretion"

RPE1 secretome limma output

| Gene name | Fold change (log) | Fold change (log)*-1 | t | P value | adj. P value | adj. P value (log10) | B |
|---|---|---|---|---|---|---|---|
| TFPI2 | -4.634686667 | 4.634686667 | -14.40819034 | 0.0000452 | 0.00670112 | 2.17385262 | 2.906154396 |
| C1R | -4.092236667 | 4.092236667 | -14.10948173 | 0.0000498 | 0.00670112 | 2.17385262 | 2.826583128 |
| PTX3 | -4.546486667 | 4.546486667 | -13.92628646 | 0.0000529 | 0.00670112 | 2.17385262 | 2.776461507 |
| C1S | -3.740626667 | 3.740626667 | -12.3099374 | 0.0000933 | 0.008860572 | 2.052538233 | 2.286090695 |
| KRT1 | -3.460836667 | 3.460836667 | -11.19368157 | 0.000144024 | 0.010082295 | 1.996440609 | 1.889145311 |
| LGMN | -4.017906667 | 4.017906667 | -10.95027764 | 0.000159194 | 0.010082295 | 1.996440609 | 1.795260178 |
| LGALS3BP | -2.76477 | 2.76477 | -9.627600706 | 0.000285285 | 0.012349583 | 1.908347711 | 1.232176971 |
| CXCL8 | -6.841985 | 6.841985 | -13.47911166 | 0.000278398 | 0.012349583 | 1.908347711 | 1.190146702 |
| HLA-C | -2.720053333 | 2.720053333 | -9.400770425 | 0.000317635 | 0.012349583 | 1.908347711 | 1.125706602 |
| PLOD2 | -3.055633333 | 3.055633333 | -9.35307611 | 0.000324989 | 0.012349583 | 1.908347711 | 1.102916623 |
| IGFBP3 | -2.368876667 | 2.368876667 | -8.770633945 | 0.000433636 | 0.01498015 | 1.824483845 | 0.812729348 |
| HHIP | -2.46779 | 2.46779 | -7.203004916 | 0.001036189 | 0.029478188 | 1.530499221 | -0.093111001 |
| AGRN | -2.06159 | 2.06159 | -7.17737918 | 0.001052448 | 0.029478188 | 1.530499221 | -0.109645734 |
| C3 | -2.863896667 | 2.863896667 | -7.125907313 | 0.001086038 | 0.029478188 | 1.530499221 | -0.14304327 |
| SEMA3C | -2.331746667 | 2.331746667 | -6.75288451 | 0.001371893 | 0.034754627 | 1.458987374 | -0.392715968 |
| ESM1 | -4.074055 | 4.074055 | -7.887686022 | 0.00191182 | 0.040360655 | 1.394041795 | -0.45190821 |
| INHBA | -2.169486667 | 2.169486667 | -6.383322929 | 0.001748312 | 0.040360655 | 1.394041795 | -0.654012738 |
| TMEM132A | -2.77205 | 2.77205 | -7.375770336 | 0.002421159 | 0.041738359 | 1.379464632 | -0.686424608 |
| PRSS23 | -2.493846667 | 2.493846667 | -6.295801302 | 0.001854792 | 0.040360655 | 1.394041795 | -0.718039422 |
| CPA4 | 1.630776667 | -1.630776667 | 6.033749606 | 0.002223105 | 0.041738359 | 1.379464632 | -0.914892117 |
| IL11 | -3.55853 | 3.55853 | -6.783260188 | 0.00324354 | 0.045649829 | 1.340560843 | -0.984467537 |
| CTSL | -2.112873333 | 2.112873333 | -5.911664794 | 0.002424134 | 0.041738359 | 1.379464632 | -1.009320055 |
| PTN | -2.65815 | 2.65815 | -5.884847957 | 0.002471145 | 0.041738359 | 1.379464632 | -1.030299543 |
| GDF6 | 1.69568 | -1.69568 | 5.836419337 | 0.00255882 | 0.041738359 | 1.379464632 | -1.068406176 |
| RPS27 | 1.784846667 | -1.784846667 | 5.795336529 | 0.002636107 | 0.041738359 | 1.379464632 | -1.100956061 |
| SERPINE1 | -1.874283333 | 1.874283333 | -5.717258332 | 0.002790778 | 0.042419828 | 1.372431094 | -1.163388705 |
| CTSS | -3.476296667 | 3.476296667 | -5.557755625 | 0.00314159 | 0.045649829 | 1.340560843 | -1.293298891 |
| CST1 | -2.092333333 | 2.092333333 | -5.319159266 | 0.003769242 | 0.048923528 | 1.31048223 | -1.493749548 |
| B2M | -1.9026 | 1.9026 | -5.293652948 | 0.003844768 | 0.048923528 | 1.31048223 | -1.515623781 |
| YWHAH | 2.224176667 | -2.224176667 | 5.287789068 | 0.003862384 | 0.048923528 | 1.31048223 | -1.520665025 |
| SFRP1 | -1.423446667 | 1.423446667 | -5.085192555 | 0.004534071 | 0.05557893 | 1.255089819 | -1.69772167 |
| RNH1 | 2.86104 | -2.86104 | 5.299548077 | 0.007534575 | 0.069408814 | 1.158585374 | -1.882381104 |
| B4GALT1 | -1.69823 | 1.69823 | -4.868998172 | 0.005408794 | 0.063009319 | 1.200595216 | -1.892998558 |
| GREM1 | -1.6954 | 1.6954 | -4.855029254 | 0.005471862 | 0.063009319 | 1.200595216 | -1.905846618 |
| CTSC | -2.001683333 | 2.001683333 | -4.700573615 | 0.006230299 | 0.068384835 | 1.1650402 | -2.049815219 |
| CDH6 | 1.470146333 | -1.470146333 | 4.656324603 | 0.00646995 | 0.068384835 | 1.1650402 | -2.091711496 |
| IGFBP6 | -1.533466667 | 1.533466667 | -4.654770226 | 0.006478563 | 0.068384835 | 1.1650402 | -2.093188558 |
| SDC4 | -1.570851 | 1.570851 | -4.552406823 | 0.00707669 | 0.069408814 | 1.158585374 | -2.19126278 |
| RPS11 | 1.35936 | -1.35936 | 4.520260465 | 0.007277753 | 0.069408814 | 1.158585374 | -2.222390494 |
| CFH | -1.30585 | 1.30585 | -4.503447112 | 0.007385583 | 0.069408814 | 1.158585374 | -2.238734026 |
| RPL7 | 2.79456 | -2.79456 | 4.79401689 | 0.010501586 | 0.079158626 | 1.101501755 | -2.248170613 |
| GAPDH | 2.378166667 | -2.378166667 | 4.474218 | 0.007577535 | 0.069408814 | 1.158585374 | -2.267249673 |
| WNT5A | -1.274774667 | 1.274774667 | -4.460224613 | 0.007671501 | 0.069408814 | 1.158585374 | -2.280948039 |
| FST | -1.577906667 | 1.577906667 | -4.429097239 | 0.007885466 | 0.069685517 | 1.156857472 | -2.311527606 |
| RPL13A | 1.77861 | -1.77861 | 4.695528304 | 0.011238884 | 0.079158626 | 1.101501755 | -2.323544995 |
| RPL18A | 1.845975 | -1.845975 | 4.694251257 | 0.011248857 | 0.079158626 | 1.101501755 | -2.324531518 |

| | | | | | | | |
|---|---|---|---|---|---|---|---|
| RPL8 | 1.71387 | -1.71387 | 4.540240177 | 0.012536047 | 0.085066035 | 1.070243809 | -2.445260298 |
| IQGAP1 | 2.27769 | -2.27769 | 4.509062554 | 0.012818352 | 0.085455677 | 1.068259081 | -2.470129326 |
| CLTC | 2.713026667 | -2.713026667 | 4.203640492 | 0.009662367 | 0.079158626 | 1.101501755 | -2.537533987 |
| GLA | -1.414032333 | 1.414032333 | -4.19255445 | 0.009761189 | 0.079158626 | 1.101501755 | -2.548853962 |
| RPS13 | 1.177195667 | -1.177195667 | 4.173371024 | 0.009934996 | 0.079158626 | 1.101501755 | -2.568488421 |
| IL6 | -4.455563333 | 4.455563333 | -4.165235551 | 0.010009795 | 0.079158626 | 1.101501755 | -2.576832889 |
| RPS26 | 1.424806667 | -1.424806667 | 4.122398877 | 0.010414659 | 0.079158626 | 1.101501755 | -2.620944456 |
| PLOD1 | -1.325910333 | 1.325910333 | -4.087530881 | 0.010758364 | 0.079158626 | 1.101501755 | -2.657067197 |
| RPS18 | 1.164984 | -1.164984 | 4.058995849 | 0.011049512 | 0.079158626 | 1.101501755 | -2.686774422 |
| NUCB1 | -1.0806 | 1.0806 | -4.05303373 | 0.011111499 | 0.079158626 | 1.101501755 | -2.692998013 |
| MDK | -2.30975 | 2.30975 | -4.014867005 | 0.011518047 | 0.079579233 | 1.099200248 | -2.732974359 |
| WARS | -1.707507333 | 1.707507333 | -3.872955486 | 0.013189395 | 0.086413279 | 1.063419513 | -2.883683034 |
| CCT8 | 1.611705 | -1.611705 | 4.004706842 | 0.018688203 | 0.101119138 | 0.995166643 | -2.893417259 |
| DHFRL1 | 1.470275 | -1.470275 | 3.974849008 | 0.019129872 | 0.101119138 | 0.995166643 | -2.919757975 |
| DSC3 | 1.730795 | -1.730795 | 3.972881943 | 0.019159416 | 0.101119138 | 0.995166643 | -2.921498512 |
| RPL11 | 1.198878667 | -1.198878667 | 3.800301709 | 0.014153336 | 0.090219997 | 1.044697191 | -2.962107815 |
| RPS23 | 1.168060667 | -1.168060667 | 3.793674357 | 0.014245263 | 0.090219997 | 1.044697191 | -2.969304406 |
| IPO5 | 1.466290333 | -1.466290333 | 3.765852888 | 0.014638799 | 0.090422292 | 1.043724487 | -2.99959368 |
| TGFBI | -1.207302667 | 1.207302667 | -3.7504527 | 0.014862046 | 0.090422292 | 1.043724487 | -3.016414174 |
| CCT4 | 1.81853 | -1.81853 | 3.859295018 | 0.020964039 | 0.103458892 | 0.985232179 | -3.023099466 |
| PSAP | -1.355236333 | 1.355236333 | -3.731609748 | 0.015140586 | 0.090422292 | 1.043724487 | -3.037047574 |
| RPS9 | 1.015580333 | -1.015580333 | 3.72571123 | 0.015229018 | 0.090422292 | 1.043724487 | -3.043518471 |
| CLIC4 | 1.444125 | -1.444125 | 3.688651032 | 0.015798507 | 0.091455504 | 1.038790151 | -3.084304742 |
| ERAP1 | -0.995081667 | 0.995081667 | -3.683197149 | 0.015884377 | 0.091455504 | 1.038790151 | -3.090325871 |
| ACTR2 | 2.217405 | -2.217405 | 3.73881252 | 0.023110117 | 0.109773056 | 0.959504245 | -3.133246557 |
| FASN | 2.699975 | -2.699975 | 3.684698347 | 0.024160641 | 0.110059167 | 0.958373779 | -3.183525272 |
| HIST1H4A | -1.34279 | 1.34279 | -3.541692439 | 0.018312706 | 0.101119138 | 0.995166643 | -3.248240916 |
| RPL14 | 1.349815 | -1.349815 | 3.594005994 | 0.02605455 | 0.110059167 | 0.958373779 | -3.268921956 |
| VCAN | 1.280333667 | -1.280333667 | 3.522047811 | 0.018682595 | 0.101119138 | 0.995166643 | -3.270420766 |
| RPS17 | 1.27179 | -1.27179 | 3.502802257 | 0.019053338 | 0.101119138 | 0.995166643 | -3.292210575 |
| MMP14 | 1.29924 | -1.29924 | 3.529071815 | 0.027521835 | 0.11208364 | 0.950457774 | -3.330941637 |
| G6PD | 1.8763 | -1.8763 | 3.520371413 | 0.027725953 | 0.11208364 | 0.950457774 | -3.339307387 |
| TFRC | -1.231755333 | 1.231755333 | -3.46060162 | 0.019896317 | 0.103458892 | 0.985232179 | -3.340199343 |
| RPS20 | 1.226564667 | -1.226564667 | 3.447390647 | 0.020168972 | 0.103458892 | 0.985232179 | -3.355281218 |
| FBN1 | 1.104264 | -1.104264 | 3.43357303 | 0.020458752 | 0.103458892 | 0.985232179 | -3.371085634 |
| TXNRD1 | 1.659169 | -1.659169 | 3.41544888 | 0.020846129 | 0.103458892 | 0.985232179 | -3.3918622 |
| CAPZB | 2.68598 | -2.68598 | 3.454847208 | 0.029323494 | 0.115609519 | 0.937006407 | -3.402736648 |
| CCT5 | 1.85742 | -1.85742 | 3.410084738 | 0.030479011 | 0.116990144 | 0.931850726 | -3.44650069 |
| SEMA3A | -0.970692667 | 0.970692667 | -3.324556513 | 0.022920399 | 0.109773056 | 0.959504245 | -3.49684601 |
| PLOD3 | -0.887136667 | 0.887136667 | -3.318876898 | 0.023057675 | 0.109773056 | 0.959504245 | -3.503449609 |
| RPS4X | 1.877746667 | -1.877746667 | 3.30087757 | 0.023498973 | 0.110059167 | 0.958373779 | -3.52441068 |
| RPS16 | 0.904633 | -0.904633 | 3.271609678 | 0.024237343 | 0.110059167 | 0.958373779 | -3.558603001 |
| VASN | 0.884959 | -0.884959 | 3.264301817 | 0.024425821 | 0.110059167 | 0.958373779 | -3.567161316 |
| SDC1 | 1.273425 | -1.273425 | 3.281741555 | 0.034109927 | 0.12205857 | 0.913431723 | -3.573934203 |
| THBS1 | -0.967812 | 0.967812 | -3.249611259 | 0.024809799 | 0.110059167 | 0.958373779 | -3.584390735 |
| CTSB | -1.082736 | 1.082736 | -3.2439139 | 0.024960568 | 0.110059167 | 0.958373779 | -3.591081748 |
| RPS7 | 1.574978 | -1.574978 | 3.217124006 | 0.025683679 | 0.110059167 | 0.958373779 | -3.622611293 |
| YWHAZ | 1.006954333 | -1.006954333 | 3.207867394 | 0.025939069 | 0.110059167 | 0.958373779 | -3.633531303 |

| | | | | | | | |
|---|---|---|---|---|---|---|---|
| CFL1 | 0.931386 | -0.931386 | 3.203282259 | 0.026066645 | 0.110059167 | 0.958373779 | -3.638945257 |
| UCHL1 | 1.110633333 | -1.110633333 | 3.184139302 | 0.026607062 | 0.111106411 | 0.954260882 | -3.661583322 |
| CORO1C | 1.7949235 | -1.7949235 | 3.191841081 | 0.036965447 | 0.12205857 | 0.913431723 | -3.664923447 |
| MYH9 | 2.13583 | -2.13583 | 3.17758576 | 0.037444173 | 0.12205857 | 0.913431723 | -3.679481834 |
| ARPC1B | 1.490551 | -1.490551 | 3.17596337 | 0.037499127 | 0.12205857 | 0.913431723 | -3.681140975 |
| METRNL | -1.18992 | 1.18992 | -3.167229988 | 0.037796607 | 0.12205857 | 0.913431723 | -3.690080117 |
| RPS15 | 1.5792485 | -1.5792485 | 3.164144031 | 0.037902398 | 0.12205857 | 0.913431723 | -3.693241976 |
| HSPA4 | 1.684252667 | -1.684252667 | 3.155088997 | 0.027451743 | 0.11208364 | 0.950457774 | -3.696044321 |
| RPL13 | 1.3354465 | -1.3354465 | 3.143843073 | 0.03860725 | 0.122256291 | 0.912728784 | -3.714083836 |
| HEXB | -0.832624333 | 0.832624333 | -3.117399635 | 0.0285935 | 0.114374 | 0.941672691 | -3.740943493 |
| CCT2 | 1.504504667 | -1.504504667 | 3.088337199 | 0.029510851 | 0.115609519 | 0.937006407 | -3.775710508 |
| RPS5 | 0.957181667 | -0.957181667 | 3.065275066 | 0.03026263 | 0.116990144 | 0.931850726 | -3.803388372 |
| RPS10 | 1.669314667 | -1.669314667 | 3.025184769 | 0.031621802 | 0.120162848 | 0.920229786 | -3.851687561 |
| MMP1 | -1.454415 | 1.454415 | -3.006747364 | 0.04380016 | 0.135317567 | 0.868645819 | -3.856713291 |
| LIF | -1.143131 | 1.143131 | -2.997779327 | 0.044168001 | 0.135353553 | 0.868530341 | -3.866156791 |
| YWHAQ | 1.192433667 | -1.192433667 | 2.991732386 | 0.032809021 | 0.12205857 | 0.913431723 | -3.892167071 |
| RPL24 | 1.2928675 | -1.2928675 | 2.9363556 | 0.046788638 | 0.1399975 | 0.85387972 | -3.931207809 |
| LAMA5 | -0.793196333 | 0.793196333 | -2.957034689 | 0.034094014 | 0.12205857 | 0.913431723 | -3.934321199 |
| RPS8 | 0.831992 | -0.831992 | 2.950110581 | 0.034357195 | 0.12205857 | 0.913431723 | -3.942753463 |
| CST3 | -1.125722333 | 1.125722333 | -2.938091541 | 0.034819499 | 0.12205857 | 0.913431723 | -3.957406195 |
| RPS19 | 1.151870667 | -1.151870667 | 2.934525345 | 0.034958019 | 0.12205857 | 0.913431723 | -3.961757681 |
| PLAT | -0.825849667 | 0.825849667 | -2.931024594 | 0.035094601 | 0.12205857 | 0.913431723 | -3.966031013 |
| SEMA7A | 0.826239 | -0.826239 | 2.921915184 | 0.035452838 | 0.12205857 | 0.913431723 | -3.977158667 |
| KPNB1 | 2.3745765 | -2.3745765 | 2.883296179 | 0.04920219 | 0.144691457 | 0.83955711 | -3.987917335 |
| ALDOA | 1.850655333 | -1.850655333 | 2.908778489 | 0.035976739 | 0.12205857 | 0.913431723 | -3.993225835 |
| SPOCK1 | -0.822472 | 0.822472 | -2.904379565 | 0.036154118 | 0.12205857 | 0.913431723 | -3.998611292 |
| TALDO1 | 1.566405667 | -1.566405667 | 2.89767765 | 0.036426259 | 0.12205857 | 0.913431723 | -4.006821256 |
| CYR61 | -1.314684 | 1.314684 | -2.884251153 | 0.036978436 | 0.12205857 | 0.913431723 | -4.023287132 |
| EFEMP1 | -0.87624 | 0.87624 | -2.880088424 | 0.037151541 | 0.12205857 | 0.913431723 | -4.028397086 |
| APEX1 | 1.085587667 | -1.085587667 | 2.848571697 | 0.038492151 | 0.122256291 | 0.912728784 | -4.067159885 |
| RAN | 0.861196 | -0.861196 | 2.829320893 | 0.03933774 | 0.12354001 | 0.908192367 | -4.090900594 |
| RPS14 | 1.069897 | -1.069897 | 2.813882368 | 0.040030968 | 0.124686621 | 0.904180144 | -4.109974353 |
| PGD | 1.9559815 | -1.9559815 | 2.727934631 | 0.057164571 | 0.155463369 | 0.808371924 | -4.156659805 |
| RAB7A | 1.17578 | -1.17578 | 2.702527703 | 0.058606831 | 0.155463369 | 0.808371924 | -4.184627475 |
| CHID1 | 1.0079705 | -1.0079705 | 2.69020802 | 0.059321549 | 0.155463369 | 0.808371924 | -4.198225603 |
| PKM | 1.645704667 | -1.645704667 | 2.691551671 | 0.046034403 | 0.139944585 | 0.854043903 | -4.262149628 |
| HEXA | -0.730892667 | 0.730892667 | -2.682234262 | 0.046531379 | 0.1399975 | 0.85387972 | -4.273812551 |
| RPS2 | 2.1019875 | -2.1019875 | 2.589730018 | 0.065548494 | 0.167170655 | 0.776839955 | -4.310006888 |
| ARPC5 | 1.109004 | -1.109004 | 2.573242646 | 0.066641791 | 0.168825871 | 0.772561002 | -4.328494657 |
| WDR1 | 1.628744 | -1.628744 | 2.631133875 | 0.049365555 | 0.144691457 | 0.83955711 | -4.337947523 |
| PPIA | 0.777461667 | -0.777461667 | 2.624436714 | 0.049751031 | 0.144691457 | 0.83955711 | -4.346373836 |
| RPS6 | 1.832984 | -1.832984 | 2.622200703 | 0.049880476 | 0.144691457 | 0.83955711 | -4.349188216 |
| NPEPPS | 1.787885 | -1.787885 | 2.520594988 | 0.070278 | 0.176858544 | 0.752373955 | -4.387795312 |
| SERPINH1 | 1.296979 | -1.296979 | 2.498900562 | 0.071843199 | 0.179607999 | 0.745674327 | -4.41234625 |
| LDLR | 1.053672667 | -1.053672667 | 2.571390671 | 0.052925088 | 0.151295131 | 0.820175048 | -4.413278612 |
| UBA1 | 1.599438333 | -1.599438333 | 2.570935213 | 0.052953296 | 0.151295131 | 0.820175048 | -4.413854277 |
| CSPG4 | 0.740682667 | -0.740682667 | 2.561662256 | 0.053531226 | 0.15180497 | 0.818714011 | -4.42557898 |
| CSE1L | 2.1066285 | -2.1066285 | 2.469211268 | 0.074051295 | 0.182111733 | 0.739662074 | -4.446050607 |

| | | | | | | | |
|---|---|---|---|---|---|---|---|
| RPL18 | 0.698942 | -0.698942 | 2.52955227 | 0.055587049 | 0.155463369 | 0.808371924 | -4.466241547 |
| SRGN | -0.898109 | 0.898109 | -2.524680599 | 0.055906513 | 0.155463369 | 0.808371924 | -4.472419093 |
| RPSA | 1.838209333 | -1.838209333 | 2.519627056 | 0.056240049 | 0.155463369 | 0.808371924 | -4.478829515 |
| RNASE4 | -0.988093667 | 0.988093667 | -2.513249684 | 0.056664098 | 0.155463369 | 0.808371924 | -4.486922457 |
| HSPA1B | 1.171011667 | -1.171011667 | 2.496534275 | 0.057792391 | 0.155463369 | 0.808371924 | -4.508151317 |
| APP | 0.694026667 | -0.694026667 | 2.494742945 | 0.057914769 | 0.155463369 | 0.808371924 | -4.510427762 |
| YWHAG | 0.813388 | -0.813388 | 2.481676516 | 0.058816115 | 0.155463369 | 0.808371924 | -4.527040915 |
| ADAM19 | -1.762689667 | 1.762689667 | -2.474511972 | 0.059316891 | 0.155463369 | 0.808371924 | -4.536156195 |
| RPS25 | 0.721460333 | -0.721460333 | 2.465139047 | 0.059979126 | 0.156110054 | 0.806569125 | -4.548087434 |
| CCT7 | 1.8016835 | -1.8016835 | 2.34885841 | 0.08384494 | 0.191184462 | 0.718547406 | -4.583867874 |
| TPM3 | 1.930041 | -1.930041 | 2.341125252 | 0.084523657 | 0.191184462 | 0.718547406 | -4.592784794 |
| NID1 | -0.661117 | 0.661117 | -2.401636806 | 0.064685696 | 0.167170655 | 0.776839955 | -4.629098909 |
| CD44 | 0.645635667 | -0.645635667 | 2.396108236 | 0.065114241 | 0.167170655 | 0.776839955 | -4.636165462 |
| SH3BGRL3 | 1.338535 | -1.338535 | 2.251433184 | 0.092874185 | 0.20760112 | 0.682770308 | -4.69670067 |
| EEF2 | 1.593083667 | -1.593083667 | 2.285056255 | 0.074413688 | 0.182111733 | 0.739662074 | -4.778493554 |
| HIST1H2BK | -1.280677 | 1.280677 | -2.284274468 | 0.074484075 | 0.182111733 | 0.739662074 | -4.779497732 |
| CBR1 | 3.049634 | -3.049634 | 2.175473512 | 0.100688454 | 0.221165389 | 0.655282836 | -4.785352738 |
| PFN1 | 0.812336 | -0.812336 | 2.273894766 | 0.075425475 | 0.182111733 | 0.739662074 | -4.792832489 |
| TARS | 0.734557667 | -0.734557667 | 2.271509083 | 0.075643667 | 0.182111733 | 0.739662074 | -4.795897978 |
| CFL2 | 0.723280667 | -0.723280667 | 2.270674686 | 0.075720141 | 0.182111733 | 0.739662074 | -4.79697019 |
| SPTBN1 | 1.287061 | -1.287061 | 2.260797528 | 0.076631806 | 0.183031986 | 0.737473009 | -4.809664506 |
| APLP2 | 0.639780667 | -0.639780667 | 2.251613272 | 0.077490196 | 0.183031986 | 0.737473009 | -4.821471482 |
| GNB2L1 | 1.187273 | -1.187273 | 2.251001294 | 0.077547762 | 0.183031986 | 0.737473009 | -4.822258325 |
| RPS3 | 1.323552 | -1.323552 | 2.244946283 | 0.078119835 | 0.183244058 | 0.7369701 | -4.830044159 |
| CCT6A | 1.293698 | -1.293698 | 2.135167048 | 0.105136502 | 0.227846611 | 0.642357427 | -4.832603056 |
| FSCN1 | 1.247705333 | -1.247705333 | 2.235250813 | 0.079045389 | 0.184277594 | 0.734527468 | -4.842513531 |
| RPS3A | 1.969818 | -1.969818 | 2.227142522 | 0.079828534 | 0.184968555 | 0.732902097 | -4.852943808 |
| SPTAN1 | 1.339044 | -1.339044 | 2.207029612 | 0.081807571 | 0.188405315 | 0.72490685 | -4.878824126 |
| OAF | -0.7685525 | 0.7685525 | -2.085058127 | 0.11097902 | 0.236921503 | 0.625395521 | -4.891518643 |
| GAS6 | -0.714101667 | 0.714101667 | -2.184600742 | 0.084077093 | 0.191184462 | 0.718547406 | -4.907695245 |
| RPL22 | 0.876022667 | -0.876022667 | 2.171313239 | 0.085453534 | 0.192144042 | 0.716373078 | -4.924803473 |
| PCNA | 1.273108 | -1.273108 | 2.017829085 | 0.119399516 | 0.247933421 | 0.605664927 | -4.970815426 |
| PPIC | -1.2724155 | 1.2724155 | -1.977321834 | 0.12481684 | 0.251866462 | 0.598829659 | -5.018702669 |
| HMGA1 | -0.813877 | 0.813877 | -1.964743883 | 0.126554245 | 0.251866462 | 0.598829659 | -5.033585009 |
| NAGLU | -0.7360395 | 0.7360395 | -1.949934146 | 0.12863445 | 0.253993097 | 0.595178087 | -5.051114419 |
| DYNC1H1 | 1.6230295 | -1.6230295 | 1.947346764 | 0.129001757 | 0.253993097 | 0.595178087 | -5.054177589 |
| ACLY | 1.402632 | -1.402632 | 2.048974566 | 0.099322896 | 0.219513442 | 0.658538881 | -5.082340235 |
| LRP1 | 0.620722 | -0.620722 | 2.048682746 | 0.099358716 | 0.219513442 | 0.658538881 | -5.082715813 |
| EDIL3 | 0.7038575 | -0.7038575 | 1.911812644 | 0.134165493 | 0.258796381 | 0.5870418 | -5.096261467 |
| SRPX | 0.7112605 | -0.7112605 | 1.904786883 | 0.135213278 | 0.25950023 | 0.585862252 | -5.104584911 |
| CSF1 | -0.652242 | 0.652242 | -2.005694471 | 0.104788113 | 0.227846611 | 0.642357427 | -5.138015951 |
| PLA2G15 | -0.9061045 | 0.9061045 | -1.874701818 | 0.13980314 | 0.26696077 | 0.573552554 | -5.140232235 |
| PPIB | -0.668003333 | 0.668003333 | -2.000012213 | 0.105528957 | 0.227846611 | 0.642357427 | -5.145320859 |
| XRCC6 | 1.198114 | -1.198114 | 1.859698881 | 0.142155845 | 0.267962728 | 0.57192561 | -5.158010227 |
| RPL27 | 0.557104 | -0.557104 | 1.987333631 | 0.107201992 | 0.23015117 | 0.637986813 | -5.161615172 |
| VCP | 1.129717333 | -1.129717333 | 1.938840653 | 0.113863479 | 0.241721351 | 0.616684986 | -5.223864354 |
| DPYSL2 | 1.220403 | -1.220403 | 1.79991864 | 0.151970272 | 0.280333511 | 0.552324984 | -5.228827703 |
| HSPA8 | 0.854851667 | -0.854851667 | 1.929672838 | 0.115171035 | 0.243138852 | 0.614145639 | -5.235617524 |

| | | | | | | | |
|---|---|---|---|---|---|---|---|
| THBS3 | 0.6644865 | -0.6644865 | 1.781901638 | 0.155071325 | 0.284671998 | 0.54565525 | -5.250156612 |
| TCP1 | 0.9026595 | -0.9026595 | 1.774273752 | 0.156404876 | 0.285739678 | 0.544029449 | -5.25918353 |
| PSMA5 | -0.556117 | 0.556117 | -1.902168501 | 0.119189057 | 0.247933421 | 0.605664927 | -5.270843555 |
| FAM3C | -0.865375 | 0.865375 | -1.901360616 | 0.119309272 | 0.247933421 | 0.605664927 | -5.271877409 |
| FLNC | 1.9889865 | -1.9889865 | 1.74376325 | 0.161864485 | 0.290134454 | 0.537400695 | -5.295266842 |
| PLAU | -1.589224 | 1.589224 | -1.874926627 | 0.123313312 | 0.251866462 | 0.598829659 | -5.305676041 |
| RPL30 | 0.745823667 | -0.745823667 | 1.874859051 | 0.123323726 | 0.251866462 | 0.598829659 | -5.305762368 |
| TAGLN2 | 0.521200667 | -0.521200667 | 1.874381562 | 0.123397334 | 0.251866462 | 0.598829659 | -5.306372344 |
| LTA4H | 0.958632 | -0.958632 | 1.725093426 | 0.165306609 | 0.293096758 | 0.532988985 | -5.317324821 |
| PAM | -0.510352 | 0.510352 | -1.860048776 | 0.125628234 | 0.251866462 | 0.598829659 | -5.324672398 |
| ACTR3 | 0.6337865 | -0.6337865 | 1.716934579 | 0.166835534 | 0.293096758 | 0.532988985 | -5.326958182 |
| HNRNPA2B1 | 0.512658333 | -0.512658333 | 1.855664192 | 0.12631904 | 0.251866462 | 0.598829659 | -5.330266819 |
| LAMC1 | -0.501193667 | 0.501193667 | -1.853913062 | 0.126596037 | 0.251866462 | 0.598829659 | -5.332500624 |
| CSRP1 | 0.681704 | -0.681704 | 1.707793109 | 0.168566683 | 0.293096758 | 0.532988985 | -5.3377469 |
| PARK7 | 0.775452757 | -0.775452757 | 1.821048219 | 0.131913162 | 0.256730721 | 0.59052216 | -5.374366793 |
| PRDX6 | 0.537152 | -0.537152 | 1.819417946 | 0.132182863 | 0.256730721 | 0.59052216 | -5.376440596 |
| IGFBP4 | -1.0421747 | 1.0421747 | -1.81799336 | 0.132419003 | 0.256730721 | 0.59052216 | -5.378252511 |
| CCDC80 | -0.507718667 | 0.507718667 | -1.764197683 | 0.141662018 | 0.267962728 | 0.57192561 | -5.446496924 |
| CRIM1 | 0.6205465 | -0.6205465 | 1.612629661 | 0.18777307 | 0.317127852 | 0.498765614 | -5.449667529 |
| FSTL1 | -0.497722333 | 0.497722333 | -1.759816814 | 0.142443345 | 0.267962728 | 0.57192561 | -5.452038045 |
| DDB1 | 1.076619667 | -1.076619667 | 1.75323148 | 0.143626154 | 0.26885684 | 0.57047891 | -5.460362472 |
| PLEC | 1.416594667 | -1.416594667 | 1.713403633 | 0.150996876 | 0.280333511 | 0.552324984 | -5.510572905 |
| ANXA2 | 0.856584 | -0.856584 | 1.712103782 | 0.151243829 | 0.280333511 | 0.552324984 | -5.512207491 |
| STC1 | -0.465591667 | 0.465591667 | -1.6720897 | 0.159049911 | 0.289181656 | 0.53882926 | -5.562387608 |
| GOT1 | 1.1659113 | -1.1659113 | 1.66592902 | 0.160287495 | 0.289286427 | 0.538671942 | -5.570088551 |
| PAPPA | -0.453776 | 0.453776 | -1.664232051 | 0.160630095 | 0.289286427 | 0.538671942 | -5.572208565 |
| MAMDC2 | -0.591167 | 0.591167 | -1.493086652 | 0.215229924 | 0.355597265 | 0.449041588 | -5.588769846 |
| PGK1 | 1.0192511 | -1.0192511 | 1.640622358 | 0.165473953 | 0.293096758 | 0.532988985 | -5.601647619 |
| ARPC4 | 0.516705433 | -0.516705433 | 1.634194415 | 0.166818014 | 0.293096758 | 0.532988985 | -5.609643895 |
| YWHAE | 1.0615 | -1.0615 | 1.63034813 | 0.167627497 | 0.293096758 | 0.532988985 | -5.614424665 |
| DDX39B | 1.0622745 | -1.0622745 | 1.463053197 | 0.22276357 | 0.361752805 | 0.441588093 | -5.62335665 |
| LDHB | 0.787610667 | -0.787610667 | 1.622144403 | 0.169367234 | 0.293096758 | 0.532988985 | -5.624611491 |
| TIMP2 | -0.659784 | 0.659784 | -1.620642947 | 0.169687597 | 0.293096758 | 0.532988985 | -5.626474395 |
| IGFBP7 | -0.4788801 | 0.4788801 | -1.610366733 | 0.171896564 | 0.295568752 | 0.529341482 | -5.63921174 |
| COL12A1 | -0.451337 | 0.451337 | -1.574783478 | 0.17976974 | 0.30664104 | 0.513369721 | -5.683139075 |
| GSTP1 | 0.426710333 | -0.426710333 | 1.573987636 | 0.179949873 | 0.30664104 | 0.513369721 | -5.684118254 |
| GOT2 | 0.809279667 | -0.809279667 | 1.556614413 | 0.183927126 | 0.312019232 | 0.505818637 | -5.705456355 |
| PSMB3 | -0.5129825 | 0.5129825 | -1.382959864 | 0.244202274 | 0.373520576 | 0.427685469 | -5.714682052 |
| PROCR | -0.68948461 | 0.68948461 | -1.348484498 | 0.254062578 | 0.383110236 | 0.416676244 | -5.753522686 |
| TXNDC5 | 0.7130531 | -0.7130531 | 1.326212984 | 0.260643427 | 0.386892587 | 0.412409592 | -5.778444489 |
| PGLS | -0.521222 | 0.521222 | -1.314008048 | 0.264321465 | 0.389310685 | 0.409703677 | -5.792041892 |
| P4HB | 0.551469333 | -0.551469333 | 1.479485308 | 0.202656865 | 0.34075048 | 0.467563523 | -5.799257543 |
| DSTN | 0.545596367 | -0.545596367 | 1.471736824 | 0.204638646 | 0.342566896 | 0.465254607 | -5.808590357 |
| EXT2 | 0.425230667 | -0.425230667 | 1.45898135 | 0.207942162 | 0.34657027 | 0.460208695 | -5.823915337 |
| EXT1 | 0.473507 | -0.473507 | 1.28483427 | 0.273322658 | 0.399471577 | 0.398514116 | -5.824363016 |
| YWHAB | 0.433028333 | -0.433028333 | 1.437858039 | 0.213526902 | 0.354324117 | 0.450599286 | -5.849185079 |
| PSMD2 | 1.3601425 | -1.3601425 | 1.260598852 | 0.281028982 | 0.404511414 | 0.39306922 | -5.851008408 |
| SDF4 | -0.4588745 | 0.4588745 | -1.247726242 | 0.285208165 | 0.408977746 | 0.388300323 | -5.865081607 |

| | | | | | | | |
|---|---|---|---|---|---|---|---|
| MDH1 | 0.97101 | -0.97101 | 1.42129422 | 0.218007258 | 0.358626658 | 0.445357431 | -5.868902167 |
| CLIC1 | 0.96719 | -0.96719 | 1.413104237 | 0.220255873 | 0.360663727 | 0.442897534 | -5.878618419 |
| GANAB | 0.677857823 | -0.677857823 | 1.409892259 | 0.221143812 | 0.360663727 | 0.442897534 | -5.882422928 |
| TLN1 | 0.966986 | -0.966986 | 1.390650708 | 0.226535317 | 0.364423628 | 0.438393473 | -5.905141296 |
| CD109 | 0.374674 | -0.374674 | 1.389783932 | 0.226781124 | 0.364423628 | 0.438393473 | -5.906161712 |
| EZR | 0.840514333 | -0.840514333 | 1.386403467 | 0.227742218 | 0.364423628 | 0.438393473 | -5.910138889 |
| FLNA | 0.854563 | -0.854563 | 1.384642995 | 0.228244272 | 0.364423628 | 0.438393473 | -5.912208538 |
| TGFB2 | 0.4669645 | -0.4669645 | 1.200177743 | 0.301173889 | 0.425450103 | 0.371151367 | -5.9165557 |
| EFEMP2 | -0.451054 | 0.451054 | -1.196033723 | 0.302605437 | 0.425889134 | 0.37070344 | -5.92100208 |
| LAMB1 | -0.373836667 | 0.373836667 | -1.37571566 | 0.230806465 | 0.365649165 | 0.436935414 | -5.922686937 |
| DCBLD2 | -0.388655 | 0.388655 | -1.375081425 | 0.230989532 | 0.365649165 | 0.436935414 | -5.923430292 |
| GGH | -0.381876333 | 0.381876333 | -1.370350679 | 0.232359388 | 0.365649165 | 0.436935414 | -5.928970431 |
| LNPEP | -0.439316 | 0.439316 | -1.187018073 | 0.305742418 | 0.428716306 | 0.367829998 | -5.930652557 |
| NRP1 | 0.372290667 | -0.372290667 | 1.368625752 | 0.232860784 | 0.365649165 | 0.436935414 | -5.93098848 |
| HSP90B1 | 0.7424995 | -0.7424995 | 1.364537629 | 0.234053213 | 0.366009139 | 0.43650807 | -5.935767022 |
| NPC2 | -0.932278905 | 0.932278905 | -1.356410308 | 0.236441024 | 0.368227824 | 0.433883399 | -5.94524883 |
| ACTN4 | 0.880777 | -0.880777 | 1.344875206 | 0.239869663 | 0.372042743 | 0.429407163 | -5.95866443 |
| RPS27A | 0.571015167 | -0.571015167 | 1.337486281 | 0.242090528 | 0.373520576 | 0.427685469 | -5.967231653 |
| VCL | 0.878448333 | -0.878448333 | 1.329240004 | 0.244591954 | 0.373520576 | 0.427685469 | -5.976768319 |
| GDI2 | 0.55533587 | -0.55533587 | 1.325801574 | 0.245642133 | 0.373520576 | 0.427685469 | -5.980737038 |
| PXDN | 0.368437 | -0.368437 | 1.325490924 | 0.245737221 | 0.373520576 | 0.427685469 | -5.98109537 |
| CTSD | -0.460930667 | 0.460930667 | -1.31228427 | 0.249811799 | 0.37820113 | 0.422277178 | -5.996294087 |
| HSPA13 | -0.413942 | 0.413942 | -1.111591121 | 0.333221938 | 0.457127568 | 0.339962587 | -6.010079858 |
| EEF1A1P5 | 0.7104645 | -0.7104645 | 1.105541205 | 0.335523539 | 0.458629298 | 0.338538205 | -6.016343 |
| FLNB | 0.887189767 | -0.887189767 | 1.286372849 | 0.257990273 | 0.386892587 | 0.412409592 | -6.025910062 |
| TUBA1B | 0.738235683 | -0.738235683 | 1.282183272 | 0.259335817 | 0.386892587 | 0.412409592 | -6.030672618 |
| LMNA | 0.628587133 | -0.628587133 | 1.278518617 | 0.260518111 | 0.386892587 | 0.412409592 | -6.03483241 |
| CCBE1 | -0.575041 | 0.575041 | -1.266975239 | 0.264274962 | 0.389310685 | 0.409703677 | -6.047898128 |
| TUBB | 0.858197933 | -0.858197933 | 1.242706049 | 0.272337088 | 0.399471577 | 0.398514116 | -6.075179119 |
| ACTB | 0.544560333 | -0.544560333 | 1.231726604 | 0.276058109 | 0.400910299 | 0.396952787 | -6.087434847 |
| ACTN1 | 0.756621667 | -0.756621667 | 1.23067452 | 0.276417101 | 0.400910299 | 0.396952787 | -6.088606351 |
| HSP90AB1 | 0.6988896 | -0.6988896 | 1.224459891 | 0.278546365 | 0.402462429 | 0.395274656 | -6.095516023 |
| RPL12 | -0.3683461 | 0.3683461 | -1.200586024 | 0.286865668 | 0.409808097 | 0.387419465 | -6.121892295 |
| ARPC3 | 0.39448115 | -0.39448115 | 0.988922106 | 0.382831177 | 0.498204956 | 0.302591956 | -6.133540637 |
| COL4A2 | -0.740815467 | 0.740815467 | -1.17960627 | 0.294361489 | 0.418941446 | 0.377846673 | -6.14484552 |
| MFGE8 | 0.520775 | -0.520775 | 0.964479751 | 0.393472969 | 0.508570505 | 0.293648831 | -6.157177563 |
| ENO1 | 0.586068933 | -0.586068933 | 1.166674744 | 0.299069102 | 0.424053205 | 0.37257965 | -6.158885164 |
| PRKCSH | 0.6432265 | -0.6432265 | 0.943075154 | 0.403003747 | 0.517369675 | 0.28619903 | -6.177588824 |
| ADAMTS12 | -0.348307067 | 0.348307067 | -1.129180575 | 0.313100729 | 0.437420136 | 0.359101229 | -6.199108266 |
| NID2 | 0.340482967 | -0.340482967 | 1.108027408 | 0.321271341 | 0.445839019 | 0.350821926 | -6.221472086 |
| HTRA1 | -0.3314863 | 0.3314863 | -1.107510312 | 0.321473398 | 0.445839019 | 0.350821926 | -6.22201572 |
| PSMA2 | -0.315175533 | 0.315175533 | -1.085552504 | 0.330156587 | 0.455386494 | 0.341619854 | -6.244963021 |
| PGAM1 | -0.3254206 | 0.3254206 | -1.084059349 | 0.330754401 | 0.455386494 | 0.341619854 | -6.246513602 |
| MMP11 | 0.3197325 | -0.3197325 | 0.853268191 | 0.445175003 | 0.565774251 | 0.247356822 | -6.260085316 |
| CTGF | -0.5846323 | 0.5846323 | -1.063934763 | 0.338903618 | 0.461589157 | 0.335744402 | -6.267287192 |
| PNP | 0.793836333 | -0.793836333 | 1.056282253 | 0.342047549 | 0.464207388 | 0.333287952 | -6.275124602 |
| PDGFC | 0.33304458 | -0.33304458 | 0.822816711 | 0.460282934 | 0.575317576 | 0.240092358 | -6.286823673 |
| PSMB1 | -0.285501333 | 0.285501333 | -1.030338328 | 0.352892863 | 0.477221665 | 0.321279848 | -6.301435412 |

| | | | | | | | |
|---|---|---|---|---|---|---|---|
| RELN | -0.490257933 | 0.490257933 | -1.021038824 | 0.356850921 | 0.480862943 | 0.31797869 | -6.310766803 |
| NCL | 0.654508 | -0.654508 | 1.015629604 | 0.359170428 | 0.482278313 | 0.316702267 | -6.316169956 |
| HSPG2 | -0.351574 | 0.351574 | -1.007163805 | 0.362826131 | 0.484223989 | 0.314953699 | -6.324589554 |
| PSMA3 | -0.279454767 | 0.279454767 | -1.006375808 | 0.363167991 | 0.484223989 | 0.314953699 | -6.325370958 |
| HSP90AA1 | 0.499537967 | -0.499537967 | 1.001740764 | 0.365184311 | 0.485209924 | 0.314070325 | -6.329959269 |
| PDIA3 | 0.362226767 | -0.362226767 | 0.992261571 | 0.369337133 | 0.489017807 | 0.310675327 | -6.3393003 |
| B4GAT1 | 0.30949705 | -0.30949705 | 0.760067592 | 0.492713703 | 0.594384784 | 0.225932317 | -6.339779498 |
| AXL | 0.264250333 | -0.264250333 | 0.988261796 | 0.371101227 | 0.489647452 | 0.310116501 | -6.343224484 |
| PRDX1 | -0.272520633 | 0.272520633 | -0.974895555 | 0.377047333 | 0.495466153 | 0.304986008 | -6.356262834 |
| PSMA6 | -0.261053 | 0.261053 | -0.972505386 | 0.378118906 | 0.495466153 | 0.304986008 | -6.358582054 |
| ADAMTSL1 | -0.289311315 | 0.289311315 | -0.730384725 | 0.508663192 | 0.605931076 | 0.217576774 | -6.363770899 |
| MINPP1 | 0.265281733 | -0.265281733 | 0.967044634 | 0.38057655 | 0.496972815 | 0.303667367 | -6.363866565 |
| UBE2N | -0.29273575 | 0.29273575 | -0.721647865 | 0.513431879 | 0.607800978 | 0.216238606 | -6.370697562 |
| CCT3 | 0.774705667 | -0.774705667 | 0.955248532 | 0.38593041 | 0.500524081 | 0.300575023 | -6.37521427 |
| DKK3 | -0.2787839 | 0.2787839 | -0.713376854 | 0.517977255 | 0.611277506 | 0.213761585 | -6.377197287 |
| LXN | 0.285911667 | -0.285911667 | 0.929928455 | 0.397630673 | 0.512202223 | 0.290558541 | -6.399253946 |
| COL4A1 | -0.3065745 | 0.3065745 | -0.682684899 | 0.535106209 | 0.620961346 | 0.206935433 | -6.400817606 |
| IGF2R | -0.26888447 | 0.26888447 | -0.681125118 | 0.535987688 | 0.620961346 | 0.206935433 | -6.401996699 |
| MSN | 0.3733753 | -0.3733753 | 0.877823234 | 0.422608229 | 0.540710866 | 0.267034903 | -6.447310711 |
| TPT1 | -0.245439567 | 0.245439567 | -0.863384649 | 0.429745026 | 0.547997012 | 0.261221809 | -6.460278818 |
| CHST11 | 0.216288 | -0.216288 | 0.581308887 | 0.594563356 | 0.680524324 | 0.167156348 | -6.472957664 |
| GALNT2 | -0.244294667 | 0.244294667 | -0.820749654 | 0.451366141 | 0.57028966 | 0.243904503 | -6.497652728 |
| PSMB5 | 0.221711333 | -0.221711333 | 0.820046771 | 0.451729441 | 0.57028966 | 0.243904503 | -6.498257133 |
| DKK1 | -0.379209 | 0.379209 | -0.81605275 | 0.453798053 | 0.571004173 | 0.243360718 | -6.501684223 |
| LOXL2 | -0.225636 | 0.225636 | -0.808378955 | 0.457792653 | 0.574129399 | 0.240990214 | -6.508233526 |
| PDIA6 | 0.370499833 | -0.370499833 | 0.80079203 | 0.461768054 | 0.575317576 | 0.240092358 | -6.514662824 |
| NOV | 0.265843333 | -0.265843333 | 0.788854509 | 0.468075426 | 0.579848508 | 0.236685456 | -6.524685715 |
| ACTG1 | 0.414458667 | -0.414458667 | 0.785410531 | 0.469906991 | 0.579848508 | 0.236685456 | -6.527555981 |
| PSMB4 | -0.2254172 | 0.2254172 | -0.785268809 | 0.469982475 | 0.579848508 | 0.236685456 | -6.527673888 |
| XYLT2 | 0.1809015 | -0.1809015 | 0.48904765 | 0.652345474 | 0.731242714 | 0.135938448 | -6.530159314 |
| CHRDL1 | -0.211679 | 0.211679 | -0.779833346 | 0.47288431 | 0.581540576 | 0.235419978 | -6.532183683 |
| GPI | 0.378888 | -0.378888 | 0.775192212 | 0.475372554 | 0.582714744 | 0.234543993 | -6.536015366 |
| CDH2 | -0.2332688 | 0.2332688 | -0.77190604 | 0.477140197 | 0.583000884 | 0.234330786 | -6.538717742 |
| LMAN2 | -0.18080695 | 0.18080695 | -0.465861246 | 0.667382727 | 0.743710957 | 0.12859582 | -6.543184144 |
| BMP1 | -0.22554 | 0.22554 | -0.763032932 | 0.481937203 | 0.586974799 | 0.231380544 | -6.545970118 |
| ECM1 | -0.211248133 | 0.211248133 | -0.758248783 | 0.484538223 | 0.588257267 | 0.230432699 | -6.549853373 |
| LGALS1 | -0.231152 | 0.231152 | -0.74671121 | 0.490852909 | 0.594025813 | 0.226194683 | -6.559139776 |
| MMP2 | 0.214802 | -0.214802 | 0.728736345 | 0.50080882 | 0.602238454 | 0.220231518 | -6.573383758 |
| PSMA4 | 0.368134667 | -0.368134667 | 0.717908422 | 0.506875304 | 0.605931076 | 0.217576774 | -6.58183106 |
| LTBP1 | 0.2131132 | -0.2131132 | 0.715384367 | 0.508296885 | 0.605931076 | 0.217576774 | -6.583785649 |
| PSMA1 | 0.45161 | -0.45161 | 0.708093362 | 0.512419044 | 0.607800978 | 0.216238606 | -6.589400648 |
| CALU | -0.1321118 | 0.1321118 | -0.3579873 | 0.739815073 | 0.802228594 | 0.095701862 | -6.596228098 |
| COTL1 | 0.286336667 | -0.286336667 | 0.687962712 | 0.52392166 | 0.614792936 | 0.211271131 | -6.6046622 |
| TKT | 0.258195 | -0.258195 | 0.686209149 | 0.524932038 | 0.614792936 | 0.211271131 | -6.605974679 |
| PLS3 | 0.249731267 | -0.249731267 | 0.684687724 | 0.525809748 | 0.614792936 | 0.211271131 | -6.607111197 |
| FAM20C | -0.231653333 | 0.231653333 | -0.668120015 | 0.535432816 | 0.620961346 | 0.206935433 | -6.619353456 |
| RAC1 | 0.0786285 | -0.0786285 | 0.200038073 | 0.851949447 | 0.894311574 | 0.048511149 | -6.650084604 |
| RARRES1 | -0.2304056 | 0.2304056 | -0.6234138 | 0.56198894 | 0.649105766 | 0.187684533 | -6.651141588 |

| | | | | | | | |
|---|---|---|---|---|---|---|---|
| UAP1 | 0.06963905 | -0.06963905 | 0.189258657 | 0.859804491 | 0.896090393 | 0.047648179 | -6.652680779 |
| REXO2 | -0.0639536 | 0.0639536 | -0.173718259 | 0.871164143 | 0.904487362 | 0.043597497 | -6.656175235 |
| MAN2A1 | -0.1712892 | 0.1712892 | -0.608486071 | 0.571044943 | 0.657566904 | 0.182060053 | -6.661342368 |
| ANXA5 | -0.05084285 | 0.05084285 | -0.137134565 | 0.898052821 | 0.919838469 | 0.036288432 | -6.663236565 |
| SRPX2 | 0.0131927 | -0.0131927 | 0.035731671 | 0.973338309 | 0.975906484 | 0.010591796 | -6.674164004 |
| SPARC | -0.180970667 | 0.180970667 | -0.584382799 | 0.585863089 | 0.672592066 | 0.17224826 | -6.677366522 |
| PSMB6 | -0.150569633 | 0.150569633 | -0.556370828 | 0.603382696 | 0.687290179 | 0.162859862 | -6.695284335 |
| PEBP1 | 0.155540667 | -0.155540667 | 0.555247435 | 0.604091894 | 0.687290179 | 0.162859862 | -6.695986902 |
| ALCAM | -0.191051333 | 0.191051333 | -0.543969037 | 0.611239765 | 0.693346599 | 0.15904961 | -6.702971304 |
| PSMB2 | -0.1480348 | 0.1480348 | -0.537541111 | 0.615336066 | 0.695915789 | 0.15744331 | -6.70689549 |
| VIM | 0.399143333 | -0.399143333 | 0.528576145 | 0.621076169 | 0.700323276 | 0.154701439 | -6.712299646 |
| QSOX1 | -0.1443146 | 0.1443146 | -0.520526444 | 0.626256853 | 0.704075752 | 0.152380612 | -6.71708336 |
| GPC1 | -0.131689023 | 0.131689023 | -0.477145663 | 0.654599831 | 0.731611576 | 0.135719431 | -6.741729127 |
| FUCA2 | -0.129977 | 0.129977 | -0.453902036 | 0.670071199 | 0.744523555 | 0.128121558 | -6.754134735 |
| MET | -0.133158667 | 0.133158667 | -0.441851581 | 0.678167707 | 0.751322824 | 0.124173418 | -6.760343187 |
| TAGLN | -0.186912667 | 0.186912667 | -0.425632011 | 0.689144689 | 0.761264482 | 0.118464432 | -6.768456653 |
| UBE2V1 | 0.180382333 | -0.180382333 | 0.417562481 | 0.694639182 | 0.763671806 | 0.117093243 | -6.772388706 |
| ARHGDIA | 0.160154 | -0.160154 | 0.415285291 | 0.696193644 | 0.763671806 | 0.117093243 | -6.773485688 |
| GSTO1 | 0.116460133 | -0.116460133 | 0.413588636 | 0.697352938 | 0.763671806 | 0.117093243 | -6.77429939 |
| LEPRE1 | 0.103241267 | -0.103241267 | 0.377532519 | 0.722209938 | 0.788620048 | 0.103132187 | -6.790855425 |
| LDHA | 0.261049667 | -0.261049667 | 0.36757108 | 0.729149356 | 0.793916204 | 0.100225334 | -6.795179552 |
| STC2 | 0.108913667 | -0.108913667 | 0.350649461 | 0.741005886 | 0.802228594 | 0.095701862 | -6.802274477 |
| TGM2 | 0.32082 | -0.32082 | 0.324420137 | 0.759547764 | 0.819966336 | 0.086203977 | -6.812643606 |
| SERPINB2 | 0.132035333 | -0.132035333 | 0.317785982 | 0.76426784 | 0.822724587 | 0.084745523 | -6.815144344 |
| SERPINE2 | -0.111462 | 0.111462 | -0.286236652 | 0.786872874 | 0.843525977 | 0.073901538 | -6.826357245 |
| RPL10A | 0.077570067 | -0.077570067 | 0.284629906 | 0.788030847 | 0.843525977 | 0.073901538 | -6.826898088 |
| HSPA5 | 0.084058867 | -0.084058867 | 0.277877607 | 0.792904061 | 0.846358267 | 0.072445759 | -6.829138798 |
| LAMA4 | 0.070232667 | -0.070232667 | 0.253166794 | 0.810829485 | 0.8630678 | 0.063955086 | -6.836894139 |
| CNBP | 0.0640623 | -0.0640623 | 0.23950609 | 0.82079762 | 0.869201645 | 0.060879461 | -6.840880055 |
| NME1-NME2 | 0.0734912 | -0.0734912 | 0.239001173 | 0.821166817 | 0.869201645 | 0.060879461 | -6.841023249 |
| CLSTN1 | 0.064191 | -0.064191 | 0.225601212 | 0.830984116 | 0.877149901 | 0.056926181 | -6.844715582 |
| MYDGF | 0.055066067 | -0.055066067 | 0.202045477 | 0.848327164 | 0.892975962 | 0.049160232 | -6.85070039 |
| MRC2 | 0.054345533 | -0.054345533 | 0.191611283 | 0.856041844 | 0.896090393 | 0.047648179 | -6.853144387 |
| TPI1 | -0.059173333 | 0.059173333 | -0.185298466 | 0.860718404 | 0.896090393 | 0.047648179 | -6.85456111 |
| RPL5 | -0.064970333 | 0.064970333 | -0.165557232 | 0.87538434 | 0.906392505 | 0.042683694 | -6.858689287 |
| TIMP1 | 0.043274033 | -0.043274033 | 0.156742132 | 0.881952248 | 0.909222175 | 0.041329981 | -6.860384387 |
| TGFB1 | -0.050301 | 0.050301 | -0.155467874 | 0.882902585 | 0.909222175 | 0.041329981 | -6.860621831 |
| PRDX2 | 0.046256 | -0.046256 | 0.140401948 | 0.894155436 | 0.918321799 | 0.037005106 | -6.863283664 |
| CD81 | 0.084808667 | -0.084808667 | 0.130908246 | 0.901261321 | 0.920643285 | 0.03590861 | -6.864822926 |
| GALNT10 | -0.046699833 | 0.046699833 | -0.125742054 | 0.905132669 | 0.922119073 | 0.035212995 | -6.865615618 |
| PSMA7 | 0.029525033 | -0.029525033 | 0.085392769 | 0.935463935 | 0.950471378 | 0.022060957 | -6.87071416 |
| MIF | -0.016604 | 0.016604 | -0.043565886 | 0.967038972 | 0.975906484 | 0.010591796 | -6.873947149 |
| TFPI | 0.011411667 | -0.011411667 | 0.040650813 | 0.96924293 | 0.975906484 | 0.010591796 | -6.874094383 |
| FN1 | -0.0105768 | 0.0105768 | -0.039046289 | 0.970456175 | 0.975906484 | 0.010591796 | -6.874171078 |
| CALR | -0.011427667 | 0.011427667 | -0.037712203 | 0.971465003 | 0.975906484 | 0.010591796 | -6.874232497 |
| LTBP3 | -0.005345433 | 0.005345433 | -0.019725561 | 0.985071507 | 0.985071507 | 0.006532242 | -6.874852209 |
