## Supplementary figures and images for "OPTN translocates to an ATG9A-positive compartment to regulate innate immune signalling and cytokine secretion"

### Supplemental Figure 1

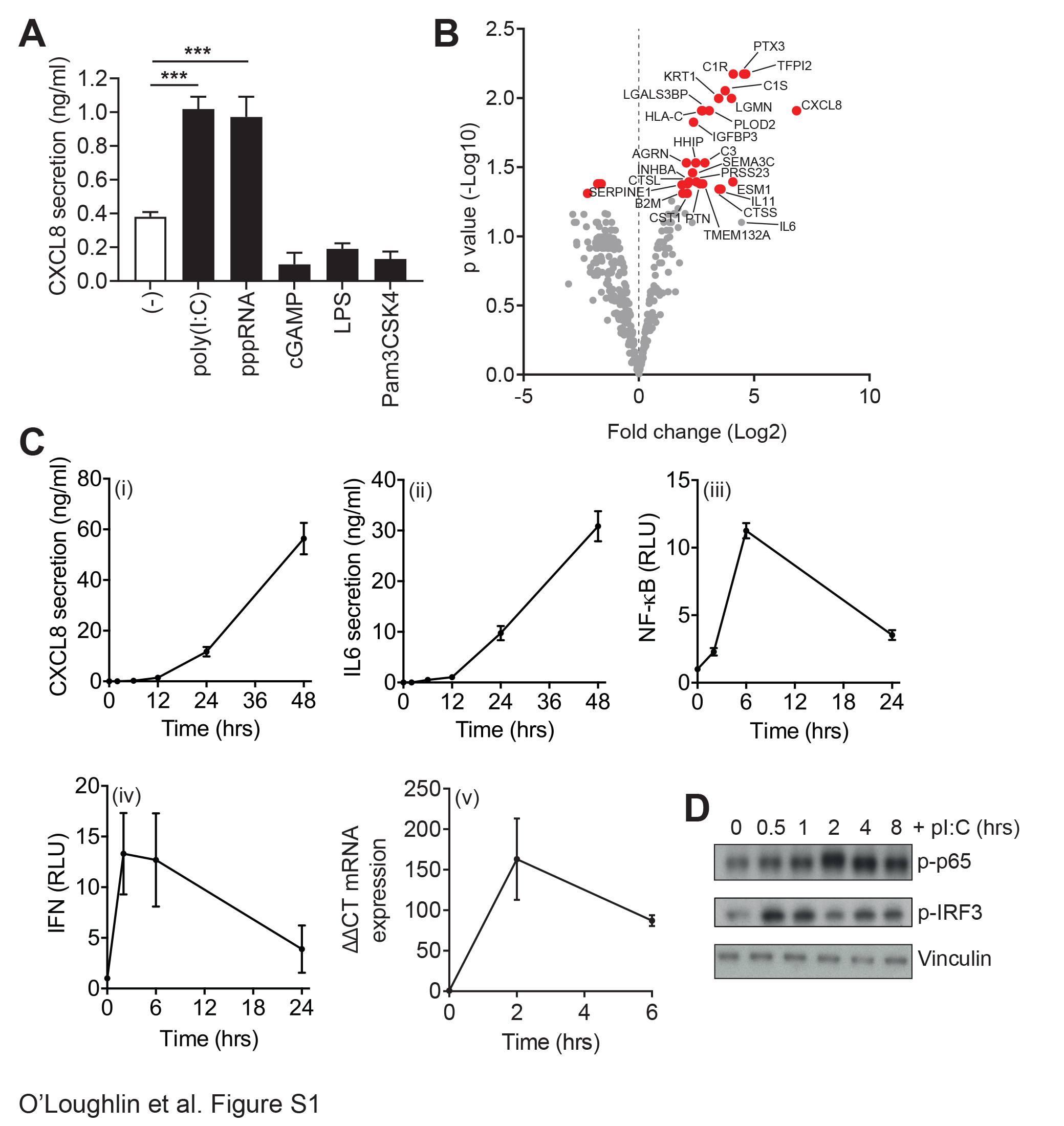

### Supplemental Figure 2

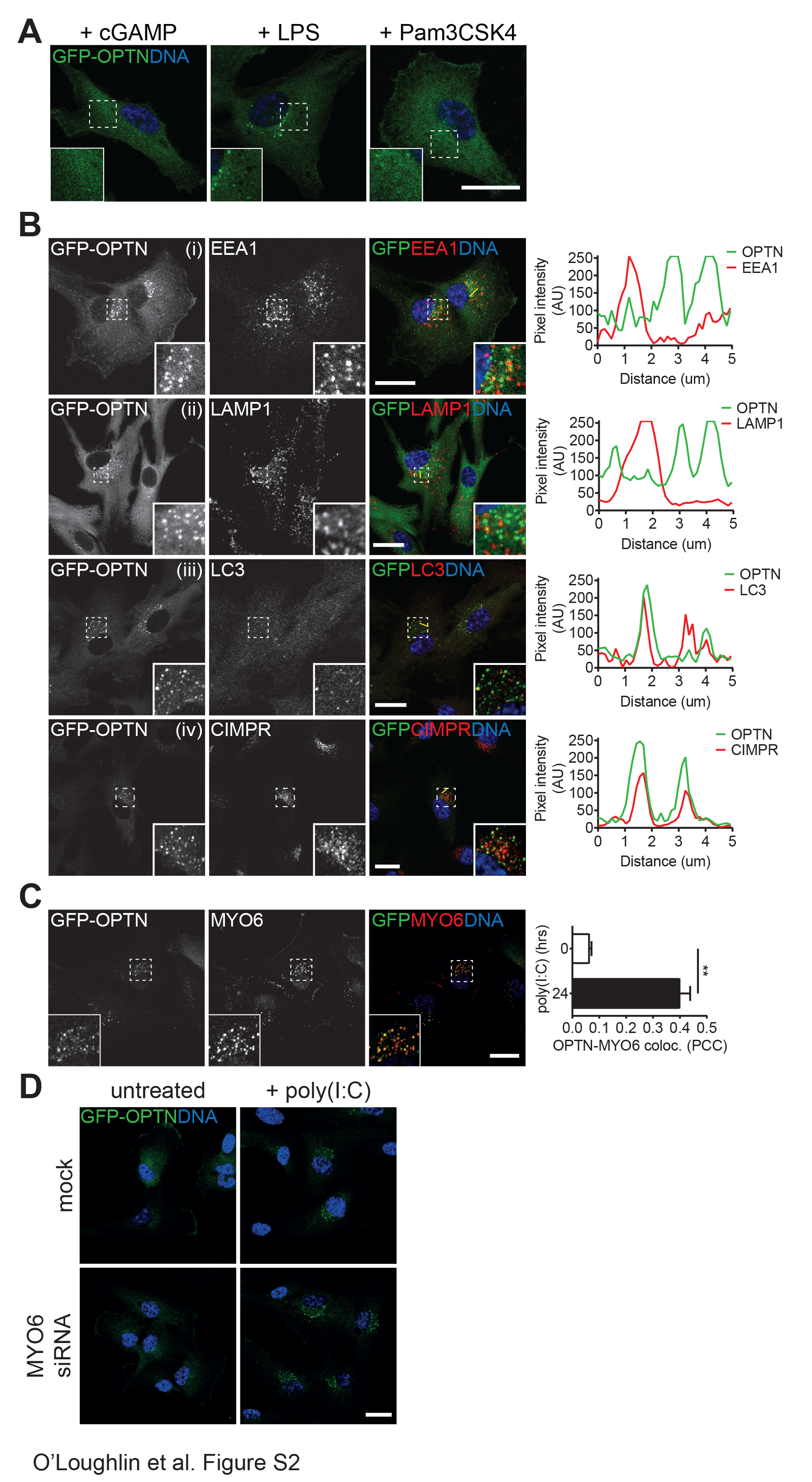

### Supplemental Figure 3

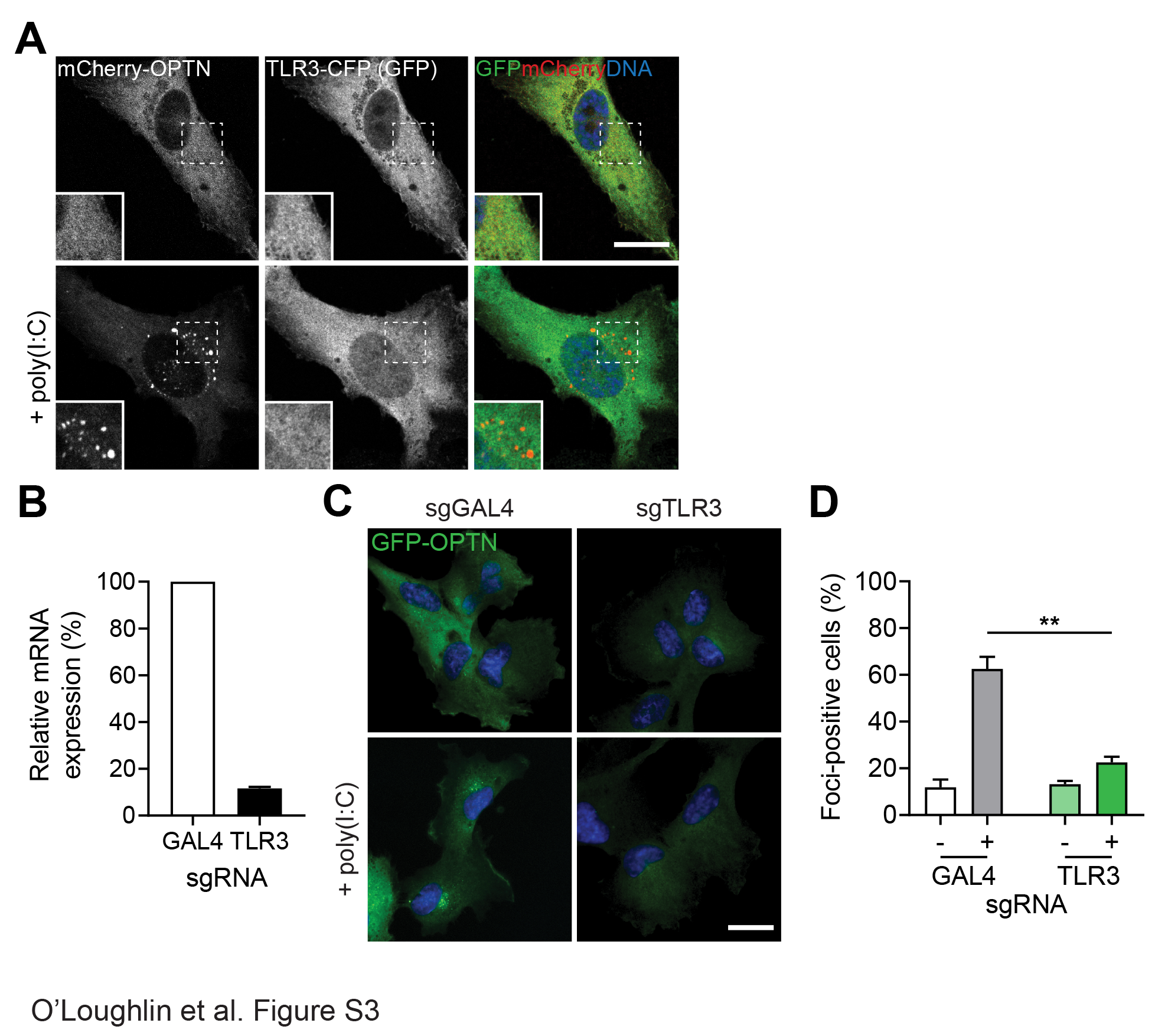

### Supplemental Figure 4

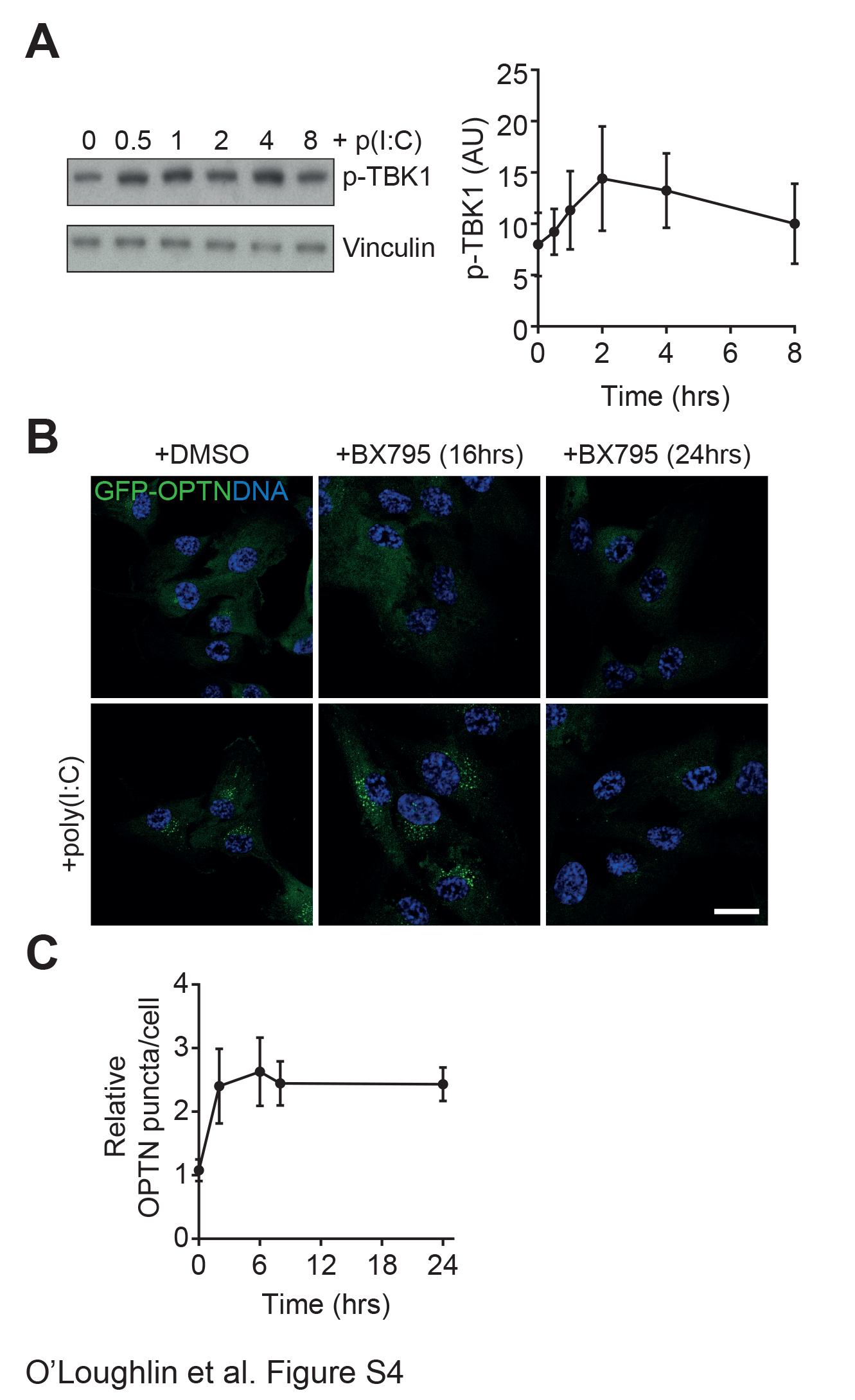

### Supplemental Figure 5

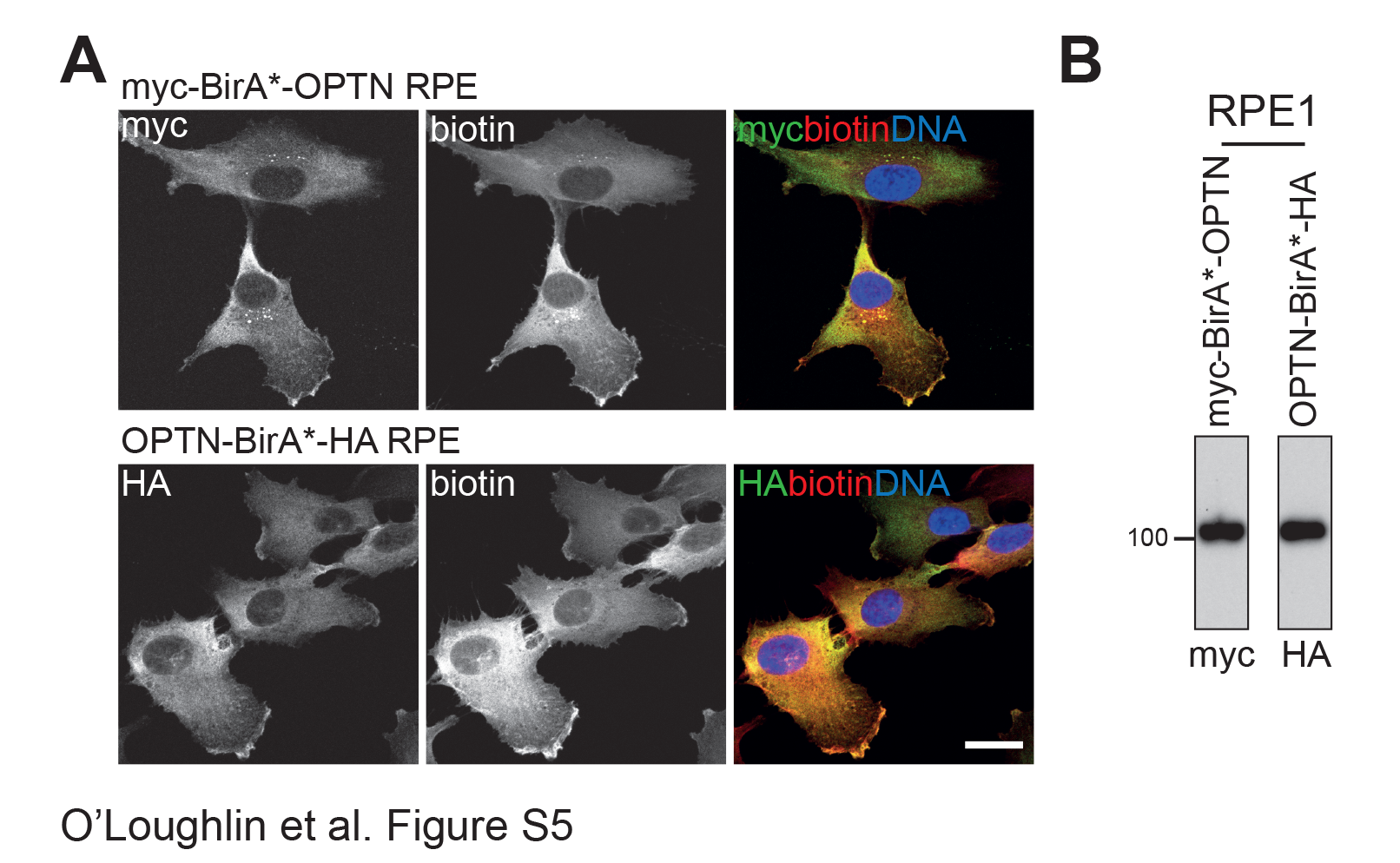

### Supplemental Figure 6

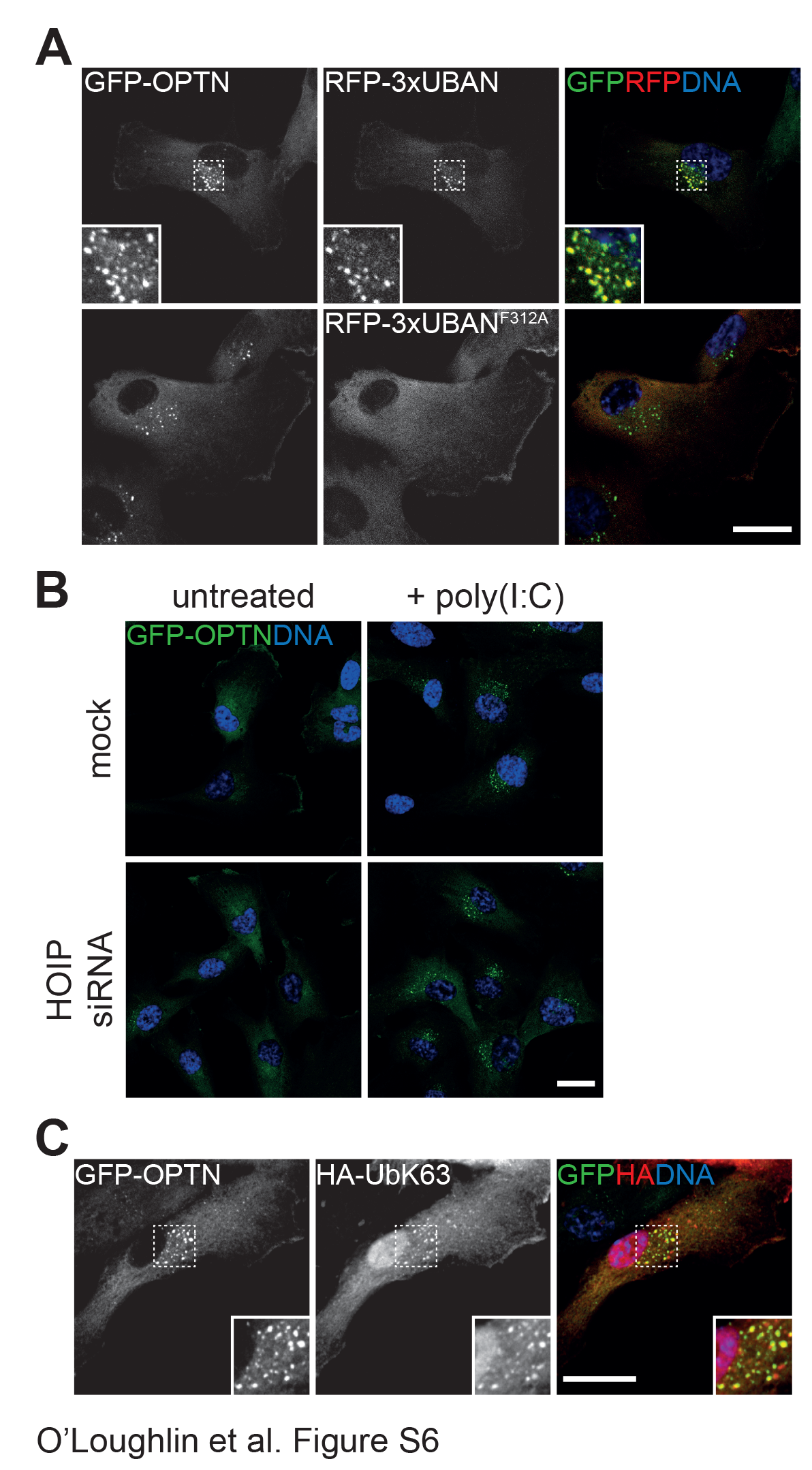

### Supplemental Figure 7

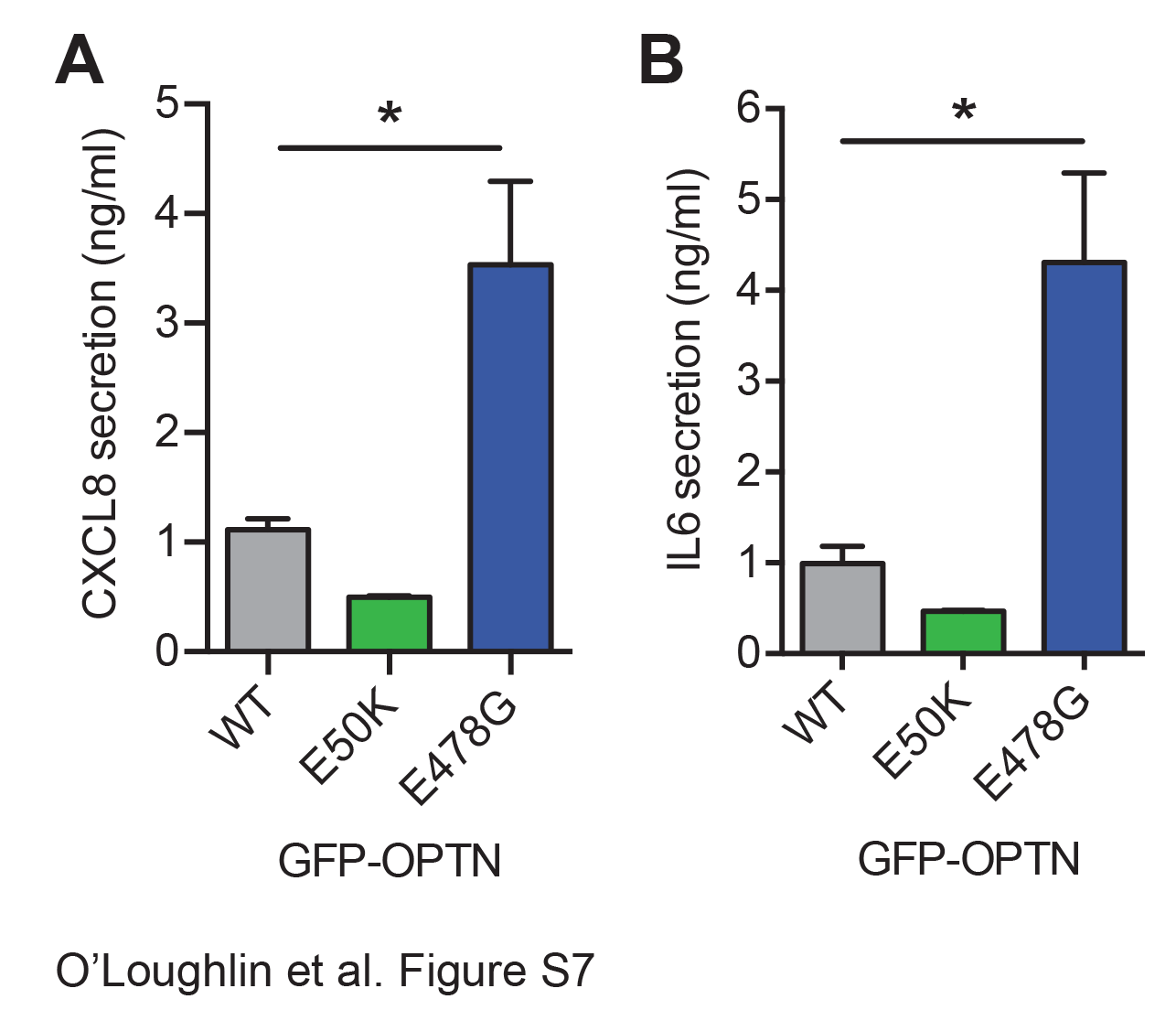
